# A spatial mRNA profiling workflow using Rapid Amplified Multiplex FISH (RAMFISH)

**DOI:** 10.1101/2024.12.06.627193

**Authors:** Tirtha Das Banerjee, Joshua Raine, Jeriel Lee Cheng Hock, Kok Hao Chen, Ajay S. Mathuru, Antónia Monteiro

**Affiliations:** Department of Biological Sciences, National University of Singapore 117557, Singapore; Department of Physiology, Yong Loo Lin School of Medicine, National University of Singapore, Singapore; Genome Institute of Singapore, Agency for Science, Technology and Research (A*STAR), Singapore; N.1 Institute for Health, National University of Singapore, Singapore; Institute for Digital Medicine (WisDM), Yong Loo Lin School of Medicine, National University of Singapore, Singapore; Healthy Longevity Translational Research Program, Yong Loo Lin School of Medicine, National University of Singapore, Singapore

**Keywords:** Multiplexed-FISH, HCR3.0, *Bicyclus anynana*, zebrafish, RAMFISH

## Abstract

Spatial localization of multiple mRNAs in intact tissues provides vital insights into development and function, yet routine multiplexed imaging remains constrained by complex workflows, specialized equipment costs, or sample preparation requirements. Here, we present Rapid Amplified Multiplexed FISH (RAMFISH), an accessible, modular, benchtop workflow for targeted spatial mRNA profiling in intact tissues and whole-mount specimens. RAMFISH achieves reliable multiplexing of over 30 transcripts through iterative cycles of standard hybridization either manually or via open-source automation and imaging in shared confocal systems. To streamline analysis, we introduce an integrated, open-source pipeline automating signal normalization, rigid and non-rigid alignment, Laplacian of Gaussian spot calling, and 30-channel composite merging. We validate the platform by mapping multiple transcripts in developing *Bicyclus anynana* butterfly wings and intact 14 days-post-fertilization *Danio rerio* larvae, resolving both historically inferred domains and newly characterized gene expression. Overall, RAMFISH methodology offers an end-to-end, completely open-source ecosystem for robust, semi-quantitative, spatial mRNA mapping on a variety of sample types.

**Graphical Abstract:** 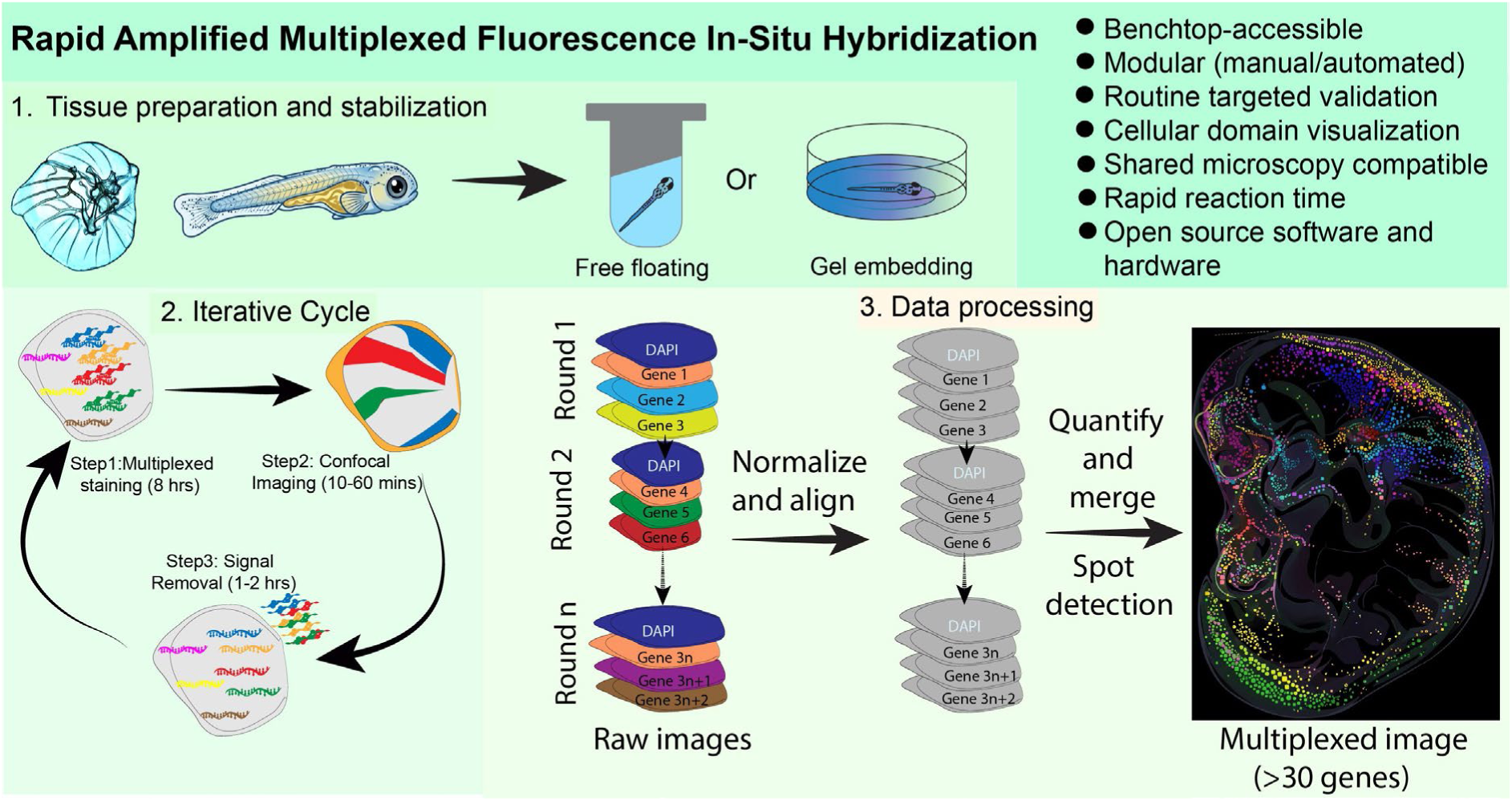

## Introduction

Multicellular organisms are organized through dynamic spatial and temporal patterns of gene expression. Visualizing the localization of RNA molecules within intact tissues provides critical insight into developmental processes, tissue organization, disease progression, and the structure of gene-regulatory networks. As a result, there is growing interest in methodologies that enable the simultaneous detection of multiple transcripts while preserving spatial context (Chen et al., 2015; Marx, 2021; Moffitt et al., 2022; Ståhl et al., 2016; Tian et al., 2023). Such approaches are particularly valuable for studying tissue differentiation, mapping regulatory interactions, and linking gene expression patterns to morphological outcomes.

To support these applications, RNA imaging workflows must provide reliable signal detection while remaining practical to implement across diverse sample types. Methods that minimize extensive sample preparation and can be applied to intact tissues, organs, or whole organisms are especially important for extending spatial transcript analysis to a broader range of biological systems.

Recent technological advances now enable spatially resolved transcriptomic profiling at single-cell or subcellular resolution while preserving tissue architecture (Moffitt et al., 2022; Moses and Pachter, 2022; Tian et al., 2023). These approaches broadly fall into two categories: sequencing-based methods such as Visium (Ståhl et al., 2016), Slide-seq (Rodriques et al., 2019), and Stereo-seq (Chen et al., 2022); and imaging-based methods that use fluorescently labelled probes for transcript detection such as MERFISH (Chen et al., 2015), seq-FISH (Lubeck and Cai, 2012), osmFISH (Codeluppi et al., 2018), STARmap (Wang et al., 2018), cycleHCR (Gandin et al., 2024), EASI-FISH (Wang et al., 2021), PRISM (Chang et al., 2024), and FISH&CHIPS (Zhou et al., 2023). While these specialized technologies have vastly expanded the horizons of spatial biology, their implementation is often tailored for large-scale profiling initiatives. For standard laboratories requiring the routine, targeted validation of specific candidate gene groups, these workflows can introduce unnecessary experimental overhead.

Rather than attempting to match the massive screening scales or single-molecule decoding complexity of these specialized platforms, our study introduces a modular, benchtop, alternative focused on routine lab work. We apply a streamlined Rapid Amplified Multiplex FISH (RAMFISH) workflow optimized specifically for semi-quantitative localization of mRNA transcripts within their native spatial domains. This is done through easy to implement, iterative cycles of hybridization via Hybridization Chain Reaction (HCR; (Choi et al., 2018)), confocal imaging, and enzymatic signal removal. We provide a straightforward software for signal normalization, image alignment at the domain level across the multiple rounds, Laplacian of Gaussian (LoG) based spot calling, and a merging tool to create the multiplexed-FISH images from up to 30 channels. By prioritizing robust tissue and cellular level domain mapping over precise subcellular, single-molecule localization, RAMFISH eliminates the need for complex probe barcoding schemes, pixel-level alignment, or dedicated proprietary instrumentation, operating entirely on a standard molecular biology lab space with access to a shared confocal microscope.

Here, we demonstrate the practical utility of RAMFISH by localizing the expression of known and novel genes across multiple developmental stages of *Bicyclus anynana* butterfly wings, and intact 14-day-post-fertilization *Danio rerio* larvae. These datasets reveal highly reproducible, distinct spatial territories for both well-characterized and previously unexamined genes, demonstrating the value of a targeted, mid-plex tool to address biological questions of gene expression relationships in intact tissues. Our results illustrate how RAMFISH provides a robust and easily adoptable framework for integrating spatial gene expression mapping into day-to-day research.

## Results

### Workflow Development Overview

The RAMFISH workflow consists of tissue collection, fixation, permeabilization, and iterative cycles of probe hybridization, imaging, and signal removal (**Figure 1A**). Depending on experimental requirements and available infrastructure, staining can be performed either manually or using an optional open source automated fluidic system (Banerjee et al., 2026). Both approaches support repeated hybridization cycles for multiplex transcript detection within the same sample.

**Figure 1:**
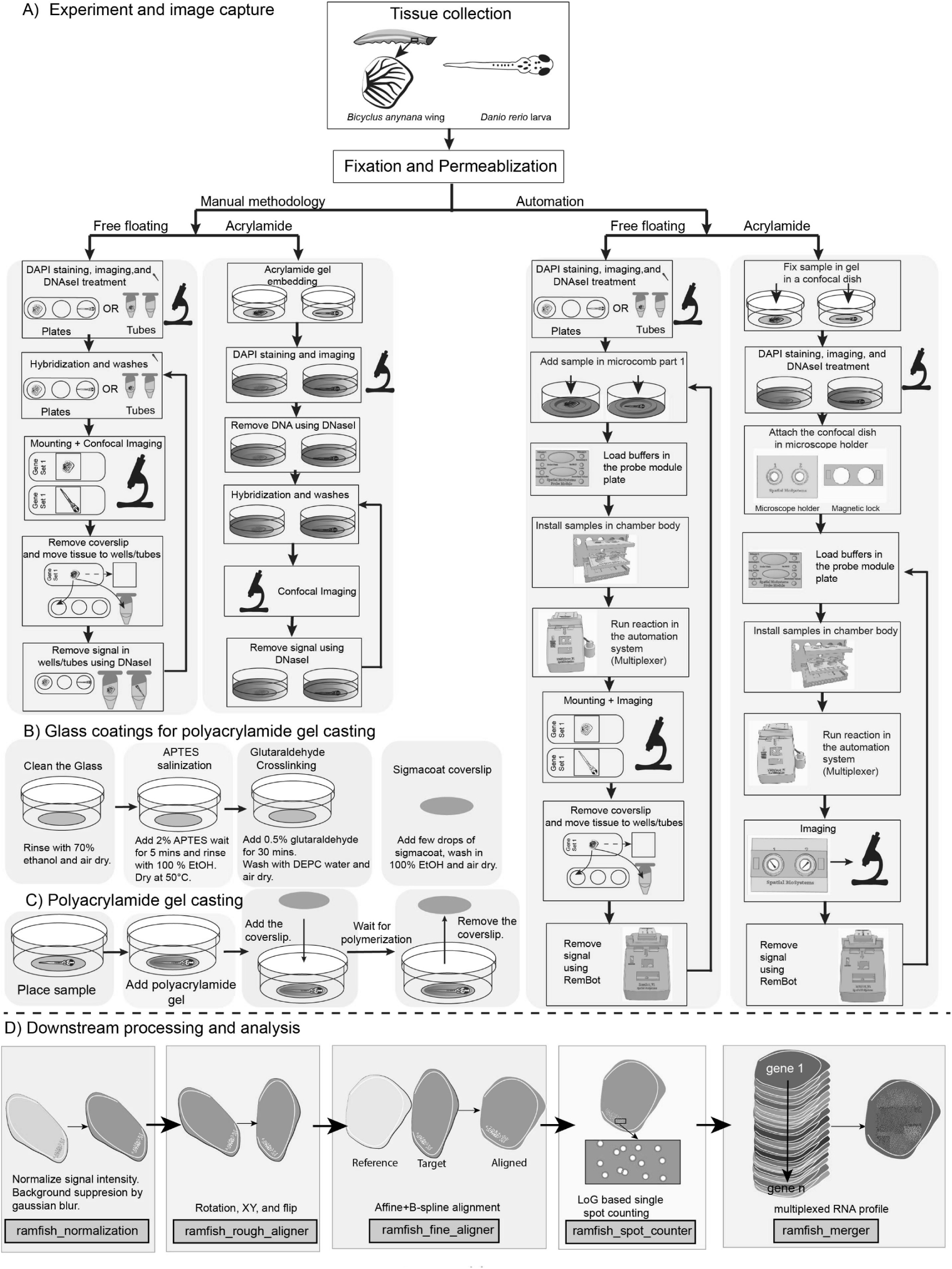
Illustration of the manual and the automation-based multiplexing protocol. **(A)** Tissues are dissected, fixed, and permeabilized. Afterwards, multiplexing can be performed either manually or using an automation system. (**B**) Simplified method of glass coating and (**C**) method for polyacrylamide gel casting. (**D**) Downstream processing and analysis on the acquired images.

### A. Manual workflow

In the manual implementation, samples are processed using standard labwares and buffers. Two configurations are supported (**Figure 1A**).

A free-floating configuration is used where tissues are maintained in solution throughout hybridization and washing steps. This approach is well suited for 2D image registration and is compatible with multi-cycle experiments when combined with domain level gene expression alignment using the RAMFISH Software Suite (described below).

For multi-cycle experiments requiring 3D image registration, samples can be immobilized by embedding them in an acrylamide-based gel within a glass-bottom imaging dish. All hybridization and washing steps are performed directly in the dish, and imaging is carried out between cycles. Immobilization improves spatial stability and facilitates alignment of gene expression patterns across sequential imaging rounds.

### B. Automated workflow

For automated, hands-free experimentation, RAMFISH can be performed using an open-source microcontroller-based fluidic system that enables programmable reagent exchange and temperature control (**Figure 1A**). The automation platform supports both free-floating and gel-embedded sample configurations. In this system, hybridization, washing, and signal removal steps are carried out under controlled fluidic conditions, after which samples are imaged and returned to the system for subsequent cycles. This configuration enables consistent execution of multi-cycle experiments while reducing manual intervention.

### Visualizing multiple transcripts in developing butterfly larval wings

We first implemented the RAMFISH workflow to investigate spatial expression patterns of multiple genes in developing larval wings of *Bicyclus anynana*. A major limitation of traditional mRNA detection protocols in butterfly evolutionary and developmental biology is their restriction to detecting only one to three genes per experiment. This constraint significantly limits the amount of information that can be extracted within a specific developmental context and hampers efforts to dissect gene regulatory networks, particularly in experiments combining CRISPR-mediated gene knockout with downstream staining analyses (Hanly et al., 2023; Matsuoka and Monteiro, 2022; Murugeshan et al., 2022).

Using RAMFISH, we successfully detected the expression of 21 previously characterized genes and 12 novel genes in *B. anynana* across multiple developmental stages ranging from 0.75 to 3.25 (wing staging as described in (Banerjee and Monteiro, 2020; Reed et al., 2007). This enabled us to systematically explore gene expression dynamics over 10-14 iterative cycles of hybridization and washing (**Figure 2 and 3; Figure S1 to S9**), demonstrating the robustness and multiplexing capacity of the workflow.

**Figure 2:**
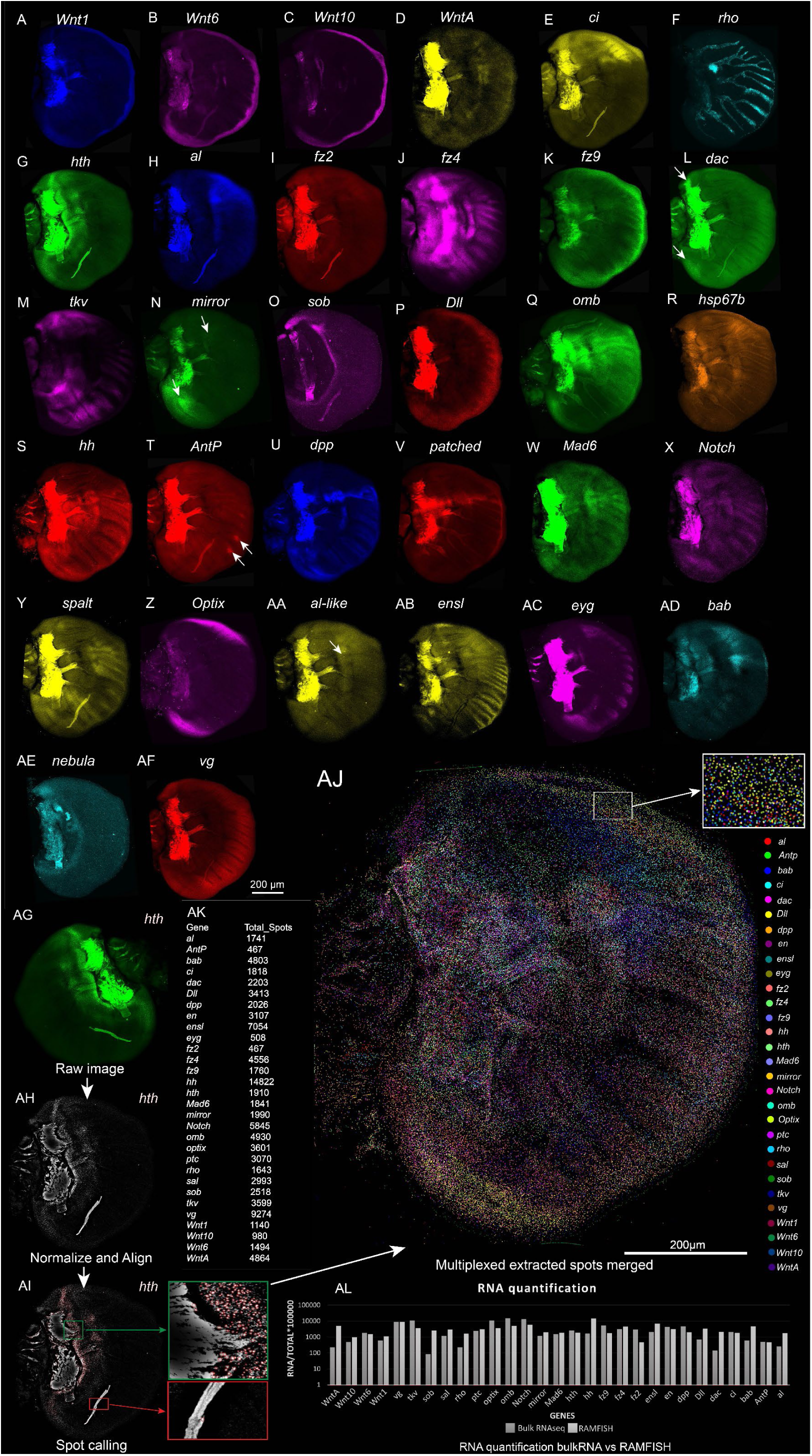
Detection of 32 genes (mRNAs) in a single early larval <u>forewing</u> of *Bicyclus anynana* (stage 1.00) using the manual, free-floating method. (**A-AF**) Raw fluorescence images of multiple target gene transcripts (white arrows indicate small, localized expression domains). Scale bar applies to all panels in this sequence. (**AG-AJ**) Step-by-step computational processing workflow of the RAMFISH pipeline illustrated using the *homothorax* (*hth*) channel: (**AG**) Raw input grayscale image showing background noise and structural tissue features. (**AH**) Normalized and aligned *hth* expression profile following background subtraction and spatial registration. (**AI**) Transcript spot calling overlay; green and red insets show high-magnification views where verified spots are encircled by red validation rings, while non-specific autofluorescence along the tracheal tissue is cleanly minimized and bypassed by the Laplacian of Gaussian (LoG) algorithm. (**AJ**) High-dimensional composite map generated by merging the extracted synthetic spots from 30 multiplexed transcript channels against a grayscale DAPI structural backdrop. Inset highlights a magnified view of tight, high-density combinatorial transcript mapping. (**AK**) Quantitative data table detailing the rough total spot counts computed across the tissue for each evaluated gene channel. (**AL**) Validation chart showing correlation analysis between bulk RNA-seq data and RAMFISH single-wing transcript counts. To account for technological variation, we used normalized gene count indicated as (RNA/Total)*100000 calculated by RNA count for specific gene by total gene count of the group multiplied by 100000 (Relative Abundance Scaling). Note: The tracheal tissue tracking along the developing wing veins and within the proximal base domain exhibits strong, non-specific autofluorescence. Gene labels for panel AJ are added separately with the same color code as spots to increase visibility.

**Figure 3.**
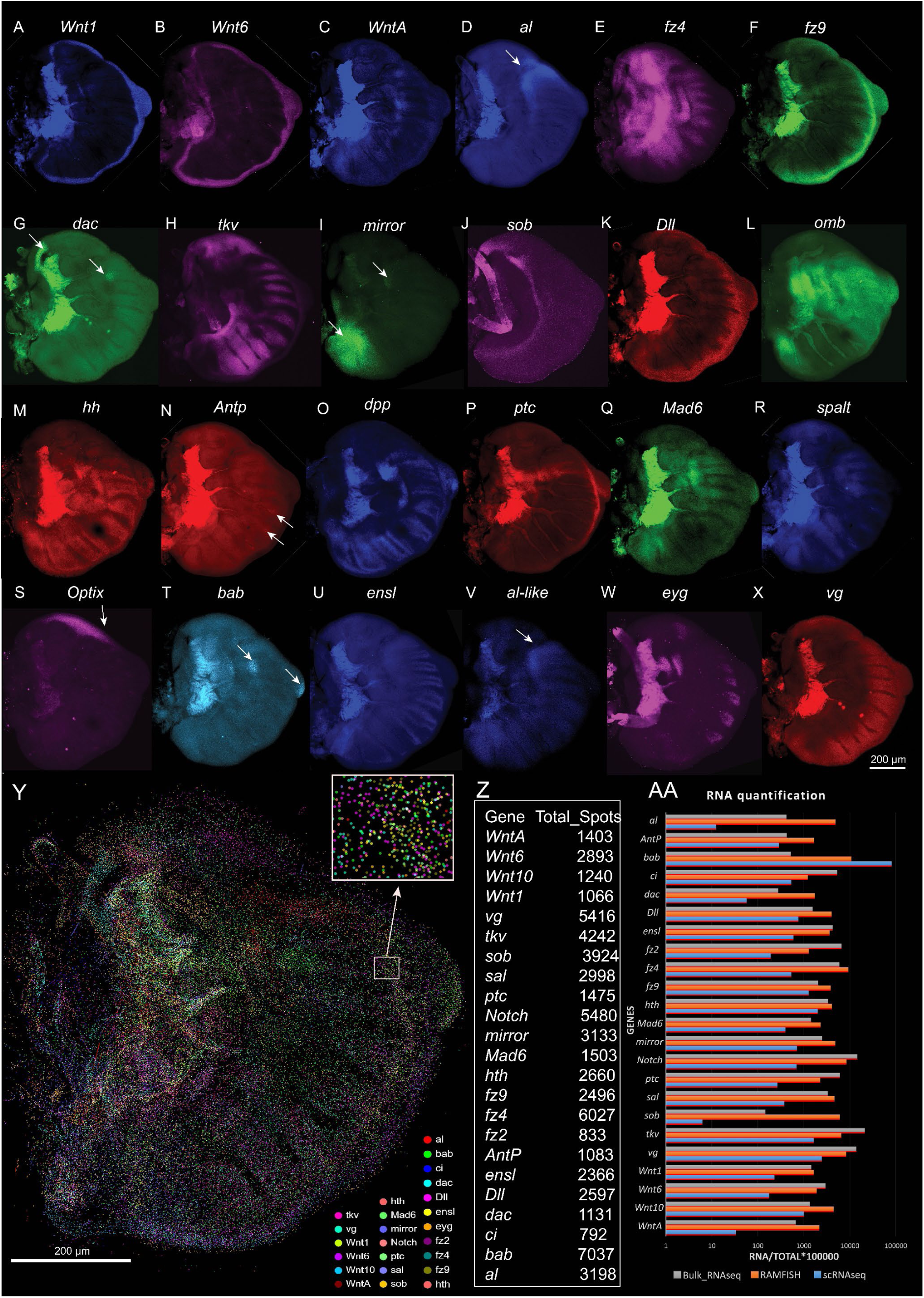
Expression of 24 genes (mRNAs) in a single larval hindwing of *Bicyclus anynana* (stage 1.00) using the manual, free-floating method. (**A-X**) Raw fluorescence images of multiple target gene transcripts (white arrows indicate small, localized expression domains). Scale bar applies to all panels in this sequence. (**Y**) High-dimensional composite map generated by merging the extracted synthetic spots from the multiplexed transcript channels against a grayscale DAPI structural backdrop. Individual target gene spot distributions are color-coded according to the adjacent matrix key. (**Z**) Quantitative data table detailing the total spot counts computed across the hindwing tissue for each evaluated gene channel using the RAMFISH pipeline. (**AA**) RNA quantification validation chart illustrating a comparative expression analysis between bulk RNAseq data and single-cell RNAseq data, and RAMFISH transcript counts. To account for technological variation, we used normalized gene count for the three technical modalities bulk RNA-seq, single-cell RNA-seq and RAMFISH indicated as RNA/Total*100000 calculated by RNA count for specific gene by total gene count of the group (here 23), multiplied by 100000 (Relative Abundance Scaling). Note: The tracheal tissue tracking along the developing hindwing veins and within the proximal base domain exhibits strong, non-specific autofluorescence. Gene labels are increased in size in panel Y to increase visibility.

Many of these genes play important functions in the development of distinct color patterns observed in the adults. During the larval stages, gene expression patterns are highly complex and dynamic (Banerjee et al., 2023; Connahs et al., 2019; Hanly et al., 2023; Mazo-vargas et al., 2017; Reed et al., 2011; Zhang et al., 2017). The known genes were *Wnt1, Wnt6, Wnt10, WntA, cubitus interruptus (ci), frizzled2 (fz2), frizzled4 (fz4), frizzled9/frizzled3 (fz9), thickvein (tkv), Distal-less (Dll), optomotor-blind (omb), hedgehog (hh), engrailed* (*en*), *Antennapedia* (*Antp*), *decapentaplegic (dpp), patched (ptc), Mothers against dpp 6 (Mad6), Notch, spalt, Optix,* and *vestigial* (*vg*) (**Figure 2 and 3; Figure S1 to S9; Video S1**). The 12 new genes for *B. anynana* were *rhomboid (rho,* at the mRNA level), *homothorax (hth), aristaless (al/al1), dachshund (dac), mirror, sister of odd and bowel (sob), heat shock promoter 67b (hsp67b), aristaless-like (al-like /al2), enhancer of split mbeta (ensl), eyegone (eyg), bric-a-bac (bab),* and *nebula.* **Table S1** contains a description of the expression domains of the 21 known genes, whether it is consistent with previous findings in other butterfly species, as well as their function, if known. The coding sequences and the probes used for the detection of the genes are provided in the **Supplementary file S2**.

Below, we provide the description of the novel genes described for the first time in *B. anynana* butterflies.

1) *rhomboid (rho)* mRNA expression was visualized in the veins of butterflies, consistent with previous protein expression results from butterflies (Banerjee and Monteiro, 2020a) and mRNA results from *Drosophila* (Guichard et al., 1999) (**Figure 2F; Figure S1, S2, S4, S5**).
2) *homothorax (hth)* expression was newly visualized along two bands along the proximal domain of the developing larval wings (**Figure 2G**, **Figure 3F, Figure S1 to S5**). A previous study has looked at the expression of *hth* in the pupal wing (Hanly et al., 2019).
3) *dachshund (dac)* was newly visualized in three distinct domains on the anterior compartment of the hindwing (stage 1.00 and 2.50) (**Figure 2G**, **Figure 3J, Figure S3, Figure S5, Figure S6)** and in an anterior proximal domain, with slightly higher expression in the lower posterior compartment on the forewing during wing development (**Figure 2L, Figure S2, Figure S4**).
4) *mirror* was expressed in the lower posterior compartment of the developing larval wings, consistent with a previous study (Chatterjee et al., 2024). Slightly higher expression of *mirror* was also observed in the proximal domain in the anterior compartment of the developing wings (**Figure 2N, Figure S1 to S6**).
5) *sister of odd and bowel (sob)* was newly visualized in a strong proximal band as well as at lower levels in a broad distal band (**Figure 2O, Figure S1 to S6**).
6) *heat shock promoter 67b (hsp67b)* was newly visualized in complex and dynamic patterns in larval wings. Major expression domains were observed along the anterior compartment and in the wing margin (**Figure 2R, Figure S1 to S6**).
7) *enhancer of split mbeta (ensl)* was newly visualized in provein cells adjacent to the veins, from early to late stages of wing development (**Figure 2AB, Figure S1 to S6**).
8) *eyegone (eyg)* was newly visualized along the wing margin in intervein cells (**Figure 2AC, Figure S1 to S6**).
9) *bric-a-bac (bab)* was newly visualized in larval wings, expressed in distinct domains in the anterior compartment (**Figure 2AD**, **Figure 3V, Figure S1 to S6**). A previous study has looked at the expression of *bab* in pupal *Colias eurytheme* butterflies (Ficarrotta et al., 2022).
10) *nebula* (uncharacterized gene: XP_023941093.2) was a newly visualized gene expressed strongly along a proximal band, and at lower levels throughout the wing tissue (**Figure 2AE, Figure S1 to S5**).
11) *aristaless (al/al1)* was expressed in the anterior compartment along a band running the proximal AP domain of the larval wings, consistent with previous protein and mRNA expression patterns in different butterfly species (Martin and Reed, 2010) (**Figure 2H, Figure S1 to S6**).
12) *aristaless-like (al-like /al2)* was expressed in a band along the proximal axis consistent with a previous study in butterflies (Martin and Reed, 2010) (**Figure 2AA, Figure S1 to S6**).

### Automated image processing and multiplexing spot merging in butterfly larval wings

To transition from qualitative imaging to semi-quantitative spatial profiling, we applied our custom Python-based computational pipeline to the multi-cycle datasets. As demonstrated in the larval forewing (**Figure 2AG-AJ**), raw confocal images from individual hybridization cycles were first subjected to automated signal normalization and rigid and non-rigid B-spline alignment to correct for major and minor tissue deformations. Following image registration, a Laplacian of Gaussian (LoG) filter was applied for automated spot extraction. This process identifies discrete fluorescent signals corresponding to individual transcripts or mRNA aggregates localized within diffraction-limited domains across the tissue architecture. This algorithm successfully extracted localized transcript coordinates while preserving the integrity of macro-structures; for instance, the algorithm accurately mapped the pro-vein and vein domains marked by *ensl* and *rho* as continuous domains. The extracted spatial coordinates from all individual cycles were then computationally integrated, generating a single, high-plex spatial map resolving distinct transcripts simultaneously across the entire wing (**Figure 2AJ**, **Figure 3Y, Figure S7**).

### Benchmarking RAMFISH Quantification Against Sequencing Modalities

To evaluate the semi-quantitative nature of our LoG-based spot extraction, we aggregated the total called spots for each mapped gene (**Figure 2AK**, **Figure 3Z**) and compared these spatial profiles to orthogonal bulk (Matsuoka et al., 2023) and single-cell RNA-seq (scRNA-seq) datasets (**Figure 2AL**, **Figure 3AA**). The single-cell dataset was generated by dissociating *B. anynana* larval hindwing wings into a single-cell suspension, followed by library preparation and sequencing using the 10x Genomics platform (10x Single Cell 3’ Gene Expression library; 8000 cells were analysed). To account for inherent technological variations in absolute counting between RAMFISH quantification and standard sequencing modalities, transcript counts were normalized using Relative Abundance Scaling. For both the RAMFISH extracted spot counts and the orthogonal RNA-seq datasets (bulk RNA-seq and single-cell RNA-seq), the raw count for each specific gene was divided by the total count of all genes probed within the multiplexed panel and then multiplied by a scaling factor of 100000. This transformation converts absolute counts into a directly comparable proportional metric across platforms.

As expected, absolute transcript counts naturally diverge across these distinct modalities due to inherent technical differences: bulk sequencing homogenizes the entire tissue volume, single-cell platforms experience transcript capture dropouts during dissociation, and spatial imaging faces optical crowding where dense fluorescent signals in highly expressing domains can be undercounted by spatial filters. Despite these cross-platform variances, the overall relative abundance trends remain largely preserved. Highly expressed genes yielded proportionally higher spatial spot counts, whereas known lowly expressed genes maintained lower detection frequencies. This multi-modal comparison confirms that RAMFISH paired with LoG spot calling provides a reliable, semi-quantitative representation of relative endogenous gene expression within the native 3D tissue space.

Crucially, this multi-modal comparison highlights a unique robustness inherent to the RAMFISH methodology. While bulk and single-cell sequencing provide deep transcriptomic profiling, they fundamentally uncouple transcript counts from their morphological context. RAMFISH bridges this gap by pairing robust, semi-quantitative spot extraction with direct visual inspection of the native 3D tissue space. For users, this dual readout is highly advantageous: the automated quantification provides a reliable measure of relative expression, while the raw optical data visually confirms that transcripts map accurately to specific developmental domains.

Note that in the case of well-known highly autofluorescent tissues such as the tracheal tissues at the proximal domain of the larval wings (Banerjee et al., 2023; Hanly et al., 2023; Matsuoka and Monteiro, 2022), a mask can be applied at the domain followed by spot counting to fully avoid non-specific spot counting (**Figure S7**).

### Multiplexing to visualize multiple transcripts in zebrafish larvae

*Z*ebrafish are commonly used vertebrate laboratory models for fundamental studies including neurobiology (Lieschke and Currie, 2007). Larval zebrafish, particularly pigment free mutants (Antinucci and Hindges, 2016; Lister James A et al., 1999), are ideal for high-throughput spatial gene expression techniques because they are abundant, transparent, require no special preparation before the procedure, and are small enough to process dozens of animals in parallel. Spatial transcriptomic profiles of the brain add unique insights into circuit-level connections between brain regions that bulk and single-cell RNA sequencing alone cannot reveal. This was achieved by registering *in situ* results from individual stainings of young larvae (< 6 dpf) to a standard reference brain (Schulze et al., 2023; Shainer et al., 2023). Maps of gene expression in late-stage larvae or juvenile fish brains, however, when complex behaviors including learning and social behaviors emerge (Gemmer et al., 2022), are limited.

We spatially profiled 9-10 genes using both the manual and the automation system on 14 days post fertilization (dpf) larvae to examine gene expression in the brain (**Figure 4; Figure S10-S14; Video 1 and 2**). We examined a number of neurotransmitter and neuromodulator associated genes such as *tph2 (*serotonin*), chatb* (acetyl choline)*, oxt* (oxytocin), and *gad2* (GABA) whose expression profile is known, as well as of receptor associated genes such as *nrp1a* (semaphorin & VEGF) and *kctd12.2* (auxiliary subunit of GABA_B_ receptor), and neurodevelopmental markers such as *omp* (olfactory marker bulb), *neurod1,* and *HuC* (*elavl3* homolog, neuronal differentiation). Overall, the expression patterns observed are consistent with previous reports available at mapzebrain (Shainer et al., 2023), or Zebrafish Information Network (ZFIN; **Table S2**) for younger larvae. For example, as seen in the mapZebrain atlas for *tph2,* the HCR technique revealed a broader mRNA expression profile compared to promoter-GAL4 fusion transgenic lines (**Figure S15**). The 10 profiled genes were: *glutamate decarboxylase 2 (gad2*), *neuropilin 1a* (*nrp1a*), *neuronal differentiation 1(neurod1), olfactory marker protein b (omp*), *tryptophan hydroxylase 2* (*tph2), oxytocin (oxt*), *potassium channel tetramerisation domain containing 12.2 (kctd12.2), cholinergic receptor, nicotinic, alpha 3 (chrna3), ELAV like neuron-specific RNA binding protein 3 (elavl3),* and *choline O-acetyltransferase b (chatb).* The details of the expression domains are described in **Table S2**.

**Figure 4.**
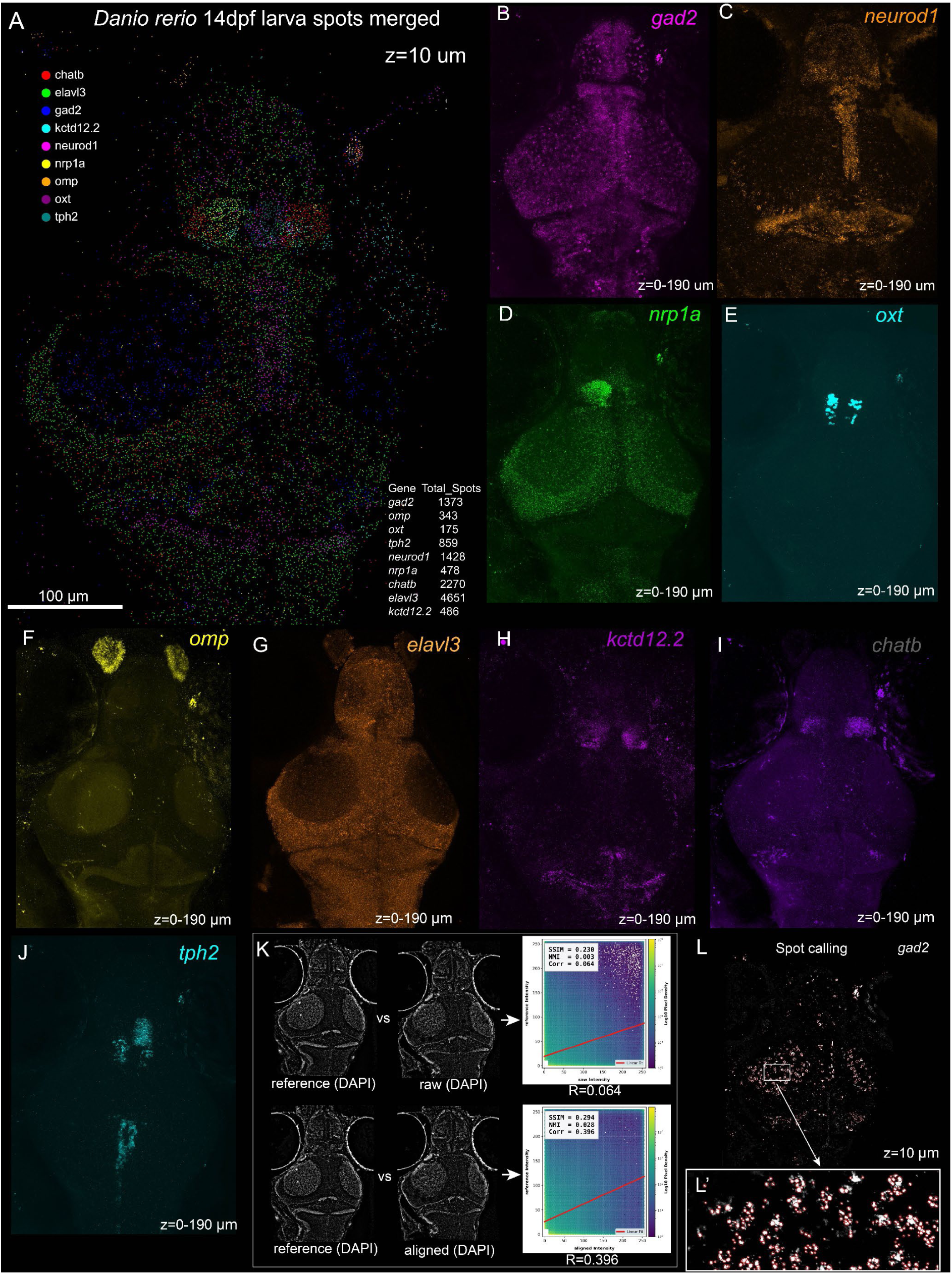
Detection of multiple genes (mRNAs) in a single 14 dpf *Danio rerio* larva across ∼190 µm depth (maximum projection of multiple z-stacks). (**A**) High-dimensional composite map displaying the spatial distribution of merged synthetic spots across the larval anatomy at a representative single focal plane (z = 10µm). An embedded quantitative table outlines the total spot counts computed across the volume for each target gene. **(B-J)** Individual maximum intensity projections (z = 0-190 µm) demonstrating the multiplexed expression profiles of target transcripts within the gel-embedded 14 dpf *Danio rerio* larva. Genes include *gad2, nrp1a*, *neurod1, oxt, omp*, *kctd12.2, chatb, tph2,* and *elavl3.* (**K**) Quantitative alignment validation framework. Dual-panel comparisons show the structural DAPI reference matched against the unaligned raw baseline (top) and the computationally corrected, registered output (bottom). Adjacent 2D density plots demonstrate a substantial correction of spatial drift, with the Pearson correlation coefficient (R) increasing from 0.064 to 0.396, Normalized Mutual Information (NMI) from 0.003 to 0.028, and the Structural Similarity Index (SSIM) improving from 0.230 to 0.294. Note the DAPI signal is acquired after DNase I treatment. (**L-L’**) Transcript validation metrics for spot calling illustrated via the *gad2* channel at z = 10 µm: (**L**) Main focal plane mapping localized spots. (**L**’) High-magnification inset detailing high-density cluster resolution, where individual verified transcripts are cleanly demarcated by red validation rings.

### Computational Alignment and Spatial Extraction in Thick Tissue Volumes

To validate the semi-quantitative capabilities of RAMFISH in thick, 3D specimens, we applied our computational pipeline to the individual z-slices of 14 dpf zebrafish brain dataset. Unlike free-floating tissues, maintaining spatial registration across a 190 μm volume over multiple hybridization cycles requires correcting for localized hydrogel expansion and optical drift. We utilized the ubiquitous DAPI nuclear stain as a constant structural anchor for our B-spline non-rigid registration against 2D image planes. As demonstrated in **Figure 4K**, applying this computational alignment significantly corrected spatial drift between cycles (ref DAPI vs raw file from subsequent round, and ref DAPI vs aligned DAPI from the subsequent round), improving both the Pearson correlation coefficient (R=0.064 to 0.396) and the Structural Similarity Index (Larkin, 2015; Wang et al., 2004) (SSIM = 0.230 to 0.294), and Normalized Mutual Information (Pluim et al., 2003; Studholme et al., 1999) (NMI = 0.003 to 0.064) of the tissue architecture. We also verified the alignment against the butterfly wing tissues by comparing the rough alignment using ramfish_rough_aligner (raw) and fine alignment using ramfish_fine_aligner (aligned) against a reference gene staining showcasing improvement in the values of SSIM, NMI, and R after the fine alignment (**Figure S16**). The alignment_score.py script was used for scoring (see Availability of resources) where in the same folder the ref, raw, and aligned images are placed and the script was executed by typing in terminal: python alignment_score.py.

Following the alignment, we applied the LoG spatial filter to extract transcript coordinates. Even within highly dense neural structures, the algorithm successfully resolved distinct spatial puncta, as shown for *gad2* at a specific optical slice (z = 10 μm) (**Figure 4L**). By integrating the extracted spots from all evaluated genes at this specific optical depth, we generated a highly multiplexed, cross-sectional transcript map of the larval brain at 10 µm, 30 µm, 60 µm, and 90 µm (**Figure 4A**; **Figure S11**).

**Video 1. A volume rendered 14 dpf zebrafish larva stained for multiple genes (mRNAs) across ∼190 µm depth (Images acquired as mirror images).**

**Video 2. Detection of multiple genes (mRNAs) in a 14 dpf zebrafish larva across ∼190 µm depth (Images acquired as mirror images).**

## Discussion

Spatial RNA profiling technologies have rapidly advanced, providing unparalleled insights into tissue architecture and accelerating discoveries across developmental biology, neurobiology, and cancer research (Choi et al., 2023; Coutant et al., 2023; Jung and Kim, 2023; Moffitt et al., 2018; Rao et al., 2021; Zhang et al., 2021). However, translating these advancements into routine laboratory practice can be challenging due to infrastructural needs. Many current platforms require cost-prohibitive specialized instrumentation, proprietary reagents, and/or complex computational overhead. Furthermore, while foundational iterative HCR methodologies such as EASI-FISH (Wang et al., 2021) and cycleHCR (Gandin et al., 2024) offer single-molecule localization and single-cell segmentation, they require permanent immobilization of the tissue within rigid hydrogel matrices and specialized computation infrastructure for post-processing. While effective, they introduce additional layers of complexity that may not be necessary for simpler, cellular and tissue-domain analysis of gene expression.

RAMFISH is an effective technique in such a scenario. It leverages a combination of established techniques such as hybridization chain reaction (HCR) chemistry (Choi et al., 2018) coupled with a gentle, non-destructive enzymatic signal removal (Codeluppi et al., 2018; Shah et al., 2016; Sternson et al., 2022; Wang et al., 2021). By integrating newly optimized permeabilization and hybridization buffer recipes along with a higher incubation temperature (37°C), RAMFISH improves reaction kinetics to be effective for both gel-embedded and entirely free-floating tissue. Processing tissues free-floating vastly accelerates sample handling and maximizes reagent accessibility without the attenuating interference of a polyacrylamide mesh. The viability, however, ultimately relies on the developed computational pipeline (the RAMFISH Software Suite) capable of running on most desktop-grade PCs to correct the large spatial shifts and elastic warping naturally experienced by untethered tissues. By shifting the focus on semi-quantitative, cellular and tissue level domain mapping instead of absolute single-molecule counting, RAMFISH allows for rapid evaluation of target gene expression.

The RAMFISH Software Suite features a robust computational pipeline designed to correct structural displacements and localized tissue deformations *in silico* through a tiered registration strategy and semi-quantification of RNA levels. This process begins with normalization of the images, followed by a visual rough alignment tool for batch-level spatial transformations, and a fully automated fine alignment combining an affine transform with a multi-resolution B-spline non-rigid registration. Afterwards, the suite streamlines downstream data processing by Laplacian of Gaussian (LoG)-based spot calling to quantify approximate levels of RNA and an image-merging tool capable of generating composites from up to 30 multiplexed-FISH channels. While this computational alignment is tailored for 2D images, the integrated wet-lab workflow operates directly on thick, whole tissues and intact specimens to extend multi-gene spatial mapping beyond the physical limitations of thin sectioning. Finally, while manual implementation is fully supported, the entire workflow has been optimized for integration with an open-source automated fluidic system (Banerjee et al., 2026) to significantly reduce hands-on time and ensure more efficient reagent utilization.

By combining iterative multiplexing with optional fluidic automation and a dedicated Python software suite, RAMFISH provides a practical balance between experimental scalability and ease of implementation. As demonstrated in developing butterfly larval wings and whole zebrafish larvae, the workflow enables routine spatial analysis of multiple genes across complex whole-mount specimens. This makes it broadly useful for developmental biologists, geneticists, and disease researchers mapping tissue patterning, embryonic development, and gene-regulatory networks. RAMFISH has already been applied to gain biological insights across several recent studies in our labs focusing on evo-devo of novel gene patterning and neuroscience (Banerjee and Monteiro, 2025; Goel et al., 2026; Raine et al., 2025). RAMFISH complements existing transcriptomic approaches, including bulk RNA-seq (Pertea et al., 2016), single-cell sequencing (Prakash et al., 2024) and spatial capture-based technologies (such as LCM-Seq (Banerjee et al., 2022)), by providing semiquantitative and image-based validation of candidate gene localizations. Furthermore, multiplexed spatial readouts generated by RAMFISH can be seamlessly integrated with functional perturbation approaches, such as CRISPR-Cas9 (Jinek et al., 2012; Matsuoka and Monteiro, 2022; Murugeshan et al., 2022), RNA interference (Fire et al., 1998), or pharmacological treatments, to efficiently assess how the manipulation of individual genes influences downstream expression programs *in-situ*.

A major historical limitation in studying the evolution and development of butterfly wing patterns is the restriction of traditional mRNA detection protocols to only one to three genes per experiment (Banerjee and Monteiro, 2025; Hanly et al., 2023; Matsuoka and Monteiro, 2022; Murugeshan et al., 2022). This requires researchers to infer the spatial overlap of gene expression by comparing disparate, individual samples. Observing multi-gene topographical expression in the same intact tissue as done through RAMFISH here reduces the uncertainty in such inference. In the larval wings, both well-characterized patterning genes and previously unexamined transcripts displayed consistent, reproducible spatial domains that aligned directly with established developmental axes. We not only recapitulated the domains of 21 known patterning genes but also identified 12 novel genes, including *hth* and *dac*, crucially anchoring these new domains against established compartment boundaries (**Figure 1 and 2**). Regulatory genes *hh* and *ci* showed precise anterior-posterior (A/P) restricted expression (Banerjee and Monteiro, 2020; Schwartz et al., 1995), while *mirror* localized to lower posterior compartment cells associated with regional differentiation (Chatterjee et al., 2024). Along the proximal-distal axis, genes including *WntA* and its receptor *fz2* marked territories implicated in the specification of central symmetry system (CSS) bands (Banerjee et al., 2023; Hanly et al., 2023). Margin-associated patterning genes such as *Wnt1, vg*, and *Dll* delineated distal boundary regions (Banerjee et al., 2023), whereas vein-related transcripts including *rho* and *ensl* were enriched in vein or provein territories (Banerjee and Monteiro, 2020; De Celis et al., 1997). By transitioning from inferred developmental models to direct, multiplexed spatial observation, RAMFISH reveals the precise, overlapping expressional territories.

Detection of multiple transcripts spatially in intact, older (14 dpf) zebrafish larvae, when complex behaviors such as social behaviors begin to emerge and crystallize, is significantly challenging to study compared to 5-7 dpf larvae. Gene expression profiles are therefore often inferred or extrapolated from younger larval brains, and expression domains are inferred from disparate samples. As shown there, the RAMFISH workflow developed uniquely enables the simultaneous visualization of multiple transcripts in the intact brain of 14 dpf larvae. Rather than inferring domain via mapping 1-3 genes at a time, we directly mapped the 3D spatial expression of neurotransmitter synthesis associated genes (*tph2, oxt, gad2, and chatb*), receptors (*nrp1a, and kctd12.2),* and neurodevelopmental markers *(elavl3, neurod1, and omp)* with spatial distributions consistent with known anatomical and functional organization of the brain (Shainer et al., 2023). Signal detection was equally robust throughout the brain tissue regardless of the depth or sparsity of expression. Genes expressed deep within the brain, such as *oxt* and *tph2* (∼100-150 µm from the dorsal surface), were clearly visible at a cellular level, on par with those localised closer to the dorsal surface, such as *neurod1* and *nrp1a*, providing confidence that RAMFISH is well suited to thick tissue and whole-mount applications.

The high reproducibility of expression patterns across samples and cycles supports the robustness of the methodology for multi-round imaging in intact tissues. By enabling routine mapping of multiple transcripts within the same specimen using accessible resources, RAMFISH offers a practical framework for linking functional genetics with spatial systems-level analysis across diverse model organisms. We envisage these results will inspire researchers to apply RAMFISH to a variety of organisms and systems (Mathuru et al., 2020) with greater ease and fuel new discoveries.

### Probes preparation

To aid the probe design process, we developed an online web-based probe designer that can generate HCR3.0 style probes. Link: https://tirthadasbanerjee.com/hcrprobedesigner/hcr_22.1.html

Input from the user requires: gene name, gene sequence, selection of amplifier, selection of GC concentration threshold, and the gap in between the probe pairs. Clicking ‘Design Probe’ and ‘Download Excel File’ exports the probe sequence formatted for orders from Integrated DNA Technologies (IDT) as oligos (100 µM) or as Pools (oPool at 50 pmol/oligo).

### RAMFISH Software Suite

The RAMFISH Software Suite is an open-source computational pipeline engineered to handle multiplexed spatial domain datasets generated from the RAMFISH workflow. The pipeline is capable of running on most desktop-grade PCs with an NVIDIA GPU as described below. It processes, high-resolution microscope channels by integrating GPU-accelerated intensity normalization, full-stack visual rough alignment, automated non-rigid B-Spline registration, and multi-core mRNA spot segmentation. By standardizing coordinate space across iterative hybridization rounds and stripping away complex background autofluorescence, the suite empowers researchers to seamlessly reconstruct publication-grade, high-dimensional composite spatial expression maps across intact tissue architectures.

### Hardware Requirements

Optimum Configuration: CPU: AMD Ryzen 5 5600X / Intel Core i5-12400F; GPU: NVIDIA RTX 3060; RAM: 32GB DDR4.

High-Performance Configuration: CPU: AMD Ryzen Threadripper 9965WX; GPU: NVIDIA RTX 5080 or 5090; RAM: 128GB DDR5.

Operating Systems Tested: Windows 11, Ubuntu 24.04.

### Prerequisite Installation

Install the following core applications prior to executing the pipeline:

Python 3.13 (or newer version): Install from the Microsoft Store or the official Python website (https://www.python.org/).

Web Browser: Google Chrome (https://www.google.com/chrome/) or Mozilla Firefox (https://www.firefox.com/).

Imaris Utilities: Imaris Viewer and Imaris File Converter from the official website (https://imaris.oxinst.com/).

Integrated Development Environment: Visual Studio Code (VS Code) from the official website (https://code.visualstudio.com/).

Run terminal and install the prerequisites. Type the following command:

`pip install numpy cupy-cuda12x[ctk] imageio imagecodecs opencv-python tifffile flask SimpleITK fastapi uvicorn python-multipart pandas pillow scikit-image tqdm`

If you are running the pipeline on an older GPU run the following command:

`pip install numpy cupy-cuda11x[ctk] imageio imagecodecs opencv-python tifffile flask SimpleITK fastapi uvicorn python-multipart pandas pillow scikit-image tqdm`

Load the image files (imaging in the current version was done at 20x lens at 2k or 4k resolution) from the microscope software (Olympus FLUOVIEW was used in this study) in Imaris file converter and convert the files into.ims format. Open the.ims file in Imaris viewer and export the individual channels including DAPI in high-resolution (4000×4000 pixels) at 600 dpi in.tiff format. There is no need to orient or scale the images at this stage as it can be done using ramfish_rough_aligner mentioned below. Make sure the lines at the boundary of your images are not exported along with your image.

### Pipeline Folder Structure and Execution

#### 1) 1_normalization

##### Description

To resolve intensity discrepancies across iterative multiplexed imaging cycles, this module uses a GPU-accelerated standardization protocol to equalize brightness and contrast. Multichannel inputs are initially converted into a unified grayscale representation by taking the maximum intensity across available color dimensions. Local background variations are resolved via background subtraction, where a heavy Gaussian blur filter estimates non-uniform illumination profiles, which are subsequently subtracted from the raw input matrix to isolate native biological signal. Finally, the 99.5^th^ (user can change from 99.0 to 99.9 depending on the non-specific signals/autofluorescence of the images) percentile of background-subtracted pixel intensities is computed to establish a robust upper-bound scaling target (filtering out hot-pixel anomalies), normalizing the overall matrix into a clean, uniform 16-bit output.

##### Usage Directories

rawdata/ (place high-resolution exports from Imaris Viewer here).

##### Script Name

ramfish_normalization.py

##### Execution Command

`python ramfish_normalization.py`

##### Configuration File (config.json)

The script execution parameters are dynamically managed via a modular config.json configuration file located in the same working directory. This file contains a settings block allowing researchers to tune the image-processing thresholds without modifying the underlying Python code:

1) raw_dir & out_dir: Define the local file path strings for the high-resolution input directory and the destination folder for the processed images, respectively.
2) output_prefix: A custom text string (default: NORM_) added to output filenames to ensure traceability and prevent accidental reprocessing.
3) valid_extensions: A list of valid image extensions (e.g.,.tif,.png,.jpg) defining the target formats the script will actively scan and parse.
4) target_intensity: The mathematical maximum value to which the target percentile is rescaled (default: 60000.0), carefully calibrated to maximize signal dynamic range while preventing bit-depth clipping or pixel saturation in 16-bit space.
5) blur_sigma: The standard deviation (expressed as a pixel radius) for the Gaussian convolution filter, dictating the spatial scale used to model and subtract non-uniform background illumination.
6) percentile: The robust intensity cutoff percentile (typically set between 99.0% and 99.9%) used to establish the true signal maximum while intentionally ignoring bright, single-pixel hot spots or transient noise artifacts during contrast scaling.

##### Output

Normalized structural outputs are automatically compiled and saved inside a generated directory named normalized_data/.

2) 2_rough_aligner

##### Description

Rigid SimpleITK-based B-Spline registration frameworks (Lowekamp et al., 2013) (https://simpleitk.org/) frequently fail when confronted with heavy geometric deformations, rotational shifts, or physical tissue inversion (flipping) between hybridization cycles. To address this, ramfish_rough_aligner provides a hybrid interactive application bridging the gap between visual micro-manipulation and high-resolution spatial data matrices. By pairing a lightweight browser UI with a Python image-processing backend, users can quickly load large-scale tissue images (such as sequential DAPI rounds) to visually align them using real-time structural adjustments for X/Y translation, overall scale, rotation, flipping, and overlay opacity. Once visual alignment is achieved on a compressed local preview, the Python backend translates those UI coordinates into exact mathematical affine transformation matrices based on the native pixel ratio, batch-transforming every associated gene channel in the directory without signal loss.

##### Usage Directories

Round_1/, Round_2/, Round_3/…Round_N/ (manually organize the corresponding normalized files into their respective round sequence folder names). Create additional folders based on the number of multiplexed rounds.

##### Script Name

ramfish_rough_aligner.py.

##### Execution command

`python ramfish_rough_aligner.py`

##### UI Execution

A local full-stack web interface will launch automatically within your default browser. Select your baseline Round_1 image as your absolute fixed structural anchor and select the corresponding moving reference images for downstream folders (e.g., matching the reference target inside the Round_2/ directory). Click “Load Images into UI” to initialize the views (the base channel mounts at 100% opacity, while the moving channel presents at 50%). Adjust the Target Opacity, Image Scale, Rotation (Deg), X Translate, Y Translate, and Flip Horizontal tools until the structural features match. Click “Apply Rough Alignment to Folder” to batch-transform every other target gene channel inside that round’s folder. An illustration of the UI is provided in **Figure S17**.

##### Output

Transformed structural files are updated directly inside their native sequence directory (e.g., Round_2/) prefixed as ALIGNED_.

3) 3_fine_aligner

##### Description

To ensure high-fidelity spatial registration across iterative hybridization cycles, a hierarchical fine-alignment architecture maps all sequential imaging datasets back to the absolute Round_1 coordinate grid. A global affine matrix transformation is calculated to fine-tune remaining structural displacements, rotational shifts, or minor scaling discrepancies. Following rigid correction, a non-rigid B-Spline deformable registration algorithm (via SimpleITK (Lowekamp et al., 2013) (https://simpleitk.org/)) tracks structural landmarks (such as localized nuclei via DAPI) to compensate for localized elastic tissue distortions. This dual-stage processing layout models complex elastic morphing phenomena, including localized non-uniform stretching, expansion, or tissue shrinkage. Executed as a clean client-server framework, the backend warps and resamples target channel arrays using linear interpolation to bind all target genes into a single, unified coordinate space.

##### Usage Directories

Round_1/, Round_2/…Round_N/. Migrate roughly aligned (ALIGNED_) entries into these active folders, ensuring only the roughly adjusted images occupy the workspace. Create additional folders based on the number of multiplexed rounds.

##### Script Name

ramfish_fine_aligner.py

##### Execution Command

`python ramfish_fine_aligner.py`

##### UI Execution

A dedicated browser alignment console will launch automatically. Specify your multiplex constraints within the “Total FISH Rounds” field. Designate your static anchor file using the “Choose File” tool inside the “Round 1 (Global Fixed Reference)” section. Navigate down to the sequential configuration interfaces (e.g., “Round 2 Alignment”) to load your “Structural Reference” and corresponding “Target Genes” (supports multi-file batch selections). Select “Run Batch Alignment” to calculate registration fields. An illustration of the UI is provided in **Figure S18**.

##### Output

Aligned channels are compiled inside a fresh directory named aligned_outputs/. To support smooth handling during the spot-calling phase, filenames are unified using standardized prefixes indexing their source imaging cycle (e.g., R2_aligned_…).

##### Note

Depending on system resources, image size, and batch count, execution spans from several minutes to an hour. For 3D image registration outside the core RAMFISH software architecture, cell boundaries can be mapped in Imaris 10.2 via the native Image Alignment toolkit (Image Processing → Image Alignment → Align Images), with 3D renderings captured using Imaris Viewer 10.2.

4) 4_spot_counter

Description: The spot detector and counter utilize a parallelized Laplacian of Gaussian (LoG) blob detection algorithm (via skimage.feature.blob_log) with automatic scale selection (Lindeberg, 1998) to identify fluorescent mRNA transcript spots and localized signal clusters. Rather than enforcing absolute single-molecule localization, the engine is optimized for robust, semi-quantitative spatial profiling where detected blobs represent either individual transcripts or overlapping multimolecular mRNA aggregates clumped within diffraction-limited domains. The engine reads multi-format spatial images using OpenCV, converts them to grayscale, and standardizes their data ranges to a floating-point matrix scale [0.0, 1.0]. It then generates a robust scale-space representation by applying Gaussian convolution kernels across a sizing spectrum bounded by your minimum and maximum spot radius settings. The initial peak-detection sensitivity functions as a gate to capture localized intensity maxima. Following blob localization, a hard intensity cutoff filter is applied to eliminate diffuse autofluorescence noise from the final transcript segmentation. The entire batch sequence runs on a multi-core architecture using a ProcessPoolExecutor pipeline for rapid local computation.

##### Usage Directories

rawdata/ (migrate your registered, high-resolution outputs from the fine-alignment directory here; it is recommended to rename files directly to their respective target gene names, e.g., ci.jpg).

##### Script Name

ramfish_spotcounter.py

##### Execution Command

`python ramfish_spotcounter.py`

##### Configuration File (config.json)

The script execution parameters are dynamically managed via a modular config.json configuration file located in the same working directory. This file contains a “settings” block allowing researchers to tune the spot-counting sensitivity thresholds without modifying the underlying Python code:

1) blob_min_sigma: The minimum expected spot radius in pixels (default: 2). Decrease this value if small, sharp points are being missed.
2) blob_max_sigma: The maximum expected spot radius in pixels (default: 5). Increase this value if larger, brighter clusters are being ignored.
3) blob_num_sigma: The number of discrete intermediate size steps checked between the minimum and maximum sigma boundaries (default: 4).
4) blob_threshold: The initial peak-detection filter sensitivity (default: 0.30). Lower values capture dimmer spots but introduce noise, while higher values capture only stark, high-contrast peaks.
5) intensity_cutoff: The absolute brightness barrier used to eliminate tissue autofluorescence, scaled strictly from 0.0 to 1.0 (default: 0.25). To calculate: Divide your target image intensity cutoff count by your images maximum bit depth (e.g., an intensity noise floor of 16,384 counts on a 16-bit image = 16384/65535 = 0.25).
6) synthetic_spot_radius: The radius (in pixels) of the rendered dots in the synthetic output images. Increase this value to make the spots and clusters larger and more visible in the final multiplexed merge.

Output: The script automatically orchestrates progress updates via a tqdm status bar and structures three distinct directories within your workspace:

1) data_sheets/: A master folder containing an individual_csvs/ subdirectory archiving coordinate logs (x, y coordinates and intensities) mapped per gene, a processing exception log (ERROR_LOG.csv), an overall count sheet (BATCH_SUMMARY_COUNTS.csv), and a master spatial coordinate file (ramfish_master_spots.csv).
2) spot_images/: Exports diagnostic validation images overlaying a sharp red circle around every verified transcript center to visually inspect detection accuracy against background tissue noise.
3) synthetic_spots/: Contains the binarized, zero-background synthetic dot images (white pixels on an isolated black canvas) used in the next step to render the final multi-channel multiplexed-FISH map.

**5) 5_merger**

Description: To reconstruct the comprehensive spatial expression map across your target anatomy, individual localized coordinates of detected spots from all sequential rounds are integrated into a single, unified coordinate system. Because all transcript channels share the precise B-Spline deformation grid anchored to the Round 1 reference, the discrete spatial coordinates of each target gene can be seamlessly overlaid. This merging process generates a high-dimensional, composite multiplexed-FISH map, providing a global view of combinatorial gene expression across the intact tissue architecture.

##### Usage Instructions

The directory includes the ramfish_merger.html application tool. Open this file locally within your browser and load your target channels from your synthetic_spots/ index using the “Choose Files” interface (supports up to 30 distinct multiplex channels simultaneously). Ensure target files follow the nomenclature rule gene_name.file_extension (e.g., ptc.jpg, hh.jpg, omb.jpg) and denote your structural template layer strictly as DAPI.jpg (the script handles the DAPI layer automatically, rendering it as a gray, semi-transparent backdrop). The application enables dynamic user configurations for background noise floors, gamma curves, and layout adjustments for figure legends across four quadrant anchors. Select “Generate Composite” to export your high-dimensional panel. An open-access web portal deployment of this module is actively available at: https://tirthadasbanerjee.com/assets/tools/ramfishmerger.html. An illustration of the UI is provided in **Figure S19**.

##### Output

Generates a publication-grade, multi-color composite spatial map matching the native resolution of your primary data inputs.

### Robustness analysis: Measurement of signal quality after multiple cycles and signal removal

We performed a robustness analysis to quantify signal degradation and stripping efficiency due to repeated multiplexing cycles. To evaluate signal retention, we examined the expression of *omb* and *rho* across different imaging rounds and independent samples (**Figure S9**). Transcript abundance was directly quantified using the ramfish_spotcounter script. We confirmed that *omb* expression remained qualitatively robust between Round 1 and Round 12, maintaining clear spatial expression domains spanning the anterior-posterior boundary and within the eyespot centers as expected (**Figure S9G-L**). While spatial fidelity was preserved, a quantitative reduction in total transcript spot counts was observed in Round 12 compared to Round 1, indicating some expected RNA degradation over multiple chemical cycles (**Figure S9G-L**). Additionally, *rho* transcripts were reliably detected and quantified during late imaging rounds (Round 12) and across independent experimental replicates, demonstrating methodological consistency (**Figure S9A-F**).

To ensure that multiplexed targets do not suffer from optical carryover, we validated chemical signal stripping efficiency across successive RAMFISH hybridization cycles (**Figure S13 and S14**). Active transcript cohorts (e.g., *gad2, kctd12.2, nrp1a*) were successfully targeted, and automated spot-calling verified high-density transcript detection (**Figure S13S and U; Figure S14T**). Following chemical signal removal steps, residual spot-calling confirmed the near-complete erasure of localized fluorescent features within identical regions of interest (**Figure S13T, V**). Quantitative evaluation contrasting total spot counts immediately after hybridization against residual traces post-stripping confirmed a robust probe re-hybridization dynamic range across multiple cycles without significant signal carryover (**Figure S13W and 14S**).

## Materials and labware used

### Equipment

1) For the manual multiplexing protocol

37°C incubator: Incubating Shaker Rocking, Brand: Ohaus, United States.

Pipettes: Micropipette, 10-1000 µL (Eppendorf, Hamburg, Germany; Cat. no.: 3123000020, 3123000055, 3123000063)

Confocal microscope: Olympus FLUOVIEW FV3000 confocal LSM (Olympus Life Science, Waltham, MA, USA; Product ID: FV3000)

2) For the automation system

Automation systems (Multiplexer and RemBot): Custom-built (Construction article available at: https://www.biorxiv.org/content/10.64898/2026.04.17.713973v1).

Pipettes: Micropipette, 10-1000 µL (Eppendorf, Hamburg, Germany; Cat. no.: 3123000020, 3123000055, 3123000063).

Confocal microscope: Olympus FLUOVIEW FV3000 confocal LSM (Olympus Life Science, Waltham, MA, USA; Product ID: FV3000).

### Consumables (common)

Pipette tips: Filter pipette tips, 10 µL, 300 µL, 1250 µL. Biotix, San Diego, CA, USA; (Cat. No.: 63300041, 63300045, 63300047).

Confocal dish: Confocal dish. 35 mm, SPL (Cat No. 101350).

Coverslips: Cover Glasses 18 mm Ø, Thickness No.1 Circular (Cat No. 0111580). Corning® cover glasses square, No. 1, W x L 18 mm x 18 mm (Cat No. CLS284518-2000EA).

Glass-slides: Slides, microscope plain, size 25 mm × 75 mm (Cat No. S8902-1PAK).

### Buffers

All the chemicals, unless mentioned, were ordered from Sigma-Aldrich (Merck).

### RAMFISH buffers

20% EC hybridization buffer:

### Composition

20% v/v ethylene carbonate (Cat No. E26258-3KG), 5× sodium chloride sodium citrate (SSC), 12 mM citric acid (pH 6.0) (Cat No. 251275-100G), 0.5% Tween 20 (Cat No. P1379-100ML), 100 µg/mL heparin (Cat No. H3393-50KU), 1.2× Denhardt’s solution (Cat No. D6001-50G), 2.5% dextran sulfate (Cat No. D6001-50G), 1.0mM EDTA (pH 8.0) (Cat No. E9884-100G).

For 50 mL of solution:

Mix in a 50 ml tube 10 mL ethylene carbonate (Cat No. E26258-3KG), 12 mL of 20× SSC, 400 µL 1 M citric acid (pH 6.0 (Cat No. 251275-100G), 200 µL of Tween 20 (Cat No. P1379-100ML), 400 µL of 10 mg/mL heparin (Cat No. H3393-50KU), 1000 µL of 50× Denhardt’s solution (Cat No. D6001-50G), 5 mL of 25% dextran sulfate (Cat No. D6001-50G), 100 µl of 500mM EDTA (pH8.0) (Cat No. E9884-100G). Fill up to 50 mL with DEPC (Cat No. D5758-25ML) H_2_O.

20% EC wash buffer:

Composition

20% ethylene carbonate (Cat No. E26258-3KG), 5× sodium chloride sodium citrate (SSC), 12 mM citric acid (pH 6.0) (Cat No., 0.5% Tween 20 (Cat No. P1379-100ML), and 100 µg/mL heparin (Cat No. H3393-50KU).

For 50 mL of EC wash solution:

Mix in a 50 ml tube 10 mL ethylene carbonate (Cat No. E26258-3KG), 10 mL of 20× SSC, 400 µL 1 M citric acid (Cat No.), pH 6.0, 200 µL of Tween 20 (Cat No. P1379-100ML), 400 µL of 10 mg/mL heparin (Cat No. H3393-50KU). Fill up to 50 mL with DEPC H_2_O.

Note

The use of ethylene carbonate accelerates the overall FISH hybridization time. Previous reports have suggested the use of 15-20% ethylene carbonate as an alternative to accelerate DNA-FISH experiments (Kalinka et al., 2020; Matthiesen and Hansen, 2012). 10% ethylene carbonate has also been used in hybridization mixture and wash buffer in MERFISH experiments (Fang et al., 2023; Moffitt et al., 2016). The buffer presented in this study will freeze at 4°C. Kindly thaw before starting the experiment or prepare fresh.

Permeabilization Solution:

Composition

4.0% Ammonium lauryl sulfate (Cat No. 09887-250ML), 5.0% Tween 20 (Cat No. P1379-100ML), Tris-HCl pH 7.5 (Cat No. 10812846001), 1.0 mM EDTA pH 8.0 (Cat No. E9884-100G), and 200.0 mM NaCl (Cat No. S9888-25G).

For 50 mL of Solution:

Mix in a 50 ml tube 10.00 mL 20% Ammonium lauryl sulfate (Cat No. 09887-250ML) (filtered), 2.50 mL Tween 20 (Cat No. P1379-100ML), 2.50 mL 1M Tris-HCl (Cat No. 10812846001), pH 7.5, 0.10 mL 0.5 M EDTA (Cat No. E9884-100G) pH 8.0, and 2.00 mL 5 M NaCl (Cat No. S9888-25G). Fill up to 50 mL with DEPC H_2_O.

Note

The use of ammonium lauryl sulfate provides an alternative to the harsh sodium dodecyl sulfate (SDS) for permeabilization of the tissue such as embryos and larvae. Store at RT for up to 6 months.

Fluorescent Mounting Buffer:

Composition

70% glycerol (Cat No. G7893-500ML) with 1.0mM EDTA (pH 8.0) (Cat No. E9884-100G). For a 50 mL solution in a 50 mL tube, add 35 mL 100% glycerol and 100 µl of 500 mM EDTA. Top up to 50 ml using DEPC water, and store at room temperature for up to 6 months.

Signal removal and wash solutions

Signal removal solution (1ml)

In a 1.5 ml tube add Tris pH7.5 (Cat No. 10812846001 (1M) – 100 µl, MgCl_2_ (Cat No. M8266-100G) (0.5M) – 50 µl, CaCl_2_ (Cat No. C4901-100G) (0.5M) – 5 µl, DNaseI (ThermoFisher-Cat No.: EN0521) – 15 µl. Add dH_2_0 to fill till 1ml. Prepare fresh.

Note

Signal removal solution can remove the signal when incubated at 37°C for 30-60 mins when performed manually and as fast as 15 mins in the robotic system in samples up to 50-100 µm thick.

### Signal wash buffer 1 (10ml)

In a 15 ml tube, add Tris pH7.5 (Cat No. 10812846001) (1M) – 1000 µl, MgCl_2_ (Cat No. M8266-100G) (0.5M) – 500 µl, CaCl_2_ (Cat No. C4901-100G) (0.5M) – 50 µl. Add dH_2_0 to fill till 10ml.

### Signal wash buffer 2 (1ml; optional)

In a 1.5 ml tube, add Tris pH7.5 (Cat No. 10812846001 (1M) – 100 µl, MgCl_2_ (Cat No. M8266-100G) (0.5M) – 50 µl, CaCl_2_ (Cat No. C4901-100G)(0.5M) – 5 µl, 500mM EDTA pH 8.0 (Cat No. E9884-100G) – 5 µl, 10% SDS (Cat No L3771-100G) - 150 µl. Add dH_2_0 to fill till 1ml.

Note

Prepare the signal removal solution fresh. Signal Wash buffer 1 and 2 can be stored at room temperature for up to 2 months.

### Robot cleaning solution

Composition

2% SDS (Cat No. 436143-25G), 1% Sodium dichloroisocyanurate (Cat No. 218928-25G), and 1% NaOH (Cat No. 221465-25G).

Alternatively, prepare 20ml of solution by mixing 5ml RNaseZap (ThermoFisher, Cat No. AM9780) and 15ml DEPC water.

**HCR3.0 Buffers (**from Choi et al., 2018; Bruce et al., 2021)

30% probe hybridization buffer

Composition

30% formamide (Cat No. F7503-1L), 5× sodium chloride sodium citrate (SSC), 9 mM citric acid (Cat No. 251275-100G) (pH 6.0), 0.1% Tween 20 (Cat No. P1379-100ML), 50 µg/mL heparin (Cat No. H3393-50KU), 1× Denhardt’s solution (Cat No. D2532-5X5ML), and 5% dextran sulfate (Cat No. D6001-50G).

For 40 mL of solution:

Mix in a 50 ml conical tube (Cat No. BS-500-MJL-S) 12 mL formamide (Cat No. F7503-1L), 10 mL of 20× SSC, 360 µL 1 M citric acid (Cat No. 251275-100G), pH 6.0, 40 µL of Tween 20 (Cat No. P1379-100ML), 200 µL of 10 mg/mL heparin (Cat No. H3393-50KU), 800 µL of 50× Denhardt’s solution (Cat No. D2532-5X5ML), and 4 mL of 50% dextran sulfate (Cat No. D6001-50G). Fill up to 40 mL with DEPC (Cat No. D5758-25ML) H_2_O.

30% probe wash buffer

Composition

30% formamide (Cat No. F7503-1L), 5× sodium chloride sodium citrate (SSC), 9 mM citric acid (Cat No. 251275-100G) (pH 6.0), 0.1% Tween 20 (Cat No. P1379-100ML), and 50 µg/mL heparin (Cat No. H3393-50KU).

For 40 mL of solution

Mix in a 50 ml tube 12 mL formamide (Cat No. F7503-1L), 10 mL of 20× SSC, 360 µL 1 M citric acid (Cat No. 251275-100G), pH 6.0, 40 µL of Tween 20 (Cat No. P1379-100ML), and 200 µL of 10 mg/mL heparin (Cat No. H3393-50KU). Fill up to 40 mL with DEPC H_2_O.

50% w/v dextran sulfate

For 40 mL of solution

Mix in a 50 ml tube 20 g of dextran sulfate powder (Cat No. D6001-50G). Fill up to 40 mL with DEPC H_2_O.

### Amplification buffer

5× sodium chloride sodium citrate (SSC), 0.1% Tween20, and 5% dextran sulfate (Cat No. D6001-50G).

For 40 mL of solution

Mix in a 50 ml tube 10 mL of 20× SSC, 40 µL of Tween 20 (Cat No. P1379-100ML), 4 mL of 50% dextran sulfate (Cat No. D6001-50G). Fill up to 40 mL with DEPC H_2_O.

### Detergent Solution

Composition

1.0% SDS (Cat No. 436143-25G), 2.5% Tween (Cat No. P1379-100ML), Tris-HCl (Cat No. 10812846001) (pH 7.5), 1.0 mM EDTA (Cat No. E9884-100G) (pH 8.0), and 150.0 mM NaCl (Cat No. S9888-25G).

For 50 mL of Solution

Mix in a 50 ml tube 5.00 mL 10% SDS (Cat No. 436143-25G) (filtered), 1.25 mL Tween 20 (Cat No. P1379-100ML), 2.50 mL 1M Tris-HCl (Cat No. 10812846001), pH 7.5, 0.10 mL 0.5 M EDTA (Cat No. E9884-100G), pH 8.0, 1.50 mL 5 M NaCl (Cat No. S9888-25G). Fill up to 50 mL with DEPC H_2_O.

### 1x PBST (for 50 ml of solution)

5ml 10x PBS, 50 µL Tween 20 (Cat No. P1379-100ML). Fill up to 50mL with water.

### 5× SSCT (for 40 mL of solution)

10 mL of 20× SSC, 40 µL of Tween 20 (Cat No. P1379-100ML). Fill up to 40 mL with DEPC H_2_O.

### DEPC H_2_O (1000 ml)

Add 1ml of DEPC to 1000 mL of MilliQ water. Mix well and store in the dark overnight. The next day, autoclave.

10x PBS

For 1 liter of solution, add 81.8 g of NaCl (Cat No. S9888-25G), 5.28 g of KH_2_PO_4_ (Cat No. P0662-25G), and 10.68 g of K_2_HPO_4_. (Cat No. P3786-100G) in a glass bottle. Fill till 1000ml using dH_2_O and autoclave. Store at RT for up to 6 months.

20x SSC

For 1000 ml of solution, add 175.3 g of NaCl (Cat No. S9888-25G) and 88.2 g of trisodium citrate (Cat No. S1804-500G). Add dH_2_O to 1000 ml and autoclave. Store at RT for up to 6 months.

Alternatively prepare, 2 mM Trolox (Cat No. 238813-5G), 10% w/v glucose (Cat No. G7021-100G), 0.1mg/ml glucose oxidase (Cat No. 345386-10KU), 50 µg/ml catalase (Cat No. C9322-5G), 5x SSC, Tris-HCl (Cat No. 10812846001), 50 mM, 60% glycerol (Cat No. G5516-100ML). Modified from ref (Zhang et al., 2021). If possible, prepare fresh. Otherwise, store at 4°C and use within 1-2 weeks after preparation.

DAPI buffer:

For 1 mL of solution, add 5 µL of stock DAPI (Cat No. D9542-5MG) (1 mg/mL in DMSO) in 995 µL of 5x SSCT in a 1.5 mL tube. Prepare fresh before use.

### Sample embedding in a confocal dish

#### Glass coatings (**Figure 1B**)

Confocal dish (35 mm with glass cover slip) (SPL, Cat No. 101350). Steps:

1. On a glass confocal dish, add 200 µL of 2% APTES (Cat No. 440140-100ML) in 100% ethanol for 5 mins, followed by two washes with 100% ethanol. Dry the dish at 50°C for 30 mins. This can also be done on a regular microscope glass slide.
2. Afterwards, add 0.5% glutaraldehyde (Cat No. G7776-10ML) in DEPC water for 30 mins and wash with DEPC treated water. Air dry the dish for 30 mins.
3. Coat coverslips (18 mm; Cat No. 0111580) with Sigmacoat (Cat No. SL2-25ML) by adding a few drops of Sigmacoat in a fume hood for 1 min, washing twice with 100% ethanol, and letting the coverslips dry.

Finally, add the fixed and permeabilized tissues and embed them in a polyacrylamide gel (see gel composition below).

#### Hydrogel casting (**Figure 1C**)

A polyacrylamide gel (**Table 1**) is necessary to keep thick 3D tissues in place during the multiple cycles of hybridization, washing, imaging, and alignment. The gel will slightly expand after buffers are added, so a gel can be cast that is narrower than the coverslip diameter of the confocal dish. The expansion factor, i.e., the diameter of the gel (Dgel) divided by the diameter of coverslips (Dcp) for different buffers, is provided in Fang et al. (2023).

**Table 1.** Composition of polyacrylamide gel.

| S.N. | Reagent | Volume | Volume (small) |
| --- | --- | --- | --- |
| 1. | NaCl (2M) (Cat No. S9888-25G) | 400 µl | 100 µl |
| 2. | Acrylamide/Bisacrylamide 19:1 (A2917-100ML) (100mg/ml) | 1200 µl | 300 µl |
| 3. | Tris-HCl 1M (Cat No. 10812846001) | 120 µl | 40 µl |
| 4. | TEMED (Cat No. T9281-25ML) | 10 µl | 2.5 µl |
| 5. | Ammonium persulfate (1% w/v) (Cat No. 248614-100G) | 20 µl | 5 µl |

Note

A smaller volume can be prepared based on user requirements. If you need the gel to solidify faster use a larger volume of TEMED and APS.

Add around 70-100 µl of the above solution on top of the glass with the tissue samples. Add the Sigmacoated coverslip on top of the gel (**Figure 1C**). After gel solidification (**Figure S20**), remove the coverslip, and transfer the confocal dish to the chamber for automation or for manual experimentation (**Figure 1A**).

#### Pause step

The samples embedded in the gel can be stored either dehydrated or in 30% PHB buffer for over 2 months before starting the reaction.

Note

For clearing tissues, these can be left in the gel in 2% SDS, 0.5% Triton-X 100 in 2x SSC at 37°C overnight (Liu et al., 2022). Alternatively, the gel can also be left in 5% SDS+2% boric acid at 37°C overnight.

## Procedure

### Manual multiplexing protocol

#### A) Probe preparation

##### Preparing the primary stock solution (10 pairs of probes: 5µM final volume)

Up to 10 pairs of oligonucleotides were used, each at a concentration of 100 µM. 100 µl of oligos from each tube were mixed to form a master mix with a final probe concentration of 5 µM in a 2 ml microcentrifuge tube (Labselect, Cat No. MCT-001-200). Three to four genes with different hairpin amplifiers (Molecular Instruments) can be designed and tested in a single cycle of HCR. The number of genes tested in each cycle depends on the number of laser lines in the confocal being used. The stock mix of primary probes can be stored at-20°C.

Prepare the 30% probe hybridization buffer or 20% EC hybridization buffer with probes complementary to the target RNA.

Preparation of DNA oligo mix: Add 10 µl of each of the 5 µM DNA oligo mixes for each gene for 8 hrs manual methodology (e.g., add 10+10+10 µl for 3 genes) in a 1.5 ml tube and adjust the volume to 1000 µl in 30% probe hybridization buffer or 20% EC hybridization buffer in a 1.5ml microcentrifuge tube (Labselect, Cat No. MCT-001-150).

##### Note

If you are targeting a lowly expressed gene, you might need to either increase the concentration of the probes for that gene or the number of probe pairs, in the DNA oligo mix. Longer primary and secondary incubation might be necessary for lowly expressed genes.

Prepare the secondary probes with fluorescent tags.

Add 5 µl of each H1 and H2 hairpins (Molecular Instruments) separately in 150+150 µl of Amplification buffer in two 200 µl PCR tubes.

Heat at 95°C for 90 secs in a thermocycler and cool down at room temperature in a dark environment for 30 mins. Mix the two tubes together in a 1.5 ml microcentrifuge tube (Labselect, Cat No. MCT-001-150) and use.

##### Note

We use the following combinations: 1) B1: AF546, 2) B2: AF647, 3) B3: AF488. We have used B4 with an AF514 tag, which sometimes cross-talks with the AF546 fluorophore.

The secondaries can be used multiple times. After using the probes, collect them from the sample and store at-20°C. For repeated use, heat the secondary probes in the dark in amplification buffer at 37-42°C for 30 mins before use.

##### B) Steps

1. Dissect the tissue in 1x PBS at Room Temperature (RT ∼ 22°C). For the dissection of butterfly larval wings, follow the protocol described in Banerjee and Monteiro, 2020b.
2. Transfer the tissue to either a 1.5ml tube or glass spot plate containing 500 µL 1x PBST supplemented with 4% formaldehyde and fix for 30-60 mins at room temperature (RT) with shaking. A smaller fixation time is required for thinner tissues, while a longer time is required for thicker tissues.
3. Wash the tissue 3 times with 500 µl 1x PBST (3mins each) at RT.
4. Add 500 µl of permeabilization solution and leave the tissue for 30 mins at 37°C.

##### Note

If clearing tissue (for transparency), the tissue can also be left in 500 µL of 10% SDS (Cat No. L3771-100G) + 2% boric acid (Cat No. B0394-100G) at 37°C for clearing for up to 10 hrs.

5) Wash 3 times with 500 µl 1x PBST (3 mins each) at RT.

6) Wash 2 times with 500 µl 5x SSCT (3 mins each) at RT.

##### Note

At this step you can either embed the sample in the acrylamide gel mentioned above or perform the experiment with the tissue free-floating in the buffer. For the free-floating experiments, the reactions are carried out on 1.5 ml microcentrifuge tubes (Labselect, Cat No. MCT-001-150) or glass spot plates (PYREX^TM^; Corning, Corning, NY, USA; Cat. No.: 722085).

7) Wash 2 times with 500 µl 5x SSCT (3 mins each) at RT.

8) Stain the tissue with DAPI by adding 5 µl of DAPI stock into 1000 µl of 5x SSCT. Incubate the sample at 37°C for 10 mins.

9) Wash the sample 3 times with 5x SSCT (3 mins each).

10) Proceed to the confocal imaging and acquire high resolution image (at 2k or 4k) of the DAPI channel using 20x lens.

11) Wash the tissue 2 times with 500 µl 5xSSCT for 3 mins at 37°C.

12) Wash 3 times with 500 µl signal wash buffer 1 at 37°C for 3 mins each.

13) Add 500 µl of signal removal solution (to create DNA nuclear footprints) and incubate at 37°C for 15-90 mins (depending on the thickness of the tissue). For wings, DNA is removed within 15 mins, and for the zebrafish embedded in acrylamide gel, DNA removal was carried out for 60-90 mins.

14) Wash 2 times with 500 µl signal wash buffer 2 at 37°C for 2 mins each (optional).

15) Wash 3 times with 500 µl signal wash buffer 1 at 37°C for 3 mins each.

16) Wash 2 times with 500 µl 5x SSCT at 37°C for 3 mins each.

17) Transfer the tissues to 500 µl 30% probe hybridization buffer or 20% EC hybridization buffer and incubate at 37°C for 30 mins.

##### Pause step

You can store the tissue in the 30% probe hybridization buffer at 4°C for over 8 weeks without any significant reduction in signal quality. Do not store samples in the 20% EC hybridization buffer, as the solution will crystallize at 4°C.

18) Prepare 1000 µl 30% probe hybridization buffer or 20% EC hybridization buffer supplemented with DNA oligos complementary to the target RNA (described above).

19) Remove the 30% probe hybridization buffer or 20% EC hybridization buffer added before and replace it with 500µl of the solution mentioned in step 9 supplemented with DNA oligos at 37°C.

20) Incubate the samples at 37°C for 4 hrs (for the detection of lowly expressed genes, the samples can be left overnight). For faster detection, primary incubation can be as short as 2 hrs.

21) Wash 6 times with 500 µl of 30% probe wash buffer or 20% EC wash buffer (5-10 mins each) at 37°C.

22) Wash 2 times with 500 µl 5x SSCT (3 mins each) at 37°C.

23) Incubate the tissue in 500 µl Amplification buffer for 10-15 mins at 37°C.

##### Pause step

You can store the tissue in Amplification buffer at 4°C for over 8 weeks without any significant reduction in signal quality.

1. Replace the Amplification buffer with 200 µl secondary probes (described above) at 37°C.

##### Note

It is critical to keep the sample away from light as much as possible from this step onwards.

25) Incubate the samples in the dark for 3 hrs at 37°C. For faster detection, secondary incubation can be as short as 1.5 hrs incubated at 37°C.

26) Wash the tissue 5 times with 500 µl 5x SSCT (in a dark environment) for 5 mins each at 37°C.

##### Note

Add 5 µL of DAPI to 500 µL 5x SSCT and incubate for 5 mins. Afterwards, wash 2 times with 500 µL 5x SSCT for 3 mins each at RT.

##### Pause step

You can store the tissue in 5x SSCT at 4°C for over 8 weeks without any significant reduction in signal quality prior to imaging.

27) Replace the 5x SSCT with the fluorescent mounting buffer and proceed for confocal imaging.

##### Note

If you are performing the experiment with free floating tissue in buffers, mount the sample on a slide, add mounting media, place the coverslip, and proceed for imaging. Do not seal the coverslip. After imaging, remove the coverslip and wash with 5x SSCT to dislodge and remove the sample from the glass slide. For experiments on microtome sectioned tissues (10-50 µm sections) on coverslips, all the reactions and washes can be directly performed on the coverslips. If you are performing the experiment in the acrylamide gel, add the fluorescent mount buffer and proceed for imaging.

##### Imaging

Take the microscope plate with the confocal dish or your samples mounted in a slide under the confocal microscope and perform the imaging. In the present work, imaging was carried out using an Olympus FV3000 confocal microscope with 405 nm, 488 nm, 555 nm, and 647 nm lasers. Images were captured at 2k or 4k resolution using 20x objective lenses. Imagining time will vary based on the sample size, lens, and resolution used for imaging the sample. A typical image with 20x lens at 4k resolution with 4 channel lasers will take approximately 1-2 min. For multiple images in z-stacks, imaging will take 10-60 mins depending on the number of layers.

##### Subsequent cycles

28) For the second cycle of hybridization with a new set of probes, you need to remove the first set of probes. To remove the signal, first wash the tissue two times with 500 µl 5xSSCT for 3 mins at RT.

29) Wash 3 times with 500 µl signal wash buffer 1 at 37°C for 3 mins each.

30) Add 500 µl of signal removal solution and incubate at 37°C for 15-90 mins (depending on the thickness of the tissue). For butterfly wings, signals were removed within 15 mins, and for the zebrafish embedded in acrylamide gel, signal removal was carried out for 60-90 mins.

31) Wash 2 times with 500 µl signal wash buffer 2 at 37°C for 2 mins each (optional).

32) Wash 3 times with 500 µl signal wash buffer 1 at 37°C for 3 mins each.

33) Wash 2 times with 500 µl 5x SSCT 1 at 37°C for 3 mins each.

##### Pause step

You can store the tissue in 5x SSCT at 4°C for over 8 weeks without any significant reduction in signal quality.

##### Note

Image the tissue after signal removal to estimate the time required for signal removal. If additional time is necessary, repeat from step 16. The acrylamide gel-based method usually requires a longer time compared to the free floating tissues. After signal removal, repeat from step 7.

### Protocol on automation platform

#### A) Probe preparation

##### Preparing the primary stock solution (10 pairs of probes: 5µM final volume)

Up to 10 pairs of oligonucleotides were ordered from IDT for one gene, each at a concentration of 100 µM. 100 µl of oligos from each tube were mixed to form a master mix with a final probe concentration of 5 µM in a 2ml microcentrifuge tube (Labselect, Cat No. MCT-001-200). Three to four genes with different hairpin amplifiers (Molecular Instruments) can be designed and tested in a single cycle of HCR (Choi et al., 2018). The number of genes tested in each cycle depends on the number of laser lines in the confocal being used. The stock mix of primary probes can be stored at-20°C.

#### Prepare the 20% EC hybridization buffer with probes complementary to the target RNA

##### Preparation of DNA oligo mix

Add 5 µl of each of the 5 µM DNA oligo mix for reactions and adjust the volume to 1000 µl in 20% EC or 30% PHB buffer in an 1.5 mL microcentrifuge tube (Labselect, Cat No. MCT-001-150).

##### Prepare the secondary probes with fluorescent tags

Add 5 µl of each H1 and H2 hairpins (Molecular Instruments) separately in 150+150 µl of Amplification buffer in two 200 µl PCR tubes.

Heat at 95°C for 90 secs in a thermocycler and cool down at room temperature in a dark environment for 30 mins. Mix the two tubes together in a 1.5 ml microcentrifuge tube (Labselect, Cat No. MCT-001-150) and use.

##### Note

You can also prepare a larger volume of H1 and H2 hairpins and store at-20°C for future use. The secondaries can be used multiple times. After the reaction, collect them from the secondary collection tube and store at-20°C. For repeated use, thaw the secondary probes in the amplification buffer before use. Longer primary and secondary incubation might be necessary for lowly expressed genes.

##### B) Steps

Fixation and permeabilization steps (done manually):

1) Dissect the tissue in 1x PBS at room temperature.

2) Transfer the tissue in 500 µl 1x PBST supplemented with 4% formaldehyde and fix for 30-60 mins at room temperature (RT) with shaking. A smaller fixation time is required for thinner tissues, while a longer time is required for thicker tissues.

3) Wash the tissue 3 times with 500 µl 1x PBST (3mins each) at RT.

4) Add 500 µl of detergent solution or alternative permeabilization buffer and leave the tissue for 30 mins at 37°C.

5) Wash 3 times with 500 µl 1x PBST (3 mins each) at RT.

6) Wash 2 times with 500 µl 5x SSCT (3mins each) at RT.

##### Note

At this step you can either embed the sample in the acrylamide gel mentioned above or perform the experiment with the tissue free-floating in the buffer. For the free-floating experiments, the reactions are carried out on 1.5 ml microcentrifuge tubes (Labselect, Cat No. MCT-001-150) or glass spot plates (PYREX^TM^; Corning, Corning, NY, USA; Cat. No.: 722085).

7) Wash 2 times with 500 µl 5x SSCT (3 mins each) at RT.

8) Stain the tissue with DAPI by adding 5 µl of DAPI stock into 1000 µl of 5x SSCT. Incubate the sample at 37°C for 10 mins.

9) Wash the sample 3 times with 5x SSCT (3 mins each).

10) Proceed to the confocal imaging and acquire a high resolution image (at 2k or 4k) of the DAPI channel using a 20x lens.

##### Note

If you are performing the experiment with free floating tissue in buffers, mount the sample on a slide, add mounting media, place the coverslip, and proceed for imaging. Do not seal the coverslip. After imaging, remove the coverslip and wash with 5x SSCT to dislodge and remove the sample from the glass slide.

11) Wash the tissue 2 times with 500 µl 5xSSCT for 3 mins at RT.

12) Wash 3 times with 500 µl signal wash buffer 1 at 37°C for 3 mins each.

13) Add 500 µl of signal removal solution (to create DNA nuclear footprints) and incubate at 37°C for 15-90 mins (depending on the thickness of the tissue). For wings, DNA is removed within 15 mins, and for the zebrafish embedded in acrylamide gel, DNA removal was carried out for 60-90 mins.

14) Wash 2 times with 500 µl signal wash buffer 2 at 37°C for 2 mins each (optional).

15) Wash 3 times with 500 µl signal wash buffer 1 at 37°C for 3 mins each.

16) Wash 2 times with 500 µl 5x SSCT at 37°C for 3 mins each.

17) Transfer the tissue to 500 µl 30% probe hybridization buffer or 20% EC hybridization buffer at RT.

##### Pause step

Tissues can be stored in the 30% probe hybridization buffer at 4°C for 8 weeks without any significant loss of signal. Do not store samples in the 20% EC hybridization buffer as the solution will crystallize at 4°C.

18) Add all the buffers to the separate wells of the probe module in the fluidics robot with thermal control – Multiplexer.

i) 5x SSCT: 15ml.

ii) 20% EC Wash buffer: 10 ml.

iii) Amplification buffer: 700 µl.

iv) 20% EC hybridization buffer: 700 µl.

v) 20% EC hybridization buffer with probes (Primaries): 700 µl.

vi) Secondary fluorescence probes in amplification buffer (Secondaries): 300 µl.

vii) DAPI buffer: 700 µl.

viii) Imaging buffer: 700 µl.

19) Transfer the confocal dish with the sample to the chamber and make sure the microcomb fits perfectly on top of the confocal dish.

20) Turn on the fluidics cycle by pressing the appropriate button (see below).

Multiplexer instruction:

Button 1: For running single sample FISH cycle: Press the “1 sample” button.

Button 2: For running dual sample FISH cycle: Press the “2 samples” button.

Button 3: For calibration: This will execute a script that is used for pump calibration.

Button 4: For cleaning: Press the Clean button. This will execute a script to flush any remaining buffer/probes in all the tubes. After each reaction, load 1 ml of the robot cleaning solution to clean all the tubing.

##### Note

The system is designed to collect the secondary fluorescent probes in a separate container. The collected probes can be reused 2-3 times (optional). The script is fully customizable based on the user’s needs (available at: https://github.com/tdblab/RAMFISH/tree/main/Hardware).

##### Imaging

Take the microscope plate with the confocal dish or your samples mounted in a slide under a confocal microscope and perform the imaging. In the present work, imaging was carried out using an Olympus FV3000 confocal microscope with 405 nm, 488 nm, 555 nm, and 647 nm lasers. Images were captured at 2k or 4k resolution using 20x objective lenses. Imaging time will vary based on the sample size, lens, and resolution used for imaging the sample. A typical image with 20x lens at 4k resolution with 4 channel lasers will take approximately 1-2 min. For multiple images in z-stacks, imaging will take 10-60 mins depending on the number of layers.

##### Note

In case of drying, rehydrate in 5x SSCT and wash 2 times in 5x SSCT, followed by the addition of the fluorescent mounting buffer before confocal imaging.

##### Pause step

You can store the tissue in 5x SSCT at 4°C after the reaction for over 8 weeks without any significant reduction in signal quality.

21) Remove the signal using RemBot. Load the following buffers in the RemBot buffer plate.

i) Signal wash buffer 1: 10 ml.

ii) Signal removal solution with DNAse I for first plate (SRS1): 500 µl.

iii) Signal removal solution with DNAse I for second plate (SRS2): 500 µl (optional).

iv) 5x SSCT: 10 ml.

RemBot instruction (buttons):

Button 1: For signal removal in a single sample setup.

Button 2: For signal removal in a dual sample setup.

Button 3: For calibration: This will execute a script that is used for pump calibration.

Button 4: For cleaning: Press the Clean button. This will execute a script to flush any remaining buffer/probes in all the tubes. After each reaction, load 1 ml of DEPC treated water to clean all the tubing.

## Authors contribution

TDB: Conceptualization, workflow architecture, hardware design and development, software design and programming, chemistry modification, methodology (butterfly wings, zebrafish), writing - original draft, writing-review and editing. JR: Methodology (zebrafish), writing – review and editing. JLCH: Methodology (butterfly wing, zebrafish). KHC: Supervision, writing – review and editing, funding. ASM: Supervision, writing – review and editing, funding. AM: Conceptualization, supervision, funding, writing – review and editing.

## Acknowledgement

We thank DBS-CBIS confocal facility and Tong Yan for access to the Olympus FV3000 confocal microscope. Shen Tian, Javen Tan, Ruijing Geng, Tanisha Goel, Linwan Zhang, and Maurice Lee for helpful tips on the methodology and improving the workflow. Kian Long Tan, Suriya Murugeshan, and Shen Tian for productive discussion and proofreading an earlier version of the manuscript. Tan Lu Wee for lab management.

## Availability of resources

RAMFISH Software Suite GitHub link: https://github.com/tdblab/RAMFISH/tree/main/Software.

Sequencing data from butterfly larval and pupal wing are available at: NCBI under Bioproject: PRJNA831205 (Matsuoka et al., 2023).

The unpublished single-cell sequencing dataset (larvalhindwing.h5) used for benchmarking in this work is available at: https://github.com/tdblab/RAMFISH/tree/main/misc/bicyclus_single_cell_larval_hindwing.

The script for alignment scores (SSIM, R, and NMI) is available at: https://github.com/tdblab/RAMFISH/tree/main/misc/alignment_score.

All the additional data mentioned in the present method are described in the supplementary files.

## Accompanying article

An accompanying article describing the assembly and operation of the Multiplexer and RemBot automation systems is available. Link: https://www.biorxiv.org/content/10.64898/2026.04.17.713973v1

## Use of Artificial Intelligence

The authors declare the use of Gemini3.1 Pro for generation of code used in this work and an early draft of the graphical abstract. All the codes and the image were afterwards verified and thoroughly tested for accuracy.

