## Supplementary File 1 for "A spatial mRNA profiling workflow using Rapid Amplified Multiplex FISH (RAMFISH)"

Affiliations

### Supplementary file 1:

In this section, additional imaging results and background studies are described.

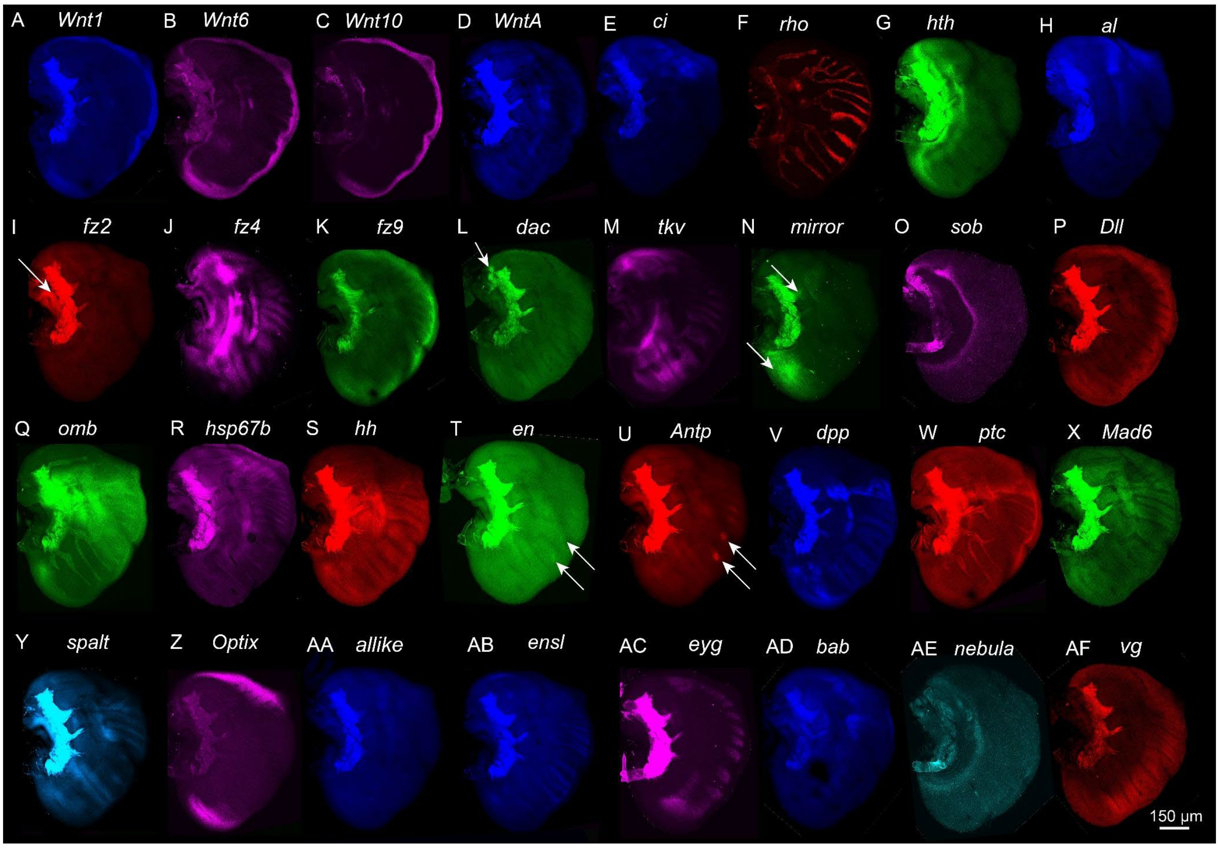

**Figure S1. Multiplexed spatial expression patterns in the larval forewing. (A-AF)** Spatial expression domains of 32 targeted genes. White arrows highlight specific localized expression features within the developing tissue. Scale bar (in AF): 150 µm. **(A)** *Wnt1,* **(B)** *Wnt6,* **(C)** *Wnt10,* **(D)** *WntA,* **(E)** *ci,* **(F)** *rho,* **(G)** *hth,* **(H)** *al,* **(I)** *fz2,* **(J)** *fz4,* **(K)** *fz9,* **(L)** *dac,* **(M)** *tkv,* **(N)** *mirror,* **(O)** *sob,* **(P)** *Dll,* **(Q)** *omb,* **(R)** *hsp67b,* **(S)** *hh,* **(T)** *en,* **(U)** *Antp,* **(V)** *dpp,* **(W)** *ptc,* **(X)** *Mad6,* **(Y)** *spalt,* **(Z)** *Optix,* **(AA)** *al-like,* **(AB)** *ensl,* **(AC)** *eyg,* **(AD)** *bab,* **(AE)** *nebula,* and **(AF)** *vg*. Note: The tracheal tissue along the veins and in the proximal domain are autofluorescent. Experiment performed using manual multiplexing in free-floating buffers.

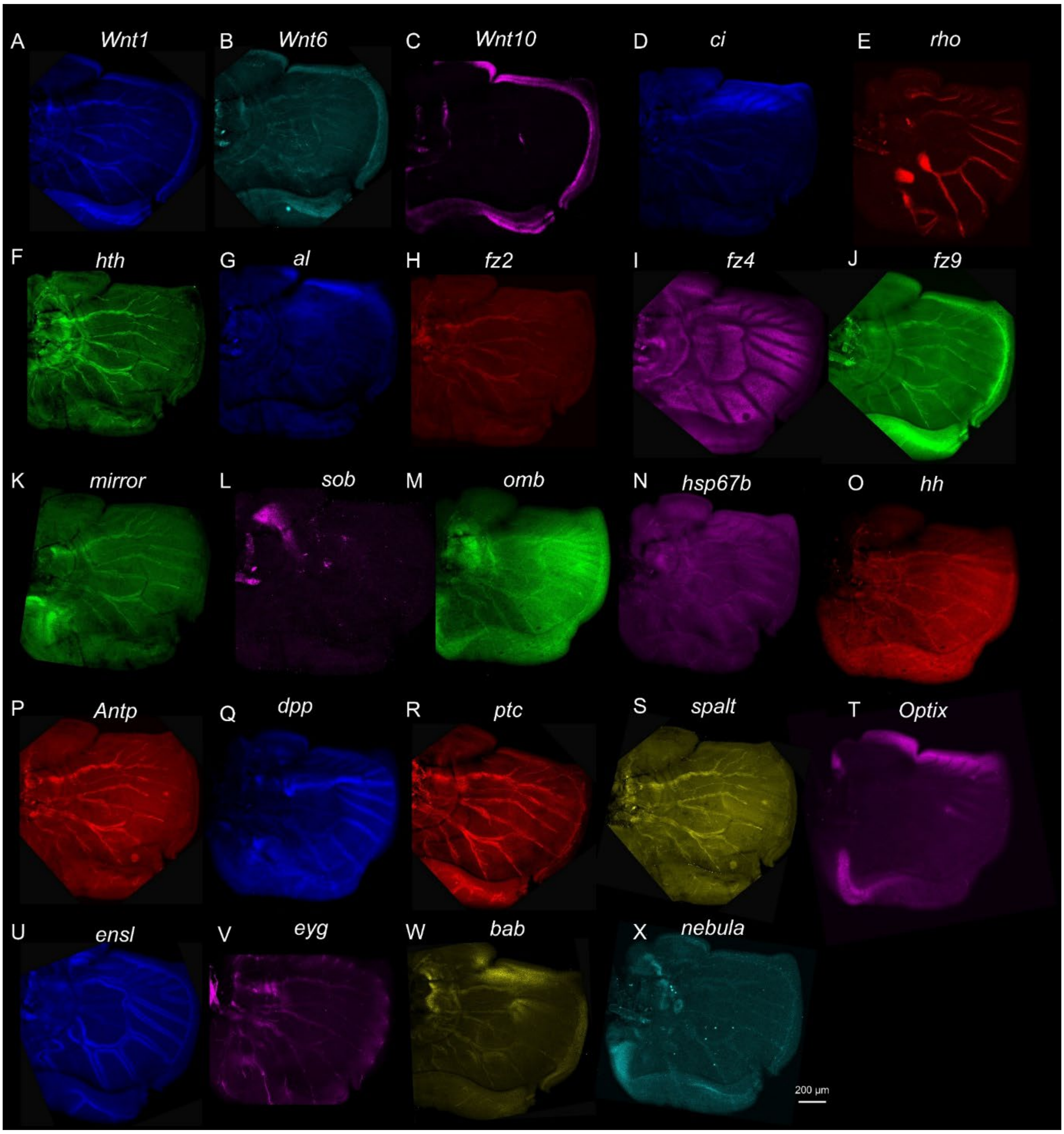


**Figure S2. Spatial expression profiles of multiplexed genes in the developing wing.** (A-X) Spatial expression domains of 24 targeted genes detected via RAMFISH. **(A)** *Wnt1,* **(B)** *Wnt6,* **(C)** *Wnt10,* **(D)** *ci,* **(E)** *rho,* **(F)** *hth,* **(G)** *al,* **(H)** *fz2,* **(I)** *fz4,* **(J)** *fz9,* **(K)** *mirror,* **(L)** *sob,* **(M)** *omb,* **(N)** *hsp67b,* **(O)** *hh,* **(P)** *Antp,* **(Q)** *dpp,* **(R)** *ptc,* **(S)** *spalt,* **(T)** *Optix,* **(U)** *ensl,* **(V)** *eyg,* **(W)**

*bab, and* **(X)** *nebula.* Scale bar (in **X**): 200 μm. Experiment performed using manual multiplexing in free-floating buffers.

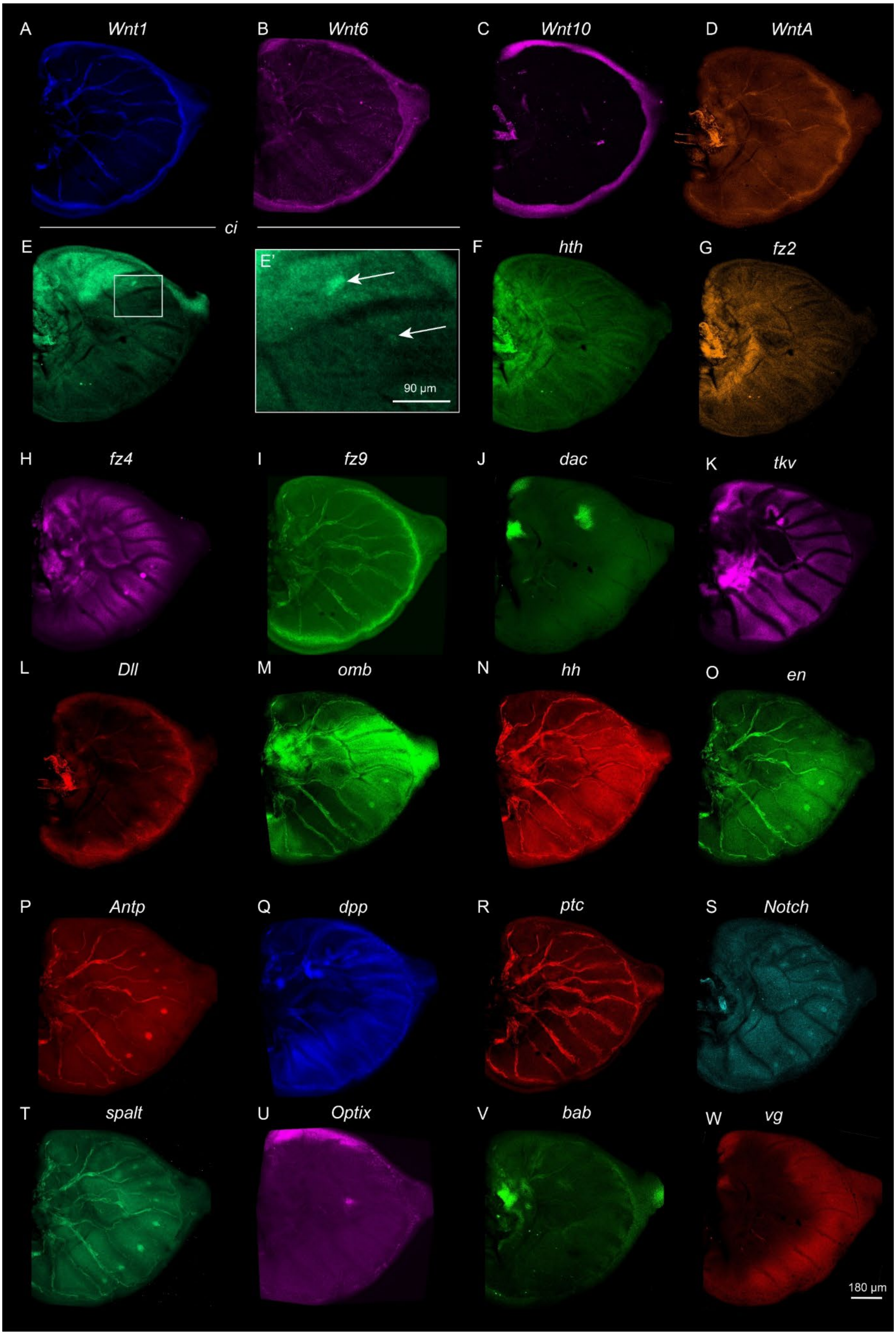
A Wnt1
B Wnt6
C Wnt10
D WntA
ci
E
E'
90 µm
F hth
G fz2
H fz4
I fz9
J dac
K tkv
L Dll
M omb
N hh
O en
P Antp
Q dpp
R ptc
S Notch
T spalt
U Optix
V bab
W vg
180 µm

**Figure S3. Multiplexed spatial expression mapping in the developing wing.** (**A-W**) Spatial expression domains of 23 targeted genes detected via RAMFISH. (**E, E'**) Detection of ci transcripts. The boxed region in (**E**) is shown at higher magnification in (**E'**), with white arrows highlighting specific localized expression features. **(A)** *Wnt1,* **(B)** *Wnt6,* **(C)** *Wnt10,* **(D)** *WntA,* **(E, E')** *ci,* **(F)** *hth,* **(G)** *fz2,* **(H)** *fz4,* **(I)** *fz9,* **(J)** *dac,* **(K)** *tkv,* **(L)** *Dll,* **(M)** *omb,* **(N)** *hh,* **(O)** *en,* **(P)** *Antp,* **(Q)** *dpp,* **(R)** *ptc,* **(S)** *Notch,* **(T)** *spalt,* **(U)** *Optix,* **(V)** *bab, and* **(W)** *vg*. Scale bars: 90 μm in **E'** and 180 μm in **W**. Experiment performed using manual multiplexing in free-floating buffers.

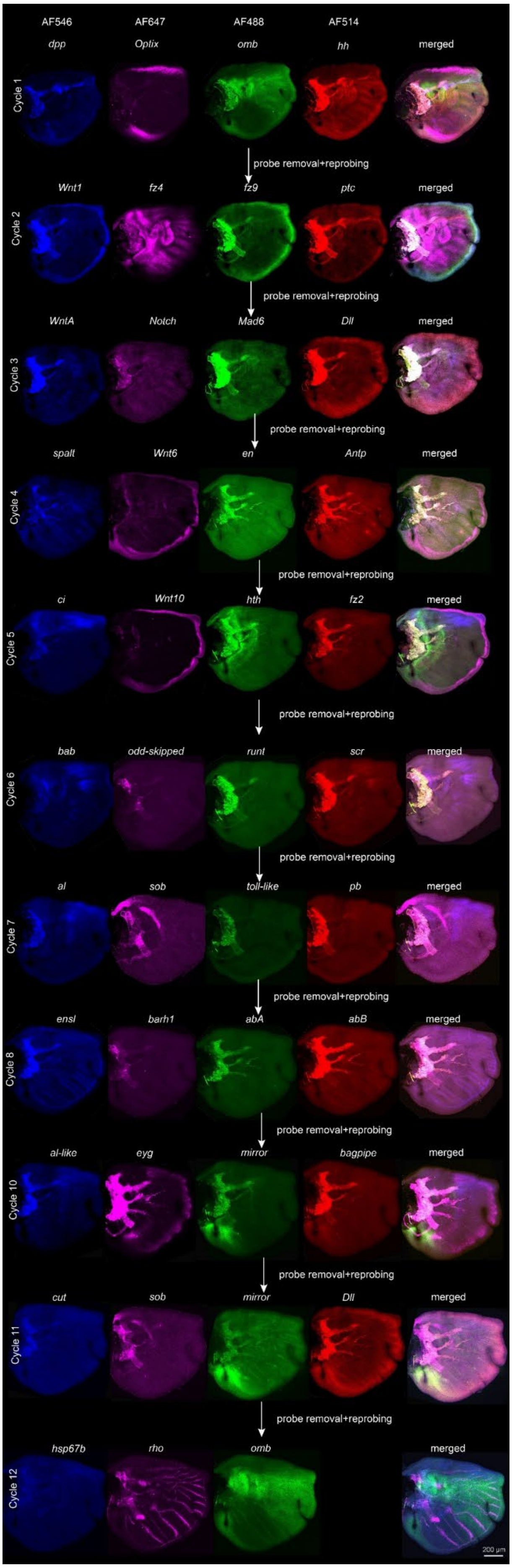
AF546
AF647
AF488
AF514
dpp
Oplix
omb
hh
merged
Cycle 1
probe removal+reprobing
Wnt1
fz4
fz9
ptc
merged
Cycle 2
probe removal+reprobing
WntA
Notch
Mad6
Dll
merged
Cycle 3
probe removal+reprobing
spalt
Wnt6
en
Antp
merged
Cycle 4
probe removal+reprobing
ci
Wnt10
hth
fz2
merged
Cycle 5
probe removal+reprobing
bab
odd-skipped
runt
scr
merged
Cycle 6
probe removal+reprobing
al
sob
toll-like
pb
merged
Cycle 7
probe removal+reprobing
ensl
barh1
abA
abB
merged
Cycle 8
probe removal+reprobing
al-like
eyg
mirror
bagpipe
merged
Cycle 10
probe removal+reprobing
cut
sob
mirror
Dll
merged
Cycle 11
probe removal+reprobing
hsp67b
rho
omb
merged
Cycle 12
200 µm

**Figure S4. Sequential multiplexed spatial transcriptomics in the larval forewing.** Representative fluorescence micrographs detailing multiple consecutive rounds of RAM-FISH hybridization, imaging, and signal removal on a single tissue sample. Rows indicate the specific imaging cycle (Cycles 1-8 and 10-12). Columns represent the distinct fluorophore channels used for detection (AF546, AF647, AF488, and AF514), followed by a merged composite overlay of all channels for that given cycle. Target genes mapped in each channel are labeled directly above their respective panels. Arrows between rows denote the intervening probe removal + reprobing steps, demonstrating the successful stripping and re-hybridization required for high-plex spatial mapping without signal carryover. Scale bar: 200 µm. Experiment performed using manual multiplexing in free-floating buffers.

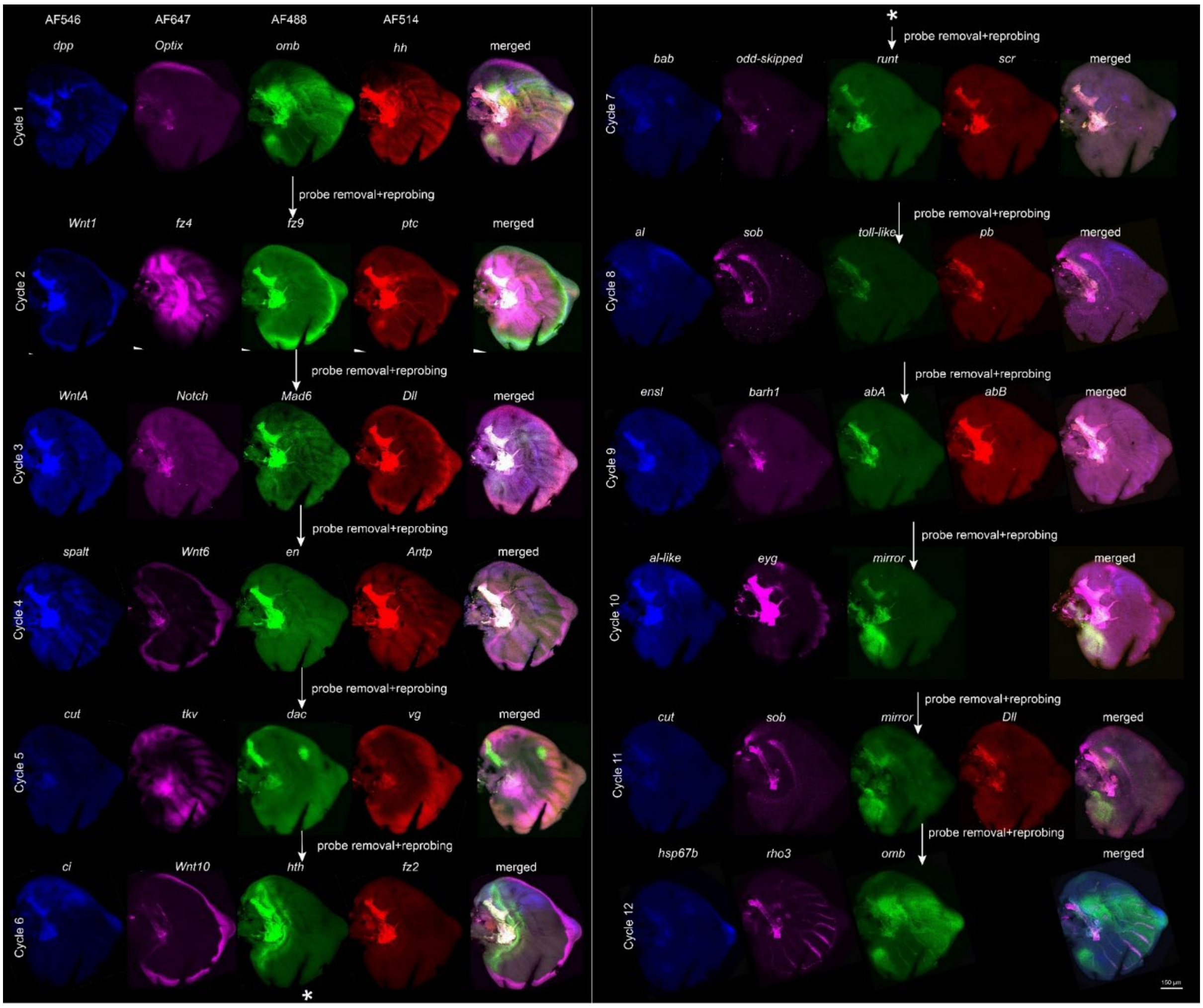


**Figure S5. Comprehensive 12-cycle sequential spatial transcriptomics in a single larval forewing.** Representative fluorescence micrographs detailing 12 consecutive rounds of RAM-FISH hybridization, imaging, and chemical signal stripping on the same tissue sample. Layout: The continuous sequence progresses through Cycles 1-6 (left block) and continues through Cycles 7-12 (right block), as denoted by the asterisk (*). Channels: Columns indicate the specific fluorophore channels (AF546, AF647, AF488, and AF514), with the targeted gene labeled directly above each panel. A merged composite overlay is provided for every cycle. Transitions: Arrows denoting probe removal + reprobing illustrate the intervening chemical stripping steps, demonstrating robust multiplexing capacity across 12 full rounds without morphological distortion or signal carryover. Scale bar (bottom right): 100 µm.

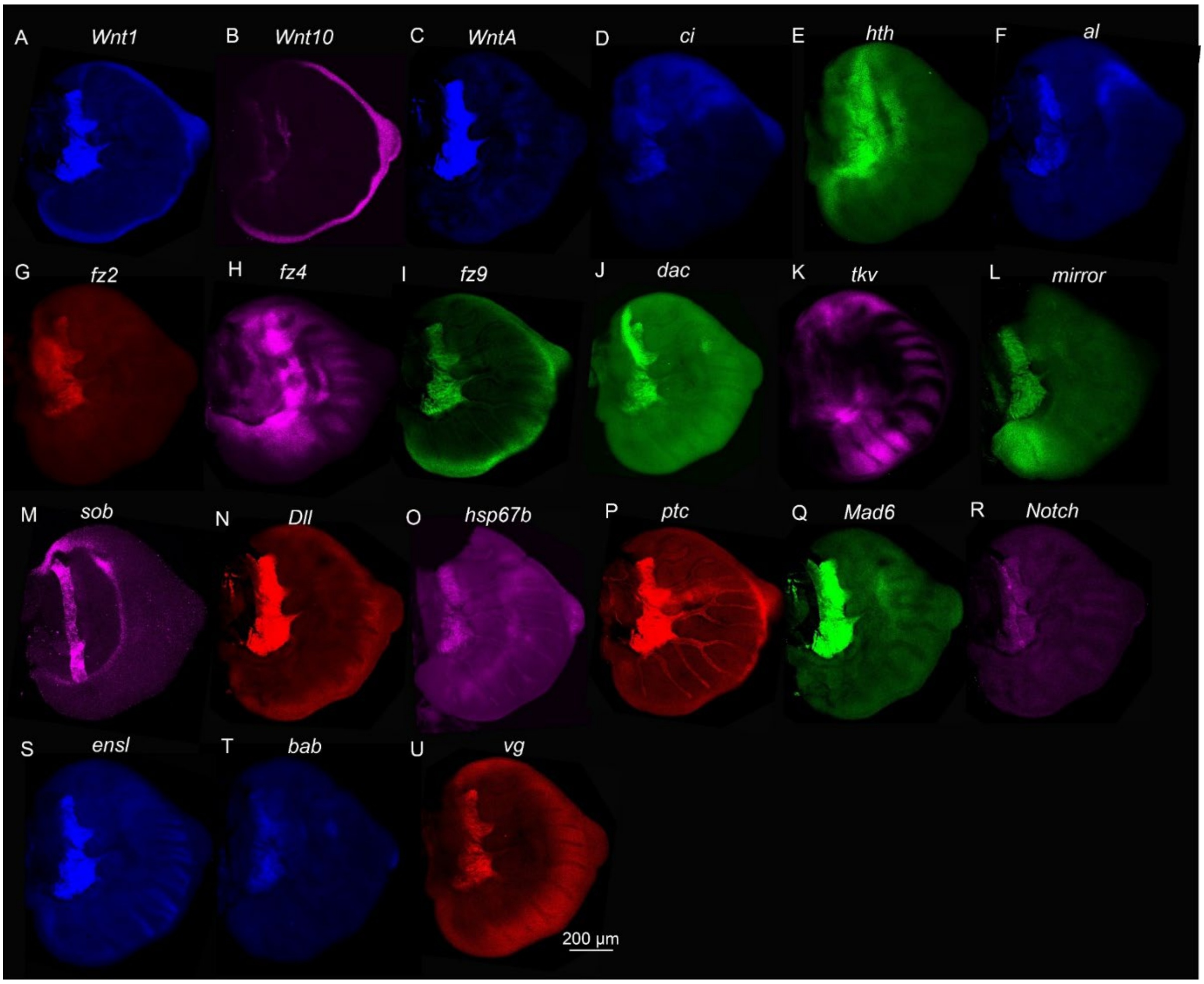


**Figure S6. Spatial expression mapping of multiplexed transcripts in the developing wing. (A-U)** Fluorescence micrographs displaying the spatial expression domains of 21 targeted genes detected via RAM-FISH. **(A)** *Wnt1,* **(B)** *Wnt10,* **(C)** *WntA,* **(D)** *ci,* **(E)** *hth,* **(F)** *al,* **(G)** *fz2,* **(H)** *fz4,* **(I)** *fz9,* **(J)** *dac,* **(K)** *tkv,* **(L)** *mirror,* **(M)** *sob,* **(N)** *Dll,* **(O)** *hsp67b,* **(P)** *ptc,* **(Q)** *Mad6,* **(R)** *Notch,* **(S)** *ensl,* **(T)** *bab,* and **(U)** *vg*. Experiment performed using manual multiplexing in free-floating buffers. Scale bar (in **U**): 200 µm.

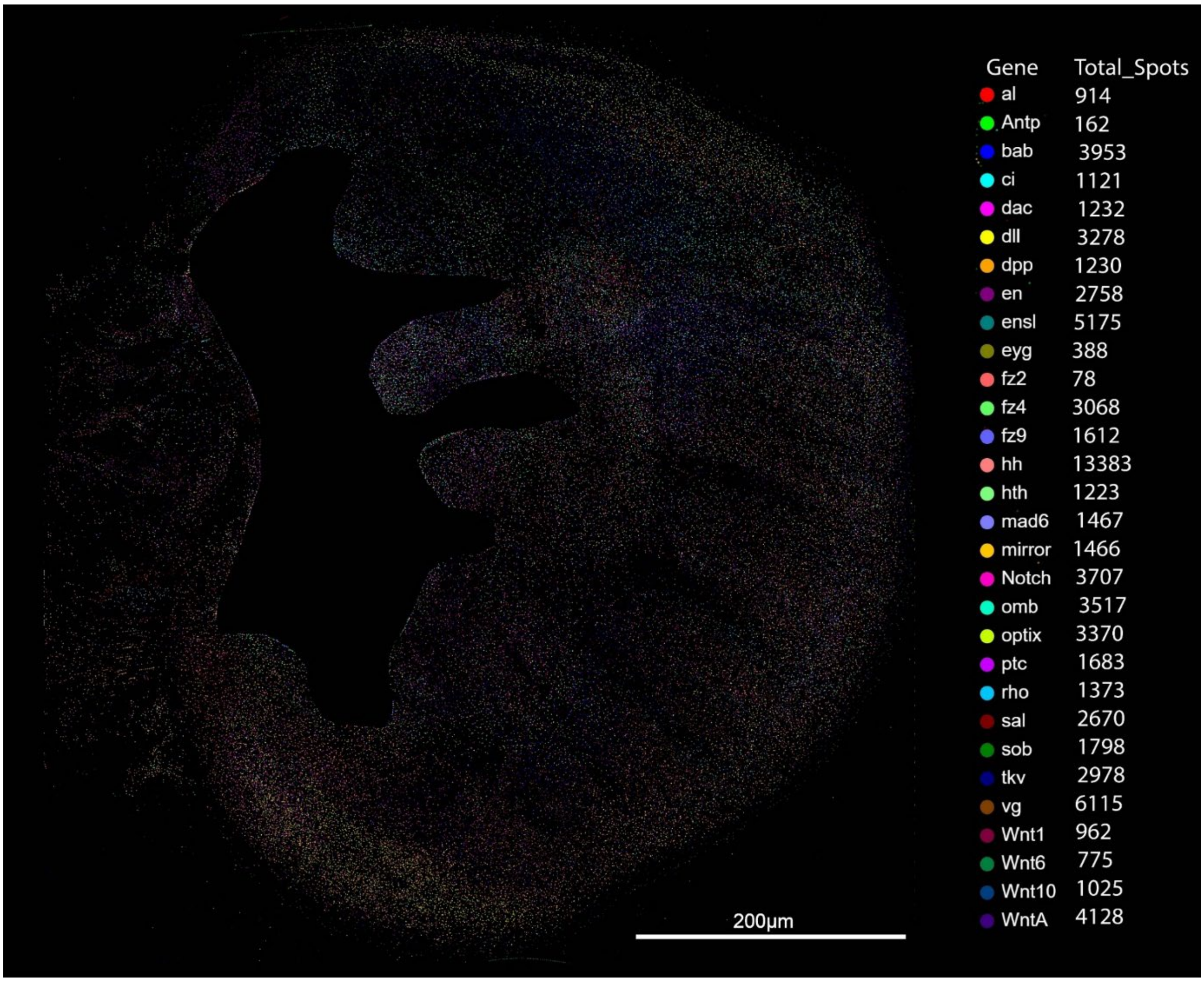


**Figure S7. Spatial mapping and quantification of multiplexed transcripts in the masked larval forewing from Figure 2.** Composite image displaying the merged spatial distribution of extracted transcript spots for all targeted genes, with a proximal tracheal tissue masked (black area). The accompanying l table provide total quantified spot counts for each respective gene across the mapped area.

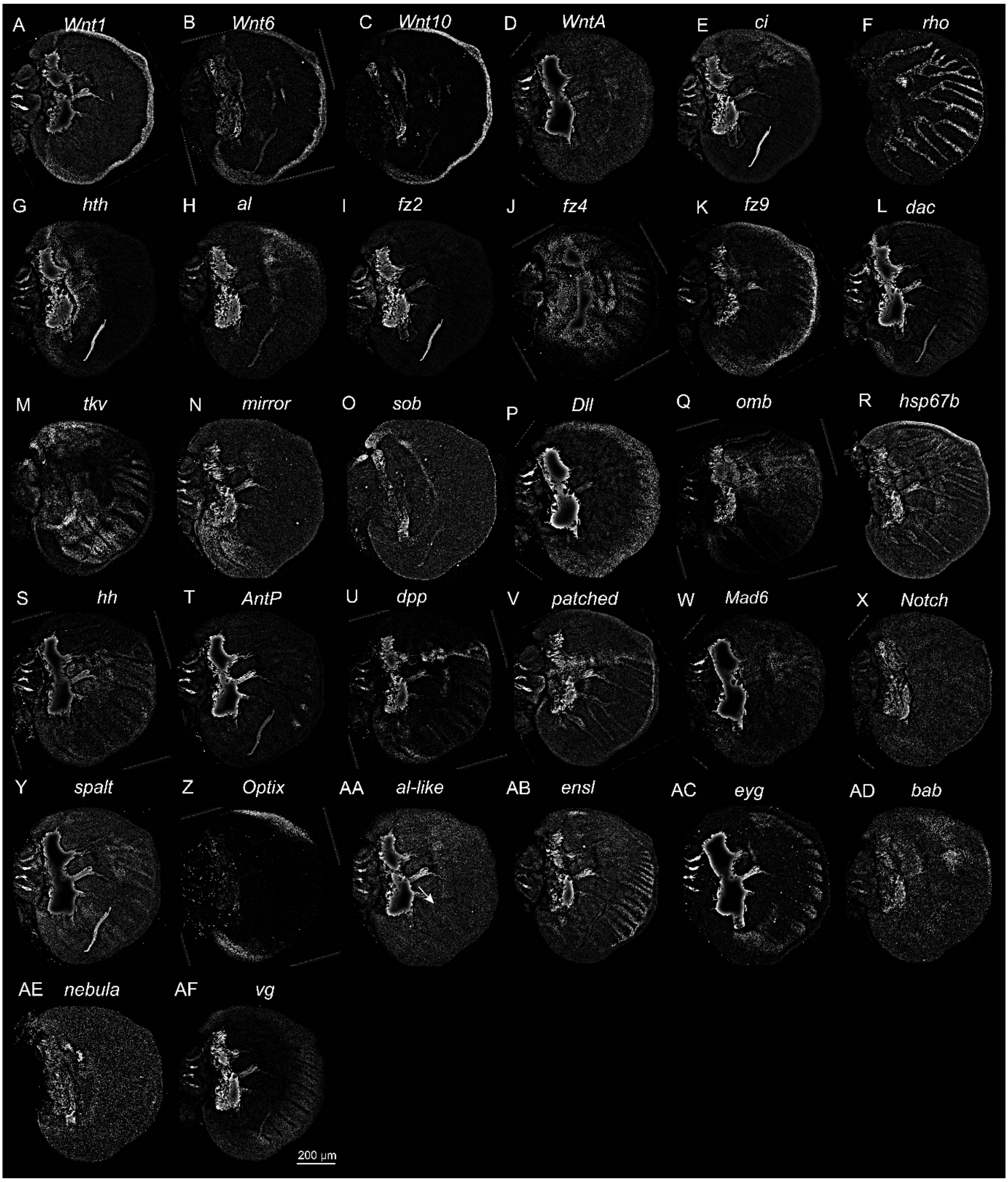


**Figure S8. Normalized gene expression using the ramfish_normalization script from the larval forewing raw files from Figure 2.** (**A**) *Wnt1*, (**B**) *Wnt6*, (**C**) *Wnt10*, (**D**) *WntA*, (**E**) *ci*, (**F**) *rho*, (**G**) *hth*, (**H**) *al*, (**I**) *fz2*, (**J**) *fz4*, (**K**) *fz9*, (**L**) *dac*, (**M**) *tkv*, (**N**) *mirror*, (**O**) *sob*, (**P**) *Dll*, (**Q**) *omb*, (**R**) *hsp67b*, (**S**) *hh*, (**T**) *AntP*, (**U**) *dpp*, (**V**) *patched*, (**W**) *Mad6*, (**X**) *Notch*, (**Y**) *spalt*, (**Z**) *Optix*, (**AA**) *al-like*, (**AB**) *ensl*, (**AC**) *eyg*, (**AD**) *bab*, (**AE**) *nebula*, and (**AF**) *vg*.

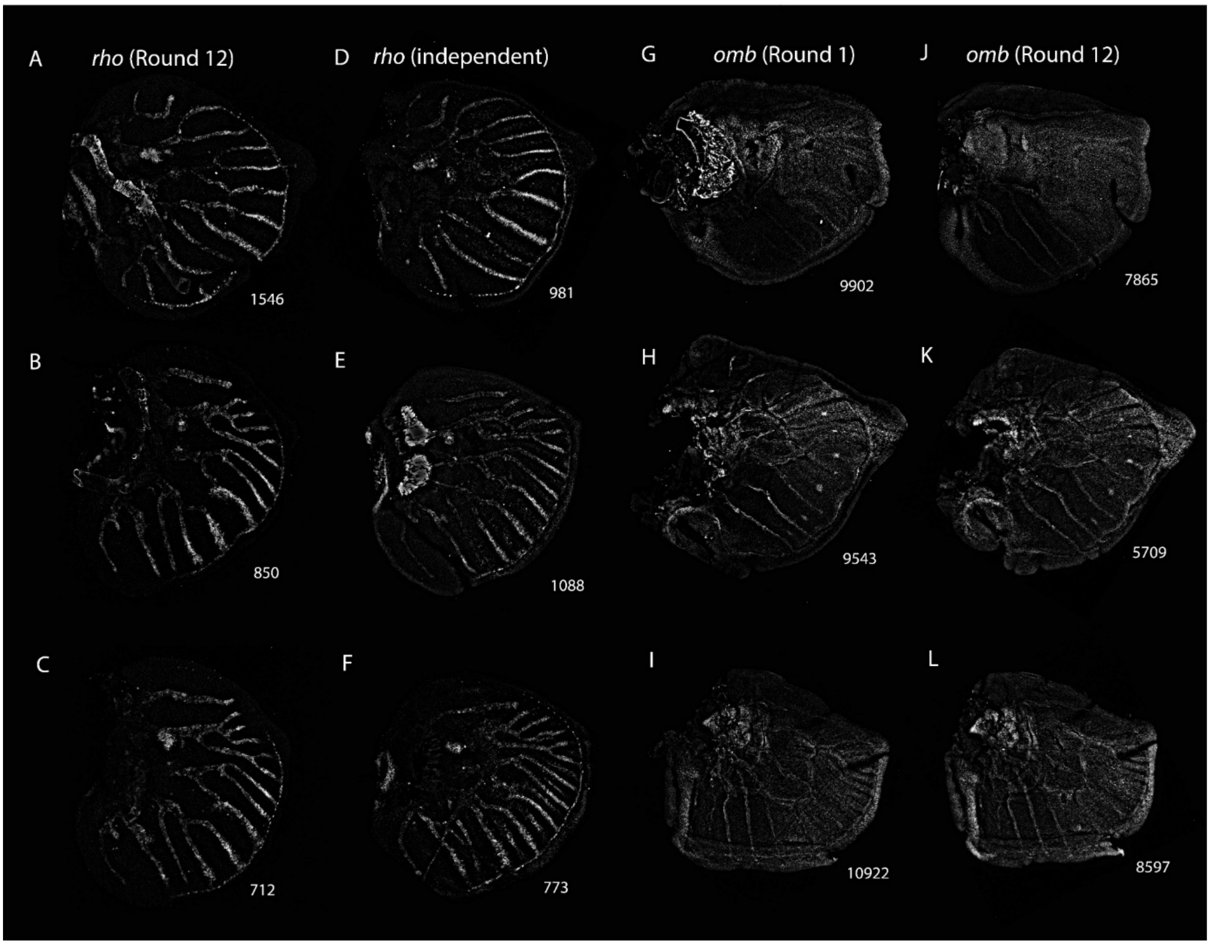


**Figure S9. Evaluation of imaging robustness and transcript quantification.** Robustness and reproducibility of signal detection for rho and omb across different imaging rounds and independent samples. (**A-C**) Detection of *rho* transcripts during Round 12 of imaging across three representative samples. (**D-F**) Detection of *rho* transcripts from independent experimental replicates. (**G-I**) Detection of *omb* transcripts during Round 1 of imaging across three representative samples. (**J-L**) Detection of *omb* transcripts from the same samples imaged during Round 12 to evaluate signal retention and decay. Numbers at the bottom-right corner of each panel indicate the total transcript spot counts quantified using the ramfish_spotcounter script.

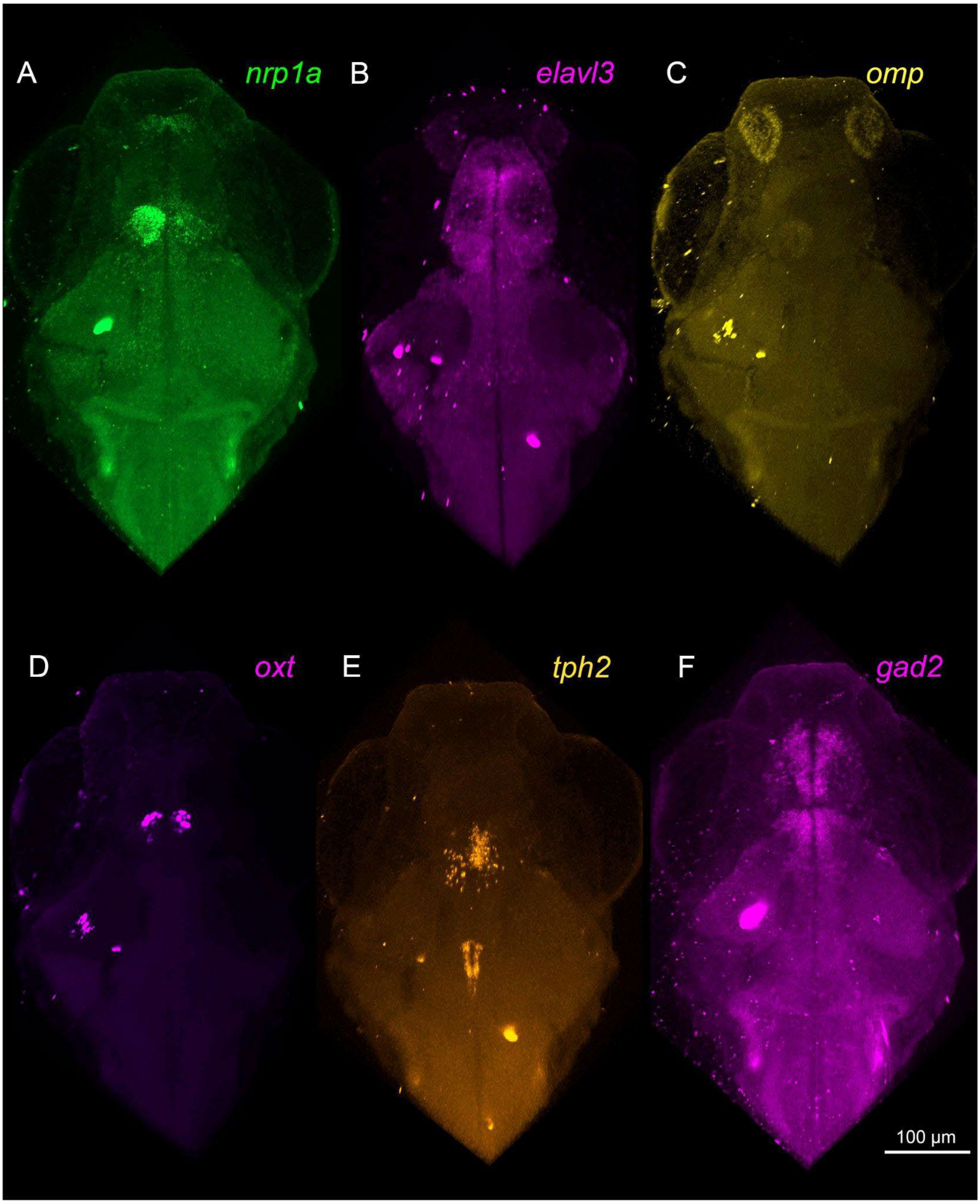


**Figure S10. Detection of multiple gene targets in a 14 dpf *Danio rerio* larva (sample 4). (A-I)** Multiplexed expression of *nrp1a*, *elavl3, omp, oxt, tph2,* and *gad2* in a 14 dpf *Danio rerio* larva. Experiment performed with the sample embedded in an acrylamide gel. Z-depth ~300 μm. Sample imaged using Zeiss LSM700.

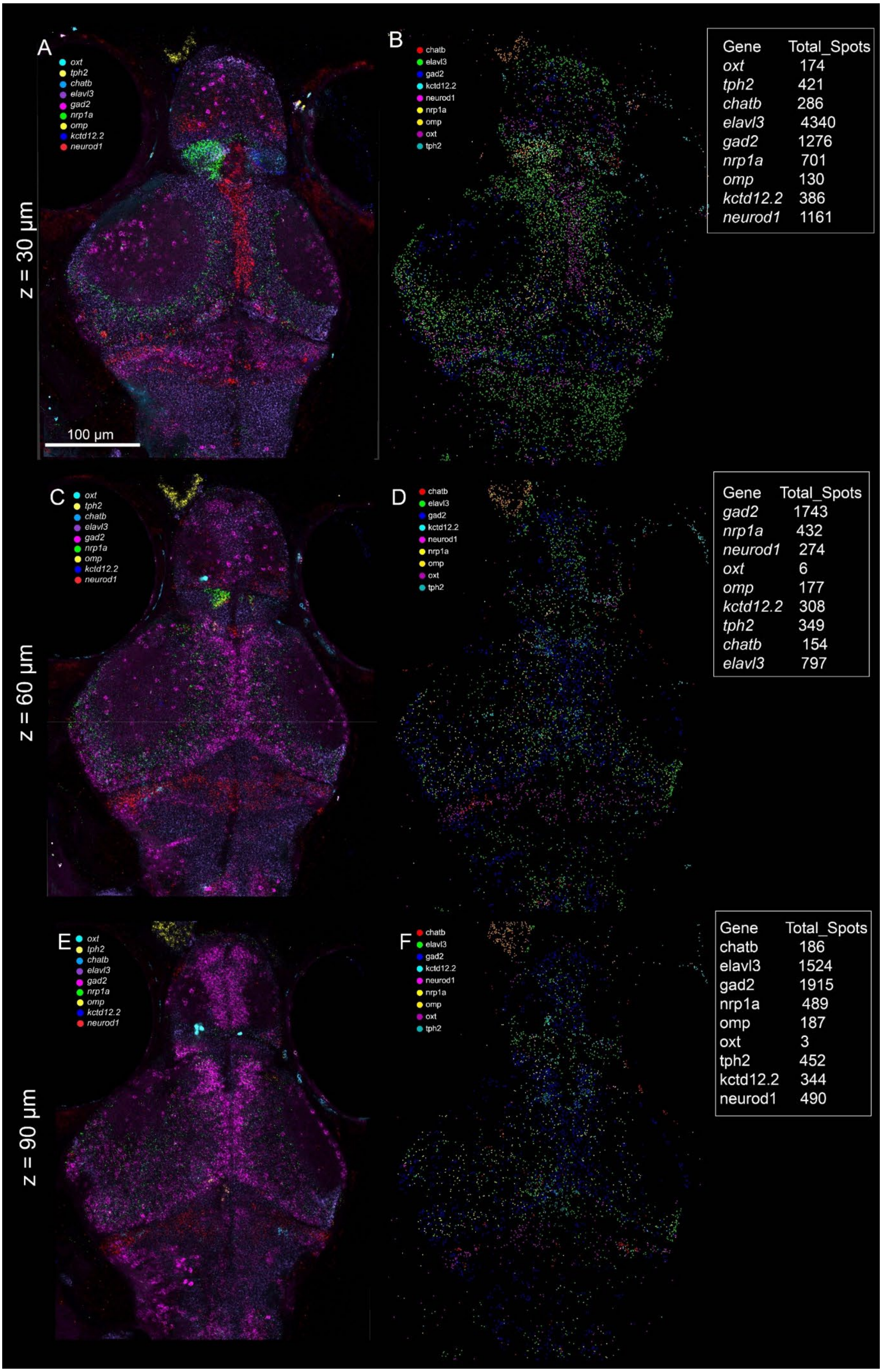
A
oxt
tph2
chatb
elavl3
gad2
nrp1a
omp
kctd12.2
neurod1
z = 30 µm
100 µm
B
chatb
elavl3
gad2
kctd12.2
neurod1
nrp1a
omp
oxt
tph2
Gene Total_Spots
oxt 174
tph2 421
chatb 286
elavl3 4340
gad2 1276
nrp1a 701
omp 130
kctd12.2 386
neurod1 1161
C
oxt
tph2
chatb
elavl3
gad2
nrp1a
omp
kctd12.2
neurod1
z = 60 µm
D
chatb
elavl3
gad2
kctd12.2
neurod1
nrp1a
omp
oxt
tph2
Gene Total_Spots
gad2 1743
nrp1a 432
neurod1 274
oxt 6
omp 177
kctd12.2 308
tph2 349
chatb 154
elavl3 797
E
oxt
tph2
chatb
elavl3
gad2
nrp1a
omp
kctd12.2
neurod1
z = 90 µm
F
chatb
elavl3
gad2
kctd12.2
neurod1
nrp1a
omp
oxt
tph2
Gene Total_Spots
chatb 186
elavl3 1524
gad2 1915
nrp1a 489
omp 187
oxt 3
tph2 452
kctd12.2 344
neurod1 490

**Figure S11: Depth-resolved multiplexed mRNA detection in a single 14 dpf Danio rerio larva.** (**A, C, E**) Raw fluorescence composite images illustrating the spatial expression of 9 target transcripts (*oxt, tph2, chatb, elavl3, gad2, nrp1a, omp, kctd12.2, neurod1*) at sequential optical depths (z = 30, 60, and 90 µm). (**B, D, F**) Corresponding synthetic spot maps generated via the RAM-FISH pipeline. Discrete transcript locations are color-coded, with the embedded quantitative tables detailing the absolute spot counts decoded for each gene at that specific focal plane. Scale bar in (**A**) applies to all panels.

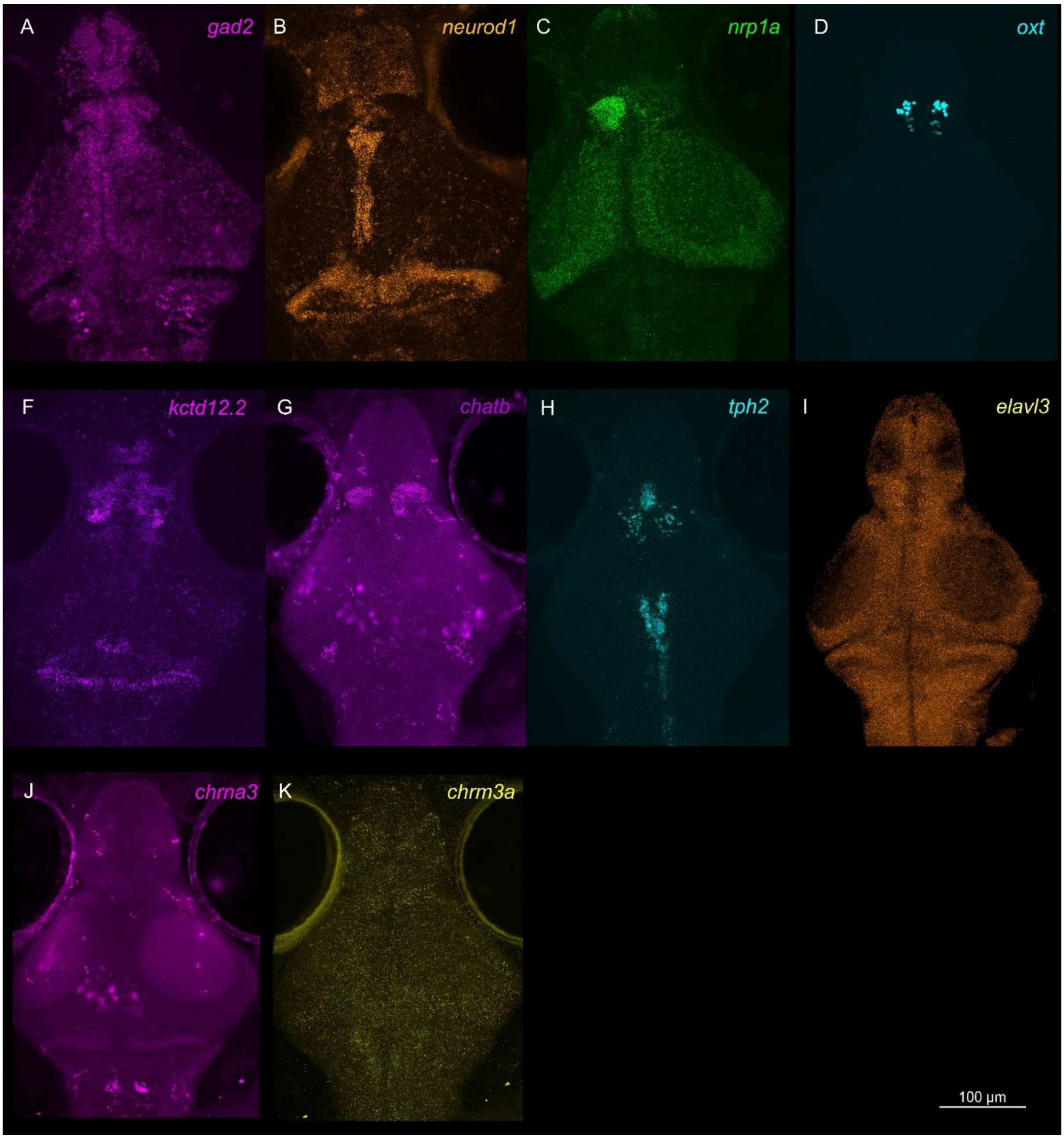


**Figure S12. Detection of multiple gene targets in a second 14 dpf *Danio rerio* larva across an axial depth of ~200 µm (sample 2). (A-I)** Multiplexed expression of *gad2, nrp1a, neuroD, oxt, kctd12.2, chatb, tph2, elavl3, chrna3,* and *chrm3a* in a 14 dpf *Danio rerio* larva. Reactions were carried out in the gel embedding format.

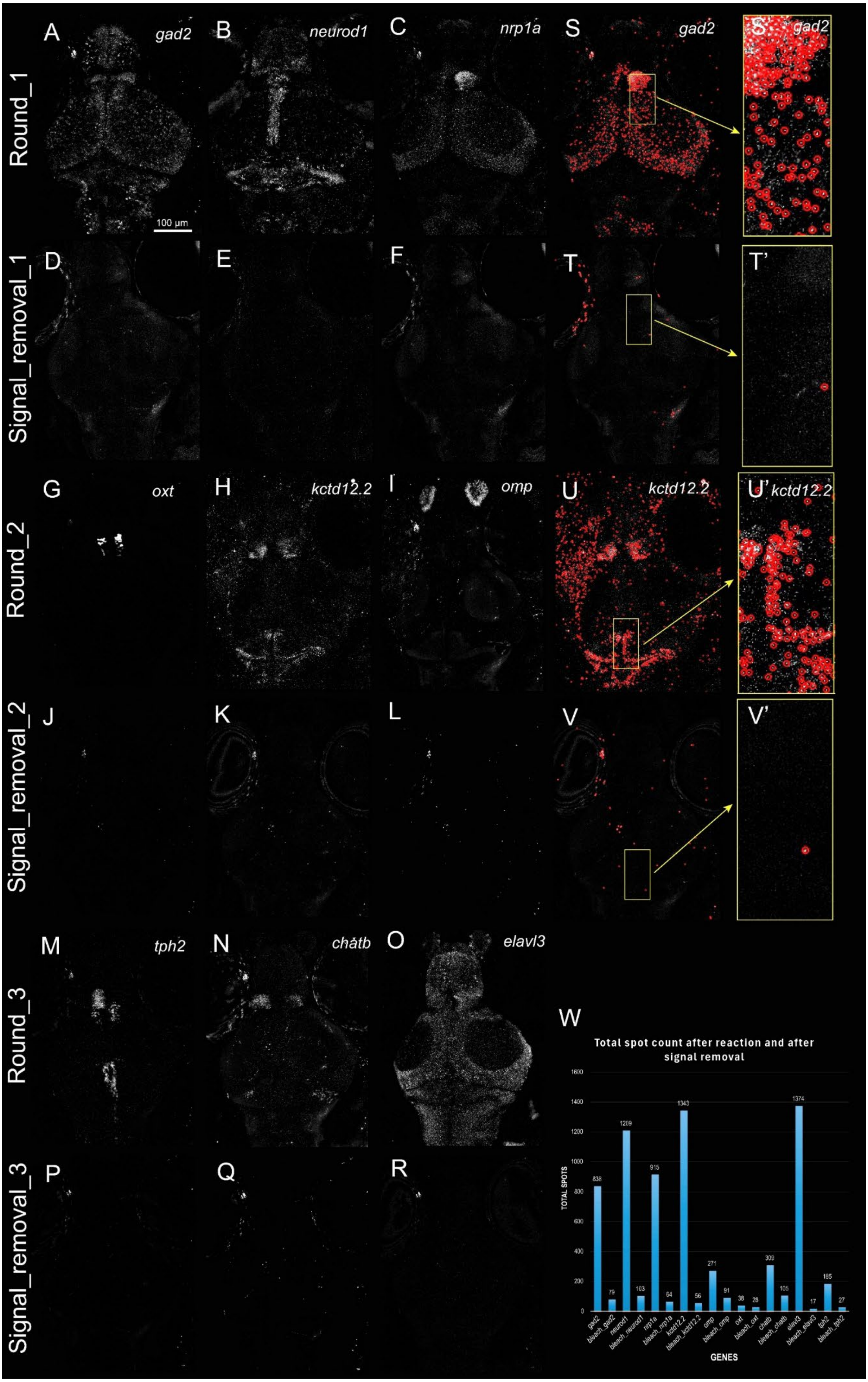
Round_1
Signal_removal_1
Round_2
Signal_removal_2
Round_3
Signal_removal_3
A gad2
B neurod1
C nrp1a
S gad2
S' gad2
100 µm
D
E
F
T
T'
G oxt
H kctd12.2
I omp
U kctd12.2
U' kctd12.2
J
K
L
V
V'
M tph2
N chatb
O elavl3
P
Q
R
W
Total spot count after reaction and after signal removal
TOTAL SPOTS
GENES
838
79
1209
103
915
64
1343
56
271
91
38
28
309
105
1374
17
185
27
gad2
bleach_gad2
neurod1
bleach_neurod1
nrp1a
bleach_nrp1a
kctd12.2
bleach_kctd12.2
omp
bleach_omp
oxt
bleach_oxt
chatb
bleach_chatb
elavl3
bleach_elavl3
tph2
bleach_tph2

**Figure S13: Validation of multiplexed chemical signal stripping and transcript removal efficiency across successive RAM-FISH hybridization cycles (sample 1).** (**A-C**) Raw fluorescence micrographs of target transcripts (*gad2, neurod1*, and *nrp1a*) detected during Round 1 of hybridization. Scale bar in panel A applies to all grayscale image panels. (**D-F**) Corresponding fluorescence imaging channels following the first Signal_removal_1 cycle, demonstrating near-complete erasure of the primary fluorescent signal before subsequent chemical steps. (**G-I**) Targeted detection of a secondary subset of transcripts (*oxt*, *kctd12.2*, and *omp*) during Round 2 of hybridization, verifying robust probe re-hybridization dynamic range without signal carryover. (**J-L**) Imaging verification channels following the second Signal_removal_2 cycle showing uniform transcript signal stripping across the region of interest. (**M-O**) Detection profiles for the final transcript cohort (*tph2, chatb*, and *elavl3*) during Round 3 of hybridization, demonstrating excellent preservation of tissue morphology and RNA integrity after multiple stripping rounds. (**P-R**) Residual fluorescence verification checks following the third Signal_removal_3 steps. (**S-V**) Algorithmic verification panels detailing spatial spot-calling efficiency before and after chemical processing: (**S-S'**) Active spot-calling profile for the *gad2* channel during Round 1, with the magnified inset (yellow box) showing high-density transcripts successfully bounded by red validation rings. (**T-T'**) Residual spot-calling check post-stripping (Signal_removal_1), confirming complete physical drop-off of localized transcripts within the identical region of interest. (**U-U'**) Active spot-calling profile for *kctd12.2* during Round 2, highlighting robust multi-cycle target identification within the target domain. (**V-V'**) Automated verification post-stripping (Signal_removal_2), illustrating efficient removal of localized fluorescence features. (W) Quantitative evaluation chart contrasting the total spot counts detected immediately after hybridization cycles against residual trace counts registered following subsequent chemical signal removal steps for each targeted gene.

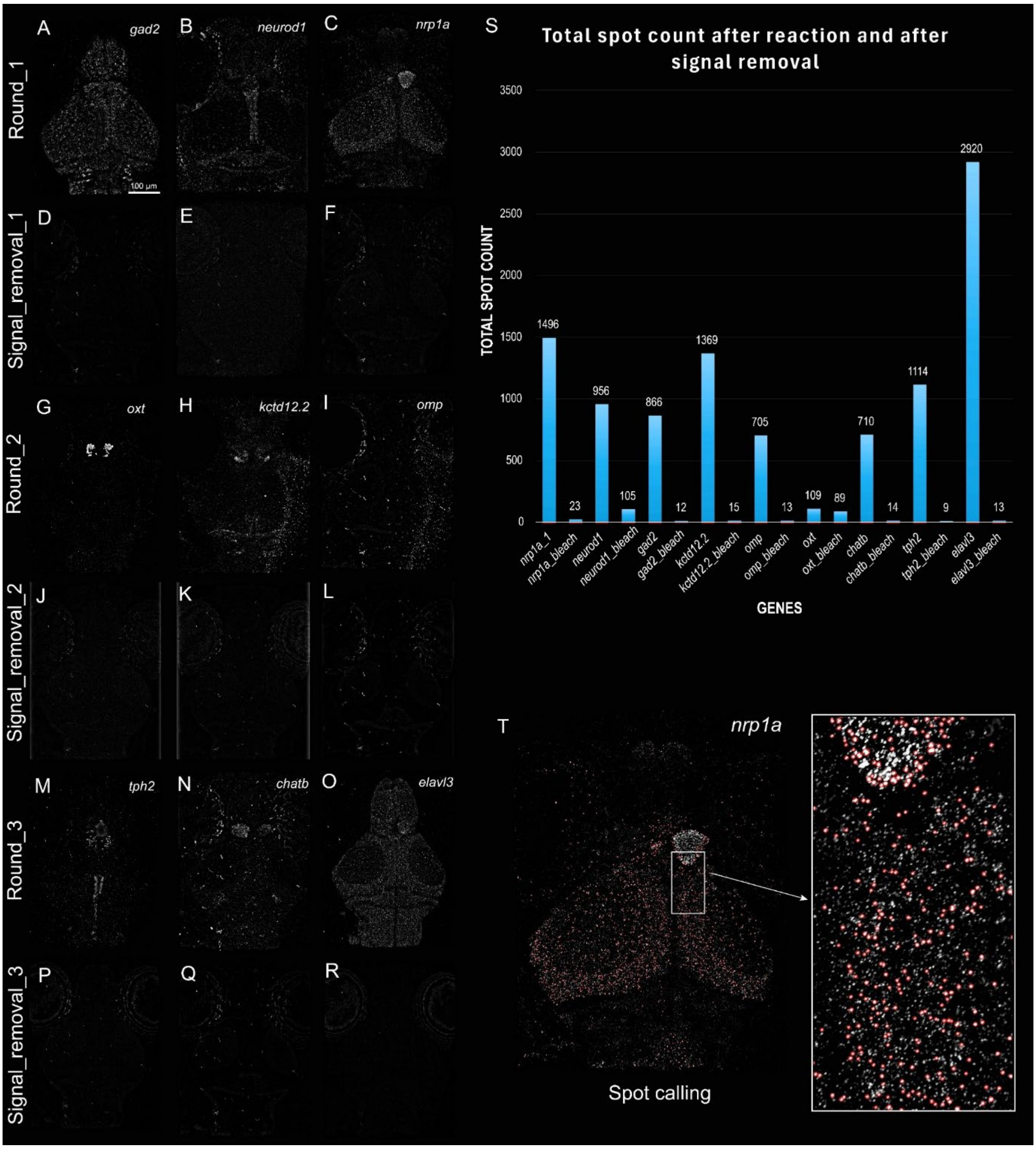


**Figure S14: Validation of multiplexed chemical signal stripping and transcript removal efficiency across successive RAMFISH hybridization cycles (sample 3).** (**A-C**) Raw fluorescence micrographs of target transcripts (*gad2, neurod1*, and *nrp1a*) detected during Round 1 of hybridization. Scale bar in panel A applies to all grayscale image panels. (**D-F**) Corresponding fluorescence imaging channels following the Signal_removal_1 cycle, demonstrating complete erasure of the primary fluorescent signal before subsequent fluidic steps. (**G-I**) Targeted detection of a secondary subset of transcripts (*oxt, kctd12.2*, and *omp*) during Round 2 of hybridization, verifying robust probe re-hybridization dynamic range without signal carryover. (**J-L**) Imaging verification channels following the Signal_removal_2 cycle showing uniform transcript signal stripping across the region of interest. (**M-O**) Detection profiles for the final transcript cohort (*tph2, chatb*, and *elavl3*) during Round 3 of hybridization, demonstrating preservation of tissue morphology and RNA integrity after multiple stripping rounds. (**P-R**) Residual fluorescence verification checks following the Signal_removal_3 step.

(**S**) Quantitative evaluation chart contrasting the total spot counts detected immediately after hybridization cycles against residual trace counts registered following subsequent chemical signal removal steps for each targeted gene. (**T**) Active spot-calling profile for the *nrp1a* channel during Round 1, with the magnified inset (white box) showing high-density transcripts successfully bounded by red validation rings.

#### Mapzebrain

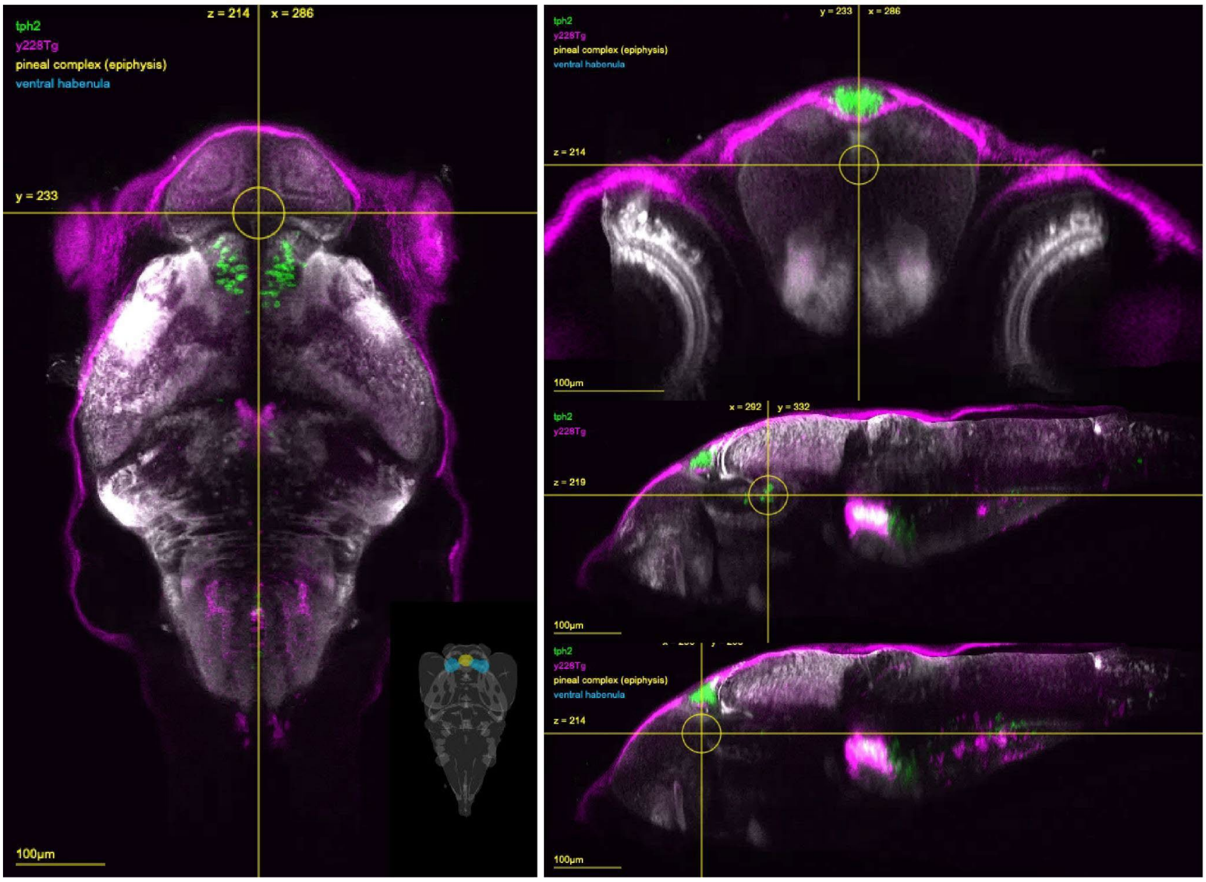


**Figure S15. Expression of *tph2* (green) and y228Tg [tph2:Gal4, Tg(tph2:Gal4FF)y228] (magenta) from mapZbrain.** Note the broader expression of the y228Tg compared to the *tph2* HCR mRNA expression.

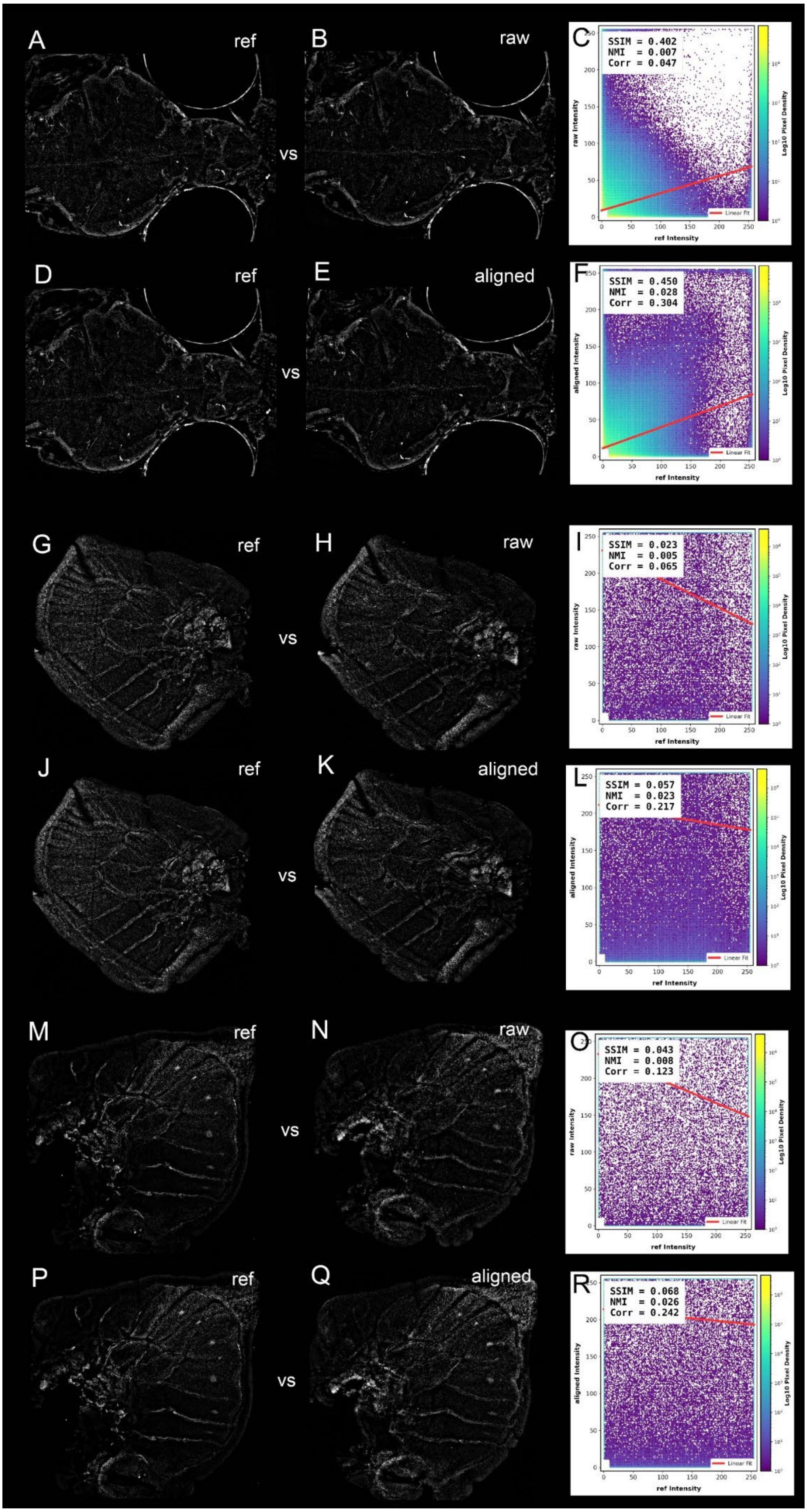
A
ref
B
raw
vs
C
SSIM = 0.402
NMI = 0.007
Corr = 0.047
raw Intensity
ref Intensity
Log10 Pixel Density
Linear Fit
D
ref
E
aligned
vs
F
SSIM = 0.450
NMI = 0.028
Corr = 0.304
aligned Intensity
ref Intensity
Log10 Pixel Density
Linear Fit
G
ref
H
raw
vs
I
SSIM = 0.023
NMI = 0.005
Corr = 0.065
raw Intensity
ref Intensity
Log10 Pixel Density
Linear Fit
J
ref
K
aligned
vs
L
SSIM = 0.057
NMI = 0.023
Corr = 0.217
aligned Intensity
ref Intensity
Log10 Pixel Density
Linear Fit
M
ref
N
raw
vs
O
SSIM = 0.043
NMI = 0.008
Corr = 0.123
raw Intensity
ref Intensity
Log10 Pixel Density
Linear Fit
P
ref
Q
aligned
vs
R
SSIM = 0.068
NMI = 0.026
Corr = 0.242
aligned Intensity
ref Intensity
Log10 Pixel Density
Linear Fit

**Figure S16. Computational correction of spatial drift across multiplexed hybridization cycles via the ramfish_fine_aligner.** In the images “ref” indicates a reference image used as to align the next rounds of images. First rough alignment is carried out on the subsequent rounds of images using the ramfish_rough_aligner (indicated as “raw”) followed by alignment using ramfish_fine_aligner (indicated as “aligned”). The graphs show the improvement of the alignment via the ramfish_fine_aligner using SSIM, NMI, and R metrics. (**A-F**) Zebrafish 14 dpf larval DAPI stainings. (**G-L**) *Bicyclus anynana* larval forewing *omb* staining. (**M-R**) *Bicyclus anynana* larval hindwing *omb* staining.

Note: The Pearson correlation coefficient (R) establishes baseline spatial alignment by evaluating the global linear relationship of pixel intensities between hybridization cycles. To verify that computational registration successfully corrected physical tissue warping without introducing distortion, the Structural Similarity Index Measure (SSIM) (Larkin, 2015; Wang et al., 2004) was calculated to assess local spatial cohesion and architectural integrity. Finally, Normalized Mutual Information (NMI) (Pluim et al., 2003; Studholme et al., 1999) robustly quantifies alignment accuracy independent of the non-linear fluorescence fluctuations and background changes inevitably induced by iterative chemical stripping.

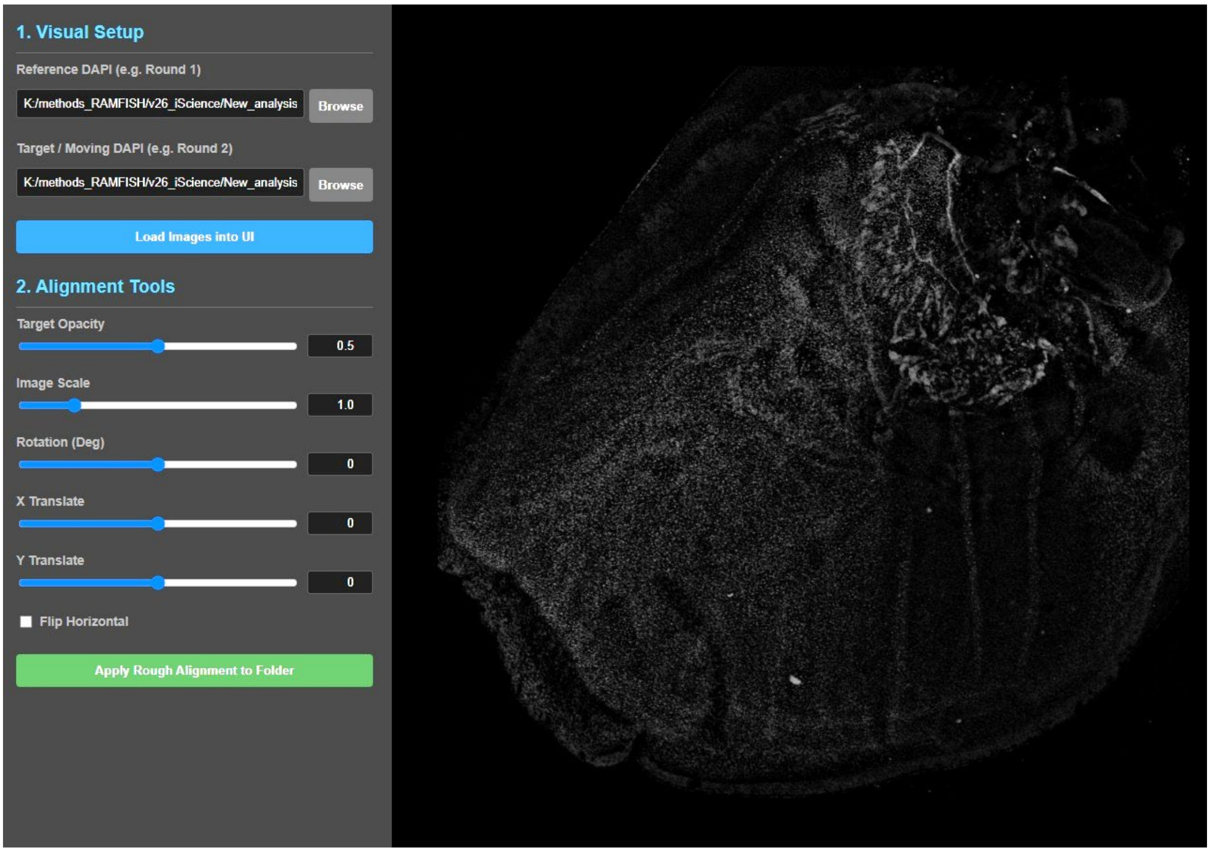


**Figure S17. The ramfish_rough_aligner interactive interface.** This web-based tool allows for the manual spatial alignment of tissue samples that have shifted, rotated, or inverted between sequential hybridization cycles. The central canvas displays a real-time overlay of the fixed Round 1 structural reference and the moving target image. Using the left-hand control panel, users adjust target opacity, scale, rotation, translation, and horizontal orientation to visually align the tissues. Clicking "Apply Rough Alignment to Folder" converts these manual adjustments into an affine transformation matrix. This matrix is then applied in batch to all associated gene channels within the target directory, maintaining native image resolution.

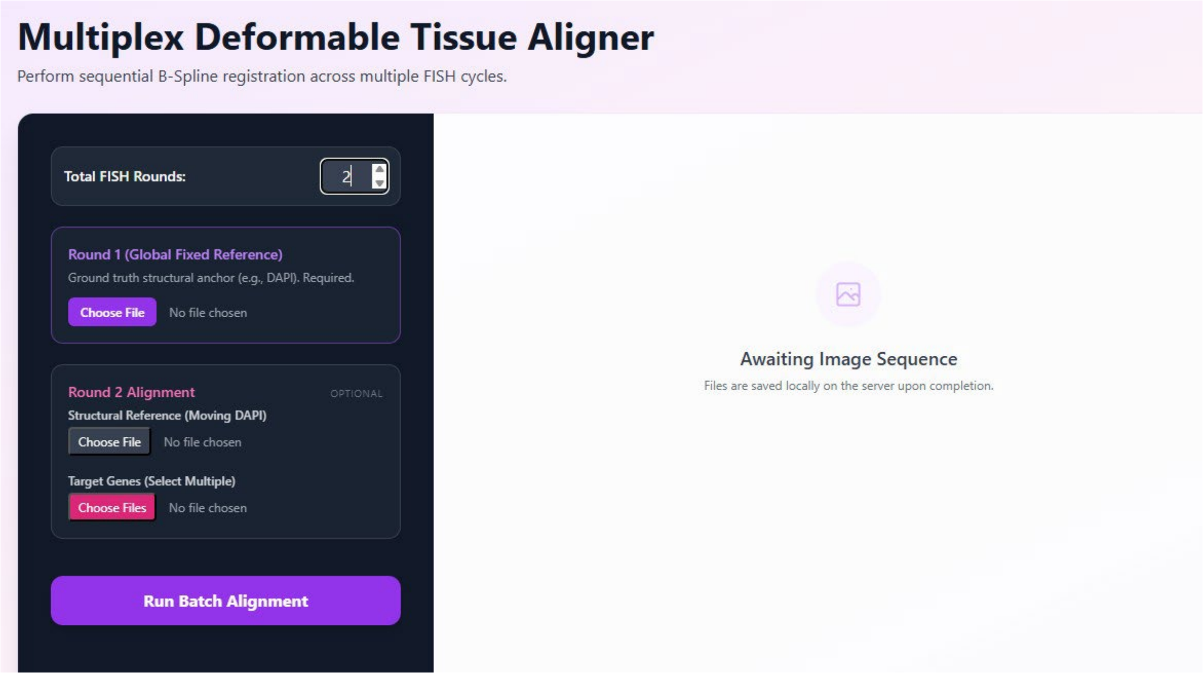


**Figure S18. The ramfish_fine_aligner interface.** This web-based interface manages the automated registration of multiplexed datasets following the initial rough alignment step. Users define the total number of hybridization cycles and establish the global coordinate system by uploading the Round 1 structural reference (e.g., DAPI). For subsequent cycles, the moving structural reference is paired with its corresponding batch of target gene channels. Selecting "Run Batch Alignment" initiates a Python-based dual-stage registration (an affine transformation followed by non-rigid B-Spline elastic deformation) that aligns the structural anchors and warps all linked target genes to the Round 1 coordinate space. The terminal console tracks registration metrics and processing status in real time.

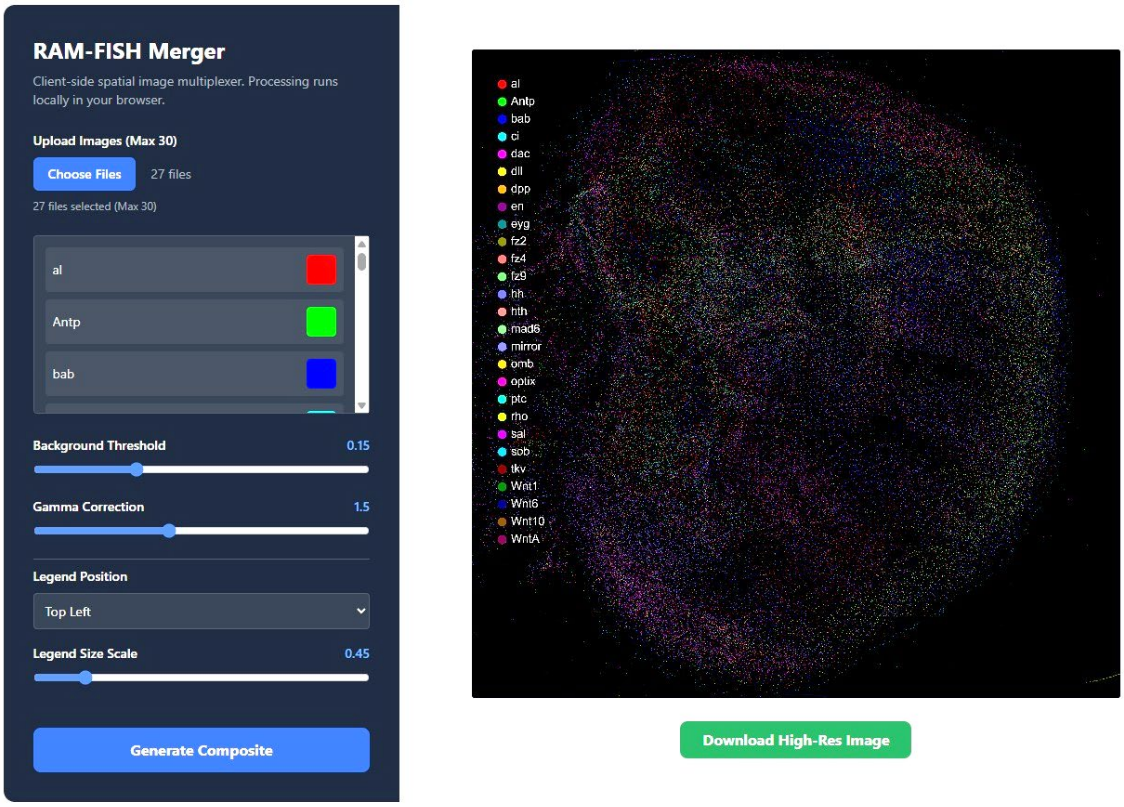

**Figure S19. The ramfish_merger interface.** This utility combines individual transcript spatial maps into a single composite image. Users upload their target gene channels (e.g., *ci, ptc, hh, omb*) along with an optional global structural reference (e.g., DAPI). The application renders the structural reference as a grayscale background and assigns distinct colors to each transcript layer. The left-hand panel provides global adjustments for gamma correction and background noise thresholds to improve the visibility of dense expression domains. The interface also includes annotation tools for automated legend placement and scaling. Selecting "Generate Composite" produces the final overlaid multiplexed-FISH spatial map. The resulting file is saved using the "Download High-Res Image" button.

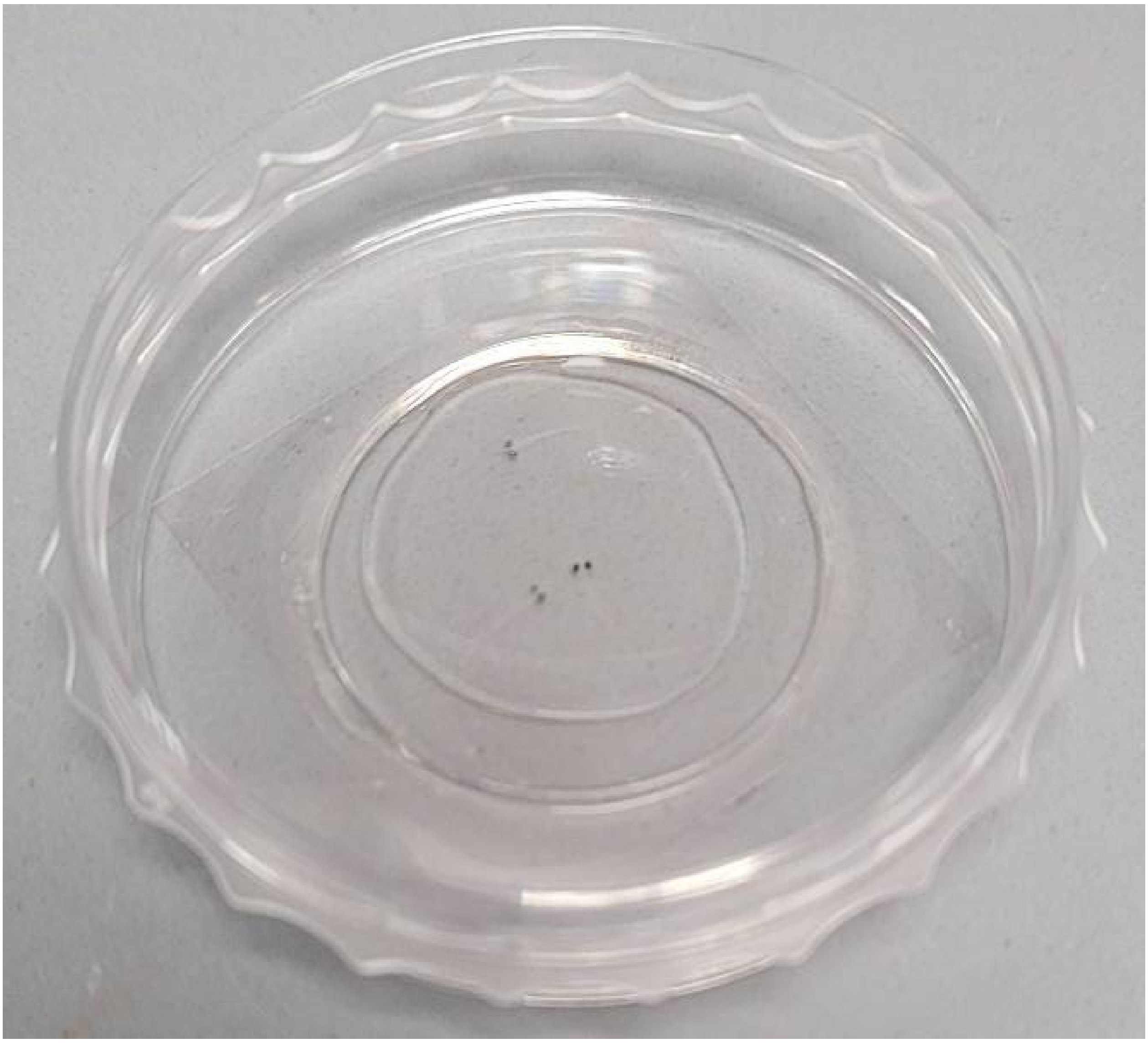

**Figure S20:** 14 dpf zebrafish larvae embedded in acrylamide gel on a confocal microscopy friendly dish.

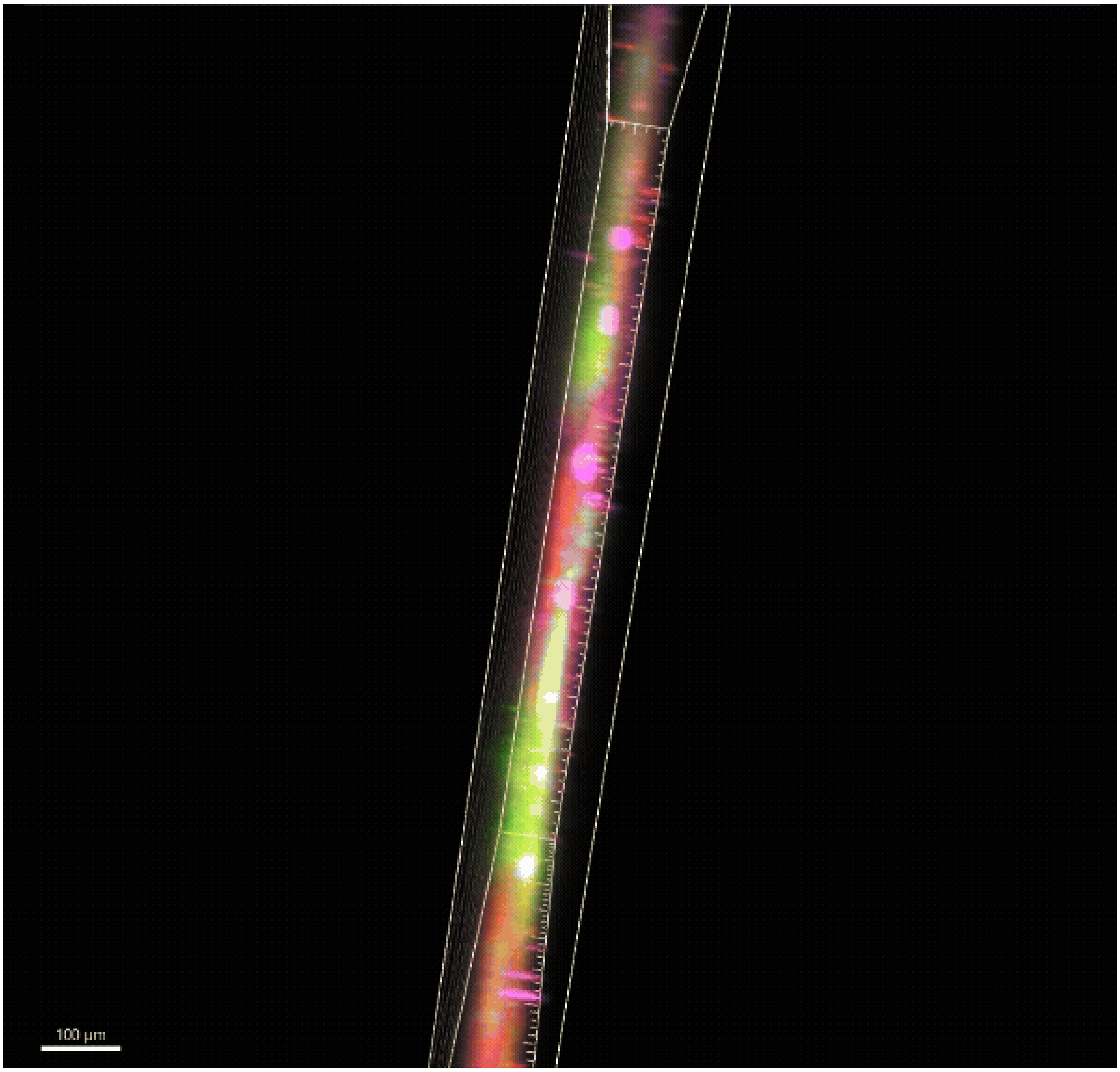


**Video S1. A volume-rendered mid larval *Bicyclus anynana* hindwing stained with *omb* (green), *frizzled4* (red), and *spalt* (magenta) across an axial depth of ~120 µm.** Experiment performed using manual multiplexing with the samples in free-floating buffers.

**Table S1: Genes examined in *Bicyclus anynana* wings**

| **Sl. No.** | **Gene name** | **Expression pattern** | **Species tested** | **Functional testing in butterflies** |
|---|---|---|---|---|
| 1 | *Wnt1* | *Wnt1* was expressed along the margin and along the discal s overlapping the expression pattern *Wnt10* and *Wnt6*, consistent previous findings (**Figure 2A, Figure 3A,** | *Junonia coenia, Euphydryas chalcedona, Agraulis vanillae, Vanessa cardui,* and *Bicyclus* | *Wnt1* has been functionally tested via RNAi where the study found Wnt1 is involved in the size of |

| | | | | |
|---|---|---|---|---|
| | | **Figure S1 to S6**). | *anynana* (Banerjee and Monteiro, 2020; Banerjee et al., 2023; Martin and Reed, 2014). | eyespots in *Bicyclus anynana* (Özsu et al., 2017). |
| 2 | *Wnt6* | *Wnt6* was expressed along the wing margin and in the discal spots of butterflies, consistent with previous findings (Banerjee et al., 2023; Martin and Reed, 2014) (**Figure 2B, Figure 3B, Figure S1 to S6**). | *Junonia coenia, Euphydryas chalcedona, Agraulis vanillae, Vanessa cardui*, and *Bicyclus anynana* (Banerjee and Monteiro, 2020; Banerjee et al., 2023; Martin and Reed, 2014). | Not tested. |
| 3 | *Wnt10* | *Wnt10* was observed along the wing margin as well as in the discal spot, consistent with previous findings (Banerjee et al., 2023; Martin and Reed, 2014) (**Figure 2C, Figure S1 to S6**). | *Junonia coenia, Euphydryas chalcedona, Agraulis vanillae, Vanessa cardui*, and *Bicyclus anynana* (Banerjee and Monteiro, 2020; Banerjee et al., 2023; Martin and Reed, 2014). | Not tested. |
| 4 | *WntA* | *WntA* was expressed in bands along the central symmetry system and along the marginal band system consistent with previous studies (Banerjee et al., 2023; Hanly et al., 2023; Martin and Reed, 2014; Mazo-vargas et al., 2017) (**Figure 2D, Figure 3C, Figure S1 to S6**). | *Junonia coenia, Euphydryas chalcedona, Agraulis vanillae, Vanessa cardui,* and *Bicyclus anynana* (Banerjee et al., 2023; Hanly et al., 2023; Martin and Reed, 2014; | *WntA* has been functionally tested in multiple butterfly species, including *Junonia coenia, Pararge aegeria, Vanessa cardui, Heliconius erato, Heliconius sara, Agraulis vanillae*, and *Bicyclus anynana*. Studies showed |

| | | | | |
|---|---|---|---|---|
| | | | Mazo-vargas et al., 2017). | that WntA controls multiple wing pattern elements (Banerjee et al., 2023; Hanly et al., 2023; Mazo-vargas et al., 2017). |
| 5 | *ci* | *cubitus interruptus (ci)* was expressed in the anterior compartment of the developing wing discs and in the eyespot centers, consistent with previous studies (Banerjee and Monteiro, 2023; Keys et al., 1999) (**Figure 2E, Figure S1 to S6**). We noticed a strong expression of *ci* in M1 and a fainter expression in M2 eyespots, in hindwings at stage 2.5, whereas the remaining eyespots did not express *Ci* (**Figure S2E)**. This absence of *Ci* expression in eyespot centers positioned away from the anterior-posterior (AP) boundary of the wing is consistent with the finding from Reed et al., 2020. | *Junonia coenia* (Keys et al., 1999) and *Bicyclus anynana* (Banerjee and Monteiro, 2023; Keys et al., 1999; Reed et al., 2020) | Not tested. |
| 6 | *fz2* | *frizzled2 (fz2)* was expressed in the proximal domain of the developing larval wing, consistent with previous results in other butterfly species (Banerjee et al., 2023; Hanly et al., 2023) (**Figure 2I, Figure S1 to S6**). | *Vanessa cardui, Agraulis incarnata* and *Bicyclus anynana* (Banerjee et al., 2023; Hanly et al., 2023). | *fz2* plays a role in receiving WntA ligand expressed in the Central Symmetry System (CSS). The functional role has been studied in *Vanessa cardui, Junonia coenia, Agraulis incarnata*, and *Bicyclus anynana* (Banerjee et al., 2023; Hanly et al., 2023). |

| | | | | |
|---|---|---|---|---|
| 7 | *fz4* | *frizzled4 (fz4)* was expressed in the intervein cells during the early stages of development (stage 0.75-1.00) (**Figure 2J, Figure 3E, Figure S1 to S6**). During the later stages (2.50-3.25), *fz4* expression was observed in the intervein cells along with missing expression in the eyespot center (**Figure S1 to S6).** These results are consistent with results from previous findings (Banerjee et al., 2023). | *Vanessa cardui, Bicyclus anynana* (Banerjee et al., 2023; Hanly et al., 2023). | *fz4* plays a role in eyespot center formation and in the orientation of scale cells. The role has been studied in *Vanessa cardui, Junonia coenia, Agraulis incarnata*, and *Bicyclus anynana* (Hanly et al., 2023; Banerjee et al., 2023). |
| 8 | *fz9/fz3* | *frizzled9/frizzled3 (fz9)* was expressed along the wing margin, consistent with previous findings (Banerjee et al., 2023; Hanly et al., 2023) (**Figure 2K, Figure 3F, Figure S1 to S6**). | *Vanessa cardui, Bicyclus anynana* (Banerjee et al., 2023; Hanly et al., 2023). | *fz9/fz3* plays a role in the differentiation of veins, wing margin color patterns and discal eyespots/bands. Knockout of this gene results in ectopic veins, altered discal eyespots, and deformed wing margin (Hanly et al., 2023). |
| 9 | *tkv* | *thickvein (tkv)* was expressed around the intervein cells with stronger expression in cells distant from *dpp* expressing cells, consistent with previous findings in butterflies (Banerjee and Monteiro, 2023) and *Drosophila* (Funakoshi et al., 2001) (**Figure 2M, Figure 3H, Figure S1 to S6**). | *Bicyclus anynana* (Banerjee and Monteiro, 2023) | Not tested. |

| | | | | |
|---|---|---|---|---|
| 10 | *Dll* | *Distal-less (Dll)* was expressed along the wing margin with finger like projections during early wing development (Stages 0.75-1.00) (**Figure 2P, Figure 3K, Figure S1 to S6**). During the later stages, *Dll* expression was observed along the wing margin and in the eyespot centers (**Figure S2L).** These results are consistent with previous findings (Carroll et al. 1994; Brakefield et al. 1996; Banerjee et al., 2023; Reed and Gilbert, 2004; Reed et al., 2020; Shirai et al. 2012, Oliver et al. 2012; Wee et al., 2022). | *Junonia coenia, Vanessa cardui, Bicyclus anynana, Tithorea tarricina, Danaus Plexippus, Morpho peleides forewings, Caligo Memnon, Consul fabius, Hypna clytemenstra, Dryadula phaetusa, Hamadryas amphinome, Hamadryas februa, Catonephele numilia, Nessaea aglaura, Myscelia cyaniris, Vanessa virginiensis forewings, Vanessa cardui forewings, Polygonia interrogationis, Colobura dirce, Siproeta stelenes, Anartia fatima,* and *Chlosyne janais* (Carroll et al., 1994; Monteiro et al., 2013; Oliver et al., 2012; Reed and Serfas, 2004; Wee et al., 2022). | *Dll* has been functionally tested in multiple butterfly species. *Dll* plays a role in scale development, pigmentation, marginal band development, and eyespot center formation, and eyespot size (Connahs et al., 2019a; Monteiro et al., 2013; Murugeshan et al., 2022; Zhang and Reed, 2016). |
| 11 | *omb* | *optomotor-blind (omb)* was expressed along a broad domain along the AP boundary as well as in the lower posterior compartment (**Figure 2Q, Figure 3L, Figure S1 to S6**). During the later stages of development, expression in the eyespot centers was also observed | *Bicyclus anynana* (Banerjee and Monteiro, 2023). | Not tested. |

| | | | | |
|---|---|---|---|---|
| | | (**Figure S3M)**. These results are consistent with a previous finding (Banerjee and Monteiro, 2023). | | |
| 12 | *hh* | *hedgehog (hh)* expression was observed in the posterior compartment, consistent with previous studies (Keys et al., 1999; Saenko et al., 2011) (**Figure 2S, Figure 3M, Figure S1 to S6**). | *Junonia coenia* (Keys et al., 1999) and *Bicyclus anynana* (Saenko et al., 2011) | Not tested. |
| 13 | *en* | *engrailed* (*en*) was expressed in the posterior compartment and in the eyespot centers consistent with previous studies (Banerjee et al., 2020; Keys et al., 1999; Monteiro et al., 2006; Reed et al., 2020) (**Figure S1 to S6**). | *Junonia coenia, Vanessa cardui, Bicyclus anynana, Tithorea tarricina, Danaus Plexippus, Morpho peleides forewings, Caligo Memnon, Consul fabius, Hypna clytemenstra, Dryadula phaetusa, Hamadryas amphinome, Hamadryas februa, Catonephele numilia, Nessaea aglaura, Myscelia cyaniris, Vanessa virginiensis forewings, Vanessa cardui forewings, Polygonia interrogationis, Colobura dirce, Siproeta stelenes, Anartia fatima,* and *Chlosyne janais* (Banerjee and Monteiro, 2020; Banerjee et al., 2020; Brunetti | *en* has been functionally tested using RNAi in *Papilio* butterflies, where it plays a role in the location and identity of color pattern elements (Vankuren et al., 2023). |

| | | | | |
|---|---|---|---|---|
| | | | et al., 2001; Monteiro et al., 2006; Oliver et al., 2012; Reed et al., 2020) | |
| 14 | *Antp* | *Antennapedia* (*Antp*) was expressed in the eyespot centers consistent with previous studies (Oliver et al., 2012; Saenko et al., 2011; Shirai et al. 2012) (**Figure 2T, Figure 3N, Figure S1 to S6**). | *Junonia coenia, Vanessa cardui, Bicyclus anynana, Tithorea tarricina, Danaus Plexippus, Morpho peleides forewings, Caligo Memnon, Consul fabius, Hypna clytemenstra, Dryadula phaetusa, Hamadryas amphinome, Hamadryas februa, Catonephele numilia, Nessaea aglaura, Myscelia cyaniris, Vanessa virginiensis forewings, Vanessa cardui forewings, Polygonia interrogationis, Colobura dirce, Siproeta stelenes, Anartia fatima,* and *Chlosyne janais* (Matsuoka and Monteiro, 2021; Saenko et al., 2011) | *Antp* plays a role in eyespot development (forewings), eyespot white center differentiation (hindwings), eyespot size (hindwings), and silver scale development in *Bicyclus anyana* (Matsuoka and Monteiro, 2022; Prakash et al., 2022). |
| 15 | *dpp* | *decapentaplegic (dpp)* was expressed along the AP boundary and lower posterior compartment consistent with a previous study | *Pieris candida, Bicyclus anynana* (Banerjee and | *dpp* is involved in wing venation, eyespot center, and ring |

| | | | | |
|---|---|---|---|---|
| | | (Banerjee and Monteiro, 2020). *dpp* was also expressed in the intervein cells spanning the veins with missing expression in the eyespot centers during later stages (2.50-3.25), consistent with a previous study (Connahs et al., 2019b) (**Figure 2U, Figure 3O, Figure S1 to S6**). | Monteiro, 2023; Wee et al., 2022) | differentiation in *Bicyclus anynana* (Banerjee and Monteiro, 2020; Banerjee and Monteiro, 2023). |
| 16 | *ptc* | *patched (ptc)* was expressed along the AP boundary, consistent with previous studies in butterflies (Banerjee and Monteiro, 2023; Keys et al., 1999; Saenko et al., 2011) and *Drosophila* (Phillips et al., 1990) (**Figure 2V, Figure 3P, Figure S1 to S6**). | *Junonia coenia* (Keys et al., 1999), *Bicyclus anynana* (Banerjee and Monteiro, 2023; Saenko et al., 2011). | Not tested. |
| 17 | *Mad6* | *Mothers against dpp 6 (Mad6)* was expressed in two large clusters of cells along the AP domain and in the posterior compartment consistent with previous results (Banerjee and Monteiro, 2023) (**Figure 2W, Figure 3Q, Figure S1 to S6**). | *Bicyclus anynana* (Banerjee and Monteiro, 2023). | Not tested. |
| 18 | *spalt* | *spalt* was expressed in four distinct domains during the early larval stages (0.75-1.00), and in the eyespot center during the later stages (2.50-3.25), consistent with previous studies (Banerjee and Monteiro, 2020; Matsuoka and Monteiro, 2022; Oliver et al., 2012; Wee et al., 2022) (**Figure 2Y, Figure 3R, Figure S1 to S6**). | *Junonia coenia, Vanessa cardui, Bicyclus anynana, Tithorea tarricina, Danaus Plexippus, Morpho peleides forewings, Caligo Memnon, Consul fabius, Hypna clytemenstra, Dryadula phaetusa, Hamadryas amphinome, Hamadryas februa, Catonephele numilia, Nessaea aglaura, Myscelia* | *sal* plays a role in eyespot center formation, venation patterning, and eyespot and marginal band melanin pigmentation in multiple butterfly species (Banerjee and Monteiro, 2020; Matsuoka and Monteiro, 2022; Stoehr et al., 2013; Zhang and Reed, 2016; Wee et al., 2022). |

| | | | | |
|---|---|---|---|---|
| | | | *cyaniris, Vanessa virginiensis forewings, Vanessa cardui forewings, Polygonia interrogationis, Colobura dirce, Siproeta stelenes, Anartia fatima,* and *Chlosyne janais* (Banerjee and Monteiro, 2020; Brunetti et al., 2001; Monteiro, 2015; Monteiro et al., 2006; Oliver et al., 2012; Wee et al., 2022) | |
| 19 | *Notch* | *Notch* was expressed in the intervein cells during the earlier stages (0.75-1.00) and had a stronger expression in the eyespot centers during later stages (2.50), consistent with previous studies (Oliver et al., 2012; Reed and Serfas, 2004) (**Figure 2X, Figure S1 to S6**). | *Junonia coenia, Vanessa cardui, Bicyclus anynana, Tithorea tarricina, Danaus Plexippus, Morpho peleides forewings, Caligo Memnon, Consul fabius, Hypna clytemenstra, Dryadula phaetusa, Hamadryas amphinome, Hamadryas februa, Catonephele numilia, Nessaea aglaura, Myscelia cyaniris, Vanessa virginiensis forewings, Vanessa cardui forewings,* | Not tested. |

| | | | | |
|---|---|---|---|---|
| | | | *Polygonia interrogationis, Colobura dirce, Siproeta stelenes, Anartia fatima,* and *Chlosyne janais* (Oliver et al., 2012; Reed and Serfas, 2004; Saenko et al., 2011) | |
| 20 | *Optix* | *Optix* was expressed in two distinct domains in the upper anterior and in the lower posterior compartment in the forewing and in the upper anterior compartment in the hindwing, consistent with previous studies (Banerjee and Monteiro, 2020; Banerjee and Monteiro, 2023) (**Figure 2Z, Figure 3S, Figure S1 to S6**). *Optix* expression has been widely studied in the pupal wings of *Heliconius* butterflies (Reed et al., 2011). | *Heliconius erato, Heliconius melpomene, Heliconius cydno, Heliconius doris, Vanessa cardui, Bicyclus anynana* (Banerjee and Monteiro, 2023; Martin et al., 2014; Reed et al., 2011). | *Optix* plays a role in wing patterning, silver scale structural coloration, and ommochrome and melanin pigmentation in multiple butterfly species (Banerjee and Monteiro, 2023; Banerjee et al., 2024; Lewis et al., 2019; Prakash et al., 2022; Thayer et al., 2020; Zhang et al., 2017). |
| 21 | *vg* | *vestigial* (*vg*) was expressed in a broad domain along the wing margin consistent with a previous study in butterflies (Banerjee et al., 2023) and in *Drosophila* (Neumann and Cohen, 1997) (**Figure 2AF, Figure 3X, Figure S1 to S6**). | *Bicyclus anynana* (Banerjee et al., 2023) | Not tested. |

**Table S2: Genes examined in *Danio rerio* brain.**

| Sl. No. | Gene name | Expression (previously described) | Expression description |
|---|---|---|---|
| 1 | *gad2* | Zfin link<br><br>Expression pattern in Zebrafish UCL 6dpf | *Glutamate decarboxylase 2 (gad2)* was expressed widely throughout the brain, with strong expression in subpallial regions, preoptic area, and cerebellar regions (Filippi et al., 2014) (**Figure 4A, Figure S12A**).<br><br>. |
| 2 | *ompb* | Zfin link | *Olfactory marker protein b (omp)* was expressed in approximately 40% of the olfactory epithelial neurons. In addition, faint expression was present in a few neurons in the telencephalon (forebrain) around the eyes (Çelik et al., 2002) (**Figure 4E, Figure S12E**). |
| 3 | *chatb* | Zfin link | *choline O-acetyltransferase b (chatb)* exhibited the strongest expression in the habenulae (**Figure 4G, Figure S12G**), distinct from the expression of *chata* in mid and hind-brain motor neurons (Förster et al., 2017). |
| 4 | *tph2* | Zfin link<br><br>Expression pattern in 6dpf Zebrafish | *Tryptophan hydroxylase 2* (*tph2)* expression was seen in median and dorsal raphe and pineal complex cells as reported previously (Teraoka et al., 2004). In addition, we could also detect expression in the synaptic regions of the ventral habenula region. (**Figure 4H, Figure S12H**). |
| 5 | *nrp1a* | Zfin link | *Neuropilin 1a (nrp1a)* was asymmetrically expressed in the left dorsal habenula and sparsely in the diencephalic or the midbrain neurons (Kuan et al., 2007) (**Figure 4C, Figure S12C**). |
| 6 | *neurod 1* | Zfin link | *Neuronal differentiation 1 (neuroD)* expression was widespread throughout the brain, including the torus longitudinalis, cerebellum, diencephalon and telencephalon (Mueller and Wullimann, 2002) (**Figure 4B, Figure S12B**). |

| | | | |
|---|---|---|---|
| 7 | *oxt* | Zfin link | *Oxytocin* (*oxt*) was expressed symmetrically in the anterior hypothalamus, in a restricted cluster of cells (Unger and Glasgow, 2003) (**Figure 4D, Figure S12D**). |
| 8 | *kctd12.2* | Zfin link | *Potassium channel tetramerisation domain containing 12.2 (kctd12.2)* was expressed predominantly in the right dorsal habenula, particularly within the medial subnuclei, and a more restricted area in the left dorsal habenula (Doll et al., 2011) (**Figure 4F, Figure S12F).** |
| 9 | *elavl3* | Zfin link | *ELAV like neuron-specific RNA binding protein 3 (elavl3)* was expressed widely throughout the brain. It is known to be expressed in post-mitotic neurons (Park et al., 2000) (**Figure 4I, Figure S12I).** |
| 10 | *chrna3* | Zfin Link | *Cholinergic receptor, nicotinic, alpha 3 (chrna3)* was expressed predominantly in the hindbrain motor neurons, especially the facial motor nucleus, and cerebellum, matching the protein expression patterns observed in transgenic zebrafish (Hua *et al.*, 2025) and mRNA expression (Raine et al., 2025) (**Figure S12J**). |
| 11 | *chrm3a* | Zfin Link | *Cholinergic receptor, muscarinic 3a (chrm3a)* expression was observed throughout the brain, with expression matching previous expression data available at mapZebrain (link). The expression is also expected in the Retinoganglion epithelial cells (Nuckels *et al.*, 2011) (**Figure S12K**). |

**Table S3. Comparative metrics of the different multiplex-FISH technologies.**

| Metric | RAMFISH (This Study) | EASI-FISH/cycleHCR | MERFISH/seqFISH | Commercial (Xenium/CosMx) |
|---|---|---|---|---|
| **Primary Niche** | **Macro-domain mapping** | Subcellular/Single-molecule | Single-molecule profiling | Single-molecule profiling |

| | | | | |
|---|---|---|---|---|
| **Achievable Plex** | **Mid-plex (~30–40 targets)** | High-plex (50-300+) | Ultra-high (100-10,000) | Ultra-high (100-5,000+) |
| **Tissue State** | **Free-floating/Intact 3D hydrogel embedding** | Hydrogel-embedded/Cleared | Thin sections (mostly 5-15 μm) | Thin sections (strictly 2D) |
| **Hardware Cost Elements** | **Standard benchtop + shared confocal. Optional Open-source hardware (<$10k to build)** | Standard Confocal + Fluidics | Custom/Specialized Optics | > $250,000 (Proprietary) |
| **Cycle Time** | **~8 hours/ cycle** | ~10-24 hours/ cycle | Minutes to Hours | Continuous (Fully Automated) |
| **Alignment Strategy** | **Manual rigid + Affine and B-Spline/ *In-silico* anchoring** | Physical (Hydrogel Restraint) | Physical/Fiducial Markers | Proprietary/Fiducial |
| **Computational Overhead** | **Local/Moderate** | High (Cluster recommended) | High (Cluster recommended) | Integrated/Very High |
