## Supplementary File 2 for "A spatial mRNA profiling workflow using Rapid Amplified Multiplex FISH (RAMFISH)"

Affiliations

**Supplementary file 2**

**Probe sequences**

***Bicyclus anynana* specific genes**

**>*Wnt1***

lcl|XM_024099417.2_cds_XP_023955185.1_1 [gene=LOC112058530] [db_xref=GeneID:112058530] [protein=protein Wnt-1] [protein_id=XP_023955185.1] [location=57..1250] [gbkey=CDS]

ATGTCGGGTCCGCCCATAATGAAGTGGTTGTGCTTGTTTGTGCTGTTTCTGTGTATGAGGTGCGAGGCCAACAAGCCGAGGCGAGGCCGAGGCAGCATGTGGTGGGGCATAGCAAAGGCAGGCGAACCAAATAACTTATCACCCTTGTCTCCAAGTGTCCTATACATGGACCCGGCTGTTCACGCCACCTTGAGAAGGAAACAGAGAAGGCTAGCGAGGGAGAACCCTGGGGTTCTCGCAGCAATATCCAAGGGAGCCAGCATGGCTGTGGCCGAATGCCAGCATCAGTTCAAATACAGGAGATGGAACTGTTCTACAAGAAATTTTTTGCGAGGGAAGAATCTATTTGGAAAAATTGTTGACAGAGGTTGCCGTGAAACCGCCTTCATCTACGCCATCACAAGCGCGGGGGTGACGCACGCGGTGTCGCGCGCATGCGCCGAAGGTTCCATCGAGTCCTGCACGTGCGACTATTCTCATGTGGACCGTTCGCCGCACCGCGCGCGCGCCGCCGCCGCCGCCAACGTGAGGGTCTGGAAATGGGGCGGGTGCAGCGACAACATCGGCTTCGGCTTCAAGTTCAGCCGAGAGTTCGTTGACACCGGGGAAAGGGGCAAGACGCTTAGGGAGAAGATGAACTTGCACAACAATGAGGCTGGCAGAATGCACGTGCAAACGGAGATGCGCCAGGAGTGCAAGTGCCACGGTATGTCTGGGTCCTGCACGGTGAAGACGTGCTGGATGAGGCTGCCGACGTTCCGGTCTGTAGGCGACGCCCTGAAAGACAGCTTCGACGGGGCGTCGCGGGTCATGATGCCCAATACCGAGGTGGAGGCGCCGTCGCAGAGGAACGACGCCGCACCTCACAGGGTCCCGCGCCGTGACCGCTACAGGTTCCAACTTCGGCCGCACAACCCTGACCACAAAACACCCGGGGTCAAGGACCTTGTATACTTGGAATCTTCACCAGGTTTCTGCGAAAAGAACCCCAGACTGGGCATCCCGGGTACGCACGGGCGTGCCTGCAACGACACTAGCATCGGCGTCGACGGTTGCGACCTGATGTGCTGCGGGCGCGGCTACCGGACCGAGACCATGTTCGTAGTGGAACGATGCAACTGCACGTTCCACTGGTGCTGCGAGGTCAAATGCAAATTGTGTCGCACGGAAAAGGTAGTTAACACGTGTTTATAG

Wnt11_HCR_P1B1: gAggAgggCAgCAAACggAACCTAGCTGGACTGACAAAATCCAGC

Wnt11_HCR_P2B1: TTCGCCTGCCTTTGCTATGCCCCCCTAgAAgAgTCTTCCTTTACg

Wnt12_HCR_P1B1: gAggAgggCAgCAAACggAAGTATAGGACACTTGGAGACAAGGGT

Wnt12_HCR_P2B1: CAAGGTGGCGTGAACAGCCGGGTCCTAgAAgAgTCTTCCTTTACg

Wnt13_HCR_P1B1: gAggAgggCAgCAAACggAACCCAGGGTTCTCCCTCGCTAGCCTT

Wnt13_HCR_P2B1: GGCTCCCTTGGATATTGCTGCGAGGTAgAAgAgTCTTCCTTTACg

Wnt14_HCR_P1B1: gAggAgggCAgCAAACggAATATTTGAACTGATGCTGGCATTCGG

Wnt14_HCR_P2B1: TTTCTTGTAGAACAGTTCCATCTCCTAgAAgAgTCTTCCTTTACg

Wnt15_HCR_P1B1: gAggAgggCAgCAAACggAAGTCAACAATTTTTCCAAATAGATTC

Wnt15_HCR_P2B1: CGGGGGGGCTTTGTCCGGCGAAACTTAgAAgAgTCTTCCTTTACg

Wnt16_HCR_P1B1: gAggAgggCAgCAAACggAAGCGTGTCGACGTGTATATTAGTAAT

Wnt16_HCR_P2B1: ATGAAGATGTCGCGTCAATCGTCAATAgAAgAgTCTTCCTTTACg

***>Wnt6***

lcl|XM_052891217.1_cds_XP_052747177.1_1 [gene=LOC112058531] [db_xref=GeneID:112058531] [protein=protein Wnt-6] [protein_id=XP_052747177.1] [location=196..1251] [gbkey=CDS]

ATGGATAATACAAGAGAGACGGGGTTTGTGAACGCGATCACTGCAGCCGGAGTGACATACGCGATCACCCGCGCCTGTACTGCGGGCTCACTGCTCGAGTGCTCATGTGAAAAGGGTGTACCAAAACCGCGTCGTGGAAGAACTCAAACGCCCCAGCCCCCAGCACCAACTCAGACAGAGCAGTGGCAGTGGGGCGGATGCAGTGACAACGTCCGCTTCGGCCTGCAGAAGTCCAGGGAATTCATGGACAGTAGATACAGGAAGAGGAGCGACATCAAAACGATGATAAAGCTGCATAACCACAACGCTGGGAGGTTGGCAATCAAAAATAACATGAAAGTAGACTGTAAATGTCACGGCCTATCTGGCTCATGCACACTGCGAACTTGTTGGTGGAGAATGCCCACCTTTAGAGAAGTGGGGGACCGATTGAGAGACAACTTTGAAGGTGCTGCTAAGGTGATCTCAAGTAATGATGGCGACAGTTTTATGCCCGAAAGTCCTAACATCAAGCGACCTGGGAAAAAAGATATCATATACTCTGAAGAATCACCCGATTTCTGCGGACCTAACATGAAGACAGGGTCACTCGGCACTGAAGGGCGCCAGTGCAATATAAGTTCTGCGGGAACTGACAGTTGCGATCAACTTTGTTGTAGAAGAGGGTACATACAAACATCTATAAAGGAGGCTGAAAATTGCAATTGTCAATTTAAGTGGTGTTGCGAAGTCATTTGTAAAACATGCTATGTGAAGCGAGACATACAAACGTGCCTTTAA

Wnt61_HCR_P1B2: CCTCgTAAATCCTCATCAAACGTTCACAAACCCCGTCTCTCTTGT

Wnt61_HCR_P2B2: CGTATGTCACTCCGGCTGCAGTGATAAATCATCCAgTAAACCgCC

Wnt62_HCR_P1B2: CCTCgTAAATCCTCATCAAACTGTCTGAGTTGGTGCTGGGGGCTG

Wnt62_HCR_P2B2: CACTGCATCCGCCCCACTGCCACTGAAATCATCCAgTAAACCgCC

Wnt63_HCR_P1B2: CCTCgTAAATCCTCATCAAATGCCAACCTCCCAGCGTTGTGGTTA

Wnt63_HCR_P2B2: TACAGTCTACTTTCATGTTATTTTTGAAATCATCCAgTAAACCgCC

Wnt64_HCR_P1B2: CCTCgTAAATCCTCATCAAAAGATCACCTTAGCAGCACCTTCAAA

Wnt64_HCR_P2B2: GCATAAAACTGTCGCCATCATTACTAAATCATCCAgTAAACCgCC

Wnt65_HCR_P1B2: CCTCgTAAATCCTCATCAAATGCACTGGCGCCCTTCAGTGCCGAG

Wnt65_HCR_P2B2: AACTGTCAGTTCCCGCAGAACTTATAAATCATCCAgTAAACCgCC

Wnt66_HCR_P1B2: CCTCgTAAATCCTCATCAAATTTATAGATGTTTGTATGTACCCTC

Wnt66_HCR_P2B2: AATTGACAATTGCAATTTTCAGCCTAAATCATCCAgTAAACCgCC

***>Wnt10***

lcl|XM_024099419.2_cds_XP_023955187.2_1 [gene=LOC112058532] [db_xref=GeneID:112058532] [protein=protein Wnt-10a] [protein_id=XP_023955187.2] [location=663..1784] [gbkey=CDS]

ATGAGGAAGTTGAAAGTTGCAGTTGGAAAACGCAGAATGCACAGGACGTACCCGGGGGCGATATATTTTGTGCTGATTGCATTTTTTGAGGTTGTGAGCTCAAGAGACAACATGCTGCCACACCACCTGAAGCTCAGTTCTACTCTAACTTGTCGACTCATCGGTGGTCTGACCAGAGAACAGAGATCTGTCTGCCACGACGCGGCCGATACAGCAGCCATCGCCTTCGAGGGTCTGCAGATGGCGGTCAAGGAGTGCCAGCATCAGTTCCGCTGGCACAGGTGGAACTGCTCCAGTCTGCTGGTCAAGAGTTCCAATCCTCACGCCAGTGCTATTATGAAGAGAGGATTCCGGGAAACCGCGTTCCTGTACGCCCTAACAGCGGCAGGAGTAGCTCACGCAGTGGCCCGGGCGTGCGCCCAGGGCCGGCTCATATCCTGCGGCTGCGACCCCCTGGGGTACCGCGCAGCCCATGAGAGGGGCCGCACGAGGACCAACAAGTGGGAGTGGAGTGGCTGCTCCCACAACCTGGCCTATGGCGTCGAGTTCTCCAAGAAATTCCTCGATGTACGGGAAAAGGTGGACGATCTGCAGTCGAAGATCAACGTACATAATAACAATGCTGGTAGATCGATTCTATCATCTCACATGGAGGTGCGGTGCAAGTGCCACGGGCTGTCAGGAAGTTGTCAACTGCGAACGTGTTGGCGCGCCACGCCCGACTTCAGGGCTGTGGCTTCTACTATTAAGAGACAATACCGCAAAGCTTTAGTAGTAGCCCAAGAAGAGCTCAATAACAGCCCTTCAGTGTTACGAGGGCGGCCACGAGGAAGAAGGAGGAGTCGAGCAAGACCTGCACCGAAGTCTAGCTTGCTGTTTTTTGAGAAGTCCCCAAGTTTTTGTGAAGCAGACCCCAAATTTGATTCCGCGGGTACATCAGGAAGAGTCTGCCGCATCGGAAGGACAACAAGGACTGGATCCTGTGACCTGCTGTGCTGTGGACGAGGACACGCCCTCATCAGAAAGTCAAGTATCAAACCATGTAACTGCACCTTTCACTGGTGCTGTAGAGTCGATTGCCAGAGGTGCCAGGATGATAAATGGATTTCAATTTGCAAGTAA

Wnt101_HCR_P1B2: CCTCgTAAATCCTCATCAAACACAACCTCAAAAAATGCAATCAGC

Wnt101_HCR_P2B2: GTGTGGCAGCATGTTGTCTCTTGAGAAATCATCCAgTAAACCgCC

Wnt102_HCR_P1B2: CCTCgTAAATCCTCATCAAACCGATGAGTCGACAAGTTAGAGTAG

Wnt102_HCR_P2B2: ACAGATCTCTGTTCTCTGGTCAGACAAATCATCCAgTAAACCgCC

Wnt103_HCR_P1B2: CCTCgTAAATCCTCATCAAACTGGAGCAGTTCCACCTGTGCCAGC

Wnt103_HCR_P2B2: TGAGGATTGGAACTCTTGACCAGCAAAATCATCCAgTAAACCgCC

Wnt104_HCR_P1B2: CCTCgTAAATCCTCATCAAACGCGGTTTCCCGGAATCCTCTCTTC

Wnt104_HCR_P2B2: TCCTGCCGCTGTTAGGGCGTACAGGAAATCATCCAgTAAACCgCC

Wnt105_HCR_P1B2: CCTCgTAAATCCTCATCAAAGTCCTCGTGCGGCCCCTCTCATGGG

Wnt105_HCR_P2B2: GAGCAGCCACTCCACTCCCACTTGTAAATCATCCAgTAAACCgCC

Wnt106_HCR_P1B2: CCTCgTAAATCCTCATCAAAAATTTCTTGGAGAACTCGACGCCAT

Wnt106_HCR_P2B2: TCGTCCACCTTTTCCCGTACATCGAAAATCATCCAgTAAACCgCC

***>frizzled4 (fz4)***

lcl|XM_024081478.2_cds_XP_023937246.1_1 [gene=LOC112045330] [db_xref=GeneID:112045330] [protein=frizzled-4] [protein_id=XP_023937246.1] [location=337..2052] [gbkey=CDS]

ATGAAGTGTTTTGTGGTATTAATAACTGTGACCCTCGTCTACGAGATCGTAGCCGAGGCGTCCGTGAGGACATGTGAACCGATCAAGGTGGCTATGTGCAAGAACATCGGATACAACCAGACGGGAATGCCCAACTTGGCTCGGCACACTCTCCAAGCGGACGCCGACGTCACACTACAGACCTTCAGCCCCCTGGTGCAGTATGGATGCTCGTCCCAGTTGCATTTGTTTTTGTGTGCGGTGTACGTGCCCATGTGCACTGATAAGGTCGCGTTACCGATAGGTCCGTGTAGGGGTTTGTGCGAGAGTGTTTACGCCAGATGTTACCCCGTGTTGCGGGGTTTCGGGTTCCCTTGGCCGGCGGAGTTGGACTGTTCTCTATTCCCGGCGGAGAATAACCATGAACACATGTGTATGGAGGGTCCTGGGGAGCGCGCGCCGCCGATCGGTACGATACCGTTGGACACGACGAGGGAAGGACGAGGGACTTGCAGACGTTTGGTGAAGCCGAACAGTTGGGTGTGGGTTAGGGGGTCGGGGCGGTGCGCGCAGTTCTGTGACGCTGAAGTGTTGTGGGAGGTGGGGGAGAGGCGGGCGGCTGAGGTGTGGCTGGCGACTTGGGCAGCGTTGAGCTTCGCGTGTACTCTTGCGGCGGTGGCGGCGCAGCTGGCGTGTGGAGCCAGGGGCGGAGCTGGCGAGCGCGCGTTGGTGCTGGTGGCACTGTGCCGGTGCGCGGCGGCGGCGGGCTGGGGCGTGCGCGCGGCTGCGGGCCGGACGGCGGCGGGCTGCGCCAAGGACTCCACGTCTCCCACCAGGATGCTGTTAGCGCATGACGGCTTAGCGAACCCCAACTGCGCGGTCGTCTTCTTGCTGCTGTATTACTTCGGGCTGGCTGCGTCAGTTTGGTGGGTGGTAGTGACGGGTGCGTGGCGAGCGAGCGTGCTGCGCCCCCCAGCCAGTAGCGGCGCCAGGAACGACCGCCACTCCTCCCTGCTGCAGCTGGCTGCGTGGGGCGTGCCCGCCGCGCTGGCCGCCGCCGTGCTGGTCACAAGGGACGTGGATGCTGATGAGCTTACAGGCACATGCTTCGTGGGCAATCAGTCTAGCAAATCTTTGCTAGCGTTGGTTATCGTTCCCGAAGCGATATGCTTGCTTCTGGGCAGCGTGTTCCTCGCTTCCGGCCTTCGCGCAGTGCTTCGTAAACCTGTACCGATTCCAGCTCCGGCAACGCTTCTGAATTCTGCACCGCAAGCGCACCCTGATCAGAGTCTGCTGAGGCTAGGAGCCTTCGCTGCGTTGTACGCTGTGCCATCTGCCTGCATACTGGCGACATGGGTGTATGAATATATTCTGAGAGAAAATTGGTTGGCTGCCCCCGTCCCTTCGACGGAACCTTCGACGCAACCGCGTCCAGCATTCTGGGTGTTCCTTTTTAGGATATTCGCATCACAAATACTGGGTGTCATGGTGGCGGTTTGGATAGCCACGCCACGGTTGAAGGCGCTGTGGAGGAGAATCAGTGGGCCAAGAAAACCAGCGTTAGCAAAATGCCCACCTGGTCCGACGCCTACACCGTTAACATTGCATTGCTACGCTACGCATCCGCATACGTTGACGAGACACCCACAGAAATACGCCACATACCGACCACCACAGCAACAGTCGTATAGAAAACCTCGACATTATCATTATTCTGCTGGAGAAACTATCCTATGA

Fz4_1_P1B2	CCTCgTAAATCCTCATCAAATGATCGGTTCACATGTCCTCACGGA

Fz4_2_P1B2	CCTCgTAAATCCTCATCAAACATACTGCACCAGGGGGCTGAAGGT

Fz4_3_P1B2	CCTCgTAAATCCTCATCAAAAACATCTGGCGTAAACACTCTCGCA

Fz4_4_P1B2	CCTCgTAAATCCTCATCAAACAGCGTCACAGAACTGCGCGCACCG

Fz4_5_P1B2	CCTCgTAAATCCTCATCAAACCCTGGCTCCACACGCCAGCTGCGC

Fz4_6_P1B2	CCTCgTAAATCCTCATCAAAACGTGGAGTCCTTGGCGCAGCCCGC

Fz4_1_P2B2	ATCCGATGTTCTTGCACATAGCCACAAATCATCCAgTAAACCgCC

Fz4_2_P2B2	AAAACAAATGCAACTGGGACGAGCAAAATCATCCAgTAAACCgCC

Fz4_3_P2B2	GGAACCCGAAACCCCGCAACACGGGAAATCATCCAgTAAACCgCC

Fz4_4_P2B2	GCCTCTCCCCCACCTCCCACAACACAAATCATCCAgTAAACCgCC

Fz4_5_P2B2	GCACCAACGCGCGCTCGCCAGCTCCAAATCATCCAgTAAACCgCC

Fz4_6_P2B2	CATGCGCTAACAGCATCCTGGTGGGAAATCATCCAgTAAACCgCC

***>frizzled2 (fz2)***

lcl|XM_052885856.1_cds_XP_052741816.1_1 [gene=LOC112043399] [db_xref=GeneID:112043399] [protein=frizzled-2] [protein_id=XP_052741816.1] [location=440..2101] [gbkey=CDS]

ATGAAAAAAGAAAAACCGACGTTTATGAGGCTGTTGTGTACAATTTTGAATCCGCGGCTCGTGGCGCTGCGCGCGCGCAACGCCAGCGCGCCCGGCGTGCCGGCGCCCGCCTGCGGCCTGCCGTGCCGCGGCGCCTTCTTCTCGCGCGAGGAGAAGGAGTTCGCCGCCGTGTGGGTCGCGCTGTGGGCCGGCCTGTGCGCCGCCTCCACGCTCATGACGCTCACCACCTTCGCCATCGACTCGCAGCGCTTCAAGTACCCCGAGCGGCCCATCGTGTACCTGTCGGCCTGCTACTTCATGGTGGCGCTGGGCTACCTGGCGCGCCTGGCGCTGGGCCACGAGGCCGTGGCGTGTGACGGGCCGCTGCTGCGGACGTCGGCGGGCGGGCCCGGCGCCTGCACGCTGGTGTTCGTGCTGGTGTACTTCTTCGGCATGGCGTCGTCCATCTGGTGGGTGGTGCTGTCGTTCGCGTGGTTCCTGGCCGCCGGGCTCAAGTGGGGCAACGAGGCCATCGCCGGGCACGCGCAGTACTACCACCTGGCGGCGTGGCTGGTGCCGGCCGCCAAGACCGTGGCCGTGCTGCTGGCGGGCGCCGTGGACGGCGACCCCGTGGCCGGCGTGTGCTACGTGGGCAACTCGTCGCCCGAGCACCTGCGGCGCTACGTGCTGGCGCCGCTCGTGGTGTACTTCGCGCTGGGCGCCTCCTTCCTGCTGGCCGGCTTCGTGTCGCTGTTCCGCATCCGCAGCGTCATCAAGCGGCAGGGCGGCGCGGGCGCGGGCTCCAAGGCCGACAAGCTGGAGAAGCTCATGATCCGCATCGGCGTGTTCAGCGTGCTGTACGCCGTGCCGGCCGGCGTGGTGATCGGCTGCCTGGCGTACGAGGCGGGCGGGCGCGAGCGCTGGCTGCGGCGCGTGGCGTGCGGCGCGGCGTGCGGCCCGCGCCCCGTCTACTCGGCGCTCATGCTCAAGTACTTCATGGCGCTGGCCGTGGGCATCACGTCGGGCGTGTGGATCTGGTCGGGCAAGACGCTGGAGTCGTGGCGCCGCGTGTGGCGCGGGGGGCGCGCGCCGCCGCCCGCGCAGCGCGCGCTG

Ftz21_HCR_P1B4: CCTCAACCTACCTCCAACAAGAACTCCTTCTCCTCGCGCGAGAAG

Ftz21_HCR_P2B4: GGCCCACAGCGCGACCCACACGGCGATTCTCACCATATTCgCTTC

Ftz22_HCR_P1B4: CCTCAACCTACCTCCAACAAAAGTAGCAGGCCGACAGGTACACGA

Ftz22_HCR_P2B4: CGCGCCAGGTAGCCCAGCGCCACCAATTCTCACCATATTCgCTTC

Ftz23_HCR_P1B4: CCTCAACCTACCTCCAACAAAGAAGTACACCAGCACGAACACCAG

Ftz23_HCR_P2B4: CCCACCAGATGGACGACGCCATGCCATTCTCACCATATTCgCTTC

Ftz24_HCR_P1B4: CCTCAACCTACCTCCAACAACCGGCACCAGCCACGCCGCCAGGTG

Ftz24_HCR_P2B4: CCAGCAGCACGGCCACGGTCTTGGCATTCTCACCATATTCgCTTC

Ftz25_HCR_P1B4: CCTCAACCTACCTCCAACAAGAAGTACACCACGAGCGGCGCCAGC

Ftz25_HCR_P2B4: GGCCAGCAGGAAGGAGGCGCCCAGCATTCTCACCATATTCgCTTC

Ftz26_HCR_P1B4: CCTCAACCTACCTCCAACAAGCCGATGCGGATCATGAGCTTCTCC

Ftz26_HCR_P2B4: CGGCACGGCGTACAGCACGCTGAACATTCTCACCATATTCgCTTC

***> frizzled9 (fz9)***

lcl|XM_024087045.2_cds_XP_023942813.2_1 [gene=LOC112049235] [db_xref=GeneID:112049235] [protein=frizzled-9] [protein_id=XP_023942813.2] [location=223..1767] [gbkey=CDS]

ATGATGGGGAGGATTTTGTGTGTTTTTCTCCTGGTCTGGATGGTCGCTGCTGGCCAGGAGGAGGAGGGTGGGAAATGCGAGAGGATCACCCTGTCCCAGTGCCAGGATCTGGGGTACAATTGGACCGCTATGCCCAATCTCATTGGACATAGAGACCAGAAGGAGGCAGAAGAAGCGATGACTGCGTTCACGAGCATCCTTGCGAGCGAATGCTCGGTGCACGCTCGCTTCCTGCTGTGCTCGGCCTTCGCCCCGCTGTGCTCCGAGCAGGTGTCGGGCTCCGTCAGCGCCTGCCGCGCGCTCTGCGACAAGGTCGTCCGCGACTGCAAGAACCAGATCGCGGCCCTGCCCGACGGCATCAAGCTCGACTGCTCCGCCTTCCCGCTCCGCCCGGACTGGCGGCTCTGCATGCGGCCGCCCAACGCCAGCGAGGAGCCCGAGCCGCCGCCGGTGCCGCGCTGGCCCTTCAACGAGCAAGATCTGAAAGAGCACGCGTGTCCCCCGGGCTACGCGCACTCCCCCACGGGCTCGTGCTGGCCCGCCTGCGACAAGCCCGCGCGCTACACGCAAGCTAACAAGAGAAAAGCGGAGATATGGATGCTCACGCTCGCCTGGTTCTCTCTGCTCTCCACTTCTTTCGCGCTGCTCACGTTCTGCGCTGAACCGTCGCGATACCGCTACCCCGAGAGGCCCGTCGTCTGGATGGCCGCGTGCCACGCTGTCGTCGCGCTCGCCTACGTCACGAGAGGCTGGCTCGGTGCGAAGACCGTCTCCTGCACTGGAAACCTGTTGGCCGTCGACGGAATGGGTTCGACGATTTGCGTCGCATTCTTCTCTCTTACATACTACTTCACGTTAGCGGCGGACGCGTGGTTCGCGAACGCGTGCGTAGCGTGGTACTTGACCGCTGCGAGCGAGTGGTCCACCGAGGCGTTGGAGCGGGCGGCGGCGTACCTGCACGCGGTGGCGTGGGGCTGGGCGGGGGCGTGGACGGCGGCGGCGCTGGCGCTGCGGCGGGTGACGGCGGACGAGCTGACCGGGACCTGCGGCGTGTCGGACGAAGCGGCCGCGGCGCTGATCGGCGTCCCGCGAGGCGCTCTGCTCATCACCTCAATAGCGTTGGCTATCGGCGCGTGTCCGGGCATAATGAGAGTGCGCCGCGCGTTGGACTCGCGAGGAGCGAAACGAGTGGGACGGTTGGCCGCTCGAGCGGCCGCTGGAGGCCTGCTCTACCTGTTCTTAGCGGCGTGCGCGACCGGCGCGCGCGTGATTGAAGCCAGAAACAGAGCGGCGCAAAAGAGCTTGCCGCTTCTGGCGGCGATGGGCGCGGGCAGTCTGGATGACTCGGCGGGGTCGCGCGCCGTGGCCGTGAGCCTGTGGCTGTGCTGCGGGGTCGCCGCGGGGGCGTGGTCGTGGTCGCGCAAGTCGGCGGTGGTGTGGCGCAAGGCGCTGTGTCCGCCGCGCAAGGCCCCGTGCTGCGCGCCGCCGCTGCTCGGACCCCATCACCCGTACTATAAAAGACCTCTGCACGTGTCTAGAGTGTGA

Ftz91_HCR_P1B4: CCTCAACCTACCTCCAACAACTGGTCTCTATGTCCAATGAGATTG

Ftz91_HCR_P2B4: CGCAGTCATCGCTTCTTCTGCCTCCATTCTCACCATATTCgCTTC

Ftz92_HCR_P1B4: CCTCAACCTACCTCCAACAAAGGCGCTGACGGAGCCCGACACCTG

Ftz92_HCR_P2B4: GGACGACCTTGTCGCAGAGCGCGCGATTCTCACCATATTCgCTTC

Ftz93_HCR_P1B4: CCTCAACCTACCTCCAACAAGGCGTTGGGCGGCCGCATGCAGAGC

Ftz93_HCR_P2B4: CACCGGCGGCGGCTCGGGCTCCTCGATTCTCACCATATTCgCTTC

Ftz94_HCR_P1B4: CCTCAACCTACCTCCAACAAGCGCGGGCTTGTCGCAGGCGGGCCA

Ftz94_HCR_P2B4: CTTTTCTCTTGTTAGCTTGCGTGTAATTCTCACCATATTCgCTTC

Ftz95_HCR_P1B4: CCTCAACCTACCTCCAACAAACGGGCCTCTCGGGGTAGCGGTATC

Ftz95_HCR_P2B4: ACAGCGTGGCACGCGGCCATCCAGAATTCTCACCATATTCgCTTC

Ftz96_HCR_P1B4: CCTCAACCTACCTCCAACAATGCGACGCAAATCGTCGAACCCATT

Ftz96_HCR_P2B4: CGTGAAGTAGTATGTAAGAGAGAAGATTCTCACCATATTCgCTTC

***>WntA***

lcl|XM_052886643.1_cds_XP_052742603.1_1 [gene=LOC128198014] [db_xref=GeneID:128198014] [protein=protein Wnt-1-like] [protein_id=XP_052742603.1] [location=381..1460] [gbkey=CDS]

ATGGATGACATAAACAAGATCCCTTCCCCCCATAAACAAGTTGAATGCGCTTCCCCGATGCCGCAGTCTCTGCACGTGCGGCACTCAAGGAATTTAGCAGCACCGAATAGGCCTGTACAATCGTCTAACACCTCTGTGGAAACCTTTACAATACTGCACAAAGAAAGCTGCCATAGATTAGAGTATCTCGTCGAACGACAAAAGCAATTATGTATGCTTTCTGATAAAATGGTACAGGTGATACAAACAGGAGCGCAACAGGCAATTGATGAATGTCAGCATCAATTTCGGAATAGCCGTTGGAACTGTAGTACCGTCGACAATTCCACTGATATATTCGGCGGAGTGCTAAAATTTAAATCTCGCGAGTCTGCATTCGTCCACGCTCTGTCAGCAGCAGCATTGGCTCACACAGTTGCTCGCGCGTGCAGTCGGGGCGAACTAAACGAGTGTTCCTGTGACGCTCGTGTTAGAAAGCGAACGCCGCGGCATTGGCAGTGGGGTGGTTGTTCTGAGGATATAAGATATGGAGAAAAGTTCAGTCGTGACTTTGTAGATGCTAAAGAAGACAAGGATAATGATGAAGGTCTCATGAACTTACATAACAATGAAGCTGGCCGCAGAGCAGTCCGCGGCAGGATGCAGCGCGTGTGCAAATGCCACGGCATGTCGGGCTCGTGCTCCGTGCGCGTGTGCTGGCGCCGCCTGCCGCAGCTGCGGCTGGTGGGCGACGTGCTGAGCACCAGATACGAGGGCGCCTCTCATGTTAAGGTTGTAGAGAGGAAGAGAGGCAAGAATATAAGAAAACTGCGACCGCTGCATCCTGATATAAAGAAACCGAACAAAACCGATCTAGTCTATCTCGAGGACTCTCCCGATTACTGTGAACCGAACGACGAGTAA

Wnta1_HCR_P1B1: gAggAgggCAgCAAACggAACAACCACCCCACTGCCAATGCCGCG

Wnta1_HCR_P2B1: TTTTCTCCATATCTTATATCCTCAGTAgAAgAgTCTTCCTTTACg

Wnta2_HCR_P1B1: gAggAgggCAgCAAACggAACCTTGTCTTCTTTAGCATCTACAAA

Wnta2_HCR_P2B1: GTAAGTTCATGAGACCTTCATCATTTAgAAgAgTCTTCCTTTACg

Wnta3_HCR_P1B1: gAggAgggCAgCAAACggAACCGCGGACTGCTCTGCGGCCAGCTT

Wnta3_HCR_P2B1: TGGCATTTGCACACGCGCTGCATCCTAgAAgAgTCTTCCTTTACg

Wnta4_HCR_P1B1: gAggAgggCAgCAAACggAAGCACACGCGCACGGAGCACGAGCCC

Wnta4_HCR_P2B1: CAGCCGCAGCTGCGGCAGGCGGCGCTAgAAgAgTCTTCCTTTACg

Wnta5_HCR_P1B1: gAggAgggCAgCAAACggAACGCCCTCGTATCTGGTGCTCAGCAC

Wnta5_HCR_P2B1: TCCTCTCTACAACCTTAACATGAGATAgAAgAgTCTTCCTTTACg

Wnta6_HCR_P1B1: gAggAgggCAgCAAACggAACGGTCGCAGTTTTCTTATATTCTTG

Wnta6_HCR_P2B1: GTTCGGTTTCTTTATATCAGGATGCTAgAAgAgTCTTCCTTTACg

Wnta7_HCR_P1B1: gAggAgggCAgCAAACggAAGGGAGAGTCCTCGAGATAGACTAGA

Wnta7_HCR_P2B1: TTACTCGTCGTTCGGTTCACAGTAATAgAAgAgTCTTCCTTTACg

***>Optix***

lcl|XM_024080404.2_cds_XP_023936172.1_1 [gene=LOC112044525] [db_xref=GeneID:112044525] [protein=homeobox protein SIX6-like] [protein_id=XP_023936172.1] [location=39..851] [gbkey=CDS]

ATGCGCGGCTCCTGGGACGAGTCCACGACGGCGGCGCTGCACGCGCGCATCCTGGAGGCGCACCGCGGGTCCGCCGCGCCCGACCGCGCCGAGCCCGCGTGCGAGCCTCCGCCGCTGACGCTGGGCGCGCTGGAGCTGGCGGCGCCCACGCCGCTGCTGCCGCTGCCCACGCTGAGCTTCAGCGCCGCGCAGGTGGCCACCGTGTGCGAGACGCTGGAGGAGAGCGGCGACGTGGAGCGCCTGGCGCGCTTCTTGTGGTCGCTGCCCGTGGCGCACCCCAACGTGGCCGAGCTGGAGCGCTGCGAAGCCGTGCTGCGCGCGCGCGCCGTCGTCGCCTTCCACGCCGGCCGCCACCGCGAGCTGTACGCCATCCTCGAGCGCCACCGCTTCCAGCGCTCCAGCCACGCCAAGCTGCAAGCGCTGTGGCTGGAGGCGCACTACCAGGAGGCTGAGCGCCTGCGCGGCCGTCCGCTGGGCCCCGTCGACAAGTACCGCGTGCGGAAGAAGTTCCCGCTCCCGAGGACGATCTGGGACGGCGAGCAGAAGACGCACTGTTTCAAGGAGCGGACGCGATCTCTACTCCGAGAATGGTACCTCCAAGATCCCTACCCGAACCCGACGAAGAAGAGGGAATTGGCGGCGGCGACGGGTCTGACGCCGACGCAAGTCGGCAACTGGTTCAAAAACCGACGGCAAAGAGACCGAGCGGCCGCCGCCAAGAACCGCTCCGCCGTGCTGGGCAGAGGATAA

Optix1_HCR_P1B2: CCTCgTAAATCCTCATCAAACGCCGTCGTGGACTCGTCCCAG

Optix1_HCR_P2B2: CGCCTCCAGGATGCGCGCGTGCAGCAAATCATCCAgTAAACCgCC

Optix2_HCR_P1B2: CCTCgTAAATCCTCATCAAAGGCTCGGCGCGGTCGGGCGCGGCGG

Optix2_HCR_P2B2: AGCGTCAGCGGCGGAGGCTCGCACGAAATCATCCAgTAAACCgCC

Optix3_HCR_P1B2: CCTCgTAAATCCTCATCAAAGGCAGCAGCGGCGTGGGCGCCGCCA

Optix3_HCR_P2B2: GCGGCGCTGAAGCTCAGCGTGGGCAAAATCATCCAgTAAACCgCC

Optix4_HCR_P1B2: CCTCgTAAATCCTCATCAAACCGCTCTCCTCCAGCGTCTCGCACA

Optix4_HCR_P2B2: AAGAAGCGCGCCAGGCGCTCCACGTAAATCATCCAgTAAACCgCC

Optix5_HCR_P1B2: CCTCgTAAATCCTCATCAAAGCGGTGGCGGCCGGCGTGGAAGGCG

Optix5_HCR_P2B2: GTGGCGCTCGAGGATGGCGTACAGCAAATCATCCAgTAAACCgCC

Optix6_HCR_P1B2: CCTCgTAAATCCTCATCAAAACAGCGCTTGCAGCTTGGCGTGGCT

Optix6_HCR_P2B2: CAGCCTCCTGGTAGTGCGCCTCCAGAAATCATCCAgTAAACCgCC

Optix7_HCR_P1B2: CCTCgTAAATCCTCATCAAAACAGTGCGTCTTCTGCTCGCCGTCC

Optix7_HCR_P2B2: GAGTAGAGATCGCGTCCGCTCCTTGAAATCATCCAgTAAACCgCC

Optix8_HCR_P1B2: CCTCgTAAATCCTCATCAAACGTCGGGTTCGGGTAGGGATCTTGG

Optix8_HCR_P2B2: CGTCGCCGCCGCCAATTCCCTCTTCAAATCATCCAgTAAACCgCC

Optix9_HCR_P1B2: CCTCgTAAATCCTCATCAAATTTGAACCAGTTGCCGACTTGCGTC

Optix9_HCR_P2B2: GGCCGCTCGGTCTCTTTGCCGTCGGAAATCATCCAgTAAACCgCC

***>spalt***

lcl|XM_024083373.2_cds_XP_023939141.1_1 [gene=LOC112046662] [db_xref=GeneID:112046662] [protein=homeotic protein spalt-major isoform X3] [protein_id=XP_023939141.1] [location=154..3339] [gbkey=CDS]

ATGCCGCGCGTCAAGCCCGCCTGCGTCCGCCGCGTCTCCATCGGTGAAAGCTCGGGATCTTGTTCGGAGGAAGATGTTGGCAATGCCATGCCGGATGAAGCGAGAGATAGGCCAGAGGCGCACATGTGTCCACGCTGTCAGGAACAGTTCGAAAACCTTCACGATTTCTTGTATCATAAGCGACTTTGCGATGAGAAAGCAATGCAAATGGGTGAAGAGAGGATGCACTCCGATCCAGAGGATATGGTAGTGTCGGGGGATGAAGAGATGGATGGTCCCAATAAACGACTAGAACAAGTCAGGAGGCATCGACAAGATGCTGAAAATAATAATAGTCTCGAAGACGGCGAGGCCGAAATACCTGAAGCCGACATGCCCCCCGTGGGCTGCCGTTCCCTTTGGCAGGACACGTTACTCTTGAGGCTCTACAAAATACGAGAGTAGCGGTCGCCCAATTCGCTGCAACAGCGATGGCAAATAATGCGAATAACGAAGCTGCTATACAAGAATTACAAGTGTTACACAACACTCTATACACTTTACAGTCACAACAAGTATTTCAACTTCAGTTAATACGTCAGCTTCAGAATCAGTTATCTCTAACTCGACGGAAAGAAGACGATCCACACAGCCCACCGCCAAGTGAACCAGAACAGAATGCCCCGTCAACGCCGGCTCGATCACCGTCGCCGCCGCGTCCGCCACGGGAGCCGTCGCCTGTTATACCCTCTCCTCCTACTAGCCAAAGTTTGCCGTCGACTCACACACATCACACACCCAAAACTGAACAGATATCTATCCCTAAGATTCCAACTTCCTCACCATCTTTAATGACCCACCCACTTTATAGTTCAATTTCTTCGTCATTAGCATCTTCCATCATAACAAACAATGATCCTCCACCGTCCCTAAATGAACCAAACACACTTGAAATGCTTCAAAAACGGGCACAGGAAGTACTCGACAATGCATCACAGGGCCTTCTAGCAAACAATCTTGCCGACGAATTAGCTTTTCGAAAATCCGGAAAAATGTCACCTTATGATGGAAAAAGTGGTGGCCGTAACGAACCTTTCTTTAAACATCGCTGTAGATATTGTGGAAAAGTGTTCGGTAGCGACTCTGCACTTCAAATTCACATTCGTTCTCACACAGGGGAAAGACCTTTCAAATGTAACGTCTGTGGCTCTCGATTTACAACCAAAGGAAATCTTAAAGTTCATTTCCAAAGGCATACTTCGAAATTTCCACATGTCAAAATGAACCCTAATCCCGTTCCAGAACATTTGGATAAATATCACCCACCGTTATTAGCGCAATTGTCGCCGGGGCCCATTCCTGGAATGCCGCCACATCCACTTCAGTTTCCCCCAGGAGCCCCAGCTCCCTTTCCGCCAAACTTGCCATTATACAGGCCACCGCATCACGATTTATTGCCTCCACGCCCTCTGGGTGATAAGCCTCTCTCACATCACCCACTTTTTGCTATGCGAGAAGAACAAGACGCACCAGCTGATCTCAGTAAACCTTCCGCACCAAGCCCTCCTCGACCCGCGTCTGATATTTTTAAGTCTGAACCTCAAGACGAAGAGAGTCAACGAGATTCCAGTTTTGAAGAGACTGATCGTATATCACCTAAGCGAGAAATCGAAGACAATGATATAGGACAAGATGCAGAACAAGATCGATACCCATCCACATCACCGTACGATGACTGCAGTATGGATTCCAAATACAGCAATGAAGATCAAATCGGCAGAGATAGTCCACACGTGAAGCCCGATCCAGATCAACCGGAAAATCTCTCAAGTTCGGAGAGCGGGCGGAGTGCACGGGGGTCGCCACCGTCGCCGTCGCCGTCGCCGTCGGCGCTGTCCACGCCGCCGCGTCTGCCGCACCACTCGCCGCTGCCGTCGCCCCCGACGCCCCTGGCGGCGCTCGGCGCGCTCGGCGGATCGCCCTTCAGCCCGCTCGGACTTGCCTTTCCTCCCGCAGTGCGCGGCAACACAACGTGTACCATCTGCTACAAGACATTCGCCTGCAACTCGGCACTGGAGATCCACTATCGAAGCCACACCAAGGAACGGCCATTCAAGTGCACCGTCTGCGATAGAGGCTTTTCTACCAAGAGCAGTGGCGGCGGTTGTCAGTGCGGAAGGCGTGCGCGCGCACCCCGCCCGCCGCACGCCACTGCTTTGGACCTCTGGAACGCCTTCGTCTACCCGGGCAACATGAAGCAGCACATGCTAACGCACAAGATCAGAGACATGCCGCCTGGTTTTGACAAGGGGCCGGGAGGACCTTCCGGACCCCCAAGCGAGGAAGGGCGGGACCCCAGCCCGGACAGACGGTCGTCCCCAGAAAAGCTGGATCTGAAAAGATCACCCCCGGTGCATCCTCCACCGCCAATGTCACACCCACCTATTGACATGCCACCTCTACCAAAAAGACCTACAGTGCCCAGTATCCCGAGTCACCCCCCACCGTCGCGTCGTCGAAGCACCTGTGCGGCGTGTGTCGCAAGAACTTCTCCTCATCATCAGCGCTGCAGATACACATGCGCACGCATACCGGAGACAAACCCTTCCGATGTGCTGTCTGTCAGAAGGCGTTTACCACCAAAGGCAATCTTAAGGTGCACATGGGCACGCACATGTGGAGCGGCGGCGCGTCGCGGCGCGGGCGGCGCATGTCGCTGGAGCTCCCGCCGCGGCCGCTGCACGAGCCGCACGAGCTGCTGCGGCGCCCCGACCTCTTCTACCCCTACCTGCCGGCGCCTTTCCTCAACGGCATGCAACAGAAGCTGAACGAGATATCTGTAATACAGCAGAACGCCGGACAAAACGGCGTAGCTGGAAAATTCCCCGGTCTGCTCGGCTTCGGAGCGTTCGGGGCCGGGAGACCGGGCGCCGCGTCCCCGCTCGAGAGGCCTCCCTCGCTGGAGGGGGGAGACGAGCGACAGGCGGCGATGCGTGAGCTGGCCGAGAGGGGACGGGAGCTGGCGGAGAGGAGTCGGCAGATGCGCGAGGAGAGCGAGCGGGAGCACTACAGGGCCGCGGGCGGACTGCCCGCGCACGCGCACGCGCCCAACCCCGCGCAGGCCTCGCCGCCCGCGCCGCACGCGCACCCGCACCCCCTCGCGTCGCTGCCGCCGCCCGCGCGGACAGAAGGCCTCACCGTATAA

spalt1_HCR_P1B1: gAggAgggCAgCAAACggAAATCCGGCATGGCATTGCCAACATCT

spalt1_HCR_P2B1: GTGCGCCTCTGGCCTATCTCTCGCTTAgAAgAgTCTTCCTTTACg

spalt2_HCR_P1B1: gAggAgggCAgCAAACggAAGTGCATCCTCTCTTCACCCATTTGC

spalt2_HCR_P2B1: CGACACTACCATATCCTCTGGATCGTAgAAgAgTCTTCCTTTACg

spalt3_HCR_P1B1: gAggAgggCAgCAAACggAAATTTCGGCCTCGCCGTCTTCGAGAC

spalt3_HCR_P2B1: CCCACGGGGGGCATGTCGGCTTCAGTAgAAgAgTCTTCCTTTACg

spalt4_HCR_P1B1: gAggAgggCAgCAAACggAACGTTATTCGCATTATTTGCCATCGC

spalt4_HCR_P2B1: ACACTTGTAATTCTTGTATAGCAGCTAgAAgAgTCTTCCTTTACg

spalt5_HCR_P1B1: gAggAgggCAgCAAACggAAGTGTGGATCGTCTTCTTTCCGTCGA

spalt5_HCR_P2B1: CTGTTCTGGTTCACTTGGCGGTGGGTAgAAgAgTCTTCCTTTACg

spalt6_HCR_P1B1: gAggAgggCAgCAAACggAAGAGTCGACGGCAAACTTTGGCTAGT

spalt6_HCR_P2B1: GTTCAGTTTTGGGTGTGTGATGTGTTAgAAgAgTCTTCCTTTACg

spalt7_HCR_P1B1: gAggAgggCAgCAAACggAAAGGATCATTGTTTGTTATGATGGAA

spalt7_HCR_P2B1: TGTGTTTGGTTCATTTAGGGACGGTTAgAAgAgTCTTCCTTTACg

spalt8_HCR_P1B1: gAggAgggCAgCAAACggAAATTTTTCCGGATTTTCGAAAAGCTA

spalt8_HCR_P2B1: CCACCACTTTTTCCATCATAAGGTGTAgAAgAgTCTTCCTTTACg

***>decapentaplegic (dpp)***

lcl|XM_052883655.1_cds_XP_052739615.1_1 [gene=LOC112044871] [db_xref=GeneID:112044871] [protein=protein decapentaplegic] [protein_id=XP_052739615.1] [location=321..1424] [gbkey=CDS]

ATGCGTGGGGCGTGCGCGTGCGCGGTGGTGTGCGCGTTGGTGGCGCTGTGCGCGGCGCGGCTGGACGAGTCCGCGCGCGCCGCCGCCGAGAAGCAGCTGCTGGCACTGCTGGGCCTGCCGCGCCGGCCGCCGCCGCGCGCCCGCCCGCCGCCCGTGCCGCGCGCGCTGCGCGTGCTGTACGACTCGCGCGCGCTGCCCGCCGCCGCCGCCAACACGGCGCGCTCCTTCCACCACACGCCCACGCCGCTCGACGAGCGCTTCCCCGGCGACCACCGCTTCCGCCTGTTCTTCAACGTGAGCGGCGTACCGGCCGACGAGGTGGCGCGCGGCGCCGACCTCTCGTTCCAACGAGCCGTCGGCACCACCGGCAGACAGAGACTGTTGTTGTACGACGTGGTGCGCCCCGGCCGCCGCGGCCACTCCGAGCCGATCCTGCGGCTGCTGGACTCCGTGCCGCTCCGGCCCGGGGAGGGAATCGTCAACGCCGACGCTCTGGGAGCGGCGCGACGGTGGCTCAAAGAGCCCAAACATAATCACGGACTATTAGTGCGAGTGTTAGAAGAAGACGCCGCGAGTGTGAGCAGGGACGCGAAGTTCCCGCACGTGCGCGTGCGCAGACGCGTCACGGACGAGGAGGAGGAGTGGCGGACGGCGCAGCCGCTGCTCATGCTGTACACGGAGGACGAGCGCGCGCGCGCGTCGCGGGAGACGAGCGAGCGGCTGACGCGCAGCAAGCGCGCGGCGCAGCGGCGGGGGCACCGCGCGCACCACCGCCGCAAGGAGGCGCGCGAGATCTGCCAGCGCCGCCCGCTGTTCGTCGACTTCGCGGACGTGGGCTGGAGCGACTGGATCGTGGCCCCGCACGGCTACGACGCGTACTACTGCCAGGGCGACTGCCCCTTCCCGCTGCCGGACCACCTCAACGGCACGAACCACGCGATAGTGCAGACTCTGGTCAACTCAGTGAACCCCGCGACGGTGCCCAAAGCGTGCTGCGTGCCGACGCAACTCTCATCTATATCTATGTTATATATGGACGAAGTGAACAATGTGGTGCTTAAAAACTATCAGGACATGATGGTGGTAGGCTGTGGCTGCCGATGA

Dpp1_HCR_P1B1: gAggAgggCAgCAAACggAAACCTCGTCGGCCGGTACGCCGCTTA

Dpp1_HCR_P2B1: TGGAACGAGAGGTCGGCGCCGCGCGTAgAAgAgTCTTCCTTTACg

Dpp2_HCR_P1B1: gAggAgggCAgCAAACggAACAGTCTCTGTCTGCCGGTGGTGCCG

Dpp2_HCR_P2B1: GCCAGGGCGCACCACGTCGTACAACTAgAAgAgTCTTCCTTTACg

Dpp3_HCR_P1B1: gAggAgggCAgCAAACggAACAGCCGCAGGATCGGCTCGGAGTGG

Dpp3_HCR_P2B1: CCCGGGCCGGAGCGGAACGGAGTCCTAgAAgAgTCTTCCTTTACg

Dpp4_HCR_P1B1: gAggAgggCAgCAAACggAAGCCGCTCCCAGAGCGTCGGCGTTGA

Dpp4_HCR_P2B1: TGTTTGGGCTCTTTGAGCCACCGTCTAgAAgAgTCTTCCTTTACg

Dpp5_HCR_P1B1: gAggAgggCAgCAAACggAACGCCACTCCTCCTCCTCGTCCGTGA

Dpp5_HCR_P2B1: TACAGCATGAGCAGCGGCTGCGCCGTAgAAgAgTCTTCCTTTACg

Dpp6_HCR_P1B1: gAggAgggCAgCAAACggAATCTCCCGCGACGCGCGCGCGCGCTC

Dpp6_HCR_P2B1: GCTTGCTGCGCGTCAGCCGCTCGCTTAgAAgAgTCTTCCTTTACg

Dpp7_HCR_P1B1: gAggAgggCAgCAAACggAAGCCCACGTCCGCGAAGTCGACGAAC

Dpp7_HCR_P2B1: GTGCGGGGCCACGATCCAGTCGCTCTAgAAgAgTCTTCCTTTACg

Dpp8_HCR_P1B1: gAggAgggCAgCAAACggAAGGGGCAGTCGCCCTGGCAGTAGTAC

Dpp8_HCR_P2B1: GCCGTTGAGGTGGTCCGGCAGCGGGTAgAAgAgTCTTCCTTTACg

Dpp9_HCR_P1B1: gAggAgggCAgCAAACggAACACTGAGTTGACCAGAGTCTGCACT

Dpp9_HCR_P2B1: GCACGCTTTGGGCACCGTCGCGGGGTAgAAgAgTCTTCCTTTACg

Dpp10_HCR_P1B1: gAggAgggCAgCAAACggAAATATAACATAGATATAGATGAGAGT

Dpp10_HCR_P2B1: AAGCACCACATTGTTCACTTCGTCCTAgAAgAgTCTTCCTTTACg

***>optomotor-blind (omb)***

>lcl|XM_052882932.1_cds_XP_052738892.1_1 [gene=LOC112048640] [db_xref=GeneID:112048640] [protein=optomotor-blind protein isoform X2] [protein_id=XP_052738892.1] [location=168..1883] [gbkey=CDS]

ATGCATCATCTCGAGAATTTCTCAATAAGCCGTGGTGCCTGTGGGGCCGACGCGGGCGCCGCCCCGCGGCGAGTCGGTGGCGCCTCTATCGATTATTCGGGCCTCTCGGTCGCGCGTAGCGCCGCGTCCCGCTCGCGCCGCGCGGGTGGCGCCCGCCGCGTCATGCGCGACCACCGCGCGACTGGCGCGACCTCGCGCGAGCCGCTCTCGGGCTATATCATTAATCTATTACTCCGGGAAGGTGGGCTCGTACGTAAGGAGCTCATCTATAGCCAAAGTCAACGGATGTTGTCAAAGTTGCTGTGGCCAAACCCAGTGCCTAGTCCGCCCAGAGACCCCCACTCAGATCATTTGACGATCCTGAACTCGATGCACAAGTACCAGCCACGGTTCCACCTGGTGCGAGCCAACGACATCCTCAAGCTGCCCTACTCCACCTTCCGCACCTACGTATTCAAGGAGACCGAGTTCATCGCCGTCACCGCCTACCAGAACGAGAAGATAACGCAGCTGAAAATCGACAACAACCCCTTCGCGAAAGGCTTCCGAGACACGGGGGCGGGGAAGCGGGAGAAGAATCTGTCCGTGTACCGCAGGCAGGCGCTGCTGACGGCGCGGTCGGACGCCCGCGAGGACGACGACGAACGTCCACTAGACGTTGGCGGACCCTCCAGCCCGCCGCCGCCCGCGACGCAACACACGAGCAGCTCGTGGTTCAGTTCGAGTGGAGGCGGCGCGGACTCCGGTCCAGAGGAAGCCGGCTCGGACTCGTCGTGCTCCGGCCCCGCGCGCGCTCCCTCGCCCCCCGCCGGGCCCTCGCGGCCTTCTCCCGCCCCTGACGTGTCTCTCGGCCCGCCGGTGCAGCCCCCCCTCCTGCCCTACCTGTACCCGCCTTCGCTGTACCCGCCACCGTTCTTCCCGCCACACCAAATGCCGCCAGGTCTGTTGTTCAACCTGCATCCCCTCCTGCAGCAGTACTCGTTGCCCCCTCCCCTCGCGCCGCCGACCCCCACGTCCGCGCCTTCTCTGAGCAAGACGCACAGGTTCGCGCCGTACGCGCTGCCCGGGTTAGGGTCTGCGTTCGAGCAGGTCGCGCCCAGAGCGAGGAGCCTCAGTTCGTCGCCGGCGCGGCCGCGGGTGGGGTCGCCTCCCACGAGAGCGGCGTCAGCGGACCCCCCGCCGGACGCGCCGACGTCGACCACCTCCGCCGCGACCCCTCCCGCGTCCGACCTCAAGAGCATCGAGCGTATGGTCAACGGCTTGGACGTGGAGACGCAGGACTGA

omb1_HCR_P1B1: gAggAgggCAgCAAACggAACGAATAATCGATAGAGGCGCCACCG

omb1_HCR_P2B1: CGCGGCGCTACGCGCGACCGAGAGGTAgAAgAgTCTTCCTTTACg

omb2_HCR_P1B1: gAggAgggCAgCAAACggAACATGACGCGGCGGGCGCCACCCGCG

omb2_HCR_P2B1: GGTCGCGCCAGTCGCGCGGTGGTCGTAgAAgAgTCTTCCTTTACg

omb3_HCR_P1B1: gAggAgggCAgCAAACggAACTTTGACAACATCCGTTGACTTTGG

omb3_HCR_P2B1: ACTAGGCACTGGGTTTGGCCACAGCTAgAAgAgTCTTCCTTTACg

omb4_HCR_P1B1: gAggAgggCAgCAAACggAACAGGATCGTCAAATGATCTGAGTGG

omb4_HCR_P2B1: CCGTGGCTGGTACTTGTGCATCGAGTAgAAgAgTCTTCCTTTACg

omb5_HCR_P1B1: gAggAgggCAgCAAACggAACGTTCTGGTAGGCGGTGACGGCGAT

omb5_HCR_P2B1: TGTCGATTTTCAGCTGCGTTATCTTTAgAAgAgTCTTCCTTTACg

omb6_HCR_P1B1: gAggAgggCAgCAAACggAACCGCCCCCGTGTCTCGGAAGCCTTT

omb6_HCR_P2B1: ACACGGACAGATTCTTCTCCCGCTTTAgAAgAgTCTTCCTTTACg

omb7_HCR_P1B1: gAggAgggCAgCAAACggAAGTCGCGGGCGGCGGCGGGCTGGAGG

omb7_HCR_P2B1: CTGAACCACGAGCTGCTCGTGTGTTTAgAAgAgTCTTCCTTTACg

omb8_HCR_P1B1: gAggAgggCAgCAAACggAACTTCCTCTGGACCGGAGTCCGCGCC

omb8_HCR_P2B1: GGCCGGAGCACGACGAGTCCGAGCCTAgAAgAgTCTTCCTTTACg

omb9_HCR_P1B1: gAggAgggCAgCAAACggAAGCGGGTACAGGTAGGGCAGGAGGGG

omb9_HCR_P2B1: GGAAGAACGGTGGCGGGTACAGCGATAgAAgAgTCTTCCTTTACg

omb10_HCR_P1B1: gAggAgggCAgCAAACggAAATGCAGGTTGAACAACAGACCTGGC

omb10_HCR_P2B1: GGGCAACGAGTACTGCTGCAGGAGGTAgAAgAgTCTTCCTTTACg

***>aristaless (al)***

lcl|XM_024097138.2_cds_XP_023952906.1_1 [gene=LOC112056678] [db_xref=GeneID:112056678] [protein=homeobox protein aristaless] [protein_id=XP_023952906.1] [location=244..1032] [gbkey=CDS]

ATGGGAGTATCAGAACCAAACTGCTCCTCAACTCCCGACCTTCCCCCTCACGACCCGGAGCGACCCGGTTCAGGCAGCGGCATGGATGACGAGGACATTCCCAGAAGGAAGCAGAGGCGGTATAGGACCACCTTCACTAGCTACCAGCTCGACGAACTGGAGAAGGCTTTTGGGAGGACGCATTATCCTGATGTATTTACCAGGGAGGAGTTAGCTTTGAAAATTGGCCTCACAGAAGCAAGAATACAGGTGTGGTTCCAAAACCGGCGGGCGAAATGGCGTAAGCAGGAGAAGGTGGGGCCCCATGCACATCCGTATAGCGGATACTTAAGCAGCGGGCAGCCATTACCGACTACATCTATGCCAGTTCCACCGCATTCGTTCAGTCAGCTCGGATTTGGATTGAGAAAGCCTTTTGACAACGCTCTAGCGTCTTTCAGGTACGCCAACAGTCCTCTGTTTGGAGCACAGTACCTTCCGCCGCTTTCTCGTCCTCCACTTTTCGGGGCTCCACTATACGCTACATCGCCCGCTCATTTCCACTCTCTGTTCGCCAATCTAACCGTACCCGAACTACCTCGAATTTCACCTGAACAATCACGATTGTCACCCGAAGTAACTCGATCGCCAGCACCGTCGATCTCCCCTCCAATATCACCAGGCAGCGAGACTTTACCCCCATCTGAAGATGTTAGGAGTTCCAGCATAGCAGCATTAAGATTAGCTGCTAGAGAACACGAACTTAGATTAGAATTGTTGCGACAGCGAGCGGATTTAATTTGTCAATAG

Al1_HCR_P1B1: gAggAgggCAgCAAACggAAAGTTGAGGAGCAGTTTGGTTCTGAT

Al1_HCR_P2B1: CTCCGGGTCGTGAGGGGGAAGGTCGTAgAAgAgTCTTCCTTTACg

Al2_HCR_P1B1: gAggAgggCAgCAAACggAAGAATGTCCTCGTCATCCATGCCGCT

Al2_HCR_P2B1: TCCTATACCGCCTCTGCTTCCTTCTTAgAAgAgTCTTCCTTTACg

Al3_HCR_P1B1: gAggAgggCAgCAAACggAACTTCTCCAGTTCGTCGAGCTGGTAG

Al3_HCR_P2B1: ATCAGGATAATGCGTCCTCCCAAAATAgAAgAgTCTTCCTTTACg

Al4_HCR_P1B1: gAggAgggCAgCAAACggAAGAGGCCAATTTTCAAAGCTAACTCC

Al4_HCR_P2B1: GAACCACACCTGTATTCTTGCTTCTTAgAAgAgTCTTCCTTTACg

Al5_HCR_P1B1: gAggAgggCAgCAAACggAAACCTTCTCCTGCTTACGCCATTTCG

Al5_HCR_P2B1: CCGCTATACGGATGTGCATGGGGCCTAgAAgAgTCTTCCTTTACg

Al6_HCR_P1B1: gAggAgggCAgCAAACggAAATAGATGTAGTCGGTAATGGCTGCC

Al6_HCR_P2B1: TGACTGAACGAATGCGGTGGAACTGTAgAAgAgTCTTCCTTTACg

Al7_HCR_P1B1: gAggAgggCAgCAAACggAAGGAGGACGAGAAAGCGGCGGAAGGT

Al7_HCR_P2B1: GTAGCGTATAGTGGAGCCCCGAAAATAgAAgAgTCTTCCTTTACg

Al8_HCR_P1B1: gAggAgggCAgCAAACggAATACGGTTAGATTGGCGAACAGAGAG

Al8_HCR_P2B1: TTCAGGTGAAATTCGAGGTAGTTCGTAgAAgAgTCTTCCTTTACg

Al9_HCR_P1B1: gAggAgggCAgCAAACggAAGGTGCTGGCGATCGAGTTACTTCGG

Al9_HCR_P2B1: CCTGGTGATATTGGAGGGGAGATCGTAgAAgAgTCTTCCTTTACg

Al10_HCR_P1B1: gAggAgggCAgCAAACggAAGGAACTCCTAACATCTTCAGATGGG

Al10_HCR_P2B1: AGCAGCTAATCTTAATGCTGCTATGTAgAAgAgTCTTCCTTTACg

***>cubitus_interruptus (ci)***

lcl|XM_024083241.2_cds_XP_023939009.1_1 [gene=LOC112046560] [db_xref=GeneID:112046560] [protein=transcriptional activator cubitus interruptus] [protein_id=XP_023939009.1] [location=602..5380] [gbkey=CDS]

ATGCCTGATCGGGAGTCAGTTGGTGGTCCTGGGACTGCGGGAAGCGCTGGTTTTCTACCACTACAGTTTCCGTCTGCTTTCGCCGCATTCCATGCAACTACTCCTCCAGGGGTGCCAGCGACGGCGATGCATCACGCAACACACTACCATCATCATGCGCAGTTGGCAGCGGCGGCGGCGGCGGCAGCGGGCGCTACCAGCGAGCTCAGCTACCTCGCCGCTCTCCACCCAGCGTACCGGCCCGTACCCTACGACCATCCGCTGTACGGCGCTAACACTTTACGAGGCTTAGAATATTTAAGTGCAGCGAGGAGTCTACATCCAGAACTTCATGCAGGCAGTACGTTAGCGAGTCAAGAGTTCCAACTCAGTTTAGAAGGATCAAGAATAGCCTCTCAGAACCGTCTAAGGTTATCCGGTGGGGCGATCAGTGCGTCAGCTAACAGGAAGAGGGCGGTTTCCTGGAGCCCCTACTCGGCGGAGTCCCTGGACTTGGCTGCTGTGATACGCGCGTCCCCGGCCAGCCTCGCTGTGAGGGCCCCTTCTGCGGCCTCTACTGGCAGCTATGGACATCTCAGTGCAGGAGCAATATCCCCAGCGCTGTCGCTATCGCATGCATCGTTGGCGCAACAGCTTCTGGCTCGCGGCGGCAGCGGCGTCATCCCCGGCGGCGTGCTGCTCGACCCCGCGCACCAGCAGGCAGCAGCGGCGGCGGCACACCACGCCGCGCACGCGCATCTGGTGGCCGGCATACACAGATCTCACATATCGTCACCGACTCAGCTGCTGATCGCGCCGGTGGACGTGCGGCCCGGCCTGGGGCTGGACGGCACGCCCCCTCACATGCAGCAACCGCATCAGCAACCTGAAATCACCAGTGTTATGGAGGCTGATAGTGCATCAACGGCTCTCAACCAGCGCAAGTCCCCACAAGTGCTGATATCACACCGAGAGAACATGAACAGCAACAAGCCATTGTCGGCGGCGGCAGAGAGCACGGTCCACGACGGGCTGGACTCTAAGGATGAGCCTGGAGACTTCATTGAGACTAACTGCCACTGGGTGGACTGTAAGCTGGAGTTCCCAACACAGGACGATCTAGTAAAGCACATCAACACAGACCACATCCACGCCAGCAAGAAAGCCTTCGTCTGCCGCTGGGTCAGCTGCTCCAGGGACGAGAAGCCTTTCAAAGCGCAGTACATGCTTGTGGTGCATATGAGACGCCACACTGGAGAAAAGCCTCATAAGTGTACATTCGAGGGATGCTGCAAAGCTTACTCCCGGCTCGAGAACCTGAAGACCCACCTGCGGAGTCACACCGGGGAGAAGCCCTACACGTGCGAATACCCCGGCTGTGCGAAGGCTTTCTCCAACGCCAGCGATCGGGCGAAGCATCAGAACAGAACTCATAGTAATGAGAAACCGTACGTATGTAAGGCTCCAGGCTGCACGAAGCGGTACACAGACCCGTCCTCCCTCCGCAAGCATGTGAAGACTGTACATGGTGCTGAGTTCTACGCCAGCAAGAAACATAAAGGGTGCAGTCGAGGAGACGATTCAGCCGAGTCTGGGGGTGGTGGCGCGGGATCTTCGCCACGGTCGGAGGAGGGTGGAGTTCCTATGGGGGTGAGGGGTCACACGTCCTCGGCCTCTGTCAAGAGTGAGAGCCCAGCTTCACCCTTACCTCATGGTCTACATACACCGGCTCATCAGTTATCAGCACAATGTGGTGGTGAACTGGACTTCGGTGTGTCTGGTTTGGGCGGGTTTAGTGATGAAAACGGCGCTCCCTACTTCAGACTAGATGGCGAGGTGGAACAGGAAGTGGTAGGGGAAGTGGGACAACTGCCCCTCATGCTGCGCGCTATGGTAGCGATAGGGGAGCCGCGCGCCCATCATCACGCGCCGCGCTTCGGACACAAGATGGGAGTGGGGAGACTGATGCCGCCCATACACGGGGCAGATGTTGGTGGAGGTGGTATTCAAGGGCGGACAGAAATAGGAGGTACAAACGTAGCGGTAGAACTGAAGACCGGACTGCCCAACACGAGGCGAGACTCTGGAATCTCTTCTGGAAGCAGTTTGTATAGTGCTAGATCGTCGGATATCTCGCGCAAGAGCAGCCAGGCGTCAGTGGTGTCGGGCGCGGTGGTCACCACCACAGGCGTGGCCGGGCAGCAGAGGCTAGTCGCGCAGCATGCGGCCATTTATGACCAACTGTCACCTGACAGCAGTCGGAGATCTAGTCAAGTTTCCTGCGTAGGCTACGCCCCGCCGCCGTCGTCAGCATTGGCTGCAGTTCAAGCTGTAAGGACCTCACAAGGCAATCAGGCTGTACTGCTTCGAGGAGTCACGTGTTCAGAAGTGAGAGCAGAAGAGTTAGCCCTAGAACTGGATCCTAACGTGCAAGTGAAGGAGGAAGCCAGGAGATTGTCAGAGCAATCCAACCTCAGTGACCAGGCTCAGGGATATCAGCCATATCCATGCACCAATGACGATGTGGGTGATGCTCCCTTCCCATTCAAGACAGAAGATGATCATGTGATATCGTATAAAGAATCTCGATCAAATTCCACAAACACTGTGGTTATCACCACAGCACAAGTTCATCACCCAAATCAAGAAGTCAATCTTGAACAGGTCGCGGAAGGAGAAATGGTGGAGAACAAGCTGGTGATACCAGACGAAATGATGCAATATCTTAACCAATCAATATTGGGAACCGACTCGGTAACACCAGCAAAAACGGACTCCAGCAATCCTGACCCAGAAGGAGCAAACAAAGAAAAAGATACAACAACCAAAGACATTTTATCCAGTGAAATACACCATACAAACAATTCTAGCGACAAAATCAGTGATGTTGCGACGAGTGACGATTCCCTTCTCAAAAATTTAGGTGCCATTGGAAGTGATCTCAACATAAGTGATATTCAAGTTGATTTAAGGTCATTAGATGTTAGTATGTCTGGTAACAGCGGCTCTCTATTAGCCTCCAAGTCTTTAGATGATAAGAACCCCCCTCTTTCGGAACAAGTTATACCGGAAGTTACACCAGAAACATGTGAACAAGAATTCTCAAAACCCCCGACAAGTAATGTTGTTACAAGCAACCCTTTGCAGTCTTTACAAACTATGACAGCAAATCAAACAGATCAATCCAATAGAATGCGAGTCAACAACCCCCTACCTCAAAAGCCCGCTATGAGTCCAAAAACAGTTGTCATGACACAAACTATCATGAGTCCCAGTTTAGCACATAACATGTTAAGTCCTCAGAGTCTACCGCATAGTTCAATGAGCCCACAGAGCATCAGAAGCCCTCAGCATATGCCTCAAAGCATCATGAGCCCACCAAGCGTATACAATGTCATGAGTCCGCAAAGTGTCATGAGCGTCATGTCTCCACAGCACAATGCAATGAGCCCACAAAGTATGCAGAGTTTATTGAGCCCTCAGATGCCAAATCAAATGATAATGAGTCCCCGACACAACAACATTGGAAGTCCTCATTCCCAGAACATCGCAAGTCCCATGATGAACATGGCAAGTCCAATGACGCAGAACATTGCTAGTCCGATGAGTCATGGCATGCCAAGTCCAATGCATCCTGGCCTTCAAAGCCCAATAACAAGTCCTATGGTACAAAACATGTCCAATATGACGATGCAACCAGTGCCAAATCAAAATGCGAATATAATGATGAGTAACATGAATCCTCCTCAACAAATGACGGCAGCACCACCCTATAATAACCGTCCCAACTGTCCTCCAAAACTTCCAAACAAAAACTTTAGTAACGTTCCTAATCAATACCAAAACCAAAACTACGGCCCAGCGCCACCTTATCCCATACAAAACCAAGTCAATGTAAACATGAGAAACCAAAACATTCAACAGTACCAGATGATGCAACACTATAATCCTAATCAACCACAGGTGACAATGATGCAAAACAACCATCAAAACATGGTTTATAATAATCAAATGACAAATTACGTTCAACCGATGAATTATCCAAATCAAAATGCTCAGATGCATCAAATGCAGTTATCACGATCTTCAGTAATGAGCGTCGATAACAGTGGAAACATGAGTCGTGGTGCTATGAACAGTTACTGTGAACAACAGAATGTATGTCCTCCCATGCAAAACGTACAATATAACCAGAACATGCAATATCCTCAACCGCCTCCATACAATTCTGTAGCGAACGCTGCTAACGTAATGGGTCCTCCACCACCAAAGAATAACCATCAGTACAACCAGGCAATGATGAACAATAACCAATACTACAATCATCAGAGATCTTACAACCAATGGGACTATCCTGGCAACCAATTCAACAAGCACAACATGCAAAAATCTGGTCAGAATTCAGTCAATATGTCTACAGGTAGTCAAAAACCTATAAACGGTGCGAGAATGTCAATGAATTGCAATCAGATGGTCAAAAATGCTGCGGAGCAACAAGCGGACTGCAGTATGAATAGTTTGAGAAGCCAAAACAACCAAACTGATGTTCAAGTTTGGGATATCTCTCAGTCCCAAATAGAGGCTACGAATGGAAGAAAGAAAAACCAAAATACCATGCGTCAGGAGACATATCAAAGGACCTTGGAATACGTAGAGAACTGTGAGAATTGGAAGAGTTCTGAAATGGTATCCAGTAGCACACATCCTCTACAGGGCGGTGACAATATGGTTGTGAACGATCTCCAAACTTCTCTATCTTCGTTTTATGAAGAAAATCAATATCTTCAGATGATACAATAG

ci1_HCR_P1B1: gAggAgggCAgCAAACggAAGTAGTTGCATGGAATGCGGCGAAAG

ci1_HCR_P2B1: ATCGCCGTCGCTGGCACCCCTGGAGTAgAAgAgTCTTCCTTTACg

ci2_HCR_P1B1: gAggAgggCAgCAAACggAAGCCGGTACGCTGGGTGGAGAGCGGC

ci2_HCR_P2B1: CGTACAGCGGATGGTCGTAGGGTACTAgAAgAgTCTTCCTTTACg

ci3_HCR_P1B1: gAggAgggCAgCAAACggAAATCCTTCTAAACTGAGTTGGAACTC

ci3_HCR_P2B1: TTAGACGGTTCTGAGAGGCTATTCTTAgAAgAgTCTTCCTTTACg

ci4_HCR_P1B1: gAggAgggCAgCAAACggAATGGCCGGGGACGCGCGTATCACAGC

ci4_HCR_P2B1: CCGCAGAAGGGGCCCTCACAGCGAGTAgAAgAgTCTTCCTTTACg

ci5_HCR_P1B1: gAggAgggCAgCAAACggAAGATGACGCCGCTGCCGCCGCGAGCC

ci5_HCR_P2B1: CGCGGGGTCGAGCAGCACGCCGCCGTAgAAgAgTCTTCCTTTACg

ci6_HCR_P1B1: gAggAgggCAgCAAACggAATCCACCGGCGCGATCAGCAGCTGAG

ci6_HCR_P2B1: CCGTCCAGCCCCAGGCCGGGCCGCATAgAAgAgTCTTCCTTTACg

ci7_HCR_P1B1: gAggAgggCAgCAAACggAAATATCAGCACTTGTGGGGACTTGCG

ci7_HCR_P2B1: TGTTGCTGTTCATGTTCTCTCGGTGTAgAAgAgTCTTCCTTTACg

***>thickvein (tkv)***

lcl|XM_024079796.2_cds_XP_023935564.1_1 [gene=LOC112044074] [db_xref=GeneID:112044074] [protein=bone morphogenetic protein receptor type-1B isoform X2] [protein_id=XP_023935564.1] [location=205..1653] [gbkey=CDS]

ATGATGTACTGTACCAGCCCTGCGGCTACATCTGGCTGCGTGTCCGTCATCATGGCCGGTGGCCAAGTTTGCCTAGGTCGGGGCATCGTGTGCGAGTGCACGGGCGCGGGCATGTGCCCCGGCGGCGCGCCCAACGGCACGTGCGGCACGCAGCCCGGCGGGTACTGCTTCGTGGCCGTGGAGGAGCTCTACGACGACAGCGGGCTCGTGGTGCTGGAGCGCACCGCCGGCTGCCTGCCGCCCGACGAGTCGGGCCTCATGCAGTGCAAGAAAGTGCCCCACCAGAACCCGAAGGCGATAGAGTGCTGCGAGAAAGACTACTGCAACCGCCGCCTGCGGCCGCAGCTGCCGGAGCCCCCGCCCGACGTCACCGAGACCCCCGGCCTGCGCCCCGCGGGCTCCGTGCCGCACACGGCGCTGGTCGCCGCGGCGCTGTGCGCGGCCCTGCTCGCCTTCCTCGCGGCCTTCTGGCTGCTCTTCAGGATGCGCAGAAGAGGATGCAAGCGACCGCCTTCCCCGCCCGCCCCCGCGCACAGCTCGGAGATCTCCTCGGGCTCCGGGTCCGGCCTCCCGCTCCTAGTCCAAAGAACCGTCGCCAAACAGATACAAATGGTCGAGTCGATCGGCAAAGGTCGCTACGGCGAAGTCTGGTTGGCGAGATGGCGCGGCGAAAAGGTGGCCGTCAAAGTTTTCTTCACCACGGAGGAGGCTTCCTGGTTCCGCGAGACGGAGATATACCAGACGGTTCTCATGCGACACGAAAACATCCTCGGCTTCATCGCGGCGGACATCAAAGGAACGGGATCCTGGACTCAGATGCTTCTCATCACGGACTACCACGAGAACGGCTCCCTGCACGATTATTTGCAGACCGTCGTTCTGGACACGCAGGGTTTGATGACGATGGCGTACTCCATAGTGAGCGGGCTGGCCCACCTGCACATGGACATATTCGGCACCAAAGGCAAGCCCGCCATCGCTCACAGAGACATAAAGAGCAAAAACATCCTCGTCAAAAGGAACGGCCAGTGCGCGATCGCCGACTTCGGCCTCGCGGTCAGATACGTGGCGGAGAGGAACGAGGTGGACATCGCACCGAACACGCGCGTCGGCACGAGGCGGTACATGGCGCCCGAGGTGTTGGACGAGAAGTTGGACGTCACCAACTTCGAGGCGTTCAAAATGGCCGACATGTATTCTTTGGGACTGGTGATGTGGGAGATGTGCAGGCGGTGTACGACCGGGGACAAGGCGCAGTACGTGGAGGCGTACGCGCTACCGTACCACGAGCACGTGCCGTCGGACCCGTCGTTCGACGACATGCACGCGGTGGTGGTGGGCCAGCGCGCACGGCCGCCGCTGCCGGCGCGCTGGCGGGCGTCGCCCACGCTGCTGGCGCTGGCGGCGCTCATGGCGGAGTGCTGGCACCACAACCCGCCCGTGCGGCTCACGGCGCTGCGCGTCAAGAAGACGCTGGCCAAGTTCCGCGCCGAGAGCGCCGTGAAGCTCGTCTGA

tkv1_HCR_P1B2: CCTCgTAAATCCTCATCAAACGTGCACTCGCACACGATGCCCCGA

tkv1_HCR_P2B2: CGCGCCGCCGGGGCACATGCCCGCGAAATCATCCAgTAAACCgCC

tkv2_HCR_P1B2: CCTCgTAAATCCTCATCAAAGCGGTGCGCTCCAGCACCACGAGCC

tkv2_HCR_P2B2: CCCGACTCGTCGGGCGGCAGGCAGCAAATCATCCAgTAAACCgCC

tkv3_HCR_P1B2: CCTCgTAAATCCTCATCAAAGGGGCTCCGGCAGCTGCGGCCGCAG

tkv3_HCR_P2B2: GGCCGGGGGTCTCGGTGACGTCGGGAAATCATCCAgTAAACCgCC

tkv4_HCR_P1B2: CCTCgTAAATCCTCATCAAACTGCGCATCCTGAAGAGCAGCCAGA

tkv4_HCR_P2B2: GGGGAAGGCGGTCGCTTGCATCCTCAAATCATCCAgTAAACCgCC

tkv5_HCR_P1B2: CCTCgTAAATCCTCATCAAACGATCGACTCGACCATTTGTATCTG

tkv5_HCR_P2B2: ACCAGACTTCGCCGTAGCGACCTTTAAATCATCCAgTAAACCgCC

tkv6_HCR_P1B2: CCTCgTAAATCCTCATCAAATCGCATGAGAACCGTCTGGTATATC

tkv6_HCR_P2B2: CGCGATGAAGCCGAGGATGTTTTCGAAATCATCCAgTAAACCgCC

***>patched (ptc)***

lcl|XM_024091128.2_cds_XP_023946896.1_1 [gene=LOC112052161] [db_xref=GeneID:112052161] [protein=protein patched] [protein_id=XP_023946896.1] [location=293..4252] [gbkey=CDS]

ATGTACGACATCGAGTGGCGGCTCAAGGACCTCTGCTACAGCCCCAGCATCCCGGACTTCGAGGGCTACCACCACATCGAGTCCATCATAGACAACGTCATCCCGTGCGCAATCATCACGCCGCTCGACTGCTTCTGGGAGGGGTCCAAGTTGCTTGGTCCTGAATATCCTATATTTGTACCTCATCTAAAAAACAAACTACAATGGACTCATTTGAACCCGCTCGAAGTATTAGAGGAAGTGAAGAAGCTAAAGTTCCAGTTCCCTCTGAGCACAATGGAGGCGTACATGAAGAGGGCCGGCATCACGTCCGCTTACATGAAGAAGCCGTGCTTAGACCCCACCGACCCTCATTGTCCAGACACGGCGCCGAACAAAAAATCAGGACATATTCCAGATGTAGCGGCAGAGCTGTCCCACGGATGTTACGGTTTCGCGGCTGCGTACATGCACTGGCCGGAGCAGTTGATAGTGGGCGGAGCGACGAGGAACTCCACATCAGCTCTGAGGAGTGCGCGCGCCCTGCAGACCGTCGTCCAGTTGATGGGCGAGCGAGAAATGTACGAATACTGGGCCGACCACTACAAAGTGCACCAAATTGGTTGGAATCAAGAAAAAGCCGCCGCCGTACTTGATGCTTGGCAGAGAAAGTTTGCAGCTGAAGTAAAAAAGATGACTACCTCAAGTTCAGTGTCAGCAGCGTACAGCTTCTACCCGTTTTCGACCTCAACTTTGAATGACATACTCGGAAAATTCTCGGAAGTCTCACTAAAGAACATTATTTTGGGATACATGTTTATGTTAATTTATGTTGCTGTAACGTTAATACAATGGCGAGATCCAATTCGTTCTCAAGCTGGAGTGGGTATAGCCGGAGTATTGCTTCTGTCGATCACAGTAGCCGCTGGCTTAGGCTTTTGTGCATTATTAGGCATACCATTCAATGCATCGAGTACACAAATAGTGCCGTTCCTAGCTCTCGGACTAGGTGTTCAAGATATGTTCCTTCTCACTCACACATACGTTGAACAAGCGGGAGATGTGCCGAGAGAAGAGAGAACCGGACTGGTACTGAAAAAGAGCGGACTGAGCGTTCTACTGGCCTCTCTGTGCAATGTTATGGCGTTCTTAGCTGCAGCTCTACTGCCTATACCTGCGTTTCGGGTATTTTGTTTACAGGCGGCCATCCTCTTACTATTCAACCTGGGTTCAATGTTGTTGGTGTTTCCTGCGATGATCTCACTGGATCTCCGGCGTAGATCAGCGGCGAGAGCCGATCTATTGTGTTGTTTGATGCCAGAAAGTCCGCTTCCCAAAAAGAAAATTCCAGAGCGAGCTAAATCAAGGAGTGGAAAGACTGATAAGAACAATAGGTTAGACACGACTCGGCAACCATTAGATCCAGATGTGACGGGCGAGCAACCCAAAGCCTGTTGCCTCAGCGTGTCGCTCACCAAGTGGGTCAAGAACCAATACGCGCCCTTCATAATGCGCCCCGCTGTTAAGGTAACATCAATGCTAGCGTTGATAGTAGTGATTTTGGCGAGTGTTTGGGGAGCCACGAAAGTCAAAGATGGATTGGATTTAACAGACATAGTACCAGAACATACTGATGAACGCGAATTTCTTACGCGCCAAGAGAAATACTTCGGCTTCTACAATATGTATGCTGTGACGCAGGGCGACTTTGAATATCCCACGAATCAGAAATTGTTATACGAATATCACGATCAATTTGTTCGAATACCGAATATTATTAAAAACGACAACGGTGGACTCACAAAGTTTTGGTTGGGTCTATTTCGCGACTGGTTACTGGATTTACAGGATGCATTCGACAAAGAAGTAGCCAGTGGCTGTATCACTCAAGAATACTGGTGTAAGAATGCAACTGATGAAGGAATATTAGCTTATAAGCTTATGGTACAGACGGGTCATGTAGACAATCCGATTGACAAATCACTCATTAATTCTGGTCACAGATTAGTCGATAAAGATGGGATTATAAATCCTAAAGCGTTTTACAACTATCTATCAGCATGGGCAACGAATGACGCATTAGCGTATGGAGCATCTCAAGGAAATCTGAAACCACAACCCCAAAGATGGATTCATTCTCCGGAAGACGTCCATTTAGAAATAAAGAAGTCATCACCGTTAATCTACACTCAATTACCGTTTTATTTATCGGGGCTGAGTGATACTGACAGTATAAAGACATTGATAAGGTCAGTTCGAGAGCTTTGTTTGAAGTACGAGGCGAAAGGATTACCGAACTTTCCTTCTGGAATACCGTTCTTGTTTTGGGAACAGTATCTGTATTTGAGGACATCACTGTTACTGGCTCTCGCGTGTGCTTTGGGTGCTGTTTTTATTGCGGTGATGGTGCTGCTCTTGAATGCATGGGCGGCGGTGTTAGTGACTCTATCATTAGCCACTTTGGTGCTGCAACTGTTAGGAGTAATGGCTATTCTGGGCGTCAAGCTCTCAGCAATGCCTGCGGTGTTACTGGTACTGGCTGTTGGACGAGGCGTACATTTTACTGTCCATTTATGTTTGGGTTTCGTTACTTCAATAGGTTGCAAGCGTCGCCGAGCGTCGCTAGCTCTAGAGGCAGTGCTCGCGCCGGTGGTGCACGGAGCGCTGGCCGCTGCGCTGGCAGCCTCCATGCTAGCTGCCAGCGAGTTCGGCTTCGTGGCACGACTGTTCCTCAGACTGCTACTGGCACTCGTGGTGCTTGGATTAGTTGACGGATTGCTGTTCTTCCCTATCATACTGTCGATATTGGGACCGGCTGCTGAGGTGCAACCTTTGGAGCATCCCGAACGATTGTCAACACCTTCACCGAAAAGTTCGCCCGTCCATCCTCGGAAATCAAGTTCTGGTTCAAACGGAGGCGACAAATCCAGTCGAACCAGCAAAACGGCTCCACGACCTTCCGGACCATCATTGACAACTATAACAGAAGAGCCATCCAGTTGGCACAGCTCCAACCACTCTGTTCAGTCGTCAATGCAGTCCATAGTGGTTCAACCGGAGGTGGTGGTAGAGACGACTACGTATAACGGAAGCGATTCTACTTCTGGCAGATCAACGCCGACTTCGAAATCGACGCACACTGGAGCTGTCACTACAACTAAGGTAACCGCAACAGCGAATATAAAAGTTGAGGTAGTAACGCCAAGCGACAGAAAATCACGACGCTCATATCATTACTACGACCGTCGAAGGGATCGAGATGATGACAGAGATAGAGAAAGAGATCGTGATCGGGATCGAGATAGAGACAGAGATCGTGAAAGAGATCGTGATAGAGACAGAGAGAGATCTAGAGAACGGGACCGAAGGGACAGGTATAGGGAAGAGAGAGATCATCGCGCTTCGCCGAGAGAAAACGGTCGTGATTCTGGGCATGAAAGTGATTCTTCCCGACACTGA

Ptc1_HCR_P1B4: CCTCAACCTACCTCCAACAATCTAATACTTCGAGCGGGTTCAAAT

Ptc1_HCR_P2B4: AACTGGAACTTTAGCTTCTTCACTTATTCTCACCATATTCgCTTC

Ptc2_HCR_P1B4: CCTCAACCTACCTCCAACAACGCCGTGTCTGGACAATGAGGGTCG

Ptc2_HCR_P2B4: TGGAATATGTCCTGATTTTTTGTTCATTCTCACCATATTCgCTTC

Ptc3_HCR_P1B4: CCTCAACCTACCTCCAACAATGATGTGGAGTTCCTCGTCGCTCCG

Ptc3_HCR_P2B4: CTGCAGGGCGCGCGCACTCCTCAGAATTCTCACCATATTCgCTTC

Ptc4_HCR_P1B4: CCTCAACCTACCTCCAACAAAGTACGGCGGCGGCTTTTTCTTGAT

Ptc4_HCR_P2B4: GCTGCAAACTTTCTCTGCCAAGCATATTCTCACCATATTCgCTTC

Ptc5_HCR_P1B4: CCTCAACCTACCTCCAACAAGAGACTTCCGAGAATTTTCCGAGTA

Ptc5_HCR_P2B4: ATGTATCCCAAAATAATGTTCTTTAATTCTCACCATATTCgCTTC

Ptc6_HCR_P1B4: CCTCAACCTACCTCCAACAACGGCTACTGTGATCGACAGAAGCAA

Ptc6_HCR_P2B4: CTAATAATGCACAAAAGCCTAAGCCATTCTCACCATATTCgCTTC

Ptc7_HCR_P1B4: CCTCAACCTACCTCCAACAATCTCCCGCTTGTTCAACGTATGTGT

Ptc7_HCR_P2B4: AGTCCGGTTCTCTCTTCTCTCGGCAATTCTCACCATATTCgCTTC

***>Mothers_against_decapentaplegic_homolog_6 (Mad6)***

lcl|XM_024089045.2_cds_XP_023944813.2_1 [gene=LOC112050707] [db_xref=GeneID:112050707] [protein=mothers against decapentaplegic homolog 6] [protein_id=XP_023944813.2] [location=83..1012] [gbkey=CDS]

ATTTCAGAACACGGCGAAATTACACAAGACGCGAGGAGGACGAATCATGAGCGGCTTGCTACAGGCTCCCTCGCTACTGATGGCGAAGAGCGGCAGAGCTGGGAGACCGAGTGGTGCAGGCTGGCGTACTGGGAGCTGACGCAGCGTGTGCGAAAAACTCGAGCGAAGATTGGCCTAGGTGTCACACTGTCCTTAGAATCTGATGGCGTCTGGCTCTACAATAGAAGCCAAGAACCCGTGTTCGTCAGCTCCCCCGCGTTAGACGCTGCTGCTGCGAAAGCTCTTCTTGTATGGAGGGTTGCACCAGGACACTGTCTCTGCATCTTCGACCCCTCGTCGCCCCCGCCCGCTGTGTCGCTACCCCACGTGGGGCCAGTTGACCCCAGATCTGTGAGGATATCGTTCGCGAAAGGCTGGGGCCCCAAATACTCGAGGCGTGACGTCACCGCCTGCCCCTGTTGGCTCGAAGTCCTGCTGGCGCCTCCGAGCTGA

Mad61_P1_B1	gAggAgggCAgCAAACggAACGTCTTGTGTAATTTCGCCGTGTTC

Mad61_P2_B1	CAAGCCGCTCATGATTCGTCCTCCTTAgAAgAgTCTTCCTTTACg

Mad62_P1_B1	gAggAgggCAgCAAACggAACCCAGCTCTGCCGCTCTTCGCCATC

Mad62_P2_B1	AGTACGCCAGCCTGCACCACTCGGTTAgAAgAgTCTTCCTTTACg

Mad63_P1_B1	gAggAgggCAgCAAACggAAGCCAATCTTCGCTCGAGTTTTTCGC

Mad63_P2_B1	AGATTCTAAGGACAGTGTGACACCTTAgAAgAgTCTTCCTTTACg

Mad64_P1_B1	gAggAgggCAgCAAACggAACAGAGACAGTGTCCTGGTGCAACCC

Mad64_P2_B1	GGCGGGGGCGACGAGGGGTCGAAGATAgAAgAgTCTTCCTTTACg

Mad65_P1_B1	gAggAgggCAgCAAACggAATCACAGATCTGGGGTCAACTGGCCC

Mad65_P2_B1	GGCCCCAGCCTTTCGCGAACGATATTAgAAgAgTCTTCCTTTACg

***>engrailed (en)***

lcl|MT648682.1_cds_QZA76182.1_1 [protein=engrailed] [protein_id=QZA76182.1] [location=1..1059] [gbkey=CDS]

TTGAAGACCGTTGCAGTCCGAACCAGGCCAACAGCCCCGGTCCGGTCACCGGCAGAGTCCCTGCGCCTCACTCCGAAGTAAGAAACGNGTACCAAAGTCAATACACTTGCACGACTATCGATCAAAGGTTTGACAGAACGATGACAGTGGTGAAAGTGCAGCCGAATTCACCACCGATGAGTCCACTGACGTGAAGCCCATAATCCCTGAGTTTGAAGACAAGAGAAACCGACAACCACCACCAACCATACCCTTCTCTATCAGCAACATATTACACCCAGAATTCGGTTTGACAGCGATTCGAAAAACGAACAAAATCGAAGGACCAAAACACGTCGGCCCCAACCACAGCATTTTGTACAAACCTTATTTGTCGAACGAGTTATCGAGTTCGAAATTCAATTTCGATTATTTAAAATCTAAGGATGATTTCGGTGCATTACCTCCACTTGGCGGTTTGAGGCAGACCGTGTCGAATATTGGAGAACAGAAGGAGGCACCAAAGATTATAGAGCAGCAGAAGAGGCCAGATTCAGCCAGCTCTATTGTCTCTTCCACATCTAGCGGGGCTTTATCGACGTGTGGCAGCACTGACGCCAACAGCAGTCAAAGCGGGAACAGCAATCTA

en1_HCR_P1B1: gAggAgggCAgCAAACggAATGGCCTGGTTCGGACTGCAACGGTC

en1_HCR_P2B1: CTCTGCCGGTGACCGGACCGGGGCTTAgAAgAgTCTTCCTTTACg

en2_HCR_P1B1: gAggAgggCAgCAAACggAATTTGGTACNCGTTTCTTACTTCGGA

en2_HCR_P2B1: GATCGATAGTCGTGCAAGTGTATTGTAgAAgAgTCTTCCTTTACg

en3_HCR_P1B1: gAggAgggCAgCAAACggAAGCTGCACTTTCACCACTGTCATCGT

en3_HCR_P2B1: TCAGTGGACTCATCGGTGGTGAATTTAgAAgAgTCTTCCTTTACg

en4_HCR_P1B1: gAggAgggCAgCAAACggAAGTTTCTCTTGTCTTCAAACTCAGGG

en4_HCR_P2B1: GAAGGGTATGGTTGGTGGTGGTTGTTAgAAgAgTCTTCCTTTACg

en5_HCR_P1B1: gAggAgggCAgCAAACggAACTGTCAAACCGAATTCTGGGTGTAA

en5_HCR_P2B1: CTTCGATTTTGTTCGTTTTTCGAATTAgAAgAgTCTTCCTTTACg

en6_HCR_P1B1: gAggAgggCAgCAAACggAATGTACAAAATGCTGTGGTTGGGGCC

en6_HCR_P2B1: TCGATAACTCGTTCGACAAATAAGGTAgAAgAgTCTTCCTTTACg

en7_HCR_P1B1: gAggAgggCAgCAAACggAAATCATCCTTAGATTTTAAATAATCG

en7_HCR_P2B1: ACCGCCAAGTGGAGGTAATGCACCGTAgAAgAgTCTTCCTTTACg

en8_HCR_P1B1: gAggAgggCAgCAAACggAA CTCCTTCTGTTCTCCAATATTCGAC

en8_HCR_P2B1: CTTCTGCTGCTCTATAATCTTTGGTTAgAAgAgTCTTCCTTTACg

en9_HCR_P1B1: gAggAgggCAgCAAACggAATAGATGTGGAAGAGACAATAGAGCT

en9_HCR_P2B1: TGCTGCCACACGTCGATAAAGCCCCTAgAAgAgTCTTCCTTTACg

***>enhancer of split (ensl)***

lcl|XM_024090728.2_cds_XP_023946496.2_1 [gene=LOC112051893] [db_xref=GeneID:112051893] [protein=enhancer of split mbeta protein-like] [protein_id=XP_023946496.2] [location=42..791] [gbkey=CDS]

ATGATCCTCCCCGCTCGCACCATGTCCCCTCCGCACGGGGCCTTCCACCCGCCGCAGCACCCTGACGAAG

AACCCGTCTCGCGCACCTACCAGTACCGCAAAGTGATGAAACCCATGCTCGAGAGGAAGCGACGAGCACG

CATCAACCGATGCCTCGACGAACTCAAAGACCTCATGGTCACAGCCCTACAAGCGGAAGGCGAAAATGTT

TCCAAACTGGAAAAAGCAGATATCCTCGAGCTGACCGTCCGCCACCTCCACAGTTTAAAACGAAGAGGAC

AGTTAGTGTTAAACCCCGAAATGTCTTACGCGGAACGTTTTCGAGCAGGCTTTGCTCAGTGTGCTACTGA

AGTGTCACAGTTCATCACCAACGCAACGGTAGCGGCTAACGCTATGCACAGACAAGGGCCAGTGGATCCT

CAGGCCGGTGCTCGGTTGTTGCAGCACTTGAGCAACTGTATACGAAGGTTGGAAGCTCCGCAAGTGGTTC

AGCCTCAACCTATAACACCAGTCGCGAGGCCTACGCCAGTGCAAGCGGTAGCTCAACCTCAAGCGCTTCA

AGCGCCTCAACCTCCAGTACCTCAAGCGCCGCAGCCGGTTCCAAGGACGTCCTTGGAGAGGAAGAGACCA

GCTGAGGAGTTGATCGACGTGGAAACGATCGCTTCGAAGAGACTGTACGCTCCACCCTCGCCGCCTCACA

GCCCCGCCTCTGAGACTGACTCCGCCCAGTCGATGTGGCGACCGTGGTGA

B1_P1 (ensl)_1	GAGGAGGGCAGCAAACGGAAAGGGGACATGGTGCGAGCGGGGAGG

B1_P1 (ensl)_2	GAGGAGGGCAGCAAACGGAATGCGGTACTGGTAGGTGCGCGAGAC

B1_P1 (ensl)_3	GAGGAGGGCAGCAAACGGAATCTTTGAGTTCGTCGAGGCATCGGT

B1_P1 (ensl)_4	GAGGAGGGCAGCAAACGGAACTCGAGGATATCTGCTTTTTCCAGT

B1_P1 (ensl)_5	GAGGAGGGCAGCAAACGGAACGTAAGACATTTCGGGGTTTAACAC

B1_P1 (ensl)_6	GAGGAGGGCAGCAAACGGAAACCGTTGCGTTGGTGATGAACTGTG

B1_P2 (ensl)_1	CTGCGGCGGGTGGAAGGCCCCGTGCTAGAAGAGTCTTCCTTTACG

B1_P2 (ensl)_2	TCCTCTCGAGCATGGGTTTCATCACTAGAAGAGTCTTCCTTTACG

B1_P2 (ensl)_3	TCCGCTTGTAGGGCTGTGACCATGATAGAAGAGTCTTCCTTTACG

B1_P2 (ensl)_4	TAAACTGTGGAGGTGGCGGACGGTCTAGAAGAGTCTTCCTTTACG

B1_P2 (ensl)_5	GAGCAAAGCCTGCTCGAAAACGTTCTAGAAGAGTCTTCCTTTACG

B1_P2 (ensl)_6	CCTTGTCTGTGCATAGCGTTAGCCGTAGAAGAGTCTTCCTTTACG

***>homothorax (hth)***

lcl|XM_024094433.2_cds_XP_023950201.1_1 [gene=LOC112054591] [db_xref=GeneID:112054591] [protein=homeobox protein homothorax isoform X3] [protein_id=XP_023950201.1] [location=301..1596] [gbkey=CDS]

ATGGCTCAGCCTAGGTACGACGAGAGCCTCCACGGCGGGGGCTACATGGAGGGCGGCGCCATGTACCACGAGCACCGGCTCACGCACCCGCACATCCCGCCGGTGCACTACCCGCCGCCCGCCGCGCCGGCGCACGCGTTGCCCGGCGAGCCGCTAGTGCACAAGCGCGACAAGGACGCCATATACGGGCATCCCCTGTTTCCCCTGCTGGCGCTGATCTTCGAGAAGTGCGAGCTGGCAACGTGTACCCCCCGCGACCCCGGCGTAGCCGGCGGTGACGTCTGTTCCTCAGAGTCCTTTAACGAGGACATCGCGGTGTTCAGTAAACAGATACGTCAAGAAAAACCTTATTACATAGCGGACCCCGAGGTAGACTCATTAATGGTGCAAGCAATACAAGTCCTACGGTTTCACCTATTAGAATTAGAAAAAGTGCACGAGCTGTGCGACAACTTCTGCCACCGCTACATCAGCTGCCTGAAGGGCAAGATGCCCATCGACCTGGTGATCGACGAGCGGGAGTCAGCCCGGCCGCCGACACCAACGGGGAGCCGCGGTCGGCGCCTGACAGCAACCACGACGGCGCATCGACCCCCGACGTCAGGCCGCCATCGTCGTCGCTATCATACGGCGGTGCGGTGAACGATGACGTCCGCTCACCGGGCTCCGGTGGCACCCCCGGTCCCCTCAGCCAGCCCCCGCCGCAGACCCTCGACGCGACAGATCCAGATGCCATGGGCAAATGGTGCGGGTCGCGGCGGGAATGGTCATCCCCTCCCGACGTGGCGCGGCGGGTCTACTCCTCAGTGTTCCTGGGCAGTCCCGGGGAATACCCAGGGGATGCCAGTAACGCGAGTATCGGCTCCGGCGAGGGTACGGGGGAGGAAGACGACGACACGAACGGAAAGAAGAACCAAAAGAAACGGGGAATCTTTCCGAAGGTCGCCACCAACATCCTTAGAGCGTGGCTCTTTCAGCACTTAACGCATCCCTACCCTTCGGAAGACCAGAAGAAACAGTTGGCACAAGACACAGGGTTAACGATACTACAAGTAAATAATTGGTTCATCAACGCGAGACGTAGGATAGTACAGCCAATGATAGACCAGTCGAATAGAGCAGTGTTCTACCCCGCAGTGTTCCCGCACGCGGGCCCCAGCGGCGCCTACAGCCCGGAGGCCACCATGGGCTACATGATGGACGGCCAGCAGATGATGCACAGGCCGCCGGCCGACCCCGCCTTCCACCAGGGCTACGCGCACTACCCCGCCGAGTACTACGGACACCATCTTTAA

Htha_HCR_P1B1: gAggAgggCAgCAAACggAAGCGACCCGCACCATTTGCCCATGGC

Htha_HCR_P2B1: CGTCGGGAGGGGATGACCATTCCCGTAgAAgAgTCTTCCTTTACg

Hthb_HCR_P1B1: gAggAgggCAgCAAACggAATTCCTCCCCCGTACCCTCGCCGGAG

Hthb_HCR_P2B1: GTTCTTCTTTCCGTTCGTGTCGTCGTAgAAgAgTCTTCCTTTACg

Hthc_HCR_P1B1: gAggAgggCAgCAAACggAACTGTTTCTTCTGGTCTTCCGAAGGG

Hthc_HCR_P2B1: TATCGTTAACCCTGTGTCTTGTGCCTAgAAgAgTCTTCCTTTACg

hth3_HCR_P1B1: gAggAgggCAgCAAACggAAACTGCGGGGTAGAACACTGCACGGT

hth3_HCR_P2B1: GCGCCGCTGGGGCCCGCGTGCGGGATAgAAgAgTCTTCCTTTACg

hth4_HCR_P1B1: gAggAgggCAgCAAACggAAGTCCATCATGTAGCCCATGGTGGCC

hth4_HCR_P2B1: CGGCGGCCTGTGCATCATCTGCTGGTAgAAgAgTCTTCCTTTACg

hth5_HCR_P1B1: gAggAgggCAgCAAACggAATAGTGCGCGTAGCCCTGGTGGAAGG

hth5_HCR_P2B1: AGATGGTGTCCGTAGTACTCGGCGGTAgAAgAgTCTTCCTTTACg

***>hedgehog (hh)***

lcl|XM_024078060.2_cds_XP_023933828.2_1 [gene=LOC112042874] [db_xref=GeneID:112042874] [protein=tiggy-winkle hedgehog protein] [protein_id=XP_023933828.2] [location=228..1376] [gbkey=CDS]

ATGTCGATAAAGCGGTGCAAAGAGAAGTTGAACACGCTCGCCATCAGTGTGATGAACCAGTGGCCGGGGGTTCGACTCCGGGTCATCGAGGGCTGGGACGAGGAGAACTCGGCTCACCTGGAAAACTCACTGCACTACGAGGGCCGGGCAGTGGACATCACCACCAGCGACCGGGATCGCAGCAAGTACGGCATGCTGGCACGCCTTGCTGTGGAAGCCGACTTCGACTGGGTGTTCTATGAGAGCCGGTCCTACATACATTGTTCTGTCAAGACAGAATCATCAGTGGGCACTGGAGCTGGTTGTTTTCCTTCTGGGTCTGTTGTGCATACGGAAGAAGGTCCTCGGGACATCGCTACTCTCAAGAAAGGCGATCGAGTTCTGGCTGCAGATGACGATGGCAAGATGGTCTATTCAGAGGTTTTAACATTTATTGATCGAGAACCAAACGCGACGCGACAGTTCGTCGAAGTGACAGCAGAGAACGGCGTGAGTATAACAACCACGCCGTCACATTTGCTGCTGCTAGCTGCCGCAGACGGATGGCGGGAGTCCTTCGCCACCAACATAGAAGCCGGTGACGTCCTCCTGACGAGAGGGCAGGGCAGCGTCATGCGACCGTCGAGAGTTGTCAAAACTCGACTGGTATCGAAGAGAGGGGTTTATGCACCCCTCACAAGGACAGGGACTATCATTGTAGACGATGCATTGGCGTCTTGCTACGCTCTCGTGCGAAGCCATGCTCTAGCGCACGCTGCAATGGCTCCACTGCGCTGGATGGCCGGCTGGACTTCCAACGAAGTCTCCCGCGGAGTCCATTGGTACGCTAATGCCCTCTACAACGTCGGCGACTATGTCTTACCCGCCTCGTACAAGTATCGCTGA

Hh1_HCR_P1B4	CCTCAACCTACCTCCAACAAGTCCCAGCCCTCGATGACCCGGAGT

Hh1_HCR_P2B4	GTTTTCCAGGTGAGCCGAGTTCTCCATTCTCACCATATTCgCTTC

Hh2_HCR_P1B4	CCTCAACCTACCTCCAACAACAGTCGAAGTCGGCTTCCACAGCAA

Hh2_HCR_P2B4	ATGTAGGACCGGCTCTCATAGAACAATTCTCACCATATTCgCTTC

Hh3_HCR_P1B4	CCTCAACCTACCTCCAACAAAGAGTAGCGATGTCCCGAGGACCTT

Hh3_HCR_P2B4	GCAGCCAGAACTCGATCGCCTTTCTATTCTCACCATATTCgCTTC

Hh4_HCR_P1B4	CCTCAACCTACCTCCAACAAACTCACGCCGTTCTCTGCTGTCACT

Hh4_HCR_P2B4	CAGCAAATGTGACGGCGTGGTTGTTATTCTCACCATATTCgCTTC

Hh5_HCR_P1B4	CCTCAACCTACCTCCAACAATCTCGACGGTCGCATGACGCTGCCC

Hh5_HCR_P2B4	CTTCGATACCAGTCGAGTTTTGACAATTCTCACCATATTCgCTTC

Hh6_HCR_P1B4	CCTCAACCTACCTCCAACAATTGCAGCGTGCGCTAGAGCATGGCT

Hh6_HCR_P2B4	AGCCGGCCATCCAGCGCAGTGGAGCATTCTCACCATATTCgCTTC

***>eyegone (eyg)***

lcl|XM_052882095.1_cds_XP_052738055.1_1 [gene=LOC112055116] [db_xref=GeneID:112055116] [protein=paired box protein Pax-6-like isoform X2] [protein_id=XP_052738055.1] [location=312..1763] [gbkey=CDS]

ATGAACCTCCTCAAGCGCGGTGGCGGCTCCCCCTCGCACCGCTTGCCCCACTCCCCCTCCCGCTCCCGTTCGCGTTCTTTGTCCCCCGCGCGCGTCCCCTACCATACCCCCCAGATGGGCGGCGAGAATAGTGCGTTCAAGGCTCTGGCGCACCAGGACCCCAACGCGCTGAAGGCTCTCTCGCAGCAGGCTCAGTTCGACAGTACGACGCTGAAAGCGCTGTCGCAGCAGCAGTTCGACTCGTTCCAGCCGCACCCAGCGCTGGAGAGCAGCGCTTTCAAAGCGCTCGTCCCCAACTCGGCCGCCGCCGCGCTTCTGGCAGCACAATCGATACAATTAGCCCGCGGATACGAATCTCATTCAGATTCCGACGAGGAAATAAACGTCCACGACTCGAGTGACGACGAAGCCGAGAAACAGATTAACAAGACGAGATCTAGATCTAGATCTCCGAGCCCCGAGCGGCGCAGAATAGCGGCCAACGACTTGCCACTACAATTGACTAAACATGACCGTTGA

B2_P1 (eyg)_1	CCTCGTAAATCCTCATCAAAGGAGCCGCCACCGCGCTTGAGGAGG

B2_P1 (eyg)_2	CCTCGTAAATCCTCATCAAACCGCCCATCTGGGGGGTATGGTAGG

B2_P1 (eyg)_3	CCTCGTAAATCCTCATCAAAAGCCTGCTGCGAGAGAGCCTTCAGC

B2_P1 (eyg)_4	CCTCGTAAATCCTCATCAAAGCGCTGGGTGCGGCTGGAACGAGTC

B2_P1 (eyg)_5	CCTCGTAAATCCTCATCAAAATCGATTGTGCTGCCAGAAGCGCGG

B2_P1 (eyg)_6	CCTCGTAAATCCTCATCAAAGTCACTCGAGTCGTGGACGTTTATT

B2_P2 (eyg)_1	GGGGGAGTGGGGCAAGCGGTGCGAGAAATCATCCAGTAAACCGCC

B2_P2 (eyg)_2	GCCAGAGCCTTGAACGCACTATTCTAAATCATCCAGTAAACCGCC

B2_P2 (eyg)_3	CGCTTTCAGCGTCGTACTGTCGAACAAATCATCCAGTAAACCGCC

B2_P2 (eyg)_4	CGAGCGCTTTGAAAGCGCTGCTCTCAAATCATCCAGTAAACCGCC

B2_P2 (eyg)_5	TGAGATTCGTATCCGCGGGCTAATTAAATCATCCAGTAAACCGCC

B2_P2 (eyg)_6	CTTGTTAATCTGTTTCTCGGCTTCGAAATCATCCAGTAAACCGCC

***>aristaless_like (al-like)***

(LOC112054675)

ATGGGGCTATCAGAACCACCGAAAGACGACTCTCCTCGAACTACTCCAGAACTCTCCCGCGCCGACCAGTCGCCTGCTTCGGAAAGACCACCTTTAGGCTCTGCTGACAGCGATGATGCAGATGACTTTGCCCCCAAGAGAAAACAAAGGCGGTACAGGACGACATTCACAAGTTTCCAGTTAGAAGAATTAGAAAAGGCGTTTTCAAGGACACATTACCCTGATGTTTTCACAAGAGAGGAGCTTGCAATGAAGATCGGCCTAACGGAGGCAAGAATACAGGTATGGTTCCAGAACCGACGCGCAAAATGGCGGAAGCAGGAGAAGGTGGGCCCGCAAGCTCACCCCTACAACCCCTACCTAGGGGGTGCGGCCCCCCCGCCCTCGGTCGCGTCCATGCCGAACCCCTTCACACAACTCGGCTTCGGATTCAGGAAGCCATTCGATACTAATGCTCTGGCTACGTTCAGGTACGGCAATTCTCCACTGCTGGGCACCCAATACCTCGGAGCTCCACTCTCCCGACCACCTCTATTTACAGCGCCAATATATTCAACGACACCACCCTTTCATTCGCTGCTAGCGGGCCTCGCAGCACCCCCCCGACAGTCTCCTGACCCCCCCTCTGTCTCCCCTCCAATATCCCCAGGCAGTGAGTCTCCCCCCAGTCAACCAGGGCCGGAGGTAGAACGACGGAGTTCGAGCATCGCAGCATTGAGACTTGCTGCAAGAGAACATGAAATACGATTGGAAATGTTGAGACAGCGACACCACACTGACCTGATTAGTTGA

B1_P1 (al_like)_1	GAGGAGGGCAGCAAACGGAAGTCGTCTTTCGGTGGTTCTGATAGC

B1_P1 (al_like)_2	GAGGAGGGCAGCAAACGGAAAGCCTAAAGGTGGTCTTTCCGAAGC

B1_P1 (al_like)_3	GAGGAGGGCAGCAAACGGAAGGGGGGGCCGCACCCCCTAGGTAGG

B1_P1 (al_like)_4	GAGGAGGGCAGCAAACGGAAAGGCCCGCTAGCAGCGAATGAAAGG

B1_P2 (al_like)_1	GGAGAGTTCTGGAGTAGTTCGAGGATAGAAGAGTCTTCCTTTACG

B1_P2 (al_like)_2	AGTCATCTGCATCATCGCTGTCAGCTAGAAGAGTCTTCCTTTACG

B1_P2 (al_like)_3	GGGTTCGGCATGGACGCGACCGAGGTAGAAGAGTCTTCCTTTACG

B1_P2 (al_like)_4	TCAGGAGACTGTCGGGGGGGTGCTGTAGAAGAGTCTTCCTTTACG

***>dachshund (dac)***

lcl|XM_052883449.1_cds_XP_052739409.1_1 [gene=LOC112044740] [db_xref=GeneID:112044740] [protein=dachshund homolog 1 isoform X2] [protein_id=XP_052739409.1] [location=119..1627] [gbkey=CDS]

ATGGAGTCCGCCGTGGACTCAGCGTCGACCGCCAGCGAGGTGAGCGGCTCGTCCGGGGGCTCGCCGCGCGTCAAGGCGGCGTCGCCGGCGCGCGGCCTCAGCCCGCCGCAGCTGCTGGCGCCGCGCCTGCCGCTGCCGCCGCCGGGCCTGGGCCTGCTGGGCTCGCTGCAGATGATGCACCACTCGCCGCTCGAGCTCATGGCGGCGGCGCACCACCACCCGCACCGCTACGGGAGCCCGCCGCCCATCTCCACGTCCGACCCCTCGGCGAACGAGTGCAAGCTGGTGGACTATCGCGGGCAGAAGGTGGCCGCGTTCATCATCCAGGGCGACACGATGCTGTGCCTGCCGCAGGCCTTCGAGCTGTTCCTGAAGCACCTGGTGGGCGGGCTGCACACCGTGTACACCAAGCTGAAGCGGCTGGACATCGTGCCGCTGGTGTGCAACGTGGAGCAGGTCCGCATCCTGCGCGGGCTGGGCGCCATCCAGCCGGGCGTCAACCGCTGCAAGCTGCTCTCGTGCAAGGACTTCGACGTGCTGTACCGCGACTGCACCACGGCAAGGTGCCTGTCAATGAAAGCGCCAGACAGCTCCAGACCGGGTCGACCTCCGAAGCGTGCTTCTGGAGTTGGTCTTTCGCTTGCCGCCACGCAGTTCCCGGGGCACCCCTTCAAGAAGCACCGCCTAGAGAATGGGGACTACTCGCCGTATGAAAATGGACATATGAGCGAGATGGCCCGCATGGATAAGTCCCCGCTCCTCGCGAACGGGTACAACGCACCCCCCACCCACCTGGGGCCCATGGGCTTCATGCACCAGCACGCCCTCATGTCTCCGGGTATGCCACACCCTGGCGTGCCCAGGCCTGACGGGTCCATCATTAAGGGGCAGCCTATGCATAACATGGAAGCACTAGCAAGATCTGGTATTTGGGAGAATTGTAGAGCAGCATACGAGGACATCGTAAAACATCTCGAAAGGTTGCGTGATGAACGGGGTGATATTGAACGTGTTATAGCAATGGACAAGGCGCGCGAGGGTTCACATAATGGTTCGTCTCCAGGCCACAGTCCTGTGCTGAACCTTTCGAAGTCAGGCTCCGGTGAGCGCGAGCGGTCCGAGCGGGACCGCGAGCGCGCGGAGCGGGGCGAAGGCTCGGCGAGCGGGCGCAGCTCGGCCGCCTCGCGCCGCACGCCGCAGCCGCCGCGCATCCCCTCCACCGCCGCGCCGGTCTCCCCGCGCTCCCACTCCGACGAGAGCGACGCTGCTCTGTCTGATCAAGACGACCATAACGTCAAAGACGAGGATGACGATCTCAGCGATGGTGAACGAGACCTCGCAACGAATTCCTCGCCAGCGCCCGTCAGCTACCCCCCACAAGGCTCGCCGTCGAACGTTCCCGTGGACCCCACCGCCGACACCCTGGTCTCCTCCACCGAGACCCTCCTCAGGAACATCCAGGGCCTCCTCAAAGTAGCGGCGGACAACGCACGCCAACAAGAACGGCAAATCAGTTACGAAAAAGCGGAACTAAAAATGGACGTGTTAAGAGAAAGAGAAGTAAAAGACAACCTGGAAAGACAGTTACTGGACGAACAAAAAATGAGAGTAATGTATCAGAAGAGATTAAAGAAAGAAAGAAAGCAGAGGCAACAAATTCAAGACCAATTAGAATTGGAGTTAAAAAGACGACAGAAGATAGAAGAAGCACTAAAGCAGTCAGGAGCGCCCGGTGAAATTCTGAGAATAGTAACTGAGAATTTAACACCACCGTCCCAAGAAAATCGCGAACGTGAGAACGGTACAGAGAGCAAACCGCCAAGTACAGAGCCCCCCACCTCGTCGCCGCCGTTCCAGAGGGACCCCCCGCGCACGCCCGACAAGCCCCAGTGGAACTACCCTCCGCCGCCTGTCGACATCATGAGCGGAGGAGCTGCTTTCTGGCAGAACTACTCTGAATCCTTGGCGCAAGAGTTGGAGATGGAGCGCAAGTCCCGCCAGCAGGCCATGGAGAGAGACGTGAAGAGCCCTCTGTCGGACCGCGCCAGTTACTACAAGAACTCGGTGCTGTTCAGCTCGGCCACTTAG

Dac1_HCR_P1B3	gTCCCTgCCTCTATATCTTTCCTCGCTGGCGGTCGACGCTGAGTC

Dac1_HCR_P2B3	GCGGCGAGCCCCCGGACGAGCCGCTTTCCACTCAACTTTAACCCg

Dac2_HCR_P1B3	gTCCCTgCCTCTATATCTTTGCAGGCCCAGGCCCGGCGGCGGCAG

Dac2_HCR_P2B3	AGTGGTGCATCATCTGCAGCGAGCCTTCCACTCAACTTTAACCCg

Dac3_HCR_P1B3	gTCCCTgCCTCTATATCTTTTGCGGGTGGTGGTGCGCCGCCGCCA

Dac3_HCR_P2B3	GAGATGGGCGGCGGGCTCCCGTAGCTTCCACTCAACTTTAACCCg

Dac4_HCR_P1B3	gTCCCTgCCTCTATATCTTTGGCCTGCGGCAGGCACAGCATCGTG

Dac4_HCR_P2B3	CACCAGGTGCTTCAGGAACAGCTCGTTCCACTCAACTTTAACCCg

Dac5_HCR_P1B3	gTCCCTgCCTCTATATCTTTCTGGATGGCGCCCAGCCCGCGCAGG

Dac5_HCR_P2B3	GAGCAGCTTGCAGCGGTTGACGCCCTTCCACTCAACTTTAACCCg

Dac6_HCR_P1B3	gTCCCTgCCTCTATATCTTTAGCACGCTTCGGAGGTCGACCCGGT

Dac6_HCR_P2B3	GGCGGCAAGCGAAAGACCAACTCCATTCCACTCAACTTTAACCCg

>***vestigial (vg)***

lcl|XM_024079072.2_cds_XP_023934840.1_1 [gene=LOC112043588] [db_xref=GeneID:112043588] [protein=protein vestigial] [protein_id=XP_023934840.1] [location=405..1265] [gbkey=CDS]

ATGGCGGTGAGCTGCCCCGAGGTGATGTACGGCGCCTACTACCCCTACCTGTACGGCCGAGGCGCCGCGCGCTCCTTCCACCACGCGCCGCACTTCCAGTACGACCGGTTGAACGTGCAAAGCTCGGGCAATGCCAGCGCAGAGTACGCGGCCGCCGTCGGCTCGGGCGTGTCCGGCGCGTCGGGCTCGGCGTCGTCGGGCGCGTCGGGCGCGCAGTCGCCGGGCTCGCCGCCGCACGCGCACCACCTGGCGCACGCGCACCTGGCGCACCACGCGTCGCCGCGCCTGCGCGCCAAGGAGGAGGACCTGTCGGCGCACGGCCGCAACAGCGCCGACGCGTCCTCCGAGTCGGAGGGCGAGTGCGGCGGCGCGGCGCGCGCCCGCGCGCAGTACGTCAGCGCCAACTGCGTGGTGTTCACGCACTACTCCGGCGACGTGGCCGCCGTCGTGGACGAGCACTTCGCGCGCGCGCTGGCGCTCGACAAGCCCAAGGAGGGCGTCCCGCTGTCGGCGCGCAACCTGCCCGCGTCGTTCTTCAACGCGGCGGCGGGCGGCGGCGGCGCGTGCGCGCTGGAACTGTACGAGTACGACCCCTGGCACCAGCACTACGGGTACAACGCGCACCGGCACGCGGCGGAGTACCACGCGCACCACGGCATGGCGGCGGGCTACGGCGGCCTGCTGCTGGGCCGCGGCTCGCTGCACGCGCAGTACAAGCCGGTGGAGTGGCCGCACCACGCGCACCACGCGCACGCCGCGCCGCACCTCGACCCCGCCGCCTGCTCGCCCTACTCCTACCCGGCGCCGCCAGGTCTGGAGGCGCCGGTGCAGGACACGTCGAAGGACCTGTACTGGTTCTGA

vg1_HCR_P1B4: CCTCAACCTACCTCCAACAACGACGGCGGCCGCGTACTCTGCGCT

vg1_HCR_P2B4: AGCCCGACGCGCCGGACACGCCCGAATTCTCACCATATTCgCTTC

vg2_HCR_P1B4: CCTCAACCTACCTCCAACAACCCGGCGACTGCGCGCCCGACGCGC

vg2_HCR_P2B4: GCCAGGTGGTGCGCGTGCGGCGGCGATTCTCACCATATTCgCTTC

vg3_HCR_P1B4: CCTCAACCTACCTCCAACAAGCAGGCGCGGCGACGCGTGGTGCGC

vg3_HCR_P2B4: GCGCCGACAGGTCCTCCTCCTTGGCATTCTCACCATATTCgCTTC

vg4_HCR_P1B4: CCTCAACCTACCTCCAACAACTCCGACTCGGAGGACGCGTCGGCG

vg4_HCR_P2B4: GGCGCGCGCCGCGCCGCCGCACTCGATTCTCACCATATTCgCTTC

vg5_HCR_P1B4: CCTCAACCTACCTCCAACAAGTGAACACCACGCAGTTGGCGCTGA

vg5_HCR_P2B4: ACGGCGGCCACGTCGCCGGAGTAGTATTCTCACCATATTCgCTTC

vg6_HCR_P1B4: CCTCAACCTACCTCCAACAAAGCGCCAGCGCGCGCGCGAAGTGCT

vg6_HCR_P2B4: AGCGGGACGCCCTCCTTGGGCTTGTATTCTCACCATATTCgCTTC

***>mirror***

lcl|XM_052890551.1_cds_XP_052746511.1_1 [gene=LOC112045741] [db_xref=GeneID:112045741] [protein=homeobox protein caupolican isoform X2] [protein_id=XP_052746511.1] [location=409..1605] [gbkey=CDS]

ATGAGCAGCGCCGCGGCGGCTCGGCCGAGCTCGCCGGCGGCATCGACAGCGCGGTGTTGCGACACCGGCC

GCCCTATATTCCAGGACCCCATCACTGGGCAAACGGTCTGCTCGTGCCAGTACGAGTTCCTCAACTACCA

GCGGCTGGCGTCCGGGGTGCCGCTGTCGATGTACAGCGCGCCGTACCCGGACGCGGCCGCCGCGGCTGGC

ATGGCAGCCTACTTCCCCGCGCTCGCCGCCGACCAACCACCCTTCTACGCGAACACGGCTGCTGGTATCG

AGCTGAAGGAGAACCTGGCAGCCAGCGCGGCTAGTTGGCCTTACCCCACAGTCTACCACCCCTACGATGC

GGCTTTCGCTGGCTACCCCTTTAATGGATATGGAATGGATTTGAATGGTGCAAGAAGAAAAAATGCCACC

CGAGAAACGACGAGTACACTCAAAGCGTGGCTCAACGAGCACAAGAAGAACCCGTACCCAACCAAGGGGG

AAAAGATAATGTTGGCGATCATCACCAAGATGACGCTGACGCAGGTGTCCACGTGGTTTGCGAACGCACG

CCGGCGGCTCAAGAAGGAAAACAAGATGACCTGGGAGCCGAGAAACAGGGTCGATGATGACGACAATAAT

AATGATGATGATGACCACAAAAGCAATGACGGGAAAGATGCTTTGGATGGCAAGGACTCTGGGACAGGCT

CGAGTGAAGACGGTGAGCGACCACAACAAAGGTTAGATCTGCTGGGACAGCGAACAGAATCAGAATGGTC

GGAATCACGAGCGGACAGTGGACCCGAGTCACCCGAGCCGCCCTACGAACGGCCCATACATCCCGCGTAC

CAACACCTGCCTTCACGCGCACCGCCCGGTAGCACTCCTGCTTCAGCGAAACCCAGAATATGGTCATTAG

CTGACATGGCCAGTAAGGATGGGGAGGCTCCGGCTCCCCCAGCGGCTGCATCCGCTTTCTACCAATCAGC

TGCTGCGGCTGCGGCGGCTGCAAGGCTCGCCCATCCTTACGGCAGACCGGACTTGTACCGGGGCTTGTAC

CCCCCGACGCATCCAGCAGATGTCGCGCTCCTGGAGTACTCCCGCTCTCTAGCTCTCGCAGCCCCCGCGC

CCGCGCCGCCCGCCCTCTCCCCGTCCTCTTCGTCTACCTCGTCCCTTGCGGAGCCGCCGCCCCCTTCCCG

CGCCTGA

B3_P1(mirror)_1	GTCCCTGCCTCTATATCTTTGGCGGCGAGCGCGGGGAAGTAGGCT

B3_P1(mirror)_2	GTCCCTGCCTCTATATCTTTCCGCGCTGGCTGCCAGGTTCTCCTT

B3_P1(mirror)_3	GTCCCTGCCTCTATATCTTTTATCCATTAAAGGGGTAGCCAGCGA

B3_P1(mirror)_4	GTCCCTGCCTCTATATCTTTCCACGCTTTGAGTGTACTCGTCGTT

B3_P1(mirror)_5	GTCCCTGCCTCTATATCTTTTCTTGGTGATGATCGCCAACATTAT

B3_P1(mirror)_6	GTCCCTGCCTCTATATCTTTGTCATCTTGTTTTCCTTCTTGAGCC

B3_P2(mirror)_1	CGTGTTCGCGTAGAAGGGTGGTTGGTTCCACTCAACTTTAACCCG

B3_P2(mirror)_2	GGTAGACTGTGGGGTAAGGCCAACTTTCCACTCAACTTTAACCCG

B3_P2(mirror)_3	CTTCTTGCACCATTCAAATCCATTCTTCCACTCAACTTTAACCCG

B3_P2(mirror)_4	GTACGGGTTCTTCTTGTGCTCGTTGTTCCACTCAACTTTAACCCG

B3_P2(mirror)_5	ACCACGTGGACACCTGCGTCAGCGTTTCCACTCAACTTTAACCCG

B3_P2(mirror)_6	TCATCGACCCTGTTTCTCGGCTCCCTTCCACTCAACTTTAACCCG

***>sister of odd and bowel (sob)***

lcl|XM_024081021.2_cds_XP_023936789.1_1 [gene=LOC112044996] [db_xref=GeneID:112044996] [protein=protein sister of odd and bowel-like] [protein_id=XP_023936789.1] [location=108..1523] [gbkey=CDS]

ATGCAAGAACAAAGCACATCCCGCACCAACAAAACCACACTCAACTCACTGGAGAGCGCCATCATCAAGC

TGAAAAACAACCTGCAGATACAGGACACCAAACCCGCAGACATGACCGGTCTGTTGAGCATCGAGACACC

GGAGGACACTAAAAGCCCAAGTAAGAGCTCTAGTTCAACTACAACGAACTTAAACTTATCTATTGCATCC

GCTGCGTTGTTACAAGAGAACATGCTCCAGAGATCGTTGCAAGGCGTAGTGATATCCCTCCAAAATGCGA

TGATGAATTCTCTGCAACAGGCTGCTTTATTACCGTCGAATTCTGCGGCAGCCGCTGCCTTAAACTTGCA

AGCTTTAGAATCCTATTTAACTTTACAACGCCTCACATCTGTGTCATCTGTAAACAACACGCAATGTAGC

TCCAACGAAAGTATTACTTCAACTAAACAAGACATACTGAGGCTCGCGACGGCCAGCGCTAAGATTACTG

AAACGGACGTTTCTGAAAGTACATCTAGAGTGATATCTCCGAAAACGCCTGAAGAGTACCCCTTATCTGA

TGCTATAGCGGCTGAAGATGTAGATTTATCCGAAGAAGTGGAAGAAGACTTAGCTTTGTTAGACAACGAA

GACGAATTCACGTTGGAACAGCATCTAGAATCTACGTCAAAAGGATACGGGTACAGTGCATCACAGAGCT

TAGACAACGGTTACCCGAGCCTTCTAATGAATGCCATGTATCAGCAGCAAATGAATGGGAGCCCCTCTCA

GCCACCTCTGTCTCCTCAACCGTCCAATTCATCCAGCTCGAAGAGTACAAATAGTTCGACATCGTCGACG

GGAAGTTGTATATCTAATAAGCAGAAAACTACAGGCAGCGGACCGAGAACGAAAAAGCAGTTTATTTGCA

AGTTCTGTAATCGACAATTCACGAAATCCTACAATCTCCTGATACATGAAAGGACGCACACCGACGAGCG

GCCATACTCGTGTGACATCTGTGGAAAAGCGTTCCGACGGCAAGATCACCTCAGAGATCACAGATACATA

CACTCAAAAGAGAAACCATTCAAGTGTTTAGAATGTGGAAAAGGATTCTGCCAGTCCCGAACATTAGCGG

TACACAAAATACTACACATGGAAGAGTCTCCACACAAGTGCCCAGTCTGCAGCAGGAGTTTCAACCAACG

ATCGAATTTAAAAACCCATTTACTAACCCATACAGATATTAAGCCATACAACTGTACTTCATGCGGGAAG

GTATTCAGGCGTAACTGTGACTTGCGACGTCACAGTTTAACACACAACCTAGTTAGCATATCATCGGTGA

CATCACCCAACAGTGCTGGTAATAAAAGCCCCTTACTCAACCCTAACTTCCCTATTGACAATTTAGACAC

TGAAGACGAGAATTAA

B2_P1 (sob)_1	CCTCGTAAATCCTCATCAAAGTTGGTGCGGGATGTGCTTTGTTCT

B2_P1 (sob)_2	CCTCGTAAATCCTCATCAAATGGTGTCCTGTATCTGCAGGTTGTT

B2_P1 (sob)_3	CCTCGTAAATCCTCATCAAAGAGCTCTTACTTGGGCTTTTAGTGT

B2_P1 (sob)_4	CCTCGTAAATCCTCATCAAACTGGAGCATGTTCTCTTGTAACAAC

B2_P1 (sob)_5	CCTCGTAAATCCTCATCAAAATAAAGCAGCCTGTTGCAGAGAATT

B2_P1 (sob)_6	CCTCGTAAATCCTCATCAAACGTTGTAAAGTTAAATAGGATTCTA

B2_P2 (sob)_1	GCTCTCCAGTGAGTTGAGTGTGGTTAAATCATCCAGTAAACCGCC

B2_P2 (sob)_2	TCAACAGACCGGTCATGTCTGCGGGAAATCATCCAGTAAACCGCC

B2_P2 (sob)_3	AAGTTTAAGTTCGTTGTAGTTGAACAAATCATCCAGTAAACCGCC

B2_P2 (sob)_4	GGATATCACTACGCCTTGCAACGATAAATCATCCAGTAAACCGCC

B2_P2 (sob)_5	CAGCGGCTGCCGCAGAATTCGACGGAAATCATCCAGTAAACCGCC

B2_P2 (sob)_6	TTGTTTACAGATGACACAGATGTGAAAATCATCCAGTAAACCGCC

>***Distal-less (Dll)***

lcl|XM_052887172.1_cds_XP_052743132.1_1 [gene=LOC112055223] [db_xref=GeneID:112055223] [protein=homeotic protein distal-less isoform X1] [protein_id=XP_052743132.1] [location=898..2040] [gbkey=CDS]

ATGCATCGCGCCACGAGAGGCCGGGTAGCGTTCGGAGCAGTCCAGAAATCCCTCAAAATTACACGAATTCAGTCCCCAAACAGCAAACCCACCACGCTGAGTTTCTCAGATCCCTTTGGGCCTCCTCAGTCCACGGACGGGGGGGGCCCGTCGACCCCGCAGCCCGCCATGACCACCCAGGAGCTGGACCACCAACACCACCACCTGGGAGGTTCACAAACCCCCCACGATATATCCAACTCCACGAATTCAACCCCCACCAACGTTTCCTCGAAATCCGCCTTCATAGAGTTACAGCAACACGGTTACGGGCCTTTCAAAGGTGGTTACCAACATCCCCACCATTTCGGCAGCCCGGGGGGTCAGCAGAACCCCCACGAGGCGTCGGGGTTCCCGAGCCCTAGGTCCTTAGGTTATCCCTTTCCTCCTATGCACCAAAATACGTACGGATATCACATAGGCTCCTACGCTCCTCAATGTGCAAGTCCGCCCAAAGATGAAAAATGTGGTCTATCCGACGACCCTGGGCTGCGGGTGAACGGCAAGGGCAAGAAGATGCGCAAACCGCGCACCATCTACTCCAGCTTGCAGCTGCAGCAGCTCAACAGGCGGTTCCAGAGAACACAGTATCTAGCACTGCCGGAGCGAGCTGAGCTGGCTGCTAGCTTGGGCCTGACGCAGACGCAGAACAGTAGCGCTATTTCGTAA

Dll1_HCR_P1B2	CCTCgTAAATCCTCATCAAAGAACGCTACCCGGCCTCTCGTGGCG

Dll1_HCR_P2B2	AATTTTGAGGGATTTCTGGACTGCTAAATCATCCAgTAAACCgCC

Dll2_HCR_P1B2	CCTCgTAAATCCTCATCAAACGGGCTGCGGGGTCGACGGGCCCCC

Dll2_HCR_P2B2	GGTGGTCCAGCTCCTGGGTGGTCATAAATCATCCAgTAAACCgCC

Dll3_HCR_P1B2	CCTCgTAAATCCTCATCAAATTGCTGTAACTCTATGAAGGCGGAT

Dll3_HCR_P2B2	ACCACCTTTGAAAGGCCCGTAACCGAAATCATCCAgTAAACCgCC

Dll4_HCR_P1B2	CCTCgTAAATCCTCATCAAAGCTGACCCCCCGGGCTGCCGAAATG

Dll4_HCR_P2B2	GGAACCCCGACGCCTCGTGGGGGTTAAATCATCCAgTAAACCgCC

Dll5_HCR_P1B2	CCTCgTAAATCCTCATCAAATTCATCTTTGGGCGGACTTGCACAT

Dll5_HCR_P2B2	CCCAGGGTCGTCGGATAGACCACATAAATCATCCAgTAAACCgCC

Dll6_HCR_P1B2	CCTCgTAAATCCTCATCAAAGGTTTGCGCATCTTCTTGCCCTTGC

Dll6_HCR_P2B2	AGCTGCAAGCTGGAGTAGATGGTGCAAATCATCCAgTAAACCgCC

***> heat shock protein 67B1 (hsp67B1)***

lcl|XM_024094531.2_cds_XP_023950299.2_1 [gene=LOC112054669] [db_xref=GeneID:112054669] [protein=heat shock protein 67B1-like] [protein_id=XP_023950299.2] [location=57..1211] [gbkey=CDS]

ATGTCGCGGTATTTTGCGTTCTTCGCGCTCGTCGCGCTCGCGGCGGCCTACCCGGCCAGCGACGACTTCC

CTCGCCCCATCACAACAATAACTGAATCGGACTTCGAAGACCAATCGCCTTGGTTCCGATTCCCACCTTT

CGGAAACATCTTCGCGCCGCTAACGAAACTGTTCTCAAGTTTTGCGGAGATCGGACCGAAGATCGAAATT

GACGAAGACAAATTCCGTGTCATCGTAAACGTTAAGGATTACAAGAAGAAAGATCTGAAAGTTAAAGTGA

AAGGTGATTACATCCTCGTCCAAGGAGCGCACGAGGCTAAGCATGACGACCACGACTTGTTCGCGAGCCA

ATTCTTCCATACGTACAGCCTTCCGTTGAATGCTAGTGCATCAGATGTCACTGCTACTTTGTCTAGCGAT

GGATATTTGGATGTAACCGCTCCTGTGAATGGTGTCGATGACAAGAACAAGGTTGTTGATAGAGAAGTGC

CAATAGTTGAAAGTGGCAAGCCGTTGAAAGAGGATAAAGAAGATAGAGAACCTGTTGTACCCGTCGCTAG

TGCAGACCCAGTTGAAACTGTTGATAAGACAGAAAATGTTGATAAAATCGAAACTTTTGACACGCCAGTA

GAACCTTTAGCTAAGGTCGAAAATGCTGATAAAGTTGAAAATCTTGAACCACCCGTGGAACCGAGTGCCA

AAGCTGATAATGTTGATACGATTGAAGTTCCAAATGCACCAGTAGAACCTCTAGCTAAGGTCGAAAATAT

TAATAAAGTTGAAAATCGTATCGCGCCAGAGGAACCTATTACCAATGTTGAAAATGTTGATAAAGTTGAA

TCTCCCGCTGTACCAGAGGAGCCAGTGGCAAAGGTTGAAAATGTTGACAAAGTTGAAACTCCCGATGCAC

CAGTGGTGCCTGTTGCCAAGGATGAAAATGTCGACAATGTTGAAACTCCCGACGTATCAGTAGAACCTCT

ACCTAAGGTCGATAATGTCGATAAAGTCGAACAACTCCCCGAAGTGACAACTGCAACAGCTTCTGAAGGT

GAAGAAAAGACAGAGGCCCCGACGACCCAAGCGCCAGCACGTGAGGAGGTAAAAGAAGACATCAAAGTCC

CCCAGGACAACGAAGTAAACGAAATCCAGCCCTAA

B1_P1 (hsp67B1)_1	GAGGAGGGCAGCAAACGGAAGTTTACGATGACACGGAATTTGTCT

B1_P1 (hsp67B1)_2	GAGGAGGGCAGCAAACGGAAGCGCTCCTTGGACGAGGATGTAATC

B1_P1 (hsp67B1)_3	GAGGAGGGCAGCAAACGGAATTCAACGGAAGGCTGTACGTATGGA

B1_P1 (hsp67B1)_4	GAGGAGGGCAGCAAACGGAAATTCACAGGAGCGGTTACATCCAAA

B1_P1 (hsp67B1)_5	GAGGAGGGCAGCAAACGGAACTTTCAACGGCTTGCCACTTTCAAC

B1_P1 (hsp67B1)_6	GAGGAGGGCAGCAAACGGAAGTCTTATCAACAGTTTCAACTGGGT

B1_P2 (hsp67B1)_1	CAGATCTTTCTTCTTGTAATCCTTATAGAAGAGTCTTCCTTTACG

B1_P2 (hsp67B1)_2	AGTCGTGGTCGTCATGCTTAGCCTCTAGAAGAGTCTTCCTTTACG

B1_P2 (hsp67B1)_3	GTAGCAGTGACATCTGATGCACTAGTAGAAGAGTCTTCCTTTACG

B1_P2 (hsp67B1)_4	AACAACCTTGTTCTTGTCATCGACATAGAAGAGTCTTCCTTTACG

B1_P2 (hsp67B1)_5	CAACAGGTTCTCTATCTTCTTTATCTAGAAGAGTCTTCCTTTACG

B1_P2 (hsp67B1)_6	AAAGTTTCGATTTTATCAACATTTTTAGAAGAGTCTTCCTTTACG

***> rhomboid-related protein 3 (rho)***

lcl|XM_024085038.2_cds_XP_023940806.1_1 [gene=LOC112047794] [db_xref=GeneID:112047794] [protein=rhomboid-related protein 3 isoform X2] [protein_id=XP_023940806.1] [location=254..1180] [gbkey=CDS]

ATGAGCGGCAAGCGCACACGAAGCTATAGATGCGCCGTGCACCAGAGAGACCGCGAGGTCAGCTCTGAGA

ACGACTTCCATCTCCTTCTCGAAGACCCCACACTATTTGCCAGGATGGTGCACCTCGTAGCGATGGAGGT

GCTCCCCGAGGAGAGGGACCGCAAGTACTACCAGGAGCGGTACACGTGTTGTCCACCGCCCTTCTTCATC

ATATGTGTTACTTTACTAGAGTTAGGAGTATTCGCGTGGTACGCATGGGGTGCAGGCGGAGTTGCGGCCG

CGGCGGGCCCCGTGCCGGTCGACTCGCCTTTGGTGTATCGCCCCGACAGGAGACGAGAGCTGTGGCGGTT

TCTCACCTACAGTGTTGTGCATGCCGGATGGTTGCACCTCGCGTTCAACCTGCTTGTACAGTTAGCGGTT

GGACTACCTTTAGAAATGGTGCACGGAGCGGTGAGGTGCGGTGCGGTGTACCTTGCCGGTGTGTTGGGCG

GGTCGCTCGCAGCGTCGGTGCTAGATCCGGACGTGTGTCTCGCGGGCGCGTCGGGTGGTGTGTACGCGCT

GTTAGCCGCTCACCTAGCCAACGCGCTGTTGAACTTCCACGCGATGCGCTACGGCGCGGTGCGATTGGTC

GCCGCGCTTGCGGTCGCGTCGTGTGACGTCGGCTTTGCAGTTCACGCTAGGTATACTAAGGAAGCGCCGC

CGGTGTCGTACGCGGCGCACGTGGCGGGCGCGCTCGCCGGGCTCACCATCGGGCTGCTGGTACTGAAGCA

CGCGCAGCAGCGGCTGTGGGAGCGGCTGCTGTGGTGGGCAGCGCTGGGCGCCTACGCGGCGTGCACGCTC

TTCGCGGTACTGTACAACGTGTTCAGCGCGCCGGTGGACGAGCTGCATTACATGCCGCCCGACCCGCCGC

CCGACGCCGGCTTCTGA

B2_P1 (rho3)_1	CCTCGTAAATCCTCATCAAATGGGGTCTTCGAGAAGGAGATGGAA

B2_P1 (rho3)_2	CCTCGTAAATCCTCATCAAATAGTACTTGCGGTCCCTCTCCTCGG

B2_P1 (rho3)_3	CCTCGTAAATCCTCATCAAATACTCCTAACTCTAGTAAAGTAACA

B2_P1 (rho3)_4	CCTCGTAAATCCTCATCAAAAAGGCGAGTCGACCGGCACGGGGCC

B2_P1 (rho3)_5	CCTCGTAAATCCTCATCAAACATCCGGCATGCACAACACTGTAGG

B2_P1 (rho3)_6	CCTCGTAAATCCTCATCAAACGCTCCGTGCACCATTTCTAAAGGT

B2_P2 (rho3)_1	CGAGGTGCACCATCCTGGCAAATAGAAATCATCCAGTAAACCGCC

B2_P2 (rho3)_2	GGTGGACAACACGTGTACCGCTCCTAAATCATCCAGTAAACCGCC

B2_P2 (rho3)_3	GCCTGCACCCCATGCGTACCACGCGAAATCATCCAGTAAACCGCC

B2_P2 (rho3)_4	CTCGTCTCCTGTCGGGGCGATACACAAATCATCCAGTAAACCGCC

B2_P2 (rho3)_5	ACAAGCAGGTTGAACGCGAGGTGCAAAATCATCCAGTAAACCGCC

B2_P2 (rho3)_6	GGCAAGGTACACCGCACCGCACCTCAAATCATCCAGTAAACCGCC

***>uncharacterized protein LOC112047998 (nebula)***

lcl|XM_024085325.2_cds_XP_023941093.2_1 [gene=LOC112047998] [db_xref=GeneID:112047998] [protein=uncharacterized protein LOC112047998] [protein_id=XP_023941093.2] [location=167..1576] [gbkey=CDS]

ATGAGTCCGGGCGAGCGCGCCGCCCTTGGGCGCCGCGATGCCCCTCGCCCTCACTATGCTGCCACTGCAC

CGGGATTCCCGCCACCGTACTACCCACCCTACGGGGTGCCGCCGCATCACAACGCCTGGCTCGGAGCACC

GCTCATATCCATGACCGGGCGAACGCCACCGCCTCGCGAAAACCCCGTGTCCAAGCTCGCCATGCACGCG

CAGGGTGCTCCACTCATCATTCAGAACAGGCACGACCGCAGAAAATTTACCACCGTACCGGAGCAACGTA

ATCATCGCACAGTTTTGCCGCCAGAGAATATTACGAGAGACAGACCTGACCCAGAAAACACGAGGCACAG

AAATCAACATCAGCAACAACATCAGCAGCAAACGCGCCCCTCTCGCGGACGCAACAACAACACGGACGCG

GTCAGTGTCGCTAGTGATGAAAGTTCCGGCTCCACCAATTCTGAAACAATGTTGCCAAGAATTATAAAAC

CGCGTAAACGGCGTAAAAAAGATAGAAAACCTAATAATTTAGTGCCACACGATTTATCATCCGCTGCATT

AGAACTTGGCGATGTCGACCACTGTACAAATATGGGGTCGTCGTTGCATGGCATGTATGAGAAATCTTAT

TCCAGAGATATGGCATTACAGTACCATCTAAACGTAAATGTGGTAGACTACCCGGATCCGGCTCCAGAAA

CATATAGGGTGGCTGCGTTAGGAGAGGCTTTTGAGGCTACTCGTTTTGGATTAGCGGAGGCGGCTGCCCC

TTCAGAATCCGCCGCCAGCACGTGTCAGTGTAGATATTGCGACCCGGTTGGACAAATTTGGGATGCTAAT

TCCATTGCGCATTTACTGGACGAAAATTCAAGTGTTGACTTTGCACCCACGAAATTAGAAGACTCGATGA

AAATTGTGGGGCCACTGGGCAGGCGTGGAGCGGATACGATGTTACGGCGTAGTTGGAGTGACCCTTCGGC

GCGAACTCCGGAAAGGAAAACGAATAGCAGTGAGGCATATTCTGCTCCAAACCTCTATTCGTTATTCCCA

CCACCCAGAAGAACTAGCTCTCCGGAACCTAGATCGCCTCTAGCTTTGGAGATATCTTCTGAAATCGTCA

CTTCAATAAACGGCCACCGTGACTTGGAAATAAAGTTATTTTCGACCTCGCCACCGGCCTCGACCTCCGA

TCGTCGTTCTTTTTTCGACGATAAAAGTGAAAGTGCTGTAAACGATAGAAGTGCGAACGCGAACTCCGAG

CTAAAATCAAAAAGTGCAGTGAGCAGTGAAAAAGTGTCAAGTGCAAACAATGTTAAAAATAGTGTAAATC

TGGATTCAGGGTGCATGAAGGCACTCGGGGCCGGCGAGAACTCGCGCAACATTTGTGATTTAGCCTTAGT

TTCTGCCTAA

B2_P1 (nebula)_1	CCTCGTAAATCCTCATCAAAAGGGTGGGTAGTACGGTGGCGGGAA

B2_P1 (nebula)_2	CCTCGTAAATCCTCATCAAAGGTGGCGTTCGCCCGGTCATGGATA

B2_P1 (nebula)_3	CCTCGTAAATCCTCATCAAACCTGTTCTGAATGATGAGTGGAGCA

B2_P1 (nebula)_4	CCTCGTAAATCCTCATCAAATATTCTCTGGCGGCAAAACTGTGCG

B2_P1 (nebula)_5	CCTCGTAAATCCTCATCAAATGCTGCTGATGTTGTTGCTGATGTT

B2_P1 (nebula)_6	CCTCGTAAATCCTCATCAAAGCCGGAACTTTCATCACTAGCGACA

B2_P2 (nebula)_1	AGGCGTTGTGATGCGGCGGCACCCCAAATCATCCAGTAAACCGCC

B2_P2 (nebula)_2	AGCTTGGACACGGGGTTTTCGCGAGAAATCATCCAGTAAACCGCC

B2_P2 (nebula)_3	TACGGTGGTAAATTTTCTGCGGTCGAAATCATCCAGTAAACCGCC

B2_P2 (nebula)_4	TTTCTGGGTCAGGTCTGTCTCTCGTAAATCATCCAGTAAACCGCC

B2_P2 (nebula)_5	TTGTTGCGTCCGCGAGAGGGGCGCGAAATCATCCAGTAAACCGCC

B2_P2 (nebula)_6	TGGCAACATTGTTTCAGAATTGGTGAAATCATCCAGTAAACCGCC

***>Antennapedia (Antp)***

lcl|XM_052883780.1_cds_XP_052739740.1_1 [gene=LOC112057779] [db_xref=GeneID:112057779] [protein=homeotic protein antennapedia] [protein_id=XP_052739740.1] [location=671..1573] [gbkey=CDS]

ATGGAGGGCTGCGACCAGCAGCTCAGGCCCGGGCAGCACCACTACCCCGCGCAGCCCGCCCCTGGCATGCCTTACCCCAGGTTTCCGCCCTACGACCGGCTGGGCTACTACCAACAGATGGAACAGAACGGGTACCGGCCCGACAGTCCGACCCAGATGCACATGGGGCCGAAGTCCGACGGCTACGGGCCCAACGGCCACCAGCCGCCGACGCCGGCGGTGTACCCCTCGTGCAAGCTGCAGGCGGTGGCGGCGACGGCGGGGGGAGTGCCGGGCAGTCCGCCCCTGGAGCAAGCCCAGCAGATGCCGCACCACATGCACCCGCAGCAGCACATGCAGCACGGCATGCCGCCGCACCAGCAGCACGTCATGTACCCGGTGGACGACATGCAGCACCAGACGCAGATGCCGCCCATGCACCAGCAGTCCATGCACCCGCAGCAGGCGCCGCCTCAACAACCCCCGCCGAATACGAACGCGTCGCTCCCCAGTCCACTGTACCCTTGGATGCGAAGTCAATTTGAACGAAAGCGGGGACGGCAAACGTACACCCGGTACCAGACCCTCGAGTTGGAGAAGGAGTTCCACTTCAACCGATACCTGACGCGGAGGAGACGGATCGAGATCGCGCACGCCCTCTGTCTCACCGAGCGCCAAATCAAGATCTGGTTCCAGAACCGGCGCATGAAGTGGAAGAAAGAAAACAAGACCAAAGGCGAACCCGGCTCCGGCGATGAACCGGACAACATGAGTCCGCCCACCTCGCCCCAGTAA

antp1_HCR_P1B4: CCTCAACCTACCTCCAACAACCGGGCCTGAGCTGCTGGTCGCAGC

antp1_HCR_P2B4: GCGGGCTGCGCGGGGTAGTGGTGCTATTCTCACCATATTCgCTTC

antp2_HCR_P1B4: CCTCAACCTACCTCCAACAACATGTGCATCTGGGTCGGACTGTCG

antp2_HCR_P2B4: GGGCCCGTAGCCGTCGGACTTCGGCATTCTCACCATATTCgCTTC

antp3_HCR_P1B4: CCTCAACCTACCTCCAACAAGCTTGCTCCAGGGGCGGACTGCCCG

antp3_HCR_P2B4: GGGTGCATGTGGTGCGGCATCTGCTATTCTCACCATATTCgCTTC

antp4_HCR_P1B4: CCTCAACCTACCTCCAACAAACTGCTGGTGCATGGGCGGCATCTG

antp4_HCR_P2B4: GAGGCGGCGCCTGCTGCGGGTGCATATTCTCACCATATTCgCTTC

antp5_HCR_P1B4: CCTCAACCTACCTCCAACAACTGGTACCGGGTGTACGTTTGCCGT

antp5_HCR_P2B4: GTGGAACTCCTTCTCCAACTCGAGGATTCTCACCATATTCgCTTC

antp6_HCR_P1B4: CCTCAACCTACCTCCAACAATCCACTTCATGCGCCGGTTCTGGAA

antp6_HCR_P2B4: GTTCGCCTTTGGTCTTGTTTTCTTTATTCTCACCATATTCgCTTC

> ***Notch***

lcl|XM_052881333.1_cds_XP_052737293.1_1 [gene=LOC112052049] [db_xref=GeneID:112052049] [protein=neurogenic locus Notch protein isoform X1] [protein_id=XP_052737293.1] [location=218..7600] [gbkey=CDS]

ATGTTGTGCTACAAGATGTGGTCGAGAAACTTTATACCGGATTATGGGATTAACTTGCTCTCCATCGTCC

TGCTGACCACCTTAGCGGCCACAATCCAAGGAGCCGAAGGTTTCGTGTCGTGTTCACCGTCGCCGTGCAA

AAATAGTGGAACGTGTCTCTCTACTACGCGGGGGGAGTATTTTTGCAATTGTACATCGCGCTACGCCGGC

GAGTTCTGCCAGCATCTCAACCCTTGCCACAGCGAATCAAGCCCTCGGTGTCAGAATGGAGGCTCATGCA

GGGTCAGGCCCGGCGCCGGGGGAGGCCCGCCCTCCTTCGCCTGCGACTGCCCTCTCGGATTCAGTGCCTC

GTTGTGCGAGATAAGAGTGCCTGCTGCGTGTGACTCCGCTCCGTGTTTGAACGGTGCTACGTGTCGATTG

ACGTCACTTGACACCTTCGAATGTGATTGTCCTCCAGGGTACACAGGCGAAGAATGTTCACACGAAGACC

ACTGCGCATCCCAACCCTGCAGGAACGGAGGACGTTGCGTTGCAGACAATACAACGACAGTCGGCTACTC

CTGCGCTTGTCCACCTGGCTTCACCGGCGCGCGATGTACTGAAGACGTGTTCGAATGCTCGAGTGGATCC

GGTCCTTGTCACCATGGGAGATGCTTCAACACCCACGGTTCTTACACCTGTGTTTGTGAGCCTGGGTATA

CAGGCAGGGATTGTGACACGGAGTACGTGCCCTGTGAACCGTCCCCTTGCTTGCATGGTGGGAGGTGCAC

GCAGCTGGATCAATTACGATACGAGTGTGATTGTCCTACTGGTTACCGAGGTCAGAACTGCGAGATCGAC

ATCGACGACTGTCCGGGCCACCTCTGTCAGAACGGTGCCACGTGCGTGGACGGATTGAACTCCTACAACT

GCGAATGCCCTCCCACCTTCTCCGGCACTCTGTGCGAGACGGACGTCGACGAATGCGCTTTGAGGCCGCT

GGTATGCCAGAATGGAGCGACATGTACGAACTCAGCAGGCGGGTTTTCGTGTATCTGCGTCAACGGGTGG

ACTGGTCCAGAATGTTCCGTCAACATCGACGACTGCGCCGGCGCAGCTTGTTTCAACGGCGCAACCTGTA

TAGACAGGGTTGGGGCCTTCTACTGCAAGTGTACACCTGGGAAGACTGGTTTACTCTGCCACTTGGATGA

CGCGTGCACATCTAACCCGTGCCACGCCGACGCGATCTGCGACACTAGCCCCATCAACGGATCCTACACT

TGCTCCTGTGCGTCCGGGTATAAGGGCTTGGATTGTTCGGAAGATATTGACGAGTGTGAACAAGGCTCAC

CCTGCGAACACGACGGCATCTGCGTCAACACCCCTGGCTCCTTTGCCTGCAACTGCTCCGTCGGGTTCAC

CGGACCGCGTTGTGAGACCAACGTCAACGAGTGCGAGAGTCACCCCTGTAGGAACGATGGCTCCTGCTTG

GACGACCCCGGCACCTTCCGATGCGTATGCATGCCTGGTTTCACTGGAACGCAGTGCGAGGTGGAAATAG

ATGAATGCGCCAACAACCCTTGCCTCAACGGAGGCGTTTGCCACGACATGATCAACGCCTTCAAATGCAC

CTGCGTTATTGGATTCACCGGTGCCCGTTGTCAAGTTAACATAGACGACTGCGCGTCGAGTCCGTGTCGC

AACGCAGGTACTTGCCACGACTCCATCGCTGGATACACCTGCGAATGTCCTCCGGGATACACGGGGATGT

CTTGCGAGACCAACATCAACGACTGTCTCTCCGCGCCGTGCCACCGAGGCGAGTGCATCGACGGCGACAA

CAGCTTCACATGCAATTGCCACCCGGGCTACACCGGCCGCGTGTGCCAGACGCAGATCAATGAGTGCGAG

TCCAACCCCTGCCAGTTCGGAGGACATTGCGAGGATCTCATAGACGGATACCAGTGTAGATGCAAGCCGG

GCACGTCGGGCAGGAACTGTGAGATCAATGTCAACGAATGCTACTCCAACCCGTGCAGAAATGGAGCCAC

GTGTATTGATGGGATCAACAGGTATACGTGTGAGTGCATCCCCGGCTTCACTGGCCAGCACTGCGAGACC

AACATCAATGAATGTCTGTCCAACCCGTGCGCTAACGGCGGCAAGTGCATCGACCGCATCAACGGGTTCC

GCTGCGAGTGCCCCAGGGGATACTATGATGCGAGATGCCTTTCCGACGTGAACGAATGCGCCTCAAATCC

ATGCATCAATGGCGGCACTTGTGAAGACGGCGTCAACCAGTTCATCTGCCATTGTCTCCCGGGATACGGA

GGCCAGCGCTGCGAGCGCGACATAGACGAGTGCAGTTCCAACCCGTGCCAGCATGGCGGCACGTGCCACG

ACCGGCTCAACGCGTACAAGTGCGACTGTGTGCTGGGATTCACGGGTGTGAACTGCGAGACCAACATCGA

CGACTGCGCCGGCAACCCGTGCCTGCACGGCGGCTCGTGCATCGACCTCGTCAACGGGTACCGCTGCGTG

TGCGCGCCGCCGCACTCCGGCCGCAACTGCGAGAACACGCTCGACCCCTGCCTGCCCAACCAGTGTCGGC

ACGGTGGTCGGTGCATTGCGGAGGCGTCGTACGCAGAGTTCACGTGCCAGTGCCCGGTGGGCTGGACGGG

CGCGCTGTGCGAGCGCGACGTGGACGAGTGCGCGGTGACGGCGCCGTGCCACAACGAGGCCACGTGCATC

AACACCGAGGGCACGTACGCGTGCCTCTGCGCGCGCGGCTACGAGGGCAAGGACTGCGCCATCAACACCG

ACGACTGCGCCTCGTTCCCCTGTCAGAACGGAGCCACATGTCTGGACAGTATAGGCGACTACAACTGCGT

GTGCGCCAACGGTTTCGCGGGCAAACACTGCGAGATTGACATCGACGAGTGCCAGTCCCGACCGTGCATG

AACGGCGCCACTTGCAACCAGTACGTGGCATCCTATACGTGCACCTGCCCGCTCGGCTTCTCGGGCATCA

ACTGTCAGACCAACGACGAGGACTGCACCGAGTCCAGCTGCATGAACGGCGGCTCCTGCATCGACGGCAT

CAACTCCTACAACTGCTCCTGCCCTCCTGGCTACACGGGCTCGAACTGCCAGTTCCGCATAAACATGTGC

GACAGTTCCCCCTGCGACAACGGCGCCACGTGCCACGACCACGTCACGCACTACACCTGCCACTGCCCCT

ACGGGTACACCGGCAAGCACTGCGAGGACTTCGTGGACTGGTGCGAAAACAATCCATGCGAGAACGGCGC

GACGTGTTCACAAAAAGGCCCGCAGTACACCTGCACTTGTGCGCCCGGCTGGTCCGGCAAACTCTGCGAT

GTTGAAATGGTCTCCTGTAAAGATGCGAGCATTCGGAAAGGAGTGAAACTAAAGCAACTGTGCAACAACG

GATCTTGCGAGGATATCGGCAACTCCCATCGCTGCCACTGCCTCGATGGATACACGGGCTCTTATTGCCA

GAAGGAAATCAACGAGTGCGACTCCGCCCCGTGCCAAAACGGAGCGGTTTGCAAAGATCTAGTTGGTACC

TATCAGTGCCAATGCACCAAAGGTTTCCAAGGACAGAACTGCGAGTTGAACGTCAACGACTGCCTTCCCA

ACCCTTGTCAGAACGGAGGAACCTGCCACGATCTGATCAATAACTTCTCCTGCTCTTGTCCGTTTGGAAC

TCTTGGAAAAATTTGCGAAATCAACGTCAACGATTGCAAACAAGACGCTTGCCACAATAATGGCTCTTGT

ATCGATAAAGTTGGCGGTTTTGAATGCAAATGTCCTGCTGGTTTTGTAGGTCCGCGTTGTGAAGGCGATA

TTAACGAGTGTTTATCCAATCCGTGCTCAAATCCTGGAACGCAAGATTGCGTGCAACTAGTCAACGATTA

TCACTGTAATTGTAAACCTGGGTTTATGGGCAGGCATTGCGATGCTAAGGTCAATTTCTGTGCAAACTCT

CCGTGCCAAAATGGCGGCATTTGCACTGCTATCCAAGGTGGACACGAATGTCTGTGTGGCGATGGATTTT

ATGGCAAGAATTGTGAATATTCAGGATACGCGTGTGACTCGAATCCTTGCCAAAATGGAGGCTATTGCCG

GACAATGGAAAACGGAGACTACGTCTGCAACTGCCCGTCAGGTTTGTCCGGTGTCAACTGCGAAATAGAT

TCTATGAACGAATGCCTTTCGAATCCTTGTAAGCATCCCGAAGCCCGGTGCATAGACAAGCCAGGGGACT

ATTTGTGTTACTGTCCCAGACAGTGGACCGGTAAGAACTGTGATATACACGACCCTCATTCTAGAGGAGG

CTACGGAAGTCCGGTATCTGGTGTGTACGGCAAAAATCCAGTTCTGACTTTGCAAGAATTAGACTTGGCA

TTCCAAAGAGAACAATGCGTAAAATTGGGCTGCAAGGAGAAACAAGGCGACCATCATTGCGATGAGGAAT

GTAACACGTTCGCATGTGAATTTGACGGAAACGATTGTTCACTCGGTATTAACCCATGGGCGAACTGCAC

GGCTCCGATAAAATGCTGGGAAGTGTTCATGAACGGGGAATGTAACGAAGTTTGCAACACTCAAGCTTGT

CTCTTCGACGGCAGAGACTGTGAGAAGTCCTTGCAAAAATGTAACCCAGTGTATGACGCGTACTGTCAAA

AGCACTATGCTAACGGCCATTGCGACTATGGATGCAACAATGCTGAGTGTAACTGGGACGGCCTGGACTG

TGAAAACGAACCACCCGATCTAGCTGAAGGTGTAATGTCAGTTATATTGCTGATGGATATGCGAACGTTC

AAGGAAAATTCAATTGCATTCTTGCGAGACTTGGGCCATCAACTACGGACAACAGTACTAATAAAGAAAG

ACCACTTGGGCAACGACATGGTGTTCCCATGGAAAGGTTCTACCGATGTCGGTCTCCAAGACACTGAATT

CGGGAAGAAACACCACATCGTGTATACAGAAAGAGGTCAATCGGGAGTGCAAGTTTACCTAGAAATCGAC

AACAGAAAGTGTACAATGATGTCTGGGTCTGAATGTTTCTTCTCGGCACGTGAAGCAGCAGATTTCTTAG

CAGCTACAGCGTCGAAGCATTCCTTGTCACCAGATTTTCCAATATTCCAAGTAAAGGGGGTCACTACTCC

CGAAGATGGCGATATCCCAACAAACTCCAAATACGTATTCATCGGTGTTATCCTCGTGCTGCTCGCTGGT

CTCCTTATTGGAGTTCTAGTCACAGCACAGAGGAAACGCGCTGCCGGTATCACATGGTTCCCAGAAGGAT

TTATTCGGAATAATTCGTCAACTAGACGACGTTCCCGACGTAGAGGTCCTGATGGCCAGGAAATGCGGAA

CTTGAATAAGGGATCTATTGGTTGTATAGACGTTGATATGAATGGTGGTCATATGGGTCCTCCTCACTGG

TCTGATGATGATGAAGATGGTTCCGCTCCTCCGAGGGCTAAACGACCTCGAGGTCCAGGTGACGCGAATG

GTGCCCCAGGGGGTTATGCCTCTGATCACACAGCTATTACGGATTATGAAGAAGCTAGTCACGAGCCTAG

GGTCTGGACACAGCAACACTTAGACGCTGCAGACATCAGAGTTCCACCGATGATGACGCCGCCTGCTATC

CACGATGGTCACGTAGACGTAAACGCGAGAGGTCCGCTCGGGATGACGCCTTTAATGGTGGCCGCCATCA

GAGGCGGAGGCTTGGACACCGGCTCTGATGTGGAAGAGGAGCAAACAGCGCACATCATATCGGAGCTGGT

GGCTCAAGGAGCTCAACTCAACGCTGCCATGGATAAGACTGGAGAGACCAGTTTGCATCTCGCTGCAAGA

TACGCACGCGCGGACGCAGCGAAGCGTCTCTTGGACGCCGGCGCGGACGCTAACTCGCAGGACAACACAG

GCAGAACTCCGTTACACTCTGCGGTCGCGGCGGACGCCATGGGCGTGTTCCAGATCCTGCTGCGGAACAG

GGCGACCAATCTCAACGCTAGGATGCATGATGGAACTACACCACTTATATTGGCTGCGAGGCTCGCGATC

GAAGGCATGGTGGAGGACCTAATAAACGCCGACGCGGACATCAACGCGGCGGACAACAGCGGCAAGACAG

CGCTGCACTGGGCCGCTGCCGTCAACAACGTGGACGCGGTCAACGTACTACTCGTCCACGGCGCCAACAG

GGACGCACAGGATGACAAGGACGAGACCCCACTATTCCTCGGCGCGCGGGAAGGCTCGTACGGCGCCTGC

AAAGCGCTGCTAGACGCCATGGCGAACAGAGAGATCACGGACCACATGGACAGATTGCCGCGGGACGTCG

CACAGGAGCGCATGCATGACGACATCGTGCGGCTACTGGACGAGCACTGTCCTCGGCCGCCGCAACACCA

TCATTTGATGGCTTCACCGAACCCCCACCCGCATCTGATCAGTCAGCCGACAGTAATCAGCTCGACCGCC

AGCAAGGGGAAGCCCAAAAAGGCGCGCGCGAAACCCGGCCCCGACAGTCCACAGGAGCAGGCGTACGACA

ATAACATGCAGCCCACGCAAATAAGGCGGAAGCCCAGCGTAAAAAAGAACAATAAGAAGGTGGCGCAGGA

GGTGCCACAAAGCGTAGAAAGTTTGGGATTCAGCCTGAGTCCCGTCGAATCACCGCTGCAAAATTTACAG

GACCTACCGTCGCCGTACGACGCGACGTCGCTGTACTCGAACGCGATGGCGCAGTTCCCGGCTGTGGAGC

AGCTGGTACAGCACAAGCAGCCGCCCAGCTATGACGACTGCGTCAAGTACGTGTGTCCGCAGACGGGGCA

GACGCTGCAGTACGGCGGCGGGGCGCTCGGCGCGTCGCTCTCGCCGCCCTACTCCAACCACTCGCCCACG

CACAGCAACCAGACCACCTCGCCGCACGCATATATAGGCTCGCCGTCGCCGGGCAAGTCGCGACCGTCGC

TGCCGACGTCGCCGGCGCACATGGCGGCGCTGCGCCACCACCACCTCGACGCCGCCGCCTACTCTGCGCT

CGCGCACCTCAGTGCGAACGGGCAGCACAACGCTCAGCTGCAGGCGGTGATGACGCACGCCGCAGCGCTG

GGGCAGGTGCAGGGGGCGCAGCACCCCGCCATCACGAACCTCATGTCTGGTCTTTATGGAAGTTGGGGTA

TGGGAGACACGTTCCCGACCCCCTCGCCAGAGTCCCCGGATCACTGGTCGACCCCTTCGCCGCAGACCCC

GCTCACTCAGTCCCCCCACTCGGATTGGTCGGACCGCGCCGCCTTGTCGCCTAACGACATACAACAGACA

AACAAAGGTGCCACTGAAGCGTTTTATATATAA

B1_P1 (notch1)_1	GAGGAGGGCAGCAAACGGAAGTTTCTCGACCACATCTTGTAGCAC

B1_P1 (notch1)_2	GAGGAGGGCAGCAAACGGAACTTGGATTGTGGCCGCTAAGGTGGT

B1_P1(notch1)_3	GAGGAGGGCAGCAAACGGAACGCGTAGTAGAGAGACACGTTCCAC

B1_P1 (notch1)_4	GAGGAGGGCAGCAAACGGAAGTGGCAAGGGTTGAGATGCTGGCAG

B1_P1 (notch1)_5	GAGGAGGGCAGCAAACGGAAGCGGGCCTCCCCCGGCGCCGGGCCT

B1_P1 (notch1)_6	GAGGAGGGCAGCAAACGGAACACGCAGCAGGCACTCTTATCTCGC

B1_P2 (notch1)_1	CAAGTTAATCCCATAATCCGGTATATAGAAGAGTCTTCCTTTACG

B1_P2 (notch1)_2	GTGAACACGACACGAAACCTTCGGCTAGAAGAGTCTTCCTTTACG

B1_P2 (notch1)_3	GATGTACAATTGCAAAAATACTCCCTAGAAGAGTCTTCCTTTACG

B1_P2 (notch1)_4	ATTCTGACACCGAGGGCTTGATTCGTAGAAGAGTCTTCCTTTACG

B1_P2 (notch1)_5	CGAGAGGGCAGTCGCAGGCGAAGGATAGAAGAGTCTTCCTTTACG

B1_P2 (notch1)_6	GCACCGTTCAAACACGGAGCGGAGTTAGAAGAGTCTTCCTTTACG

***>runt***

lcl|XM_024081366.2_cds_XP_023937134.1_1 [gene=LOC112045254] [db_xref=GeneID:112045254] [protein=segmentation protein Runt-like] [protein_id=XP_023937134.1] [location=298..1530] [gbkey=CDS]

ATGCACCTTCCGCACGCAAGCCCCGCGGCCCCGAGCATGGCGGACGTCTACTCGCACATCCACGAGTACTACCGGCAGAGCCACGGCGACCTGGTGCAGACCGGCTCCCCAGCAGTGCTCTGCTCGGCTTTACCCGGACACTGGCGGTCCAACAAGTCCTTACCGGTAGCCTTCAAGGTCGTCGCCCTCGACGATGTGCAAGACGGGACCTTAGTGACCATTAAAGCTGGGAACGATGAGAATGTGATGGCGGAGATGAGGAACTGCACCGCTGTGATGA

AGAACCAGGTGGCAAAGTTCAATGACCTGAGGTTCGTGGGTCGCAGCGGGCGTGGCAAGTCCTTCAGCCT

GACCATCACCATCAGCACGTTCCCCAGCCAAGTGGCCTCCTACACCAAAGCTATCAAGGTCACCGTGGAC

GGACCCAGAGAACCCCGGACCAAGCAAAATTACGGTTACGGTCACCCTGGCGCCTTCAACCCGTTCCTGC

TCAACCCGGGCTGGTTAGACGCTGCGTATCTGAACTACGCCTGGGCGGACTATTTCAGGCCGCCGCAATT

GAGGGATCAAGCTGCCCTTGTTAAAGGTGGGGCTGCACCCATAACGACACCACCGCTGGCGTTGCCCGGA

GCGGAGCTGTTCCCGTTTCCACCGGCGATGGCGAGCTTACCGACCGGAGGGTTGATCCCTCCGCCCGGAGCGTTCCTACCGCCGAACGGACTTCTGGCTTTCCCGCCCCATCCCGCAGATCTTGCGTTAAAAAGTTTACC

ACCGGAGCTGACGTTGAAAAACGGCATCAGTCCCTACGACGCGTTAAGGCAGTTCCAAAGCAACGTCTCC

TCGATGGACACTTCGAGCGCTCGCTTGTCTCCGACGAGCAGCAGACAGAGTGGAAGCCCGAGGAGTGTCA

CAAACGCCAGTCCGAGATCGAAGGCTGATTCGAGGTCGGAAGTCAACTCGACTCACGAAGCAACGATATC

TGACGAATCGGACGAAGAGCCGATCGAAGTCGTCAAATCGGCGTTCCACCCGACGAGACCGGCAAACGTG

GAACTGCAGGAGATGAAGCAAGTCCAGGCGGCGGACTCCACCGTGGCGGACAAGCCCCGGACGCGCAACGAACTAAAAGCGCCTTCACAACGCACCACTCGTGTTCTCTCCACTAGTCCCACATCGACCAAAATTACAAA

TGGCAGCATATCGAACCATAAATCCGTGTGGAGACCATATTGA

| B3_P1(run)_1 | GTCCCTGCCTCTATATCTTTGGCCGCGGGGCTTGCGTGCGGAAGG |
| --- | --- |
| B3_P1(run)_2 | GTCCCTGCCTCTATATCTTTTCTGCACCAGGTCGCCGTGGCTCTG |
| B3_P1(run)_3 | GTCCCTGCCTCTATATCTTTGCTACCGGTAAGGACTTGTTGGACC |
| B3_P1(run)_4 | GTCCCTGCCTCTATATCTTTCTCATCGTTCCCAGCTTTAATGGTC |
| B3_P1(run)_5 | GTCCCTGCCTCTATATCTTTTCAGGTCATTGAACTTTGCCACCTG |
| B3_P1(run)_6 | GTCCCTGCCTCTATATCTTTTGGCTGGGGAACGTGCTGATGGTGA |
| B3_P2(run)_1 | GTGCGAGTAGACGTCCGCCATGCTCTTCCACTCAACTTTAACCCG |
| B3_P2(run)_2 | CCGAGCAGAGCACTGCTGGGGAGCCTTCCACTCAACTTTAACCCG |
| B3_P2(run)_3 | ACATCGTCGAGGGCGACGACCTTGATTCCACTCAACTTTAACCCG |
| B3_P2(run)_4 | GCAGTTCCTCATCTCCGCCATCACATTCCACTCAACTTTAACCCG |
| B3_P2(run)_5 | TGCCACGCCCGCTGCGACCCACGAATTCCACTCAACTTTAACCCG |
| B3_P2(run)_6 | TTGATAGCTTTGGTGTAGGAGGCCATTCCACTCAACTTTAACCCG |

***>sex combs reduced (scr)***

lcl|XM_024098318.2_cds_XP_023954086.1_1 [gene=LOC112057777] [db_xref=GeneID:112057777] [protein=homeotic protein Sex combs reduced] [protein_id=XP_023954086.1] [location=378..1439] [gbkey=CDS]

ATGAACGACCTAAATTACAACGCAGGGATGAGTTCCTACCAATTCGTCAACTCCCTGGCGTCCTGCTATG

GGAATCAGGTGCCAGGTCGCACGGGAACGCCTGTAGAACAAGCAGGTCACCCTGGATTACCGACGCCTGG

AGCGGACTATTATAATCCAAATGCTACAGCTTCGTATCCAAATACATGTTATTCACCACCACAGGTGGGT

CATCATTACCCGCAACATCCGTATGCGACGCCGGCAGCCGGTGCTCACATGCAACCACAGACTATGATCG

ACTACACACAACTGCATCCACAAAGACTTGGCGGCACGGCGAGCCATGTGCACCAACATTCAAATCCAAG

CCCTGGAGCGCTTTCCCCGAATCTAATGTCGACGCCGCCCAGTCAAGCCGCCAGTGCAAGTTGTAAATTC

GCCGATTCAACCTCTACAACAGGTTTAGCATCGCCGCAGGACTTGTCAACTTCTTCAGGACCAGGTAGAA

CATCACCGGGTTTCGGAAATGTCGGTAATCCTAGCGGCACGACCAGTACTAAACTCGGATTGACCACTCC

GATCGCTTCCCCAGTGGAGCACAAGGCGAATATCAATCAAAACATCTCGAGCCCTGCTTCTAGTACGTCT

AGCAACGAAAGCAATGAAGCGAACAATTCAAGTTCAAGCAATACGAAGAACGCGAAAGCTTCAGGCAACG

CTCAAGCAAATCCACCGCAGATTTACCCATGGATGAAAAGAGTTCACCTAGGCCAAAGTACTGTTAACGC

GAATGGAGAGACGAAACGACAGAGGACGTCGTACACCCGGTACCAGACACTCGAACTCGAGAAGGAGTTC

CACTTCAATCGGTACCTGACGCGGAGAAGACGGATCGAGATCGCGCATGCCCTCTGCCTCACAGAGCGCC

AAATCAAAATCTGGTTCCAGAACCGGCGCATGAAGTGGAAGAAGGAACATAAAATGGCATCGATGAATAT

CGTCCCCTACCATATGAATCCGTATGGCCATCCTTACCAATTCGACCTTCACCCGAGTCAATTTGCGCAC

TTGTCAGCATAG

B4_P1(scr)_1	CCTCAACCTACCTCCAACAACGTCGCATACGGATGTTGCGGGTAA

B4_P1(scr)_2	CCTCAACCTACCTCCAACAACAAGTCTTTGTGGATGCAGTTGTGT

B4_P1(scr)_3	CCTCAACCTACCTCCAACAAGACATTAGATTCGGGGAAAGCGCTC

B4_P1(scr)_4	CCTCAACCTACCTCCAACAATGCTAAACCTGTTGTAGAGGTTGAA

B4_P1(scr)_5	CCTCAACCTACCTCCAACAAGATTACCGACATTTCCGAAACCCGG

B4_P1(scr)_6	CCTCAACCTACCTCCAACAATTCGCCTTGTGCTCCACTGGGGAAG

B4_P2(scr)_1	TGGTTGCATGTGAGCACCGGCTGCCATTCTCACCATATTCGCTTC

B4_P2(scr)_2	GTTGGTGCACATGGCTCGCCGTGCCATTCTCACCATATTCGCTTC

B4_P2(scr)_3	GCACTGGCGGCTTGACTGGGCGGCGATTCTCACCATATTCGCTTC

B4_P2(scr)_4	TGAAGAAGTTGACAAGTCCTGCGGCATTCTCACCATATTCGCTTC

B4_P2(scr)_5	CGAGTTTAGTACTGGTCGTGCCGCTATTCTCACCATATTCGCTTC

B4_P2(scr)_6	GCAGGGCTCGAGATGTTTTGATTGAATTCTCACCATATTCGCTTC

***> proboscipedia (pb)***

lcl|XM_024098314.2_cds_XP_023954082.1_1 [gene=LOC112057771] [db_xref=GeneID:112057771] [protein=homeotic protein proboscipedia] [protein_id=XP_023954082.1] [location=406..2439] [gbkey=CDS]

ATGCAAGAGATATGCAATACCGCGCTGCCGCTAGACAGTAACCCCTTGGTGCGCAAGATAAAATGCGAAG

AGTTCCCCGTGAGAGGAAAGCAAGCAGCAAGAATCGGTATGGGGGTCCGCGATGATATGGAAGAGGAGTT

CGTGGTGAAGACGAGGAATACGCCCCCAATGGGGCCCGCTCCAGGACAAGAGCATCCTCGTATGGAGCAT

GGGGGTAATGGGTTCTGGCTGGCTGCGGTCACCGCAGCCGGCGCTCCTCATCCAATGCTCGAAACGTGTA

CAGAGACTGGGTTTATTAATAGTCAACCTTCTATGGCGGAGTTTATGACGGCGCTTCCACAGCTGGGAGG

AGGCGAATTAAGTCCGCAGCACACACCTCCCGGTTACGCGCCTGACCTGCCATCGCCTGGTGGCGGTCTC

AACGTCCCCGAGTACCCTTGGATGAAGGAGAAAAAAACGACAAGAAAAAACAGCCAACAAGAAAATGGCT

TGCCGCGACGATTACGAACCGCCTACACGAACACACAACTGCTGGAATTAGAAAAAGAGTTTCATTTCAA

CAAGTACCTGTGCAGGCCTAGAAGAATCGAGATTGCCGCGTCCCTTGATCTAACAGAAAGACAGGTCAAA

GTGTGGTTCCAAAATAGAAGGATGAAACACAAAAGACAAACGGTTAGCAAAAGTGAAGATGGTGACGATA

AGGATTCGACAACCTCAGAAGGAGGCAAAAGTTCTAAAACAGGGTTAGAAAAGTTCCTCGATGACGATGG

CCCATTATCTGGCAAGAAGAGTTGTCAGGGTTGTGAACTACCACCTGGTGTTCTGTGTTCTCCATCAGAG

GATCTTCCGGAGTTAGCCTCACGAACACGTAACAACAACACTCCGAGTGCTACGAATAACAACAGCTTTG

CAAGCGATGGAGCTTCCAGCGTAGCTTCATCTTCTTCCCTAGATAAACTAGCGGAGGAAGATTCCCGAGA

AGGACTTCCGCCAGCAAACCCTGTAACGACAGCACCTAGAAATCTAGCTAAAAGAATTAAACAAGAGTCA

AGAAAGCGATCCCCTTCATTGGATGCGACGGGTTGTAAAGTGTCACCGTCGTCTTCGAAAGACGGTCTCG

TCGGTATAGCTGGCCTATCAGACGGTGGGAAATTTTCTTCAGTCAATTTGACACCGTCATCTACTCCTGG

CACACCATCAAGTATGCACCAAAGTCCTCTTGGTCTCTACCCTCGACCGTCGCCGCCCCATGCACCACCA

GGACCACCCCTTCCTCAAGCTGTACCTAACTCGATGCCACCGTACGTAATTAGAGGGAACGCTCCACCTG

GCCAATTCGTACCACATCCAGACTTTCGAATGGACCCAAAGCAATTTGTCGGTAAACTGGCCCAATATCC

TCAGAGTAACAGGTCCTACGACGCTTACGGACCTGCATTACAAGGTACAGAACAACACGCTTACACAAGG

AATCAACATCACTCAAGACAACACGATAGCTCTCCTTCTACGAGACCGACAAACGGGATAGGGTCGAGAC

AATCCTACCCACATGAAATGTACCAAAACTACGCTTACGGGAGTTATGCAAAAGATCAAGTTGCGTACGG

TCATCCTAGCTACGAACAGGGCCAAGGTTACCCAGGTGAACATATTGGCTATGCAAATAGTCACTACGGA

TACCACTATCACGAAAGCGGACAACATGACGCCTCACACGGTTACTACGGTAGTGATGGACAAAAGAATG

TTCATGGCGCCGACTACACAAAGAACGCCTACTATGACGCAAGCTCATATGGAAGTCAACAAGGAAGTGC

AACGGCAGCGAGCTATGGCGCGGGCACACAAGGATCCGCGTCAGGGGAGGCTTATGGCAGCGGCGAATGTGGCGAAACCTACGGTTCTTTCCAACAGTTTTACGAAGCAACACACACAACGCCTGCCACCGGTGACAATT

CGAACTCTTCATCAGACTTTCACTTCCTGAGCAATCTAGCGAACGACTTCGCTCCGGAATACTACACCAT

TTGA

B4_P1(pb)_1	CCTCAACCTACCTCCAACAAGACCGCAGCCAGCCAGAACCCATTA

B4_P1(pb)_2	CCTCAACCTACCTCCAACAAAAGGTTGACTATTAATAAACCCAGT

B4_P1(pb)_3	CCTCAACCTACCTCCAACAAGGAGGTGTGTGCTGCGGACTTAATT

B4_P1(pb)_4	CCTCAACCTACCTCCAACAACTCCTTCATCCAAGGGTACTCGGGG

B4_P1(pb)_5	CCTCAACCTACCTCCAACAATCGTGTAGGCGGTTCGTAATCGTCG

B4_P1(pb)_6	CCTCAACCTACCTCCAACAATCGATTCTTCTAGGCCTGCACAGGT

B4_P2(pb)_1	CATTGGATGAGGAGCGCCGGCTGCGATTCTCACCATATTCGCTTC

B4_P2(pb)_2	GAAGCGCCGTCATAAACTCCGCCATATTCTCACCATATTCGCTTC

B4_P2(pb)_3	GGCGATGGCAGGTCAGGCGCGTAACATTCTCACCATATTCGCTTC

B4_P2(pb)_4	TTGGCTGTTTTTTCTTGTCGTTTTTATTCTCACCATATTCGCTTC

B4_P2(pb)_5	CTTTTTCTAATTCCAGCAGTTGTGTATTCTCACCATATTCGCTTC

B4_P2(pb)_6	TCTGTTAGATCAAGGGACGCGGCAAATTCTCACCATATTCGCTTC

***> abdominal-A(abA)***

lcl|XM_024098330.2_cds_XP_023954098.1_1 [gene=LOC112057784] [db_xref=GeneID:112057784] [protein=homeobox protein abdominal-A homolog isoform X4] [protein_id=XP_023954098.1] [location=2967..3992] [gbkey=CDS]

ATGAGTTCCAAGTTCATCATCGATAGCATGCTCCCCAAGTACCACCAGCAGTTCCATCACCAGAACTTGT

TCGCGGGAGCCGGAGCTTCGCCGATCGAGGCGTCGTTGTCGTCGTCGCTGTCGTCGTCGTTGTCGACGTC

GCTGTCGTCGTCGCTGTCTGGCGGGCTGGGGGCGGCCGCGCTGGGGGCCGGCTCGCCGGGTGCCGGGAGCCCACAGCGCTCGTCGTCGTCGTCGTCGGCGTCGCCGGGCGCGCCGGCGAGGATGTATCCGTACGTGTCGCACCACCAGCAGTTCGGGGGGTCGGTGCCGTTCCCCGCGGGCGGCGGGCTGGCGGACGACAAGAGCTGCCGGTACCCTGGGGCCGTGAGCGGCGACCCCATGGTGAACTACGCGCTGGGGCAGCACAACGGCGGCGCGGCGGTGTCGGCCGCGTCGGCCAGCATGGCGGCGGCGGCGCAGTTCTACCACCAGGCTGCGGCGTCGGCCGCGTCAGCCGCGTCGGCCGCCTCCGTCGACGCCATGGGCGGCACGTGCGCGCAGCCGTCCGCCCAGCCGCTGCCCGACATCCCGCGCTACCCCTGGATGTCCATCACCGATTGGATGAGCCCCTTCGACCGCGTGGTCTGCGGACCGAACGGCTGTCCAAGGCGACGGGGAAGGCAAACTTACACGAGGTTTCAAACGCTAGAATTAGAAAAGGAATTCCATTTTAACCATTATCTGACGCGCCGACGCAGGATAGAGATCGCGCACGCCCTGTGTCTCACAGAAAGACAAATCAAAATATGGTTCCAGAATCGACGAATGAAACTAAAGAAGGAGCTGCGCGCAGTGAAAGAGATAAACGAGCAAGCGCGCCGGGAGCGAGAGGAACAGGACCGGATGAAGCAGCAGCAGCAGGAGAAGCAGGCCAAGCTGGAGAGCCAGCACCACGGCCACCACGTGACCCACCACCACGACCCAATGAAAATGCCCATCGACAAGGGCTCCAGCGATATCCTCAAAGTGAATAAAGTGCCGACGTAA

B3_P1(abA)_1	GTCCCTGCCTCTATATCTTTCCCAGCCCGCCAGACAGCGACGACG

B3_P1(abA)_2	GTCCCTGCCTCTATATCTTTCGCCGACGACGACGACGACGAGCGC

B3_P1(abA)_3	GTCCCTGCCTCTATATCTTTACGGCACCGACCCCCCGAACTGCTG

B3_P1(abA)_4	GTCCCTGCCTCTATATCTTTATGGGGTCGCCGCTCACGGCCCCAG

B3_P1(abA)_5	GTCCCTGCCTCTATATCTTTCGCCGCCATGCTGGCCGACGCGGCC

B3_P1(abA)_6	GTCCCTGCCTCTATATCTTTTGGCGTCGACGGAGGCGGCCGACGC

B3_P2(abA)_1	GGCGAGCCGGCCCCCAGCGCGGCCGTTCCACTCAACTTTAACCCG

B3_P2(abA)_2	ATACATCCTCGCCGGCGCGCCCGGCTTCCACTCAACTTTAACCCG

B3_P2(abA)_3	CGTCCGCCAGCCCGCCGCCCGCGGGTTCCACTCAACTTTAACCCG

B3_P2(abA)_4	TTGTGCTGCCCCAGCGCGTAGTTCATTCCACTCAACTTTAACCCG

B3_P2(abA)_5	CGCAGCCTGGTGGTAGAACTGCGCCTTCCACTCAACTTTAACCCG

B3_P2(abA)_6	CGGACGGCTGCGCGCACGTGCCGCCTTCCACTCAACTTTAACCCG

***> abdominal-B (abB)***

lcl|XM_052883843.1_cds_XP_052739803.1_1 [gene=LOC112057786] [db_xref=GeneID:112057786] [protein=homeobox protein abdominal-B isoform X2] [protein_id=XP_052739803.1] [location=336..1283] [gbkey=CDS]

ATGATGAACGGCGTGGGCGGCGCGCTGTACGAGGAGGCGCACGCGTCGCCCCCGGGGGGCTCCCCGCCGGCCGCGCCCGCCGCCGCGCCGCCGTCCGCGTCCAGCGCCTCGCCGGCGTCCGTGGGCTCCAACCCGCCGCAGCCGCTGCACATCCCGGCCAAGCGCTACGAGCCCGAGCCCGGCGTCATCCGCCACGCGCAGCAGTCGTGGGGCTACCCGCCCGACGCGCCCGCACCCTTCGAGCACCAGTACCCGGCCGGGCCCACGTACTACAACCTGCCGGTGGAGCGCGAGCGCAAGGCGGGCCTGCCCTTCTGGCCCGGCGGCGGCGAGTACAAGCCCTACGCGGACGGCTGCCACCAAGGCTTCACGCAGCCGTGCTGGAACTACCCGTACGGGGCGCCGCGCGGCGACCAGCCGCTGCCGTACGTGGGCGAGGAGCGGCGCACCGCCGTGTCCGAGGCGTCCGGCTTCTCGCACGACGCGTACGGGCTGCGGAACTACGCGCCGGAGTCCGTGTCCAGCGCGCCCTACCCGCCGCCCAGCGCGCTGCCTGGATCCCTGTCCATGTCGGTCGGCGTCGGCGTCGGCTGCGGCTCCAACCCCCTAGACTGGACCGGACAAGTCACTGTGCGTAAAAAACGCAAACCCTACTCAAAGTTCCAGACCCTCGAATTAGAAAAAGAGTTCCTTTTCAACGCTTACGTGTCAAAGCAGAAGAGATGGGAACTGGCGCGGAACCTAAACCTCACAGAGAGACAGGTGAAGATATGGTTCCAGAACCGACGGATGAAGAACAAGAAGAACTCGCAGCGGCAGGCGGCGCAGGCGGCTCAGAACAACAACAACAACTCGAACGCCAATAACCACAACCACCACGGCAGCCATCACCATGCGCCGCACCACGTGGCGTTGCACCATGCGCCGCCTGCTAAGCATCATCAGTGA

B4_P1(abB)_1	CCTCAACCTACCTCCAACAATCGTAGCGCTTGGCCGGGATGTGCA

B4_P1(abB)_2	CCTCAACCTACCTCCAACAAGAAGGGTGCGGGCGCGTCGGGCGGG

B4_P1(abB)_3	CCTCAACCTACCTCCAACAAGCAGGCCCGCCTTGCGCTCGCGCTC

B4_P1(abB)_4	CCTCAACCTACCTCCAACAACACGGCTGCGTGAAGCCTTGGTGGC

B4_P1(abB)_5	CCTCAACCTACCTCCAACAAGGTGCGCCGCTCCTCGCCCACGTAC

B4_P1(abB)_6	CCTCAACCTACCTCCAACAAACACGGACTCCGGCGCGTAGTTCCG

B4_P2(abB)_1	GCGTGGCGGATGACGCCGGGCTCGGATTCTCACCATATTCGCTTC

B4_P2(abB)_2	CGTGGGCCCGGCCGGGTACTGGTGCATTCTCACCATATTCGCTTC

B4_P2(abB)_3	TGTACTCGCCGCCGCCGGGCCAGAAATTCTCACCATATTCGCTTC

B4_P2(abB)_4	CGCGGCGCCCCGTACGGGTAGTTCCATTCTCACCATATTCGCTTC

B4_P2(abB)_5	CGAGAAGCCGGACGCCTCGGACACGATTCTCACCATATTCGCTTC

B4_P2(abB)_6	CGCTGGGCGGCGGGTAGGGCGCGCTATTCTCACCATATTCGCTTC

***> BarH1***

lcl|XM_052889016.1_cds_XP_052744976.1_1 [gene=LOC112053314] [db_xref=GeneID:112053314] [protein=homeobox protein BarH-like 1b isoform X2] [protein_id=XP_052744976.1] [location=71..841] [gbkey=CDS]

ATGGAGGAGGAGTCGATGTCCGGGCACAGCTGTTCCGAGGACGACATCAGCGTCGGGCGTCCTTCGCCGG

CACCCTCGCGATCGCCTTCTCCTCAGGACTATTTCAGGCCGTTGAAACGGTTGAAGATGGCTTCCGAGCC

CGTGGAAGAGAGGAGAGGAGAAGGCGTCAAGAGCTTTTCCATACTGGACATTCTATCACACAGCCCCCGG

ACTCCTCCGCGGATCGTAAGACCCTGGGATGCGTCAGGGTTCGAGCGTTTGCAGCGATTGCAGCGCTTAG

CCGGTGCTGCGTTATTGGACGGCGAGCGCTGGAGACTGGCGGCTCTAGGATTGTTGAGGCCGAGGGAGAT

GCCAGCGGAGTGCTGTGATTCAGCGTCTGAAAGGTCTTCGTCCGCCTCGGATTGCTGCTCGGAGCCGCGG

CGGGTGTCGCAGGGAGGATCTACGGCGCCGCACACACCGCTAGATGCTCTGTTCCACATGACCAGCAAAA

CCTTCGAGGCGAATAATGCTGACAGTGGAGATGGTCAAAATCACTTGAACCTCTTCAACTCGCGGCCACA

GGCGAAGAAGAAACGCAAGTCCCGGACGGCGTTCACCAACCACCAGATCTTCGAGCTGGAGAAACGGTTC

CTCTACCAGAAGTACCTCTCCCCGGCTGACAGAGACGAGATCGCCAGCTCATTGGGACTTTCCAATGCGC

AAGTCATCACCTGGTTCCAAAACAGACGAGCAAAATCTAAGCGAGACGTTGAAGAATTTTGTATAACTTG

A

B2_P1 (barh1)_1	CCTCGTAAATCCTCATCAAATTGACGCCTTCTCCTCTCCTCTCTT

B2_P1 (barh1)_2	CCTCGTAAATCCTCATCAAAATCCCAGGGTCTTACGATCCGCGGA

B2_P1 (barh1)_3	CCTCGTAAATCCTCATCAAAAGCGCTCGCCGTCCAATAACGCAGC

B2_P1 (barh1)_4	CCTCGTAAATCCTCATCAAATCAGACGCTGAATCACAGCACTCCG

B2_P1 (barh1)_5	CCTCGTAAATCCTCATCAAACGGCGCCGTAGATCCTCCCTGCGAC

B2_P1 (barh1)_6	CCTCGTAAATCCTCATCAAACTCCACTGTCAGCATTATTCGCCTC

B2_P2 (barh1)_1	GATAGAATGTCCAGTATGGAAAAGCAAATCATCCAGTAAACCGCC

B2_P2 (barh1)_2	TCGCTGCAAACGCTCGAACCCTGACAAATCATCCAGTAAACCGCC

B2_P2 (barh1)_3	TCAACAATCCTAGAGCCGCCAGTCTAAATCATCCAGTAAACCGCC

B2_P2 (barh1)_4	CAGCAATCCGAGGCGGACGAAGACCAAATCATCCAGTAAACCGCC

B2_P2 (barh1)_5	GTGGAACAGAGCATCTAGCGGTGTGAAATCATCCAGTAAACCGCC

B2_P2 (barh1)_6	TGAAGAGGTTCAAGTGATTTTGACCAAATCATCCAGTAAACCGCC

***Danio rerio* specific genes**

***>glutamate decarboxylase 2 (gad2)_***lcl|NM_001017708.2_cds_NP_001017708.2_1 [gene=gad2] [db_xref=GeneID:550403,ZFIN:ZDB-GENE-030909-9] [protein=glutamate decarboxylase 2] [protein_id=NP_001017708.2] [location=315..2066] [gbkey=CDS]

ATGGCATCACACGGGTTTTGGTTTCTGGGGGCTGAAAACGCGGCTGGAAACGGCAGTCAAAGCCCGAACACGCCTAGAGCATGGTGCCAGGCGGCGGCCCAGAAATTCAGCGGAGGCATCGGCTCCAAATTATGTGCTCTGTTAAATGTCGGGGAGGCTGAGAAAGCAGCTCAAGCCCCAGTTAAAGCCGAGGATGAGTCTACAGCCGAGAGCTGCGGCTGTAATAAACCCTGTAACTGCTCCAAAGCCACCGCGTGCTTTTCGGATCTCTACTCAACAGATCTGTTACCTGCACTGGACGGAGACGCGAAGACTATGAATTTTCTGCAGGAGGTTGTGGATATATTGCTGGCTTATATAGTGGAATCATTTGACAGGTCTACGAAAGTGATTGATTTCCACTATCCAAACGAACTGCTCCAAAGGAATAATTGGGAGCTTTCGGACGAACCCGAGACTTTAGACGATATTCTGATCAGCTGTCGCGCTACGCTTAAATATGCCATAAAAACTGCGCATCCCAGGTATTTCAATCAACTCTCCACTGGGTTAGACATGGTTGGCTTGGCTGCAGATTGGCTAACCTCCACTGCCAACACCAATATGTTCACCTATGAGGTGGCTCCAGTCTTCGTGCTGTTGGAATACGTCACGCTGAAGAAGATGAGGGAGATCATTGGCTGGCAGGACGGCCACGGTGATGGAATATTCTCTCCGGGTGGCGCCATTTCCAACATGTATGCCATGCTACTGGCTCGCTATAAAATGTTCCCTGAGGTGAAGGAGAAAGGAATGTCATCCGTACCTAGGCTGGTGGCCTTCACATCAGAACATAGCCATTTTTCAATCAAGAAAGGAGCAGCTGCTCTTGGAATCGGTACAGAGAGTGTCATTTGCATTAAAGCTGATGAGAGGGGTAAGATGATTCCATCTGACCTTGAGAGGAGGATCATTGAAGCCAAGCAGAAGGGATACGTGCCGTTCTTTGTCAGCGCGACGGCCGGCACCACGGTTTATGGTGCCTTTGATCCTTTGATCGCTATAGCGGACATCTGCAAGAAGCATGACGTCTGGATGCATGTGGATGGAGCATGGGGTGGAAGTCTGCTAATGTCCCGGAAACACCGGTGGAAGCTTAATGGAGTTGAGAGGGCTAATTCTATGACCTGGAACCCTCATAAAATGATGGCTGTGCCCTTGCAGTGCTCTGCTCTTCTGGTTCGAGAGGAGGGACTGATGCAGAGCTGCAATCAGATGCAGGCCTGTTATCTGTTCCAGCAGGACAAGCATTATGACCTGCAGTATGACACAGGAGACAAGGCCCTGCAGTGTGGACGTCATGTGGACATCTTTAAACTGTGGCTGATGTGGAGAGCTAAGGGCACGATTGGTTTTGAGGCTCAGATCGACAAATGCCTGGAGCTTTCCGAATATCTCTACAACAAGATTAAGGACAGGGAAGGATATCAGATGGTGTTTGATGGAAAGCCGCAGCATACCAATGTGTGTTTCTGGTACCTTCCACCGGGCGTGCGCTACCTGGAGGACAAAGTGGAGAGGATGAAGCGTCTGCACAAGGTTGCCCCTGTAATCAAAGCCAGAATGATGGAGTACGGCACGACCATGGTGAGCTACCAGCCACAGGGAGACAAGGTCAACTTCTTCCGCATGGTCATCTCCAATCCAGCCGCTACCTTTGAAGACATTGACTTCCTCATTGAAGAGATCGAGCGACTGGGGCAGGATCTTTAA

B1_P1 (gad2)_1	GAGGAGGGCAGCAAACGGAAGCTGCTTTCTCAGCCTCCCCGACAT

B1_P1 (gad2)_2	GAGGAGGGCAGCAAACGGAAGCAGTTACAGGGTTTATTACAGCCG

B1_P1 (gad2)_3	GAGGAGGGCAGCAAACGGAATCGCGTCTCCGTCCAGTGCAGGTAA

B1_P1 (gad2)_4	GAGGAGGGCAGCAAACGGAAGACCTGTCAAATGATTCCACTATAT

B1_P1 (gad2)_5	GAGGAGGGCAGCAAACGGAATTCGTCCGAAAGCTCCCAATTATTC

B1_P1 (gad2)_6	GAGGAGGGCAGCAAACGGAAGATGCGCAGTTTTTATGGCATATTT

B1_P1(gad2)_7	GAGGAGGGCAGCAAACGGAAGTGGAGGTTAGCCAATCTGCAGCCA

B1_P1 (gad2)_8	GAGGAGGGCAGCAAACGGAACTTCAGCGTGACGTATTCCAACAGC

B1_P2 (gad2)_1	TCATCCTCGGCTTTAACTGGGGCTTTAGAAGAGTCTTCCTTTACG

B1_P2 (gad2)_2	ATCCGAAAAGCACGCGGTGGCTTTGTAGAAGAGTCTTCCTTTACG

B1_P2 (gad2)_3	CAACCTCCTGCAGAAAATTCATAGTTAGAAGAGTCTTCCTTTACG

B1_P2 (gad2)_4	GGATAGTGGAAATCAATCACTTTCGTAGAAGAGTCTTCCTTTACG

B1_P2 (gad2)_5	GATCAGAATATCGTCTAAAGTCTCGTAGAAGAGTCTTCCTTTACG

B1_P2 (gad2)_6	CAGTGGAGAGTTGATTGAAATACCTTAGAAGAGTCTTCCTTTACG

B1_P2 (gad2)_7	TCATAGGTGAACATATTGGTGTTGGTAGAAGAGTCTTCCTTTACG

B1_P2 (gad2)_8	CTGCCAGCCAATGATCTCCCTCATCTAGAAGAGTCTTCCTTTACG

>***NeuroD***_lcl|AF036148.1_cds_AAB88820.1_1 [gene=nrd] [protein=NeuroD] [protein_id=AAB88820.1] [location=57..1109] [gbkey=CDS]

ATGACGAAGTCATACAGCGAGGAAAGCATGATGCTGGAGTCTCAGAGCAGCTCGAACTGGACCGACAAGTGTCACAGCAGCTCCCAAGACGAACGAGACGTGGACAAGACCAGCGAGCCCATGCTCAACGATATGGAAGACGACGATGATGCCGGTCTCAACCGACTCGAGGATGAGGACGACGAGGAAGAAGAAGAGGAGGAAGAAGACGGCGACGACACCAAGCCGAAAAGACGGGGTCCCAAGAAGAAGAAGATGACCAAGGCGCGTATGCAGAGATTCAAGATGCGGCGCATGAAGGCGAACGCCCGGGAGAGGAACCGCATGCACGGCCTCAACGACGCGCTTGAGAGCCTGCGCAAAGTTGTTCCGTGCTACTCCAAAACGCAGAAGCTCTCCAAGATCGAGACGCTCCGACTAGCCAAGAACTACATTTGGGCTCTTTCGGAAATCTTGAGGTCGGGCAAAAGCCCCGACCTGATGTCTTTTGTGCAGGCCTTGTGCAAGGGCTTGTCCCAACCCACGACCAACTTGGTCGCAGGATGCCTCCAACTGAACCCCAGAACTTTTCTGCCCGAGCAGAGCCAGGAGATGCCCCCTCATATGCAAACAGCAAGTGCTTCCTTTTCCGCTCTTCCCTACTCCTACCAGACGCCCGGTCTTCCCAGCCCTCCGTACGGTACAATGGACAGCTCTCACATCTTTCACGTCAAGCCGCACGCGTACGGGAGCGCACTGGAGCCGTTCTTTGACACCACCCTCACAGACTGCACCAGTCCCTCATTTGACGGACCCCTTAGCCCGCCTTTAAGCGTCAACGGGAACTTTTCGTTCAAACACGAGCCTTCTTCGGAATTCGAGAAGAACTACGCGTTTACCATGCACTATCAGGCAGCGGGTCTGGCCGGCGCGCAGGGACACGCCGCGTCTCTTTACGCGGGCTCGACGCAGCGCTGTGATATACCGATGGAGAACATTATGTCGTACGACGGGCACTCGCATCACGAACGGGTCATGAACGCCCAGTTGAACGCGATATTTCACGACTCGTGA

B2_P1 (NeuroD)_1	CCTCGTAAATCCTCATCAAATCGAGTCGGTTGAGACCGGCATCAT

B2_P1 (NeuroD)_2	CCTCGTAAATCCTCATCAAAACCCCGTCTTTTCGGCTTGGTGTCG

B2_P1 (NeuroD)_3	CCTCGTAAATCCTCATCAAAGGGCGTTCGCCTTCATGCGCCGCAT

B2_P1 (NeuroD)_4	CCTCGTAAATCCTCATCAAAGAGTAGCACGGAACAACTTTGCGCA

B2_P1 (NeuroD)_5	CCTCGTAAATCCTCATCAAATTCCGAAAGAGCCCAAATGTAGTTC

B2_P1 (NeuroD)_6	CCTCGTAAATCCTCATCAAAGTTGGGACAAGCCCTTGCACAAGGC

B2_P1 (NeuroD)_7	CCTCGTAAATCCTCATCAAATCCTGGCTCTGCTCGGGCAGAAAAG

B2_P1 (NeuroD)_8	CCTCGTAAATCCTCATCAAAACCGGGCGTCTGGTAGGAGTAGGGA

B2_P2 (NeuroD)_1	TCTTCTTCTTCCTCGTCGTCCTCATAAATCATCCAGTAAACCGCC

B2_P2 (NeuroD)_2	CGCCTTGGTCATCTTCTTCTTCTTGAAATCATCCAGTAAACCGCC

B2_P2 (NeuroD)_3	TGAGGCCGTGCATGCGGTTCCTCTCAAATCATCCAGTAAACCGCC

B2_P2 (NeuroD)_4	TCGATCTTGGAGAGCTTCTGCGTTTAAATCATCCAGTAAACCGCC

B2_P2 (NeuroD)_5	GTCGGGGCTTTTGCCCGACCTCAAGAAATCATCCAGTAAACCGCC

B2_P2 (NeuroD)_6	GGCATCCTGCGACCAAGTTGGTCGTAAATCATCCAGTAAACCGCC

B2_P2 (NeuroD)_7	CTTGCTGTTTGCATATGAGGGGGCAAAATCATCCAGTAAACCGCC

B2_P2 (NeuroD)_8	CATTGTACCGTACGGAGGGCTGGGAAAATCATCCAGTAAACCGCC

>***nrp1a***_lcl|XM_009297199.3_cds_XP_009295474.1_1 [gene=nrp1a] [db_xref=GeneID:353246] [protein=neuropilin-1a isoform X2] [protein_id=XP_009295474.1] [location=612..3371] [gbkey=CDS]

ATGCATTGTGGATTAGTGTTGATCCTCTTTACGGGAATCTTTCTCATCGTCAGTGCTCTCAAAAATGACAAATGTGGGGACAATATCAGGATCACTAGTGCCAATTATCTCACATCGCCCGGCTATCCAGTATCTTACTACCCGTCTCAGAAATGCATATGGGTGATTACAGCTCCAGGACCCAACCAGAGGATTTTGATCAACTTCAACCCACACTTTGACCTGGAAGACCGAGAGTGCAAATATGACTATGTGGAAGTGAGAGACGGCGTGGATGAGAACGGGCAGCTGGTGGGCAAATATTGTGGAAAGATCGCTCCATCTCCGGTGGTCTCGTCGGGAAACCAGCTCTTCATCAAGTTTGTGTCCGACTACGAGACTCACGGTGCCGGATTCTCCATCCGCTATGAGATCTTCAAAACGGGTCCAGAATGTTCCAGGAACTTCACCTCCAGCAGCGGAGTCATCAAGTCGCCCGGATTTCCAGAGAAATACCCCAATAATTTGGACTGCACGTTCATGATCTTTGCTCCTAAGATGTCAGAAATCGTTTTAGAGTTCGAGAGTTTTGAGCTGGAGCCAGACACGCAGCCGCCCGCCGGAGTCTTCTGCCGATACGACCGCCTGGAGATTTGGGACGGATTCCCTGGAGTTGGTCCATACATCGGCAGATACTGCGGACAGAATACTCCAGGACGGATCATATCCTACACTGGAACTTTGGCCATGACAATCAACACAGACAGCGCTATTGCTAAAGAAGGATTTTCAGCTAATTTCACTGTGCTGGAAAGGACGGTTCCTGATGACTTTGACTGCACTGAGCCTTTGGGTATGGAAACTGGGGAAATCCATTCGGACCAAATCATGGCCTCGTCCCAGTACAGCAACAGCTGGTCAGCAGAACGATCACGACTCAACAACCCAGAGAACGGCTGGACGCCTTTGGAGGACACGAATAAAGAGTGGATTCAGGTTGATCTTGGATTTCTGCGCTTTGTCTCGGCCATCGGCACACAGGGAGCCATTTCTCAGGAGACCAAGAAGAAATATTACGTGAAGGAATATAAGGTGGACGTGAGCTCCAATGGTGAAGACTGGATCACTATTAAGGATGGCCCAAAACAGAAGCTTTTCCAGGGTAACACCAACCCTACTGATGTGGTGAAAGCCAAGTTTCCAAAGCCCACGCTGACCCGATACCTGCGGATCCGACCCATTAACTGGGAAACTGGCATCGCACTGCGGTTCGAGGTCTACGGATGCAAGATCTCAGAGTATCCGTGCTCAGGAATGCTGGGAATGGTTTCAGGCCTGATCACAGACTCTCAGATCACCGTCTCGTCCCACATCGAGCGTACCTGGGTTTCAGAAAACGCTCGTCTGATGACCAGCCGTTCAGGATGGATGTTACTGCCACAGTCACAACCGTATGCAGACGAATGGCTGCAGATAGATCTGGCCGAAGAAAAACTAGTCAAAGGTTTGATCATCCAGGGAGGCAAACACCGAGATAACAAGGTGTTCATGAAGAAGTTTCGTCTGGGATACAGCAACAACGGATCCGATTGGAAATTGGTGATGGATGCTACCGGCAACAAGCCCAAGATTTTTGAGGGGAATTTGAACTATGACACCCCAGCGTTGAGGACAATGGAGCCAGTGTTGACACGCTTTGTTAGGATTTACCCAGACAGAGGCACTCCTGCTGGGATGGGACTCAGACTGGAACTTTTGGGATGTGAAATGGAAGTGCCTACAGTGCCGCCTACTACACCCGCAGCCTCCACCACACCGTCAGACGAGTGTGACGATGACCAGGCCAACTGCCATAGTGGAACAGGTGGAACCACTGCGACAGAGACGATCAGGGAGATGAGCACCATTCCAGGTGAGAAAGCATTCCTGTGGTTTGCTTGTGACTTCGGCTGGGCAAATGATCCTTCGTTCTGCGGCTGGATCTCCGAGGACAGCGGTTTCAGGTGGCAGATTCAGTCCAGTGGCACCCCGACTCTAAACACCGGCCCAAACATGGACCACACAGGCGGATCAGGGAACTTCATCTACACGCTGGCGACCGGAGCTCAGGAGACAGAAGTGGCCCGGTTAGTGAGTCCGTCGGTGTCGGGCCAGGACTCAGACCTCTGCCTGTCTTTCTGGTACCACATGTTTGGCTCTCACATCGGCACGCTGCACATTAAACAGCGCAGAGAGACCAGTCAGGGCTCGGCTGATGTGCTGCTGTGGACCGTCAGCGGACATCAGGGCAACCGCTGGAGGGAGGGACGCGTCCTCATACCACACTCCAACAAACCCTACCAGGTAATAATTGAAAGTGTGGTCGAGAGGAAGAGTTGGGGAGACATCGCTGTAGACGACATCAAAATCTTAGACAATGTGAACATGGCCGACTGCAAGGACCCAGATGTCCCAGCAGAGCCGATACAGCCAGAAGACAATTTCAATGAAATCATGGTGGACATCACCGATTTCCCAGATATCGTGGAGAACCCAGATATCGGCGGAGCTGGAAACATGCTGAAAACCCTGGACCCCATCCTCATTACTATCATCGCCATGAGTGCGCTTGGTGTTTTCCTGGGCGCCATCTGTGGCGTGGTGCTGTACTGCGCCTGCTCTCATAGCGGCATGTCAGACAGGAACTTATCAGCCCTGGAGAACTATAATTTTGAGCTGGTGGACGGCGTCAAGTTAAAGAAGGACAAGCTCAACTCACAGAACTCGTACTCGGAAGCGTGA

B3_P1(nrp1a)_1	GTCCCTGCCTCTATATCTTTGTAATCACCCATATGCATTTCTGAG

B3_P1(nrp1a)_2	GTCCCTGCCTCTATATCTTTGCACTCTCGGTCTTCCAGGTCAAAG

B3_P1(nrp1a)_3	GTCCCTGCCTCTATATCTTTTTCCACAATATTTGCCCACCAGCTG

B3_P1(nrp1a)_4	GTCCCTGCCTCTATATCTTTGTCTCGTAGTCGGACACAAACTTGA

B3_P1(nrp1a)_5	GTCCCTGCCTCTATATCTTTGGTGAAGTTCCTGGAACATTCTGGA

B3_P1(nrp1a)_6	GTCCCTGCCTCTATATCTTTTGAACGTGCAGTCCAAATTATTGGG

B3_P1(nrp1a)_7	GTCCCTGCCTCTATATCTTTTGCGTGTCTGGCTCCAGCTCAAAAC

B3_P1(nrp1a)_8	GTCCCTGCCTCTATATCTTTTGGACCAACTCCAGGGAATCCGTCC

B3_P2(nrp1a)_1	AAAATCCTCTGGTTGGGTCCTGGAGTTCCACTCAACTTTAACCCG

B3_P2(nrp1a)_2	GTCTCTCACTTCCACATAGTCATATTTCCACTCAACTTTAACCCG

B3_P2(nrp1a)_3	ACGAGACCACCGGAGATGGAGCGATTTCCACTCAACTTTAACCCG

B3_P2(nrp1a)_4	TAGCGGATGGAGAATCCGGCACCGTTTCCACTCAACTTTAACCCG

B3_P2(nrp1a)_5	GGGCGACTTGATGACTCCGCTGCTGTTCCACTCAACTTTAACCCG

B3_P2(nrp1a)_6	TTTCTGACATCTTAGGAGCAAAGATTTCCACTCAACTTTAACCCG

B3_P2(nrp1a)_7	TATCGGCAGAAGACTCCGGCGGGCGTTCCACTCAACTTTAACCCG

B3_P2(nrp1a)_8	ATTCTGTCCGCAGTATCTGCCGATGTTCCACTCAACTTTAACCCG

>***omp***_lcl|AF457189.1_cds_AAL87664.1_1 [gene=omp] [protein=olfactory marker protein] [protein_id=] [location=71..538] [gbkey=CDS]

ATGTCTCTGGAGTTGACGTTCAATCCTGATGTCCAGCTGACGGAGATGATGCGTCTGCGCGTTCAGTCTCTACAACAACGAGGACAGAAACGTCAAGACGGAGAGCGTCTCCTCAAGTCCAACGAGCACGTCTACAGTCTGGACTTCTCCGAACAGGCCTTGCATTTCACCCGCTGGAACATTCGCATTTCCAGCCCGGGACGCCTAAACATCATCGCCACTTCCCAGCTCTGGACGCCCGACCTCACACACCTGATGACCCGGCAGCTCCTGGAACCCACCGGACTCTTCTGGAGGAGCGCAGACGACGAGAACATCCAGTGTTATGAGGCCGACGCACAGGAGTTTGGTGAAAGGATAGCAGAGCTGGCCAAAGTGCGAAAGGTGATGTATTTCCTGTTCGCCTTTGAAGACGGCTTGAGTCCGGAGAGCGTGGAATGCTCCATTGAATTCCAGACCTCCAAGTGA

B3_P1(omp)_1	GTCCCTGCCTCTATATCTTTATCAGGATTGAACGTCAACTCCAGA

B3_P1(omp)_2	GTCCCTGCCTCTATATCTTTCGTCTTGACGTTTCTGTCCTCGTTG

B3_P1(omp)_3	GTCCCTGCCTCTATATCTTTGTGAAATGCAAGGCCTGTTCGGAGA

B3_P1(omp)_4	GTCCCTGCCTCTATATCTTTGGGCGTCCAGAGCTGGGAAGTGGCG

B3_P1(omp)_5	GTCCCTGCCTCTATATCTTTCGTCGTCTGCGCTCCTCCAGAAGAG

B3_P1(omp)_6	GTCCCTGCCTCTATATCTTTCGCACTTTGGCCAGCTCTGCTATCC

B3_P2(omp)_1	CAGACGCATCATCTCCGTCAGCTGGTTCCACTCAACTTTAACCCG

B3_P2(omp)_2	GCTCGTTGGACTTGAGGAGACGCTCTTCCACTCAACTTTAACCCG

B3_P2(omp)_3	GGGCTGGAAATGCGAATGTTCCAGCTTCCACTCAACTTTAACCCG

B3_P2(omp)_4	CTGCCGGGTCATCAGGTGTGTGAGGTTCCACTCAACTTTAACCCG

B3_P2(omp)_5	CGTCGGCCTCATAACACTGGATGTTTTCCACTCAACTTTAACCCG

B3_P2(omp)_6	AAGGCGAACAGGAAATACATCACCTTTCCACTCAACTTTAACCCG

>***oxt***_lcl|NM_178291.2_cds_NP_840076.1_1 [gene=oxt] [db_xref=GeneID:352920,ZFIN:ZDB-GENE-030407-1] [protein=oxytocin-neurophysin 1 precursor] [protein_id=NP_840076.1] [location=64..528] [gbkey=CDS]

ATGTCTGGAGGTCTGCTGTCCGCGGCGGCTCTGCTGTGTCTGCTGTCCGTCTGCTCGGCCTGCTACATCTCAAACTGCCCCATCGGAGGAAAACGCTCCGTTCAGGACTGGCCCATTCGACAGTGTATGCCGTGTGGCCCCGGGGACCGCGGACGCTGTTTCGGCCCCAGTATCTGCTGTGGTGAAGGCATCGGCTGCTTGGTCGGCTCTCCAGAAACCCTGCGCTGTCTGGAGGAGGATTTTCTCCCTTCTCCGTGTGAGATGTCTGGAAAGGCCTGCGGTTATGAGGGACGCTGCGCTGCTCCTGGAGTCTGCTGCGACTCGGAGGGCTGCAGTGTGGACCAGTCGTGTGTGGACGGAGATGCTGACGCTGCAGCTGTCAATCAACCGGCCAACAGCCCAGATCTGCTGCTGAAGCTCCTGCACCTGTCAAGCCACACCCACCCCTCCAGAATCCACCAATGA

B1_P1 (oxt)_1	GAGGAGGGCAGCAAACGGAAAGCCGCCGCGGACAGCAGACCTCCA

B1_P1 (oxt)_2	GAGGAGGGCAGCAAACGGAACGGAGCGTTTTCCTCCGATGGGGCA

B1_P1 (oxt)_3	GAGGAGGGCAGCAAACGGAACTGGGGCCGAAACAGCGTCCGCGGT

B1_P1 (oxt)_4	GAGGAGGGCAGCAAACGGAAATCCTCCTCCAGACAGCGCAGGGTT

B1_P1 (oxt)_5	GAGGAGGGCAGCAAACGGAACTCCAGGAGCAGCGCAGCGTCCCTC

B1_P1 (oxt)_6	GAGGAGGGCAGCAAACGGAAACAGCTGCAGCGTCAGCATCTCCGT

B1_P2 (oxt)_1	CGAGCAGACGGACAGCAGACACAGCTAGAAGAGTCTTCCTTTACG

B1_P2 (oxt)_2	TACACTGTCGAATGGGCCAGTCCTGTAGAAGAGTCTTCCTTTACG

B1_P2 (oxt)_3	CAGCCGATGCCTTCACCACAGCAGATAGAAGAGTCTTCCTTTACG

B1_P2 (oxt)_4	AGACATCTCACACGGAGAAGGGAGATAGAAGAGTCTTCCTTTACG

B1_P2 (oxt)_5	CACTGCAGCCCTCCGAGTCGCAGCATAGAAGAGTCTTCCTTTACG

B1_P2 (oxt)_6	AGATCTGGGCTGTTGGCCGGTTGATTAGAAGAGTCTTCCTTTACG

>***tph2***_lcl|NM_001310068.1_cds_NP_001296997.1_1 [gene=tph2] [db_xref=GeneID:407712,ZFIN:ZDB-GENE-040624-4] [protein=tryptophan 5-hydroxylase 2 isoform 1] [protein_id=NP_001296997.1] [location=89..1729] [gbkey=CDS]

ATGCAAGGGAAGAAAACCTCTCAAGAGACAACAGCAACTATGTCTAATGAACTGCAAACTAATCCAAAGG

GGCAGCAAACAGCCCCTAAATCACCTGTTGGGCTTCTTAGGAAAAGAGGTCAAGAAGCAATTCCGAAGAT

GCAACCTGCCATGATGATGTTCTCCAGCAAATACTGGGCTCGGAGAGGACTATCTTTGGATTCAGCTATG

TATGACCAACAGCACCTTGCATCCTCCATGCTCCGCAGGACATCCTTTAATCGGATAGATGAAAGACCTG

ACAAAGAAGAACAGAAATCCACTCATGACCTTGGGAAGCTGGCAGTGATTTTCTCTCTGAAGAATGAAGT

TGGCTTTCTGGTGAAAGCGCTGAGGCTCTTTCAGGAGAAGCATGTGAATTTGGCGCACATTGAGTCTCGG

AGATCAAAGAGACTCACCAATGAGATTGAGATATATGCAGAATGCAACTGCACAAAGAAAGAGTTCAACG

AACTGGTGCAACACCTCAAAGACCACGTTAATATTGTCTCGTATAACACACCTCAACATGTGTGGTCTGC

AGAGACCGACTGTTTGGACTGTGTGTGTGTTTTGGGTGGATTGCCAGATGGGGAAGGAATACCATGGTTT

CCCCAAAAAATCTCAGAGCTGGATCAGTGCTCCCATAGAGTGCTAATGTATGGCTCTGAACTGGATGCTG

ATCATCCAGGCTTTAAAGACAAGGTCTATCGGCAGAGGAGAAAGTACTTTGTGGAGGTTGCAATGAACTA

CAAATTTGGGCAGCCCATCCCACGGATTGAGTACACTGCAGAGGAAGTAAAGACATGGGGAGTCGTTTAC

AGAGAACTTACAAAACTCTATCCAACTCATGCCTGCCGCGAATACTTAAAAAACCTTCCTCTGCTGACAA

AGCATTGTGGGTACAGAGAGGACAACATCCCACAGCTGGAGGACGTGTCTCTATTCTTAAGAGAGAGGTC

AGGGTTTACCGTAAGGCCTGTGGCTGGATATCTCTCACCACGAGACTTCCTGGCTGGACTGGCTTATCGA

GTGTTTAATTGCACTCAGTATATTCGTCACAGCACAGACCCTCTCTATACACCAGAGCCGGACACCTGCC

ATGAACTGCTTGGTCACGTGCCGCTCCTGGCCGATCCCAAATTTGCTCAGTTTTCTCAGGAGATTGGCCT

TGCATCTCTAGGCGCTTCAGACGAAGATGTGCAAAAGCTGGCAACTTGTTATTTCTTCACAATAGAGTTT

GGGCTGTGCAAACAAGATGGTCAGTTGAGAGTCTATGGAGCAGGTCTACTGTCTTCTATTGGAGAGTTAA

GGCATGCGCTTTCTGATAAAGCAACAGTGAAGGTGTTTGACCCCAAAACCACGTGCTACCAGGAGTGCCT

CATTACCACATTTCAAGATGTGTATTTTGTCTCTGAGAGCTTTGAGGAGGCAAAAGAGAAAATGAGGGAA

TTTGCTAAATCAATAAAGAGGCCTTTTTCTGTCTACTACAACCCTTACACGCAGAGCATCGACTTACTCA

AGGACACCAGGAGCATTGAAAATGTGGTTCAAGATCTACGCAGCGACTTAACCACCGTCTGTGACGCCCT

GGGAAAAATGAACAAATACCTCGGTATCTAA

B1_P1 (tph2)_1	GAGGAGGGCAGCAAACGGAATTGCTGGAGAACATCATCATGGCAG

B1_P1 (tph2)_2	GAGGAGGGCAGCAAACGGAACATGGAGGATGCAAGGTGCTGTTGG

B1_P1 (tph2)_3	GAGGAGGGCAGCAAACGGAAGGTCATGAGTGGATTTCTGTTCTTC

B1_P1 (tph2)_4	GAGGAGGGCAGCAAACGGAAAAGAGCCTCAGCGCTTTCACCAGAA

B1_P1 (tph2)_5	GAGGAGGGCAGCAAACGGAACTCAATCTCATTGGTGAGTCTCTTT

B1_P1 (tph2)_6	GAGGAGGGCAGCAAACGGAATAACGTGGTCTTTGAGGTGTTGCAC

B1_P1 (tph2)_7	GAGGAGGGCAGCAAACGGAAACACACACACAGTCCAAACAGTCGG

B1_P1 (tph2)_8	GAGGAGGGCAGCAAACGGAAGCACTGATCCAGCTCTGAGATTTTT

B1_P2 (tph2)_1	AAAGATAGTCCTCTCCGAGCCCAGTTAGAAGAGTCTTCCTTTACG

B1_P2 (tph2)_2	TATCCGATTAAAGGATGTCCTGCGGTAGAAGAGTCTTCCTTTACG

B1_P2 (tph2)_3	GAGAGAAAATCACTGCCAGCTTCCCTAGAAGAGTCTTCCTTTACG

B1_P2 (tph2)_4	TGCGCCAAATTCACATGCTTCTCCTTAGAAGAGTCTTCCTTTACG

B1_P2 (tph2)_5	CTTTGTGCAGTTGCATTCTGCATATTAGAAGAGTCTTCCTTTACG

B1_P2 (tph2)_6	GTTGAGGTGTGTTATACGAGACAATTAGAAGAGTCTTCCTTTACG

B1_P2 (tph2)_7	CCTTCCCCATCTGGCAATCCACCCATAGAAGAGTCTTCCTTTACG

B1_P2 (tph2)_8	AGAGCCATACATTAGCACTCTATGGTAGAAGAGTCTTCCTTTACG

***>elavl3_***NM_131449.1 **elavl3** [organism=Danio rerio] [GeneID=30732]

TAGATCATATCATCTTTGTACGTCAAGAATGGTTACTATAATTAGCACCATGGAAACTCAGGTGTCCAAT

GGTCCGAGCGGAACCAGCCTGCCTAACGGCCCTGTCATTAGCACTAACGGCGCCACAGATGACAGCAAAA

CTAACCTGATCGTCAACTACCTGCCTCAGAACATGACCCAGGAAGAGTTCAAGAGCCTCTTTGGCAGCAT

CGGGGAAATCGAGTCCTGCAAATTGGTCAGAGACAAGATCACAGGCCAGAGCTTGGGATATGGCTTTGTA

AACTATGTGGATCCCAACGACGCCGACAAGGCTATCAACACGCTCAACGGTCTCAAACTGCAGACCAAAA

CAATCAAGGTGTCTTACGCCAGGCCCAGCTCAGCTTCCATCCGCGATGCCAACCTGTATGTGAGCGGCCT

GCCCAAAACCATGAGTCAGAAAGACATGGAGCAGTTGTTTTCCCAGTATGGAAGGATCATCACCTCACGC

ATCCTGGTAAACCAGGTCACAGGTATATCGCGCGGGGTAGGTTTCATTCGGTTCGACAAACGGAACGAAG

CAGAGGAGGCCATCAAGGGCCTGAACGGTCAGAAGCCACTAGGAGCAGCTGAGCCCATCACCGTAAAGTT

CGCCAACAACCCCAGTCAGAAGACAGGACAGGCTCTGCTGACCCAGCTCTACCAGACAGCCGCTCGCCGC

TACACTGGCCCTCTGCACCACCAGACCCAGCGCTTCAGATTCTCCCCCATAACCATTGACAGCATGACTA

GTCTTGCCGGGGTCAACCTGACCGGGCCCACTGGAGCCGGCTGGTGCATCTTCGTCTACAACCTGTCCCC

GGAAGCTGACGAAAGTGTCCTGTGGCAGCTCTTCGGGCCTTTTGGCGCCGTCACAAACGTCAAGGTCATC

CGTGACTTCACCACCAACAAATGTAAGGGCTTTGGCTTCGTCACCATGACCAACTACGACGAGGCAGCCA

TGGCTATCGCCAGTCTGAATGGCTACCGCCTGGGCGACCGCGTGCTGCAGGTCTCGTTCAAGACCAGCAA

GCAGCACAAGGCTTGAAGGAAGGCCTAGTCACTATTGCTCTTTAACATGCAGGGGGAGCTACTGAGCTC

B2_P1 (elavl3)_1	CCTCGTAAATCCTCATCAAATCCTCTGCTTCGTTCCGTTTGTCGA

B2_P1 (elavl3)_2	CCTCGTAAATCCTCATCAAAGTTGGCGAACTTTACGGTGATGGGC

B2_P1 (elavl3)_3	CCTCGTAAATCCTCATCAAACAGTGTAGCGGCGAGCGGCTGTCTG

B2_P1 (elavl3)_4	CCTCGTAAATCCTCATCAAAATGGTTATGGGGGAGAATCTGGGGA

B2_P1 (elavl3)_5	CCTCGTAAATCCTCATCAAAGACGAAGATGCACCAGCCGGCTCCA

B2_P1 (elavl3)_6	CCTCGTAAATCCTCATCAAATTGTGACGGCGCCAAAAGGCCCGAA

B2_P1 (elavl3)_7	CCTCGTAAATCCTCATCAAATAGTTGGTCATGGTGACGAAGCCAA

B2_P1 (elavl3)_8	CCTCGTAAATCCTCATCAAACGAGACCTGCAGCACGCGGTCGCCC

B2_P2 (elavl3)_1	TTCTGACCGTTCAGGCCCTTGATGGAAATCATCCAGTAAACCGCC

B2_P2 (elavl3)_2	AGCCTGTCCTGTCTTCTGACTGGGGAAATCATCCAGTAAACCGCC

B2_P2 (elavl3)_3	AGCGCTGGGTCTGGTGGTGCAGAGGAAATCATCCAGTAAACCGCC

B2_P2 (elavl3)_4	ACCCCGGCAAGACTAGTCATGCTGTAAATCATCCAGTAAACCGCC

B2_P2 (elavl3)_5	TTCGTCAGCTTCCGGGGACAGGTTGAAATCATCCAGTAAACCGCC

B2_P2 (elavl3)_6	TGGTGAAGTCACGGATGACCTTGACAAATCATCCAGTAAACCGCC

B2_P2 (elavl3)_7	CTGGCGATAGCCATGGCTGCCTCGTAAATCATCCAGTAAACCGCC

B2_P2 (elavl3)_8	AGCCTTGTGCTGCTTGCTGGTCTTGAAATCATCCAGTAAACCGCC

***>chrna3_***lcl|XM_001921279.5_cds_XP_001921314.1_1 [gene=chrna3] [db_xref=GeneID:568467] [protein=neuronal acetylcholine receptor subunit alpha-3] [protein_id=XP_001921314.1] [location=49..1575] [gbkey=CDS]

ATGAACACCGGAGCTCTCCTGATCTTCTTCTCCTCATTTTCTCCTCTGTGTTTTCTCCTGTCGGGGGTTT

GCTGCTCTGAGGCCGAGCACCGGCTCTTCTCTGTCATTTTCTCCAGCTATAACCAGTACATCCGACCAGT

GGAGAATGTGTCTGATCCAGTGGTCGTCCAGTTCGAGGTCTCCATGTCACAGCTGGTCAAAGTGGACGAG

GTGAACCAGATTATGGAGACCAATCTTTGGCTGAGACACATTTGGAATGACTATAAGCTCCGATGGGATC

CCAAGGATTTTGGAGGTGTTGAGTTCATCCGTGTGCCGTCCAACAAGATTTGGAAGCCAGACATTGTGTT

GTATAACAATGCGGTGGGAGATTTCCAGGTGGACGATAAAACCAAAGCCCTTCTGCGCTTCAATGGAGAC

GTGACCTGGATCCCACCGGCCATCTTCAAGAGCTCCTGCAAGATCGACGTCACCTACTTCCCCTTCGACT

ACCAGAACTGCACCATGAAGTTTGGCTCGTGGACCTACGACAAGGCCAAGATCGACCTGGTGCTCATCGG

CTCCACCATCAACCTGAAGGACTTCTGGGAGAGCGGAGAATGGACGATCATCGACGCTCCAGGATATAAA

CACGACATTAAATACAACTGCTGTGAGGAGATCTACACGGACATCACGTATTCACTGTACATCCGCCGAT

TACCGCTCTTCTACACCATCAACATGATCATCCCCTGTCTTCTCATCTCCTTCCTGACCGTTCTCGTATT

CTACCTGCCCTCCGACTGCGGAGAGAAAGTCACTTTATGCATTTCCGTCCTCCTGTCGCTCACTGTATTT

CTTCTGGTCATCACAGAAACCATCCCATCTACGTCTCTAGTCATTCCGCTAATAGGAGAGTATCTGCTTT

TCACCATGATATTCGTCACGCTCTCTATTGTGATTACGGTGTTTGTGTTAAATGTGCATTATCGTACGCC

TAAGACTCATACTATGCCCTGTTGGGTTCGTCGAGTGTTTCTGAGTCTGCTTCCTCGGGTTATGTTCATG

ACCCGGCCTGAGAAAGACCAGGAGGTTCCTGCCAAACAGTGCAGTGTTCATCCTCCGGTTTCCAGCAAAC

AGACAAGTGTGTGTCGCGGGCCGCAGCTCCTCCTGTGTCCGACTGAACTCAACGAGGCGTCCAGGACGGC

GTTTCTGTGTCGAGAGCACCACTGCTGGAAGAAACACATCTCCAACATGACCAGCGAGGGCGCAGAGGAC

GGAGGCAGCCCGTGCTCCAGCTCTGAGTCTCTGGACGGATTTCTGTGCATGTCTGCTGTTTCACCGCAGG

TCCGAGAGGCTATCGAGAGCGTCAAGTACATCGCTGAAAACATGAGGCTACAAAACGAGGCTAAAGAGGT

CCAGGATGACTGGAAGTATGTGGCGATGGTGATAGACAGGATCTTTCTGTGGGTGTTTGTGCTGGTGTGT

ATTCTGGGTACAGCGGGACTCTTCCTGCAGCCGCTCCTGCTGGGCGAAGACATGTGA

B2_P1 (chrna3)_1	CCTCGTAAATCCTCATCAAACCAAAGATTGGTCTCCATAATCTGG

B2_P1 (chrna3)_2	CCTCGTAAATCCTCATCAAAGGATGAACTCAACACCTCCAAAATC

B2_P1 (chrna3)_3	CCTCGTAAATCCTCATCAAAACCTGGAAATCTCCCACCGCATTGT

B2_P1 (chrna3)_4	CCTCGTAAATCCTCATCAAACTTGAAGATGGCCGGTGGGATCCAG

B2_P1 (chrna3)_5	CCTCGTAAATCCTCATCAAAACGAGCCAAACTTCATGGTGCAGTT

B2_P1 (chrna3)_6	CCTCGTAAATCCTCATCAAATCCCAGAAGTCCTTCAGGTTGATGG

B2_P1 (chrna3)_7	CCTCGTAAATCCTCATCAAACTCCTCACAGCAGTTGTATTTAATG

B2_P1 (chrna3)_8	CCTCGTAAATCCTCATCAAATGATCATGTTGATGGTGTAGAAGAG

B2_P2 (chrna3)_1	CTTATAGTCATTCCAAATGTGTCTCAAATCATCCAGTAAACCGCC

B2_P2 (chrna3)_2	GCTTCCAAATCTTGTTGGACGGCACAAATCATCCAGTAAACCGCC

B2_P2 (chrna3)_3	CGCAGAAGGGCTTTGGTTTTATCGTAAATCATCCAGTAAACCGCC

B2_P2 (chrna3)_4	GTAGGTGACGTCGATCTTGCAGGAGAAATCATCCAGTAAACCGCC

B2_P2 (chrna3)_5	GGTCGATCTTGGCCTTGTCGTAGGTAAATCATCCAGTAAACCGCC

B2_P2 (chrna3)_6	GCGTCGATGATCGTCCATTCTCCGCAAATCATCCAGTAAACCGCC

B2_P2 (chrna3)_7	CAGTGAATACGTGATGTCCGTGTAGAAATCATCCAGTAAACCGCC

B2_P2 (chrna3)_8	TCAGGAAGGAGATGAGAAGACAGGGAAATCATCCAGTAAACCGCC
